# TransportTools: a library for high-throughput analyses of internal voids in biomolecules and ligand transport through them

**DOI:** 10.1101/2021.06.01.445451

**Authors:** Jan Brezovsky, Aravind Selvaram Thirunavukarasu, Bartlomiej Surpeta, Carlos Eduardo Sequeiros-Borja, Nishita Mandal, Dheeraj Kumar Sarkar, Cedrix J. Dongmo Foumthuim, Nikhil Agrawal

**Affiliations:** Laboratory of Biomolecular Interactions and Transport, Department of Gene Expression, Institute of Molecular Biology and Biotechnology, Faculty of Biology, Adam Mickiewicz University, 61-614 Poznan, Poland; International Institute of Molecular and Cell Biology in Warsaw, 02-109 Warsaw, Poland; Latvian Institute of Organic Synthesis, LV-1006 Riga, Latvia

## Abstract

Information regarding pathways through voids in biomolecules and their roles in ligand transport is critical to our understanding of the function of many biomolecules. Recently, the advent of high-throughput molecular dynamics simulations has enabled the study of these pathways, and of rare transport events. However, the scale and intricacy of the data produced requires dedicated tools in order to conduct analyses efficiently and without excessive demand on users. To fill this gap, we developed the TransportTools, which allows the investigation of pathways and their utilization across large, simulated datasets. TransportTools also facilitates the development of custom-made analyses. TransportTools is implemented in Python3 and distributed as pip and conda packages. The source code is available at https://github.com/labbit-eu/transport_tools.

## 1. Introduction

At any moment, living systems contain thousands of small organic molecules that need to arrive at their sites of action to exert their function. The transport of these molecules around the cell (and beyond) is governed primarily by channels and tunnels (henceforth referred to as ‘pathways’) formed from the internal voids of biomolecules.^1^ These pathways enable the transport of ions and small molecules between different regions, connecting inner cavities with a surface, two different cavities with each other, or different cellular environments via transmembrane proteins. Operating as such, the investigation of these pathways is critical to drug discovery^2^ and protein engineering initiatives.^3^ Since pathways are often equipped with dynamic gates,^4^ they are mostly transient and challenging to study.

One of the most common approaches used to characterize these rare events of ligand transmission via transiently open pathways is to run molecular dynamics (MD) simulations,^5^ analyzing the pathway dynamics using tools like CAVER^6^ or tracking ligand migration through the biomolecules with AQUA-DUCT;^7^ see Supplementary File 1 for an overview of the state-of-the-art tools to study ligand transport pathways. The intensive development seen in computing hardware and sampling algorithms over recent years has led to considerable growth in the size and complexity of datasets typically generated for a single protein system. It is not uncommon for such datasets to consist of thousands simulations. Such high-throughput approaches, however, impose a substantial burden on researchers in establishing the identity of the pathways observed across all simulations, determining which pathways are used by particular ligands, and developing means of specific quantitative analyses. To this end, we present TransportTools: a library designed to alleviate these difficulties by providing easy, efficient access to comprehensive details on transport processes – even for large-scale simulation sets – and offering an environment for the development of novel analyses and tools.

## 2. Features

TransportTools is available as a Python3 module distributed under the GNU General Public License v3.0, and available via pip and conda managers as the *transport_tools* package. In its standard workflow (Fig. 1), TransportTools utilizes outputs from CAVER and AQUA-DUCT analyses of MD simulation, integrating their complementary insights to investigate transport pathways and corresponding ligand migration events in soluble and membrane-embedded proteins. To achieve efficiency in such a high-throughput regimen, raw data on pathway ensembles and ligand-transport events is first coarse-grained, and positioned on a spherical grid. Next, TransportTools identifies relationships between pathway ensembles from individual simulations and joins them into superclusters, to which ligand-transport events are then assigned (see and Supplementary File 2 for method details). Critical analysis parameters can be controlled via a configuration file. These parameters are thoroughly explained in the user guide, which also includes a detailed walk-through tutorial (Supplementary File 3). Aside from the ready-made workflow, the library offers many classes to process, manipulate, and analyze pathways and events, simplifying the production of custom-made analyses and, hopefully, stimulating further development of new packages (Supplementary File 4).

**Fig. 1.**
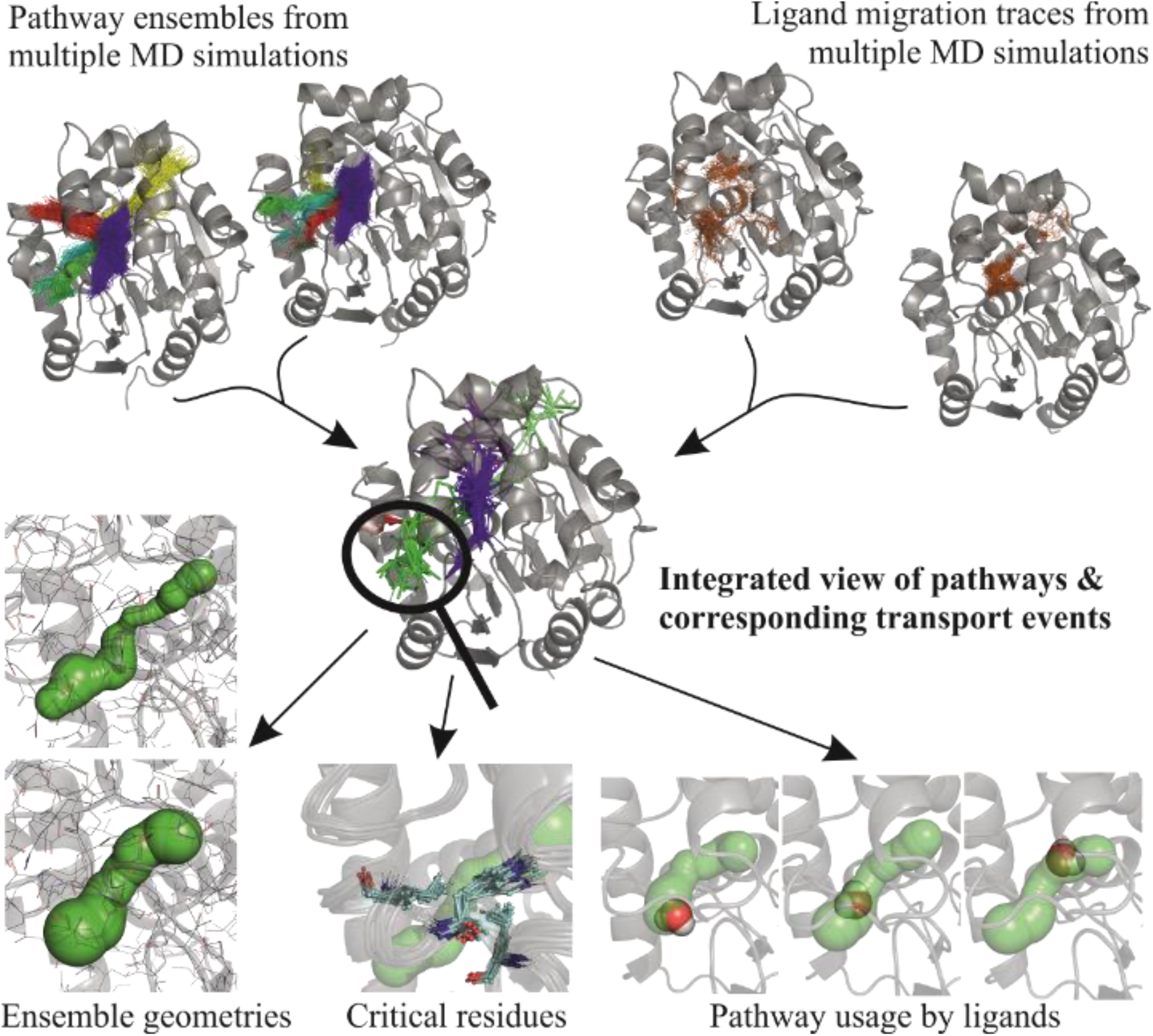
Schematic of a standard TransportTools analysis workflow.

### Outputs

The main results generated by TransportTools are presented as a set of tables stored in text files. These contain data on the composition of pathway superclusters, on their geometrical properties and utilization by transport events, and on critical protein residues. Using generated scripts, the spatial representation of superclusters and assigned events can be visualized in PyMOL.^8^ All results can be refined using various filters and split by individual simulation or by user-defined groups to facilitate their convenient comparison.

### Performance and limitations

The performance of TransportTools was analyzed on three datasets of 50 simulations (each sampling 100 ns and consisting of 10,000 frames) of up to 500 residue-long enzymes with different accessibilities of their active sites, resulting in the detection of up to 5,000,000 transport pathways and 50,000 water-transport events, which were processed within 2-21 hours on a standard workstation (Supplementary File 5). TransportTools inherits the limitations of the CAVER and AQUA-DUCT packages; their descriptions of pathway geometries and the definitions of their clusters (see Section 2.2 of Supplementary File 3 for best practice guidelines). When MD trajectories are utilized directly, usage is restricted to file formats supported by either *MDtraj* or *pytraj* packages.^9,10^

### Use cases

To illustrate the applicability of TransportTools, we applied it to the analysis of three representative examples of biological problems connected with ligand transport using an established model system – enzymes DhaA and LinB from the haloalkane dehalogenase family.^11,12^ First, we analyzed 10 simulations of DhaA in an effort to discover rare transient tunnels and their usage by water molecules (Supplementary File 6). Next, we derived an understanding of the effect of mutations on the system by contrasting simulations of LinB wild-type, LinB32 mutant with a closed primary tunnel, and LinB86 mutant with a *de novo* created tunnel (Supplementary File 7). Finally, we studied the substrate molecule selectivity of the pathways leading to the active site of LinB86 in almost 600 simulations (Supplementary File 8).

## 3. Conclusions

The TransportTools library provides users with access to (i) efficient analyses of transport pathways across extensive MD simulations, including those originating from massively parallel calculations or very long simulations; (ii) integrated data regarding transport pathways and their actual utilization by small molecules; and (iii) rigorous comparisons of transport processes under different settings, e.g. by contrasting transport in an original system against the same system perturbed by mutations, different solvents, or bound ligands.

## Acknowledgments

Computations were performed at the Poznan Supercomputing and Networking Center.

## References

(1) Kingsley, L. J.; Lill, M. A. Substrate Tunnels in Enzymes: Structure-Function Relationships and Computational Methodology. Proteins 2015, 83 (4), 599–611. https://doi.org/10.1002/prot.24772.

(2) Marques, S. M.; Daniel, L.; Buryska, T.; Prokop, Z.; Brezovsky, J.; Damborsky, J. Enzyme Tunnels and Gates As Relevant Targets in Drug Design. Med. Res. Rev. 2017, 37 (5), 1095–1139. https://doi.org/10.1002/med.21430.

(3) Kokkonen, P.; Bednar, D.; Pinto, G.; Prokop, Z.; Damborsky, J. Engineering Enzyme Access Tunnels. Biotechnol. Adv. 2019, 37 (6), 107386. https://doi.org/10.1016/j.biotechadv.2019.04.008.

(4) Gora, A.; Brezovsky, J.; Damborsky, J. Gates of Enzymes. Chem. Rev. 2013, 113 (8), 5871–5923. https://doi.org/10.1021/cr300384w.

(5) Decherchi, S.; Cavalli, A. Thermodynamics and Kinetics of Drug-Target Binding by Molecular Simulation. Chem. Rev. 2020, 120 (23), 12788–12833. https://doi.org/10.1021/acs.chemrev.0c00534.

(6) Jurcik, A.; Bednar, D.; Byska, J.; Marques, S. M.; Furmanova, K.; Daniel, L.; Kokkonen, P.; Brezovsky, J.; Strnad, O.; Stourac, J.; Pavelka, A.; Manak, M.; Damborsky, J.; Kozlikova, B. CAVER Analyst 2.0: Analysis and Visualization of Channels and Tunnels in Protein Structures and Molecular Dynamics Trajectories. Bioinformatics 2018, 34 (20), 3586–3588. https://doi.org/10.1093/bioinformatics/bty386.

(7) Magdziarz, T.; Mitusińska, K.; Bzówka, M.; Raczyńska, A.; Stańczak, A.; Banas, M.; Bagrowska, W.; Góra, A. AQUA-DUCT 1.0: Structural and Functional Analysis of Macromolecules from an Intramolecular Voids Perspective. Bioinformatics 2020, 36 (8), 2599–2601. https://doi.org/10.1093/bioinformatics/btz946.

(8) The PyMOL Molecular Graphics System, Version 2.0 Schrödinger, LLC.; 2017.

(9) McGibbon, R. T.; Beauchamp, K. A.; Harrigan, M. P.; Klein, C.; Swails, J. M.; Hernández, C. X.; Schwantes, C. R.; Wang, L.-P.; Lane, T. J.; Pande, V. S. MDTraj: A Modern Open Library for the Analysis of Molecular Dynamics Trajectories. Biophys. J. 2015, 109 (8), 1528–1532. https://doi.org/10.1016/j.bpj.2015.08.015.

(10) Roe, D. R.; Cheatham, T. E. PTRAJ and CPPTRAJ: Software for Processing and Analysis of Molecular Dynamics Trajectory Data. J. Chem. Theory Comput. 2013, 9 (7), 3084–3095. https://doi.org/10.1021/ct400341p.

(11) Pavlova, M.; Klvana, M.; Prokop, Z.; Chaloupkova, R.; Banas, P.; Otyepka, M.; Wade, R. C.; Tsuda, M.; Nagata, Y.; Damborsky, J. Redesigning Dehalogenase Access Tunnels as a Strategy for Degrading an Anthropogenic Substrate. Nat. Chem. Biol. 2009, 5 (10), 727–733. https://doi.org/10.1038/nchembio.205.

(12) Brezovsky, J.; Babkova, P.; Degtjarik, O.; Fortova, A.; Gora, A.; Iermak, I.; Rezacova, P.; Dvorak, P.; Smatanova, I. K.; Prokop, Z.; Chaloupkova, R.; Damborsky, J. Engineering a de Novo Transport Tunnel. ACS Catal. 2016, 6 (11), 7597–7610. https://doi.org/10.1021/acscatal.6b02081.

