## Supplementary File 1 for "TransportTools: a library for high-throughput analyses of internal voids in biomolecules and ligand transport through them"

### Supplementary File 1 – The overview of state-of-the-art tools to study ligand transport pathways

Investigation of the pathways in proteins enabling solvent or ligand transport constitutes a crucial source of information to understand molecular mechanisms underlying protein function as well as to guide drug discovery and protein engineering efforts (Marques *et al.*, 2017; Kokkonen *et al.*, 2019). Therefore we could witness the rapid development of such methods and tools in the last two decades (Mitusińska *et al.*, 2020; Brezovsky *et al.*, 2013). Taking into consideration the transient nature of biomolecular transport pathways, one of the principal requirements on these tools is their capability of analyzing structural ensembles, e.g., from molecular dynamics simulations, not only a single protein structure, which may have the transport pathways closed by one or more gates (McCammon, 2011; Gora *et al.*, 2013). Tools for detection of transport pathways in ensembles can be divided by their primary approach: (i) geometry-based analysis of voids within the structure of the biomolecule, and (ii) tracking of ligand movements penetrating the interior of the biomolecules. To contrast capabilities of both types of methods, the first ones are able to search for multiple pathways within a given probe radius but they do not provide information about the transport of solvent/ligands through them and frequently assume spherical geometries of the identified pathways (Brezovsky *et al.*, 2013; Mitusińska *et al.*, 2020). On the other hand, methods focused on tracking ligand movements can efficiently elucidate pathways explored by the molecule(s) of interest but they fail in exploring pathways, which were either not visited by ligand due to insufficient sampling or not used by a particular compound, e.g., due to physicochemical incompatibility, but could be used by other ligands or be engineered to do so (Brezovsky *et al.*, 2016; Gustafsson *et al.*, 2017; Kaushik *et al.*, 2018).

The first group in the vast majority is represented by tools searching for continuous empty spaces within the Voronoi diagram, which approximately represents free Van der Waals volume (Okabe *et al.*, 2000). Several tools were developed based on this approach including CAVER 3.0 (Chovancova *et al.*, 2012), Mole 1.0 (Petřek *et al.*, 2007), and MolAxis (Yaffe *et al.*, 2008) as the most popular representatives (Table 1). Methodologically, these tools mainly differ in the features such as the representation of the protein by the Voronoi diagram, the boundary between protein and solvent, way of treating multiple tunnels, cost functions, possibility to analyze trajectories/ensembles and form of outputs (Brezovsky *et al.*, 2013). Nevertheless, the most important limitations for their massive utilization for pathways detection in molecular ensembles are following: (i) Mole 1.0 exhibits non-negligible errors in tunnel geometries due to un-weighted Voronoi diagram being directly applied to atoms of different radii (Chovancova *et al.*, 2012), and (ii) MolAxis, although capable of accepting multiple structures as input, it is not clustering the tunnels from multiple snapshots, hence the analysis is provided per single structure, which is not suitable for massive analyzes of structural ensembles (Brezovsky *et al.*, 2013). The remaining tools are either web-server solutions such as BetaCavityWeb (Kim *et al.*, 2015), PoreWalker (Pellegrini-Calace *et al.*, 2009), ChExVis (Masood *et al.*, 2015), or cannot process multiple structures as Mole 2.0 (Sehnal *et al.*, 2013). Taking all aspects into consideration, CAVER 3.0 still represents the only choice for geometry-based analysis of transport pathway ensembles, and therefore its outputs are further employed by TransportTools. Importantly, all tools mentioned in this section are capable of detecting geometric pathways only, and do not provide any information about ligand transport events.

Alternatively, biomolecular pathways can be explored by tracking solvent/ligand migration through them (Mitusińska *et al.*, 2020). Several approaches were designed to study solvent/ligand behavior from molecular dynamics simulations including Streamline tracing (Vassiliev *et al.*, 2010), Visual Abstraction of Solvent Pathlines (Bidmon *et al.*, 2008), Watergate (Vad *et al.*, 2017), AQUA-DUCT (Magdziarz *et al.*, 2020), and *trj_cavity* (Paramo *et al.*, 2014). While the first three methods lack readily available implementations, both AQUA-DUCT and *trj_cavity* can be counted among well-established tools (Mitusińska *et al.*, 2020). *trj_cavity* is implemented within the GROMACS framework (Pronk *et al.*, 2013) featuring an efficient grid-based approach with a fast neighbor search to detect cavities and transmembrane tunnels. Additionally, *trj_cavity* can extract information about the time-dependent solvent capacity and volume of these cavities. However, this tool is not equipped with methods to investigate separate pathways in more complex networks of interior voids (Mitusińska *et al.*, 2020). On the other hand, AQUA-DUCT can track solvent/ligand molecules penetrating user-specified regions of interest in the protein from molecular dynamics simulation (Magdziarz *et al.*, 2017, 2020). It can analyze migration events of any kind of ligand including different solvents and ions. Its only but crucial requirement is for input trajectories to be saved with a small time-step to maintain the continuity of the traced ligand paths. AQUA-DUCT is available as an easy-to-use Python package, which makes it available to a broad community. Its outputs constitute optional inputs for TransportTools to enable coupling of ligand migration with geometries of used pathway ensembles.

**Table 1 Overview of implemented tools suitable for analyzes of transport pathways in ensembles of protein structures using geometry-based or ligand tracking approaches.**

| Tool | Principle | Limitations | Web-page | Reference |
|---|---|---|---|---|
| **Geometry-based methods** | | | | |
| CAVER 3.0 | Voronoi diagram | Lack of parallelization and unifying framework for outputs | https://caver.cz/ | (Chovancova *et al.*, 2012) |
| Mole 1.0 | Voronoi diagram | Sub-optimal pathway geometries | http://mole.chemi.muni.cz/ | (Petřek *et al.*, 2007) |
| MolAxis | Voronoi diagram | Lack of ensemble-averaged analysis | http://bioinfo3d.cs.tau.ac.il/MolAxis/ | (Yaffe *et al.*, 2008) |
| **Ligand-tracking methods** | | | | |
| AQUA-DUCT 1.0 | Tracking of molecular migration | Requires trajectories with small time-step | http://www.aquaduct.pl/ | (Magdziarz *et al.*, 2020) |
| *trj_cavity* | Grid-based analyzes of void and ligand presence | No facilities to investigate separate pathways in more complex networks of interior voids | https://sourceforge.net/projects/trjcavity/ | (Paramo *et al.*, 2014) |

#### REFERENCES


Bidmon,K. *et al.* (2008) Visual Abstractions of Solvent Pathlines near Protein Cavities. *Comput. Graph. Forum*, **27**, 935–942.

Brezovsky,J. *et al.* (2016) Engineering a de Novo Transport Tunnel. *ACS Catal.*, **6**, 7597–7610.

Brezovsky,J. *et al.* (2013) Software tools for identification, visualization and analysis of protein tunnels and channels. *Biotechnol. Adv.*, **31**, 38–49.

Chovancova,E. *et al.* (2012) CAVER 3.0: A Tool for the Analysis of Transport Pathways in Dynamic Protein Structures. *PLoS Comput. Biol.*, **8**, e1002708.

Gora,A. *et al.* (2013) Gates of Enzymes. *Chem. Rev.*, **113**, 5871–5923.

Gustafsson,C. *et al.* (2017) MD Simulations Reveal Complex Water Paths in Squalene–Hopene Cyclase: Tunnel-Obstructing Mutations Increase the Flow of Water in the Active Site. *ACS Omega*, **2**, 8495–8506.

Kaushik,S. *et al.* (2018) Impact of the access tunnel engineering on catalysis is strictly ligand-specific. *FEBS J.*, **285**, 1456–1476.

Kim,J.-K. *et al.* (2015) BetaCavityWeb: a webserver for molecular voids and channels. *Nucleic Acids Res.*, **43**, W413–W418.

Kokkonen,P. *et al.* (2019) Engineering enzyme access tunnels. *Biotechnol. Adv.*, **37**, 107386.

Magdziarz,T. *et al.* (2020) AQUA-DUCT 1.0: structural and functional analysis of macromolecules from an intramolecular voids perspective. *Bioinformatics*, **36**, 2599–2601.

Magdziarz,T. *et al.* (2017) AQUA-DUCT: a ligands tracking tool. *Bioinformatics*, **33**, 2045–2046.
Marques,S.M. *et al.* (2017) Enzyme Tunnels and Gates As Relevant Targets in Drug Design. *Med. Res. Rev.*, **37**, 1095–1139.
Masood,T.B. *et al.* (2015) CHEXVIS: a tool for molecular channel extraction and visualization. *BMC Bioinformatics*, **16**, 119.
McCammon,J.A. (2011) Gated Diffusion-controlled Reactions. *BMC Biophys.*, **4**, 4.
Mitusińska,K. *et al.* (2020) Applications of water molecules for analysis of macromolecule properties. *Comput. Struct. Biotechnol. J.*, **18**, 355–365.
Okabe,A. *et al.* (2000) Spatial Tessellations: Concepts and Applications of Voronoi Diagrams 2nd ed. Wiley.
Paramo,T. *et al.* (2014) Efficient Characterization of Protein Cavities within Molecular Simulation Trajectories: trj_cavity. *J. Chem. Theory Comput.*, **10**, 2151–2164.
Pellegrini-Calace,M. *et al.* (2009) PoreWalker: A Novel Tool for the Identification and Characterization of Channels in Transmembrane Proteins from Their Three-Dimensional Structure. *PLOS Comput. Biol.*, **5**, e1000440.
Petřek,M. *et al.* (2007) MOLE: A Voronoi Diagram-Based Explorer of Molecular Channels, Pores, and Tunnels. *Structure*, **15**, 1357–1363.
Pronk,S. *et al.* (2013) GROMACS 4.5: a high-throughput and highly parallel open source molecular simulation toolkit. *Bioinforma. Oxf. Engl.*, **29**, 845–854.
Sehnal,D. *et al.* (2013) MOLE 2.0: advanced approach for analysis of biomacromolecular channels. *J. Cheminformatics*, **5**, 39.
Vad,V. *et al.* (2017) Watergate: visual exploration of water trajectories in protein dynamics. In, *Proceedings of the Eurographics Workshop on Visual Computing for Biology and Medicine*, VCBM ’17. Eurographics Association, Goslar, DEU, pp. 33–42.
Vassiliev,S. *et al.* (2010) Tracking the flow of water through photosystem II using molecular dynamics and streamline tracing. *Biochemistry*, **49**, 1873–1881.
Yaffe,E. *et al.* (2008) MolAxis: Efficient and accurate identification of channels in macromolecules. *Proteins Struct. Funct. Bioinforma.*, **73**, 72–86.
