## Supplementary File 2 for "TransportTools: a library for high-throughput analyses of internal voids in biomolecules and ligand transport through them"

### Supplementary File 2 – The overview of operations performed within the standard TransportTools workflow and sensitivity of analyses to its main parameters

### 1. OPERATIONS WITHIN THE STANDARD TRANSPORTTOOOLS WORKFLOW

**TransportTools workflow is executed in up to ten consecutive stages:**

1. Preparatory stage including defining transformations needed for unified analyses
2. Processing of input datasets of tunnel networks
3. Layering tunnel clusters to get their simplified representation
4. Computing distances among the layered clusters
5. Clustering the layered clusters into superclusters and creating initial outputs
6. Filtering superclusters and creating filtered outputs
7. Processing datasets of transport events of ligands
8. Layering transport events to get their simplified representation
9. Assigning transport events to tunnel networks in superclusters and creating initial outputs
10. Filtering superclusters with events and creating filtered outputs with events

Note that stages 7-10 require data on transport events from AQUA-DUCT and are optional.

In stage 1, the biomolecular structures representing individual systems with tunnel networks and optionally also transport events are aligned on the reference PDB file. The whole system is translated to have its global origin in coordinates of the mean starting point of all tunnel networks, producing the respective transformation matrices. In stage 2, the tunnel networks are parsed from the CAVER results, transformed into a uniform coordinate system, and stored to enable further processing (Fig. 1A). In stage 3, the points forming the tunnel clusters processed in the previous stage are divided into layers of a user-defined thickness (Fig. 1B); see Section 2 for details on impacts of the thickness setting on overall results. Next, all points in the same layer are coarse-grained into representative nodes while filtering the outliers (Fig. 1C), using a density-based clustering based on hierarchical density estimates on their Cartesian coordinates as implemented in *hdbscan* package (Campello *et al.*, 2013). Finally, coarse-grained paths connecting those nodes from the starting point to all nodes containing the end-points of the original tunnels in a manner equivalent to the original tunnels are identified (Fig. 1D). A set of these coarse-grained paths can be used to efficiently represent the geometry of all tunnel clusters within the analyzed MD simulation (Fig. 1E).

In stage 4, the average distances between the surfaces of the closest pairs of coarse-grained nodes from all layered paths representing the original tunnel clusters are computed across all analyzed MD simulations (Fig. 2A, B). In stage 5, the coarse-grained tunnel clusters with the average closest surface distances within a user-defined distance cutoff are clustered into superclusters using hierarchical agglomerative clustering algorithms implemented in *fastcluster* package (Müllner, 2013) (Fig. 2C); see Section 2 for details on the sensitivity of formed superclusters to the main clustering parameters. Subsequently, by processing complete data on the tunnel networks belonging to each supercluster, overall statistics on the superclusters, their cumulative tunnel profiles, bottleneck residues, and details about the membership of tunnel clusters in superclusters are generated together with the respective visualization files (Fig. 2D). Finally, the supercluster profiles are filtered based on the tunnel geometry in stage 6. Then, the overall statistics of the filtered superclusters, their cumulative tunnel profiles, bottleneck residues, and details about the membership of superclusters are generated together with the respective visualization files.

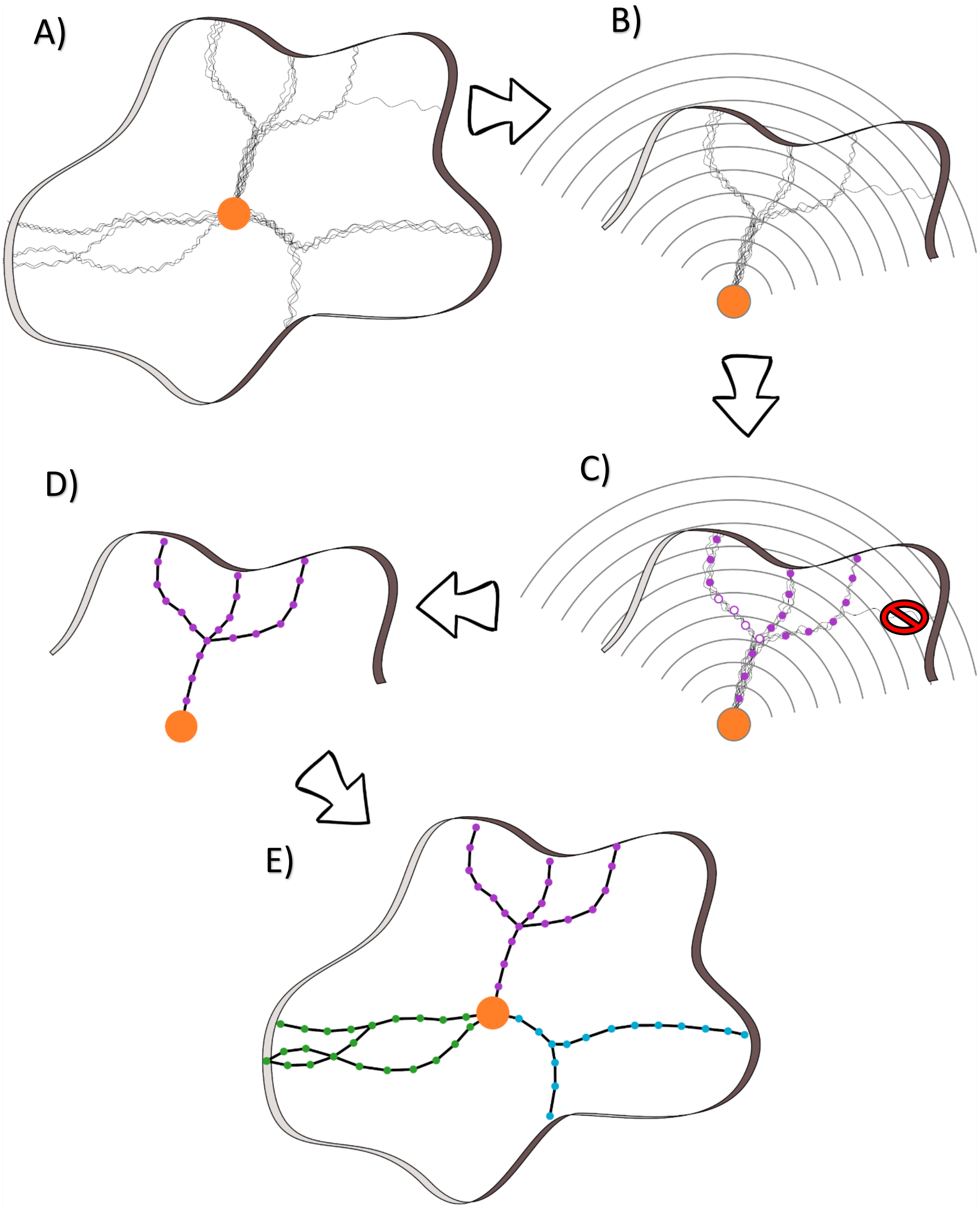


**Fig. 1. A schema of tunnel clusters transformation into their simplified representations. A**) Multiple tunnel clusters that are found in a single MD simulation formed by tunnels leading from the starting point (orange dot). **B**) Division of tunnels forming the clusters into layers of a given thickness. **C**) After layering, nodes are defined for each layer by clustering the original tunnel spheres assigned to this layer. Please note that there can be more than one node in the same layer (empty circles). Moreover, some very rare tunnels could be omitted as outliers. **D**) The nodes are joined, and the representation of the geometry of the whole tunnel cluster is obtained. **E**) Finally, the simplified representations of all tunnel clusters as sets of coarse-grained paths are obtained.

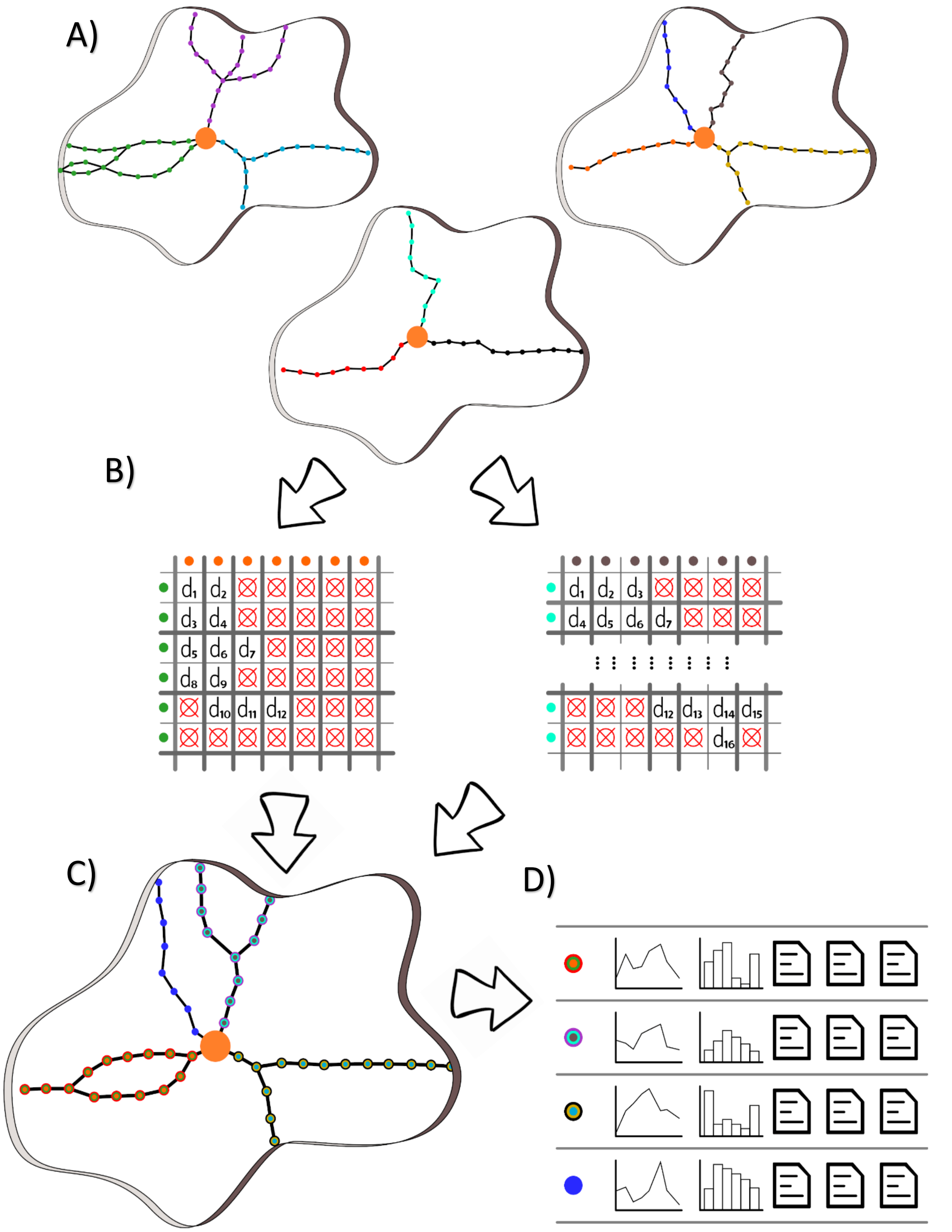


**Fig. 2. Workflow for superclusters generation.** **A**) Simplified representations of tunnel cluster geometry from different MD simulations. **B**) For each layered path representing a tunnel cluster, the distances to all paths in other clusters are calculated by averaging the shortest distances between nodes forming those paths located in the adjacent layers (distances above a user-defined cutoff are denoted as ⊗). **C**) Tunnel clusters that are within the cutoff are then merged into superclusters. **D**) Finally, each supercluster is analyzed, and outputs are generated for each one.

In stage 7, networks of transport events are parsed from the AQUA-DUCT results, transformed to a uniform coordinate system, and stored to enable their further processing. Only events that get closer than a user-defined distance to the starting point of tunnel networks are considered at this stage (Fig. 3A). In stage 8, the points forming traces of transport events processed in the previous stage are divided into layers. Next, all points in the same layer are clustered into representative nodes while filtering the outliers similarly to the coarse-graining of tunnel clusters in stage 3 (Fig. 3B). Finally, the layered paths connecting those nodes in a manner similar to the original event's trace are identified to represent the geometry of the event's trace efficiently (Fig. 3C). In stage 9, layered transport events are assigned to superclusters when the fraction of their traces buried inside the supercluster volumes are larger than a user-defined cutoff (Fig. 4). In stage 10, the supercluster profiles are filtered using additional filters based on the transport events assigned to them. Finally, the overall statistics of the superclusters, their cumulative tunnel profiles, bottleneck residues, and details about the membership of tunnel clusters in superclusters are generated together with the respective visualization files.

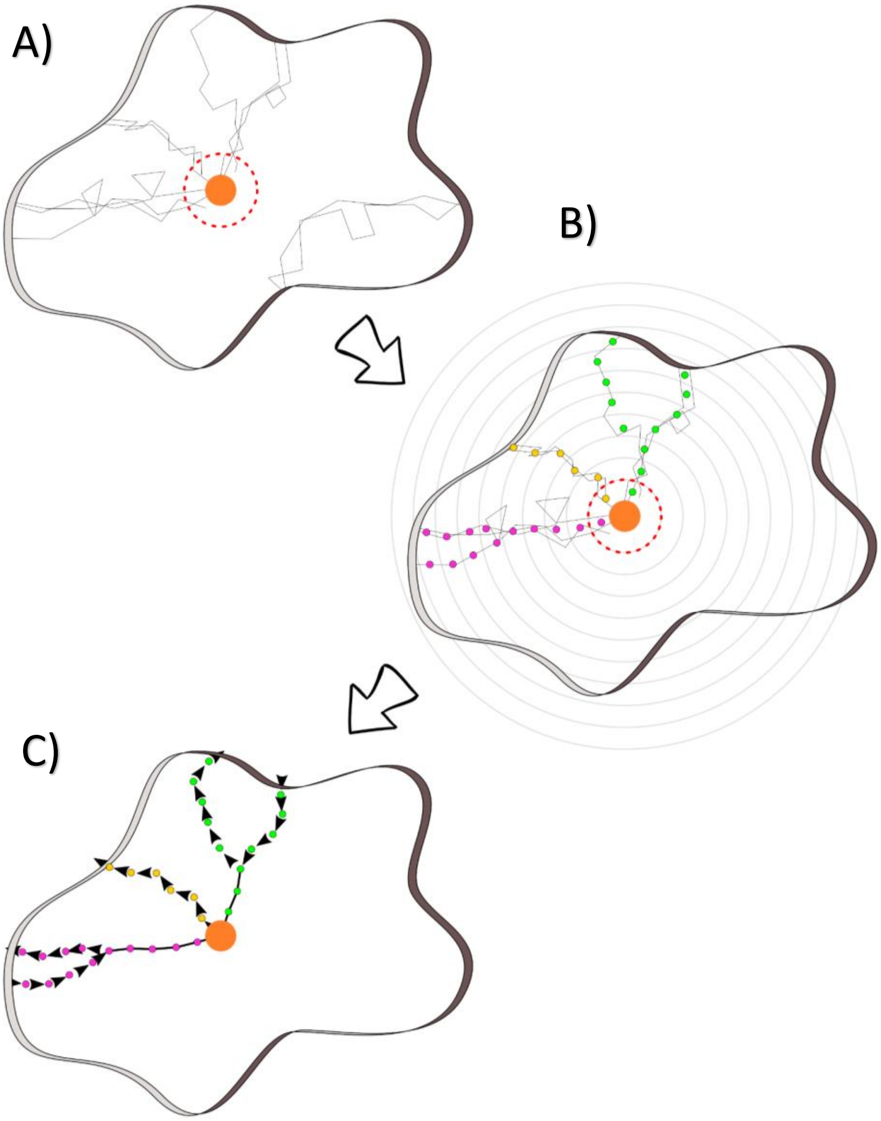


**Fig. 3. A schema of transformation of transport events into their simplified representations. A)** Transport events are considered for analysis only if they reach the internal region (dotted red circle). **B)** The layering of the clusters follows the same procedure as for tunnel clusters (see Fig. 1). **C)** Layered transport events are separated into those representing ligand entry and release, shown here with arrows.

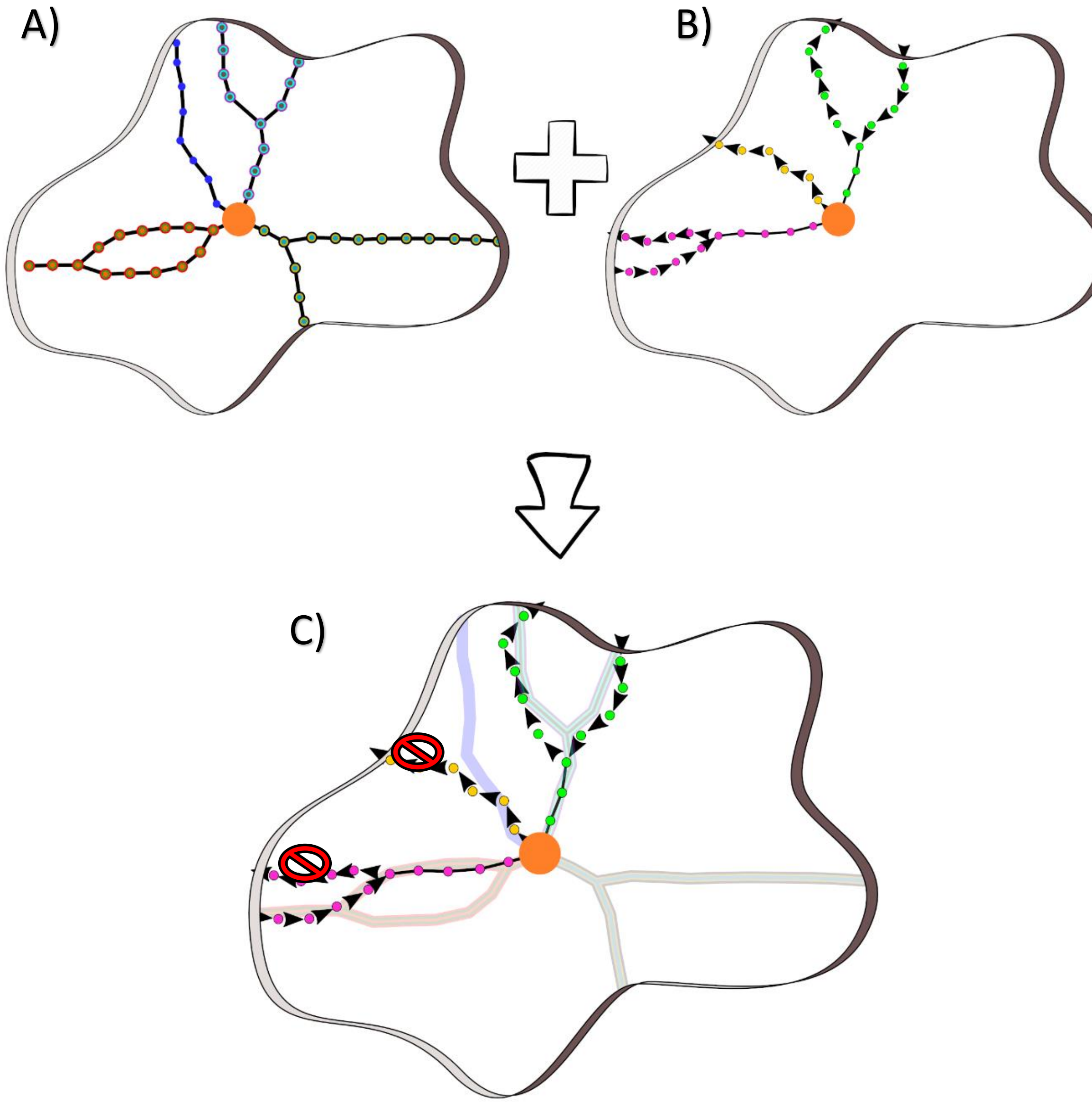


**Fig. 4. Workflow for assignment of transport events into superclusters. A)** Superclusters and **B)** information from layered transport events are joined to assign transport events to superclusters. **C)** A transport event (colored discs) is assigned to a supercluster only if a fraction of their trajectory is buried in the volume formed by a given supercluster (shaded regions) is larger than a user-defined cutoff.

#### 2. SENSITIVITY OF ANALYSES TO LAYER THICKNESS AND CLUSTERING PARAMETERS

Three simulations of the haloalkane dehalogenase from Use Case I (Supplementary file 6; subfolders e1s1_conf1, e1s2_conf1, and e1s3_conf1 in the data folder) were used as an evaluation dataset for analysis of the role of layer thickness and clustering parameters on the quality of results produced by TransportTools. Tunnel analyses of these simulations by CAVER revealed 93 tunnel clusters, among them the standard tunnels (Fig. 5), labeled as p1, p2, and p3, which are well-established for this protein family (Klvana *et al.*, 2009; Kokkonen *et al.*, 2018; Brezovsky *et al.*, 2016; Marques *et al.*, 2017). Please note that the coloring of tunnel clusters in this section reflects their throughput starting from blue, green, red, cyan, yellow, and pink for the most relevant ones, following the nomenclature established by the CAVER package (Pavelka *et al.*, 2016; Chovancova *et al.*, 2012).

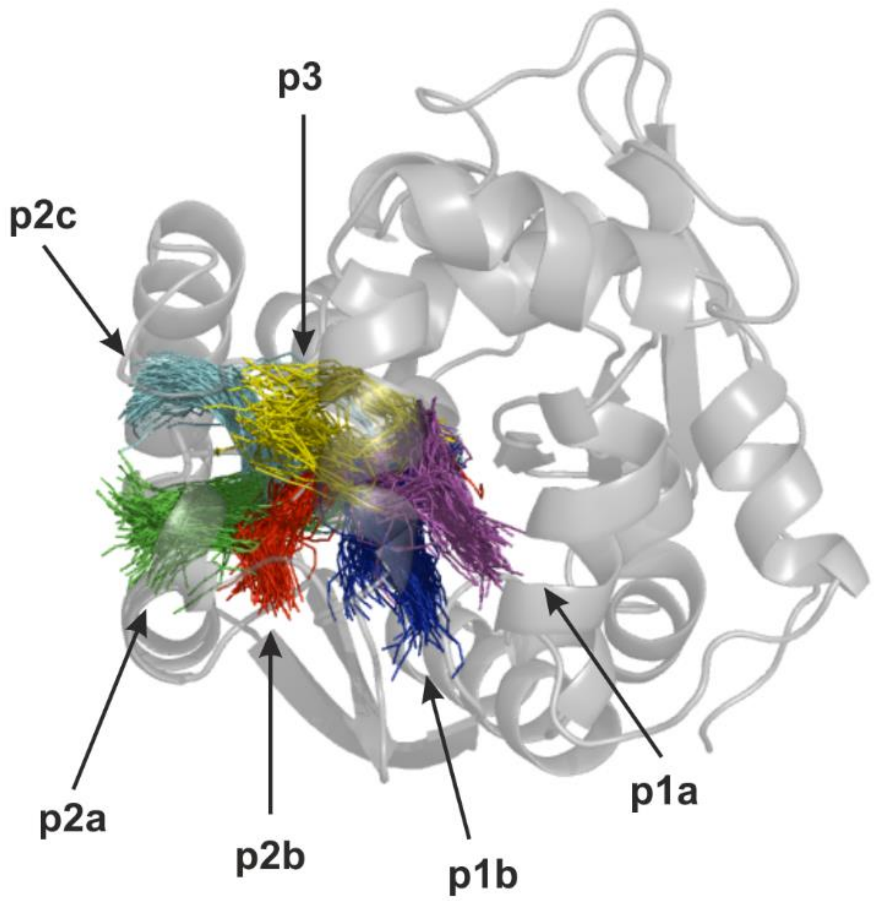


**Fig. 5. The three standard tunnel clusters and their branches recognized in the haloalkane dehalogenase family.**

Using this dataset as inputs to TransportTools, we varied the thickness of layers used during the coarse-graining of the tunnel clusters. Since the input data on tunnels used for this evaluation were obtained with the CAVER package using the default step of 0.5 Å between successive points forming a tunnel (defined by CAVER parameter *profile_tunnel_sampling_step*), we have been able to use the minimum 0.9 Å as the finest layer thickness for TransportTools that still contained enough tunnel-forming points in individual layers to compute a coarse-grained representation of at least some tunnel clusters. Conversely, the maximal value of 15.0 Å corresponds with the length of the majority of tunnels detected in this evaluation dataset, which means that during coarse-graining of tunnel clusters most tunnel-forming points will belong to a single layer and hence be processed together. When comparing the tunnel superclusters identified by TransportTools representing equivalent tunnels across the whole evaluation dataset at a given thickness, we focused on the most relevant ones, i.e., those with frequencies over 5 % (Table 1).

For the lowest thickness of 0.9 Å, only nine tunnel clusters out of 93 processed ones were layered successfully due to too few tunnel points present in individual layers, resulting in a very sparse and inaccurate description of the tunnel network. All other tested values of layer thickness produced an appropriate view of the standard tunnels with respect to the level of coarse-graining (Fig.5). Using thickness in a conservative range between 1.0 and 5.0 Å, branches of p1 tunnel were consistently identified in 82% of analyzed frames as supercluster 1. The p2a and p2b branches were detected as supercluster 2 in 25% of frames, whereas a more distinct p2c tunnel was separated to supercluster 3 in about 7% of frames. Finally, the p3 tunnel was consistently identified in about 6% of frames as supercluster 4. Also, the geometries of all tunnels were maintained across this range of layer thickness. At a more aggressive layer thickness above 10 Å, p3 tunnel was no longer detected among tunnels with more than 5% frequency since it was divided into smaller branches. Similarly, p2a was separated from the supercluster 2, which was formed by both p2a and p2b branches at smaller thicknesses, to create a smaller supercluster. Finally, the p1 tunnel also became divided into its branches at 15 Å layering thickness.

In general, when increasing the layering thickness, the coarse-grained clusters become represented by fewer coarse-grained nodes, carrying less details about the original geometry. However, this reduction is not directly proportional to the thickness used since multiple distinct coarse-grained nodes can be recognized among the original tunnel-forming points within any single layer by the applied density-based clustering. On the other hand, the complexity of the coarse-graining process increases with the thickness as illustrated by the respective coarse-graining runtimes in Table 1. The positive side of using larger thickness values is that with fewer nodes forming the coarse-grained clusters, much less time is required to calculate inter-tunnel distances between them (see Runtime distance calculation in Table 1). As such, a higher layer thickness could be employed to efficiently analyze larger biomolecular systems without overly compromising the quality of results.

**Table 1. The role of layer thickness on the coarse-grained tunnel networks.**

| **Layer thickness [Å]** | **0.9** | | | **1.0** | | |
|---|---|---|---|---|---|---|
| **Coarse-grained tunnel network** | 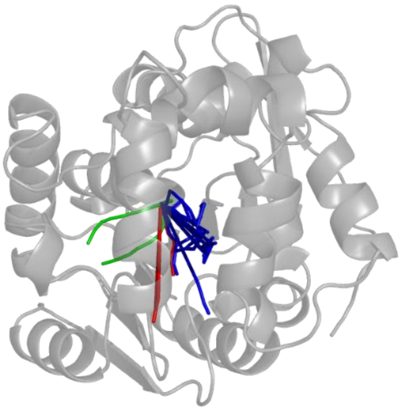 | | | 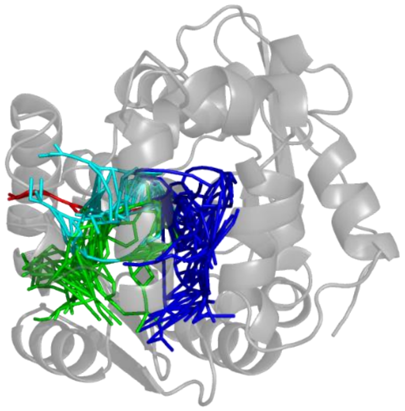 | | |
| **Supercluster /Average properties** | **#Frames** | **Bottleneck radius [Å]** | **Length [Å]** | **#Frames** | **Bottleneck radius [Å]** | **Length [Å]** |
| **1** | 8048 | 1.1±0.2 | 14±2 | 8215 | 1.1±0.2 | 14±2 |
| **2** | 1035 | 1.1±0.1 | 15±2 | 2586 | 1.0±0.1 | 16±2 |
| **3** | 773 | 1.1±0.1 | 15±2 | 695 | 1.1±0.1 | 17±1 |
| **4** | | not detected | | 589 | 0.96±0.05 | 14±2 |
| **Runtimes [s]** | | | | | | |
| **coarse-graining** | | 50* | | | 64 | |
| **distance calculation** | | <1* | | | 30 | |
| **Layer thickness [Å]** | **1.5** | | | **2.0** | | |
| **Coarse-grained tunnel network** | 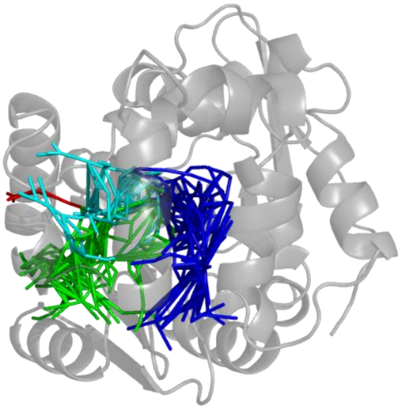 | | | 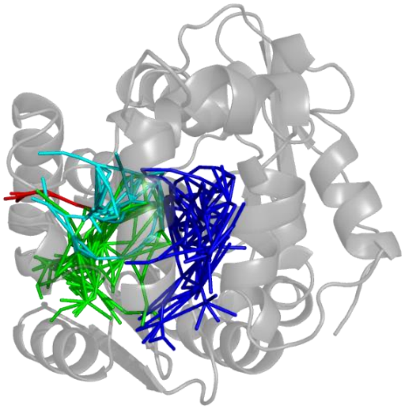 | | |
| **Supercluster /Average properties** | **#Frames** | **Bottleneck radius [Å]** | **Length [Å]** | **#Frames** | **Bottleneck radius [Å]** | **Length [Å]** |
| **1** | 8215 | 1.1±0.2 | 14±2 | 8215 | 1.1±0.2 | 14±2 |
| **2** | 2586 | 1.0±0.1 | 16±2 | 2585 | 1.0±0.1 | 16±2 |
| **3** | 695 | 1.1±0.1 | 17±1 | 695 | 1.1±0.1 | 17±1 |
| **4** | 589 | 0.96±0.05 | 14±2 | 588 | 0.96±0.05 | 14±2 |
| **Runtimes [s]** | | | | | | |
| **coarse-graining** | | 58 | | | 58 | |
| **distance calculation** | | 17 | | | 15 | |

* - only nine tunnel clusters were processed in this case

Table 1. continued

| Layer thickness [Å] | 3.0 | 5.0 |
|---|---|---|
| Coarse-grained tunnel network | 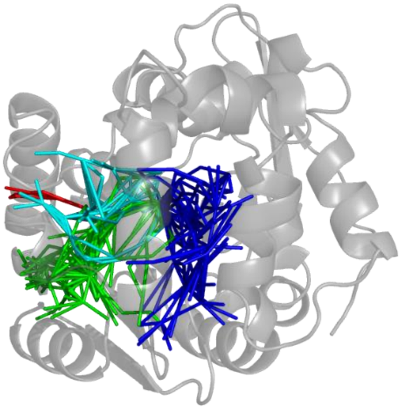 | 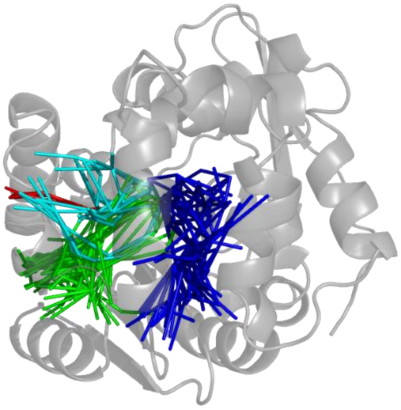 |

| Supercluster /Average properties | #Frames | Bottleneck radius [Å] | Length [Å] | #Frames | Bottleneck radius [Å] | Length [Å] |
|---|---|---|---|---|---|---|
| 1 | 8215 | 1.1±0.2 | 14±2 | 8215 | 1.1±0.2 | 14±2 |
| 2 | 2586 | 1.0±0.1 | 16±2 | 2585 | 1.0±0.1 | 16±2 |
| 3 | 695 | 1.1±0.1 | 17±1 | 695 | 1.1±0.1 | 17±1 |
| 4 | 589 | 0.96±0.05 | 14±2 | 589 | 0.96±0.05 | 14±2 |
| Runtimes [s] | | | | | | |
| coarse-graining | | 60 | | | 67 | |
| distance calculation | | 5 | | | 3 | |

| Layer thickness [Å] | 10.0 | 15.0 |
|---|---|---|
| Coarse-grained tunnel network | 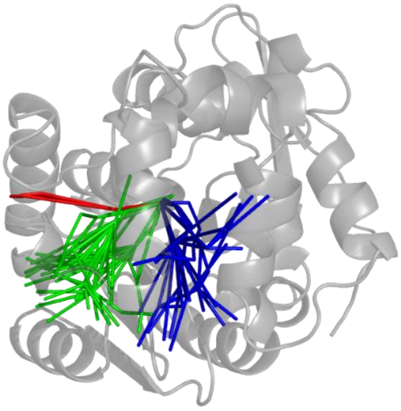 | 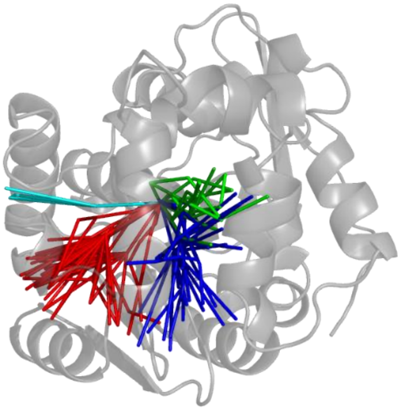 |

| Supercluster /Average properties | #Frames | Bottleneck radius [Å] | Length [Å] | #Frames | Bottleneck radius [Å] | Length [Å] |
|---|---|---|---|---|---|---|
| 1 | 8213 | 1.1±0.2 | 14±2 | 5975 | 1.1±0.2 | 14±2 |
| 2 | 2144 | 1.0±0.1 | 16±2 | 2698 | 1.1±0.2 | 14±2 |
| 3 | 695 | 1.1±0.1 | 17±1 | 2585 | 1.0±0.1 | 16±2 |
| 4 | | not detected | | 695 | 1.1±0.1 | 17±1 |
| Runtimes [s] | | | | | | |
| coarse-graining | | 101 | | | 142 | |
| distance calculation | | 2 | | | 1 | |

To illustrate the impact of the selection of clustering parameters on the superclusters formed by TransportTools, we systematically varied both the used linkage method available from the *fastcluster* package (single, average, centroid, median, complete, and Ward), as well as the clustering cutoff on the inter-cluster distances (0-15 Å) on the evaluation dataset. At 0 Å cutoff, only entirely overlapping coarse-grained tunnel clusters from 93 input ones were merged into 87 superclusters irrespective of the linkage used (Tables 2-7). At this cutoff, many alternatives of each known tunnel (Fig. 5) could frequently be identified in the respective regions (Tables 2-7). With the increasing cutoff, the examined linkage methods exhibited different capabilities to separate the distinct tunnels and recognize their branches.

Due to its aggressive tendency to chain clusters, the single-linkage method merged all known tunnels into one supercluster already at 1 Å cutoff and quickly progressed in aggregating all tunnels into a single supercluster at 3 Å cutoff (Table 2). Such behavior renders this method invalid for the most of analyses and as such TransportTools engine generally suggest against using it. However, due to its efficiency, the single-linkage method could still be beneficial in cases where only single tunnel cluster is presumed to exists.

The next group of linkage methods that often showed almost indistinguishable or even identical results on the evaluation dataset was centroid, median and average (Tables 3-5). With these methods, all known tunnels and their branches were clearly separated into different superclusters at 1 Å cutoff, with a minor difference in the formation of alternative branches. At 2 Å cutoff, all three methods merged branches of p1 tunnel into one supercluster, and for centroid and median linkages, their slight tendency towards chaining resulted in mixing of p3 tunnel with some of the p2 branches. All p3 and p2 tunnel variants were joined into one supercluster at 3 Å cutoff and later joined by p1 tunnel at 4 Å cutoff (centroid) or 5 Å cutoff (median and average). Finally, all tunnels were aggregated into a single supercluster at 10 Å cutoff, irrespective of the linkage used. In general, centroid-, median-, and average-linkage methods provide enough resolution to comfortably set up the user-required separation of tunnels, with average-linkage performing a little bit better than the other two methods Moreover, these methods enable detection of asymmetrically shaped superclusters unlike complete and Ward linkages.

The last group of linkage methods (complete and Ward) provided the most refined separations that clearly distinguished all known tunnels including their branches even at 3 and 4 Å cutoffs, respectively (Tables 6-7). As the cutoff increases, these methods merged p1 branches and then p2 and p3 tunnels into superclusters. While complete linkage joins all known tunnels to a single supercluster at 10 Å cutoff and all tunnels 15 Å cutoff, the Ward method still maintained the separation of p1 versus p2+p3 tunnels at 15 Å cutoff. The downside of these two methods is their strict differentiation of alternative variants of the same tunnels providing a possibly excessive level of details. On the evaluation dataset complete and Ward methods generated almost 50% more superclusters in contrast with centroid, median and average linkage methods at the same cutoff (Tables 3-7), hence complete and Ward methods are best suited when the details of tunnel branching is important.

**Table 2. Superclusters formed at different clustering cutoffs using the Single-linkage.**

| **Cutoff [Å]** | **0** | **1** |
|---|---|---|
| **Coarse-grained tunnel network** | 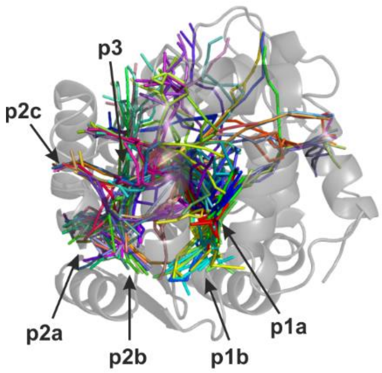  | 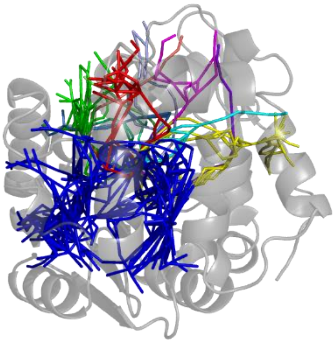 |
| **# superclusters** | 87 | 12 |
| **Cutoff [Å]** | **2** | **3** |
| **Coarse-grained tunnel network** | 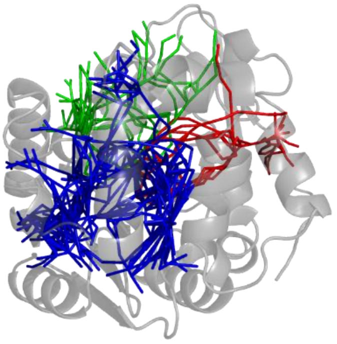 | 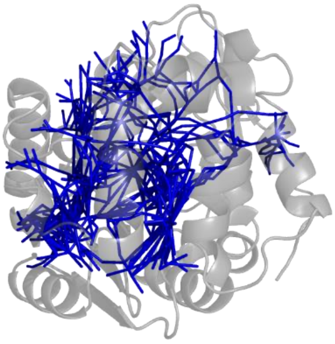 |
| **# superclusters** | 3 | 1 |
| **Cutoff [Å]** | **4** | **5** |
| **Coarse-grained tunnel network** | 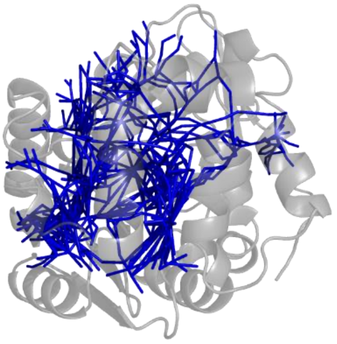 | 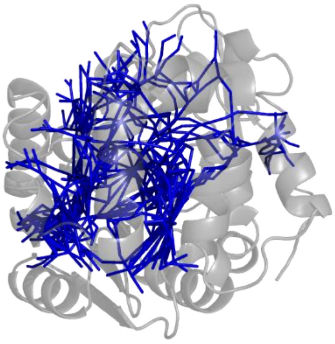 |
| **# superclusters** | 1 | 1 |
| **Cutoff [Å]** | **10** | **15** |
| **Coarse-grained tunnel network** | 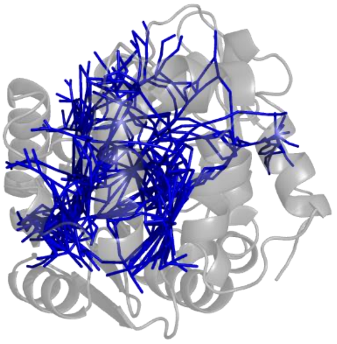 | 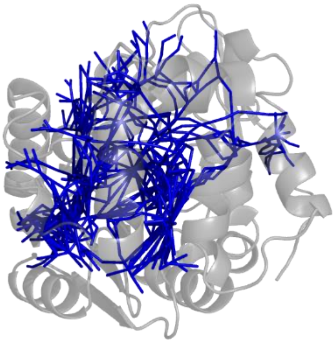 |
| **# superclusters** | 1 | 1 |

**Table 3. Superclusters formed at different clustering cutoffs using the Centroid-linkage.**

| **Cutoff [Å]** | **0** | **1** |
|---|---|---|
| **Coarse-grained tunnel network** | 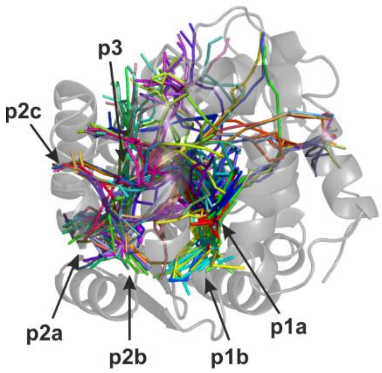  | 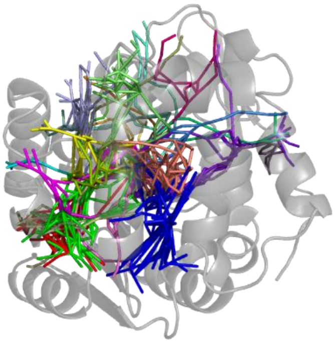 |
| **# superclusters** | 87 | 24 |
| **Cutoff [Å]** | **2** | **3** |
| **Coarse-grained tunnel network** | 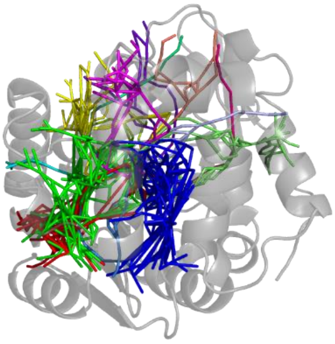 | 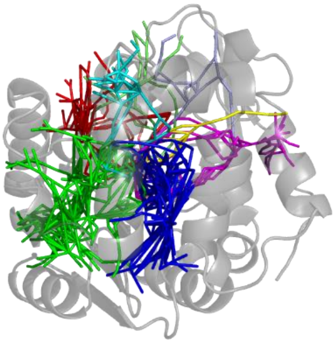 |
| **# superclusters** | 14 | 9 |
| **Cutoff [Å]** | **4** | **5** |
| **Coarse-grained tunnel network** | 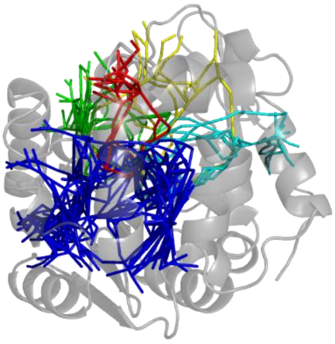 | 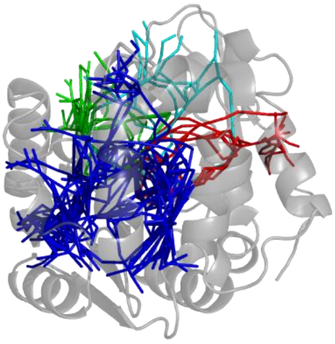 |
| **# superclusters** | 5 | 4 |
| **Cutoff [Å]** | **10** | **15** |
| **Coarse-grained tunnel network** | 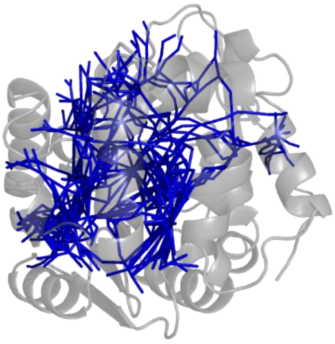 | 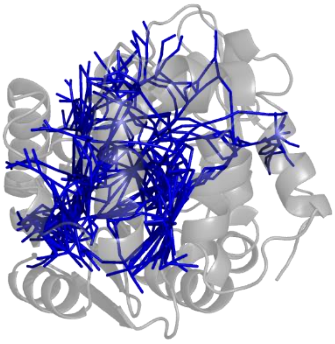 |
| **# superclusters** | 1 | 1 |

**Table 4. Superclusters formed at different clustering cutoffs using the Median-linkage.**

<table>
<tr><th>Cutoff [Å]</th><th>0</th><th>1</th></tr>
<tr><td>Coarse-grained tunnel network</td><td>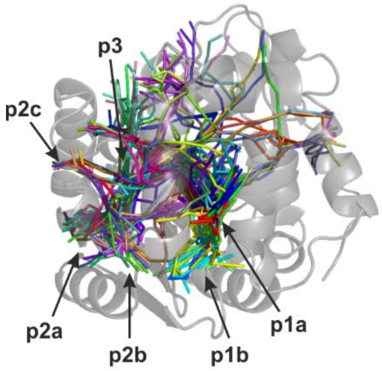
</td><td>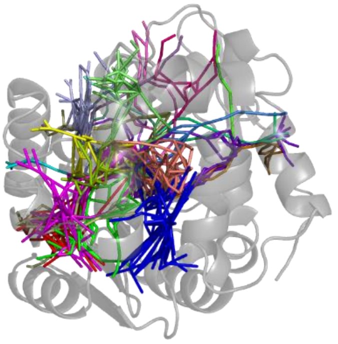</td></tr>
<tr><td># superclusters</td><td>87</td><td>23</td></tr>
<tr><th>Cutoff [Å]</th><th>2</th><th>3</th></tr>
<tr><td>Coarse-grained tunnel network</td><td>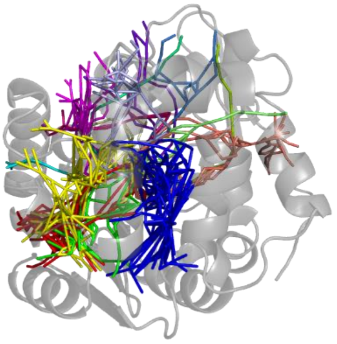</td><td>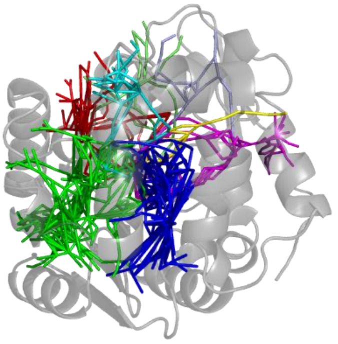</td></tr>
<tr><td># superclusters</td><td>15</td><td>9</td></tr>
<tr><th>Cutoff [Å]</th><th>4</th><th>5</th></tr>
<tr><td>Coarse-grained tunnel network</td><td>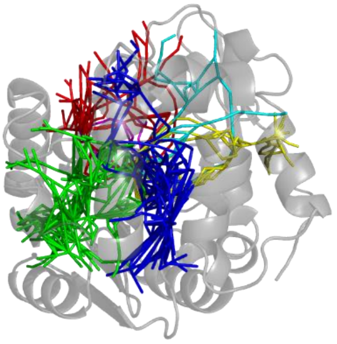</td><td>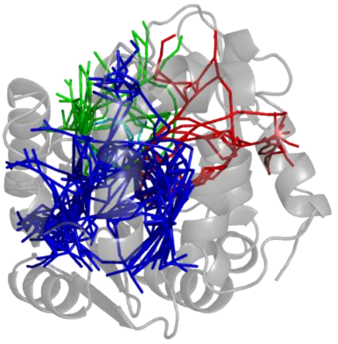</td></tr>
<tr><td># superclusters</td><td>6</td><td>4</td></tr>
<tr><th>Cutoff [Å]</th><th>10</th><th>15</th></tr>
<tr><td>Coarse-grained tunnel network</td><td>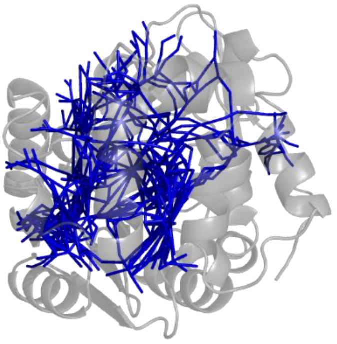</td><td>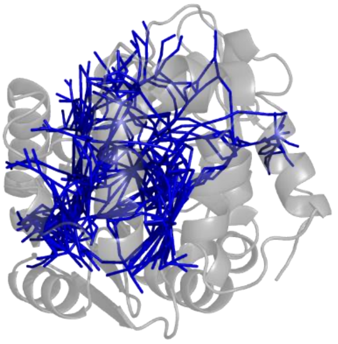</td></tr>
<tr><td># superclusters</td><td>1</td><td>1</td></tr>
</table>

**Table 5. Superclusters formed at different clustering cutoffs using the Average-linkage.**

| Cutoff [Å] | 0 | 1 |
|---|---|---|
| Coarse-grained tunnel network | 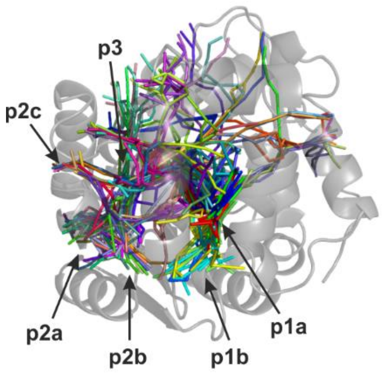 | 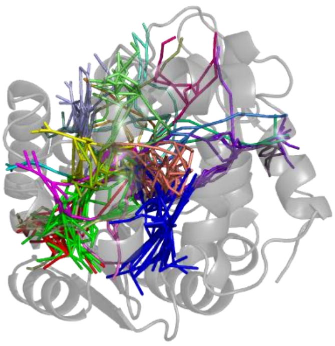 |
| # superclusters | 87 | 24 |
| **Cutoff [Å]** | **2** | **3** |
| Coarse-grained tunnel network | 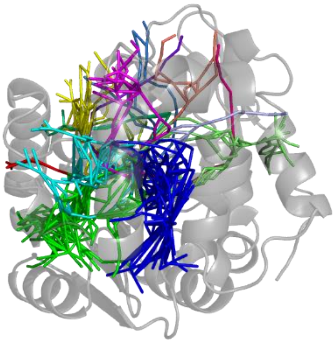 | 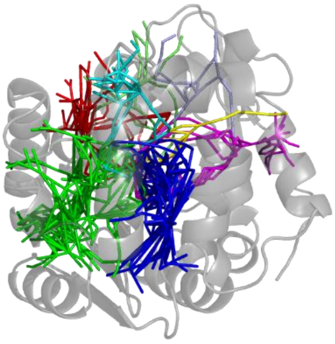 |
| # superclusters | 14 | 9 |
| **Cutoff [Å]** | **4** | **5** |
| Coarse-grained tunnel network | 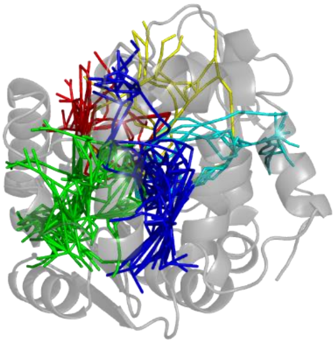 | 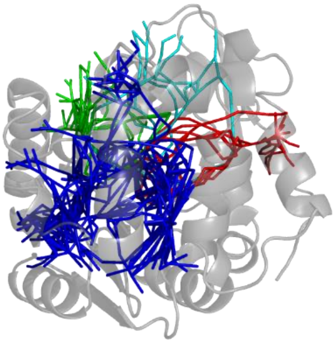 |
| # superclusters | 5 | 4 |
| **Cutoff [Å]** | **10** | **15** |
| Coarse-grained tunnel network | 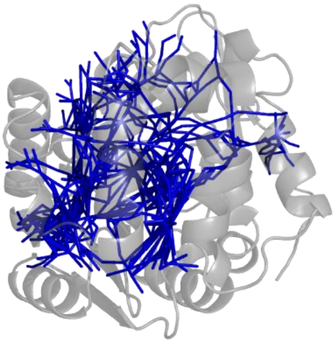 | 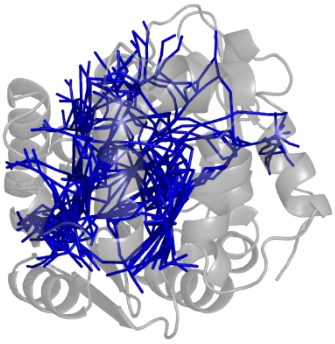 |
| # superclusters | 1 | 1 |

**Table 6. Superclusters formed at different clustering cutoffs using the Complete-linkage.**

| Cutoff [Å] | 0 | 1 |
|---|---|---|
| Coarse-grained tunnel network | 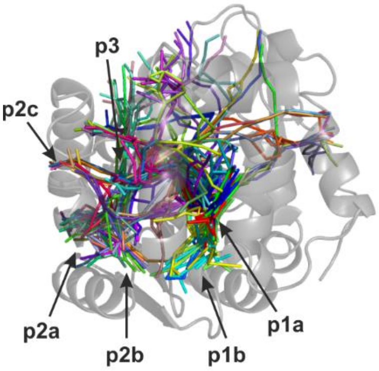 | 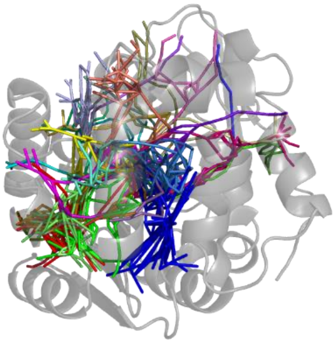 |
| # superclusters | 87 | 27 |
| **Cutoff [Å]** | **2** | **3** |
| Coarse-grained tunnel network | 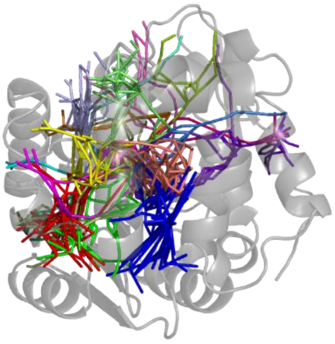 | 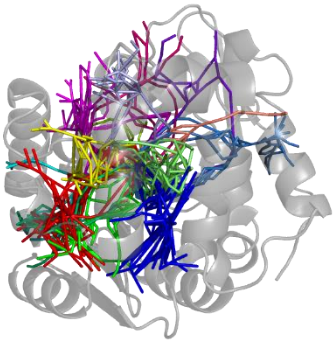 |
| # superclusters | 20 | 14 |
| **Cutoff [Å]** | **4** | **5** |
| Coarse-grained tunnel network | 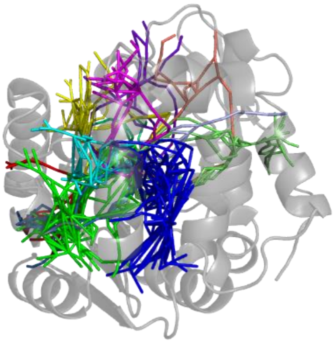 | 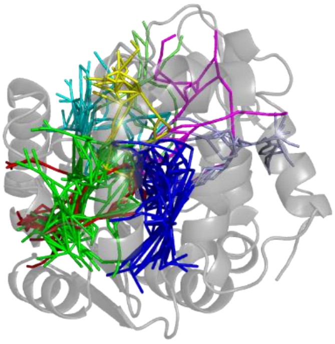 |
| # superclusters | 12 | 8 |
| **Cutoff [Å]** | **10** | **15** |
| Coarse-grained tunnel network | 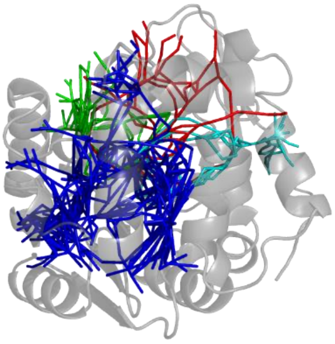 | 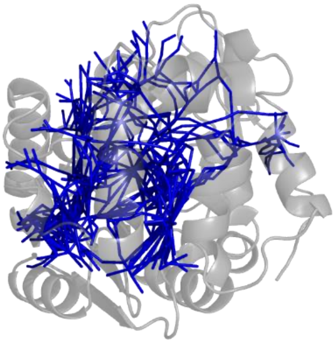 |
| # superclusters | 4 | 1 |

**Table 7. Superclusters formed at different clustering cutoffs using the Ward-linkage.**

| Cutoff [Å] | 0 | 1 |
|---|---|---|
| Coarse-grained tunnel network | 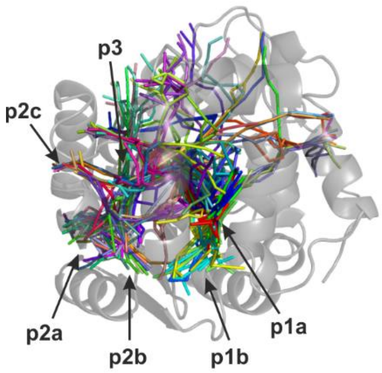  | 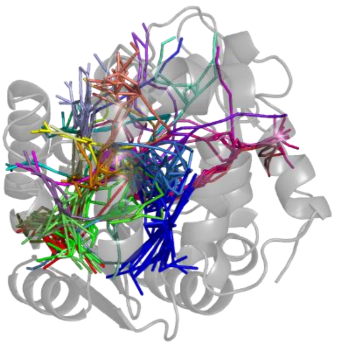 |
| # superclusters | 87 | 31 |
| **Cutoff [Å]** | **2** | **3** |
| Coarse-grained tunnel network | 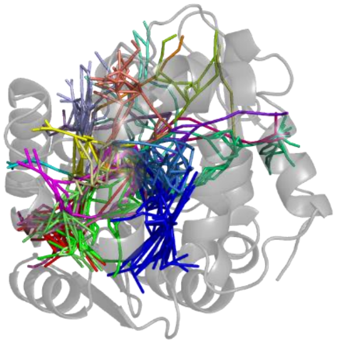 | 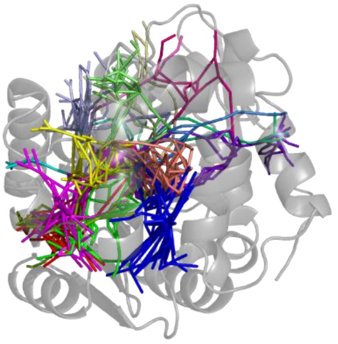 |
| # superclusters | 21 | 17 |
| **Cutoff [Å]** | **4** | **5** |
| Coarse-grained tunnel network | 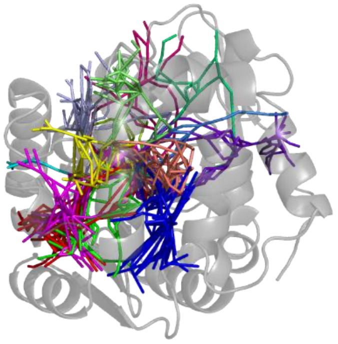 | 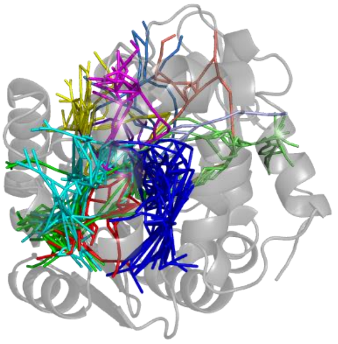 |
| # superclusters | 14 | 11 |
| **Cutoff [Å]** | **10** | **15** |
| Coarse-grained tunnel network | 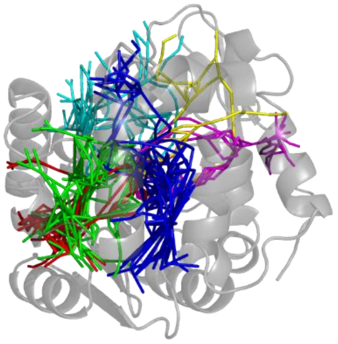 | 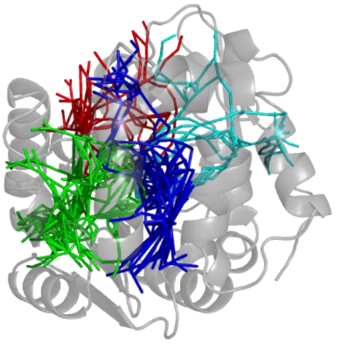 |
| # superclusters | 6 | 4 |

#### 3. REFERENCES


Brezovsky,J. *et al.* (2016) Engineering a de novo transport tunnel. *ACS Catal.*, **6**, 7597–7610.

Campello,R.J.G.B. *et al.* (2013) Density-Based Clustering Based on Hierarchical Density Estimates. In, Pei,J. *et al.* (eds), *Advances in Knowledge Discovery and Data Mining*, Lecture Notes in Computer Science. Springer, Berlin, Heidelberg, pp. 160–172.

Chovancova,E. *et al.* (2012) CAVER 3.0: A Tool for the Analysis of Transport Pathways in Dynamic Protein Structures. *PLoS Comput. Biol.*, **8**, e1002708.

Klvana,M. *et al.* (2009) Pathways and mechanisms for product release in the engineered haloalkane dehalogenases explored using classical and random acceleration molecular dynamics simulations. *J. Mol. Biol.*, **392**, 1339–1356.

Kokkonen,P. *et al.* (2018) Molecular Gating of an Engineered Enzyme Captured in Real Time. *J. Am. Chem. Soc.*, **140**, 17999–18008.

Marques,S.M. *et al.* (2017) Catalytic cycle of haloalkane dehalogenases toward unnatural substrates explored by computational modeling. *J. Chem. Inf. Model.*, **57**, 1970–1989.

Müllner,D. (2013) fastcluster: Fast Hierarchical, Agglomerative Clustering Routines for R and Python. *J. Stat. Softw.*, **53**, 1–18.

Pavelka,A. *et al.* (2016) CAVER: Algorithms for Analyzing Dynamics of Tunnels in Macromolecules. *IEEE/ACM Trans. Comput. Biol. Bioinform.*, **13**, 505–517.
