## Supplementary File 3 for "TransportTools: a library for high-throughput analyses of internal voids in biomolecules and ligand transport through them"

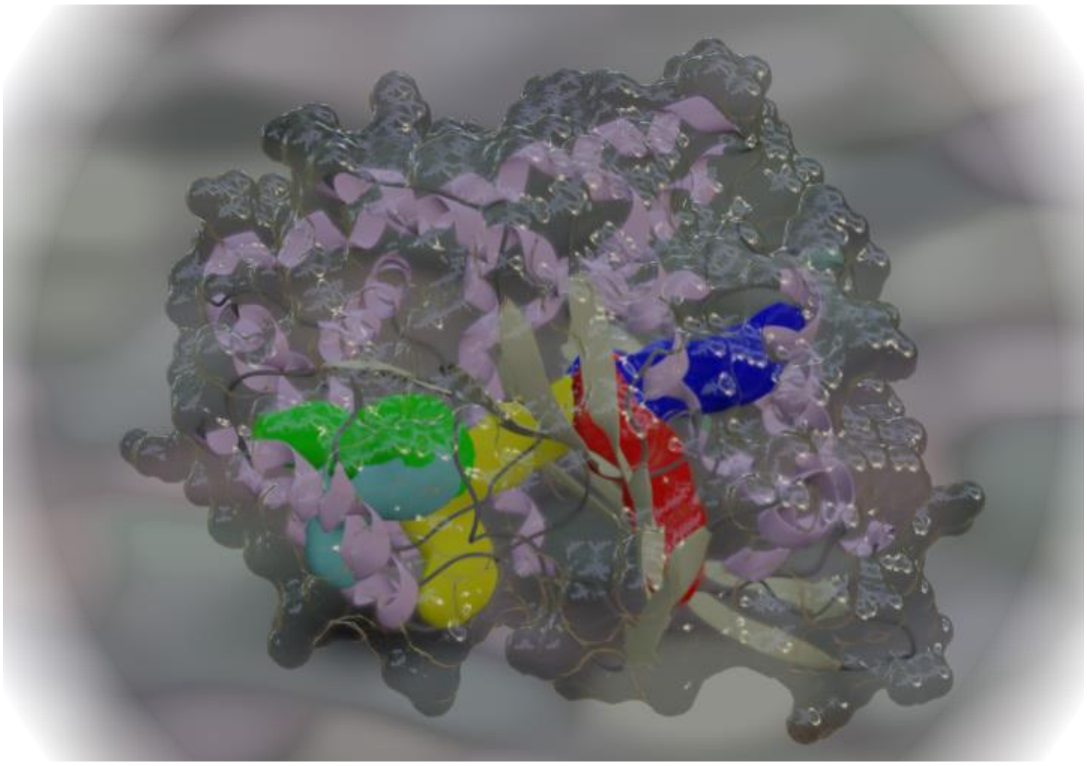

### TransportTools 0.9.0

***A library for massive analyses of internal voids in biomolecules and ligand transport through them***

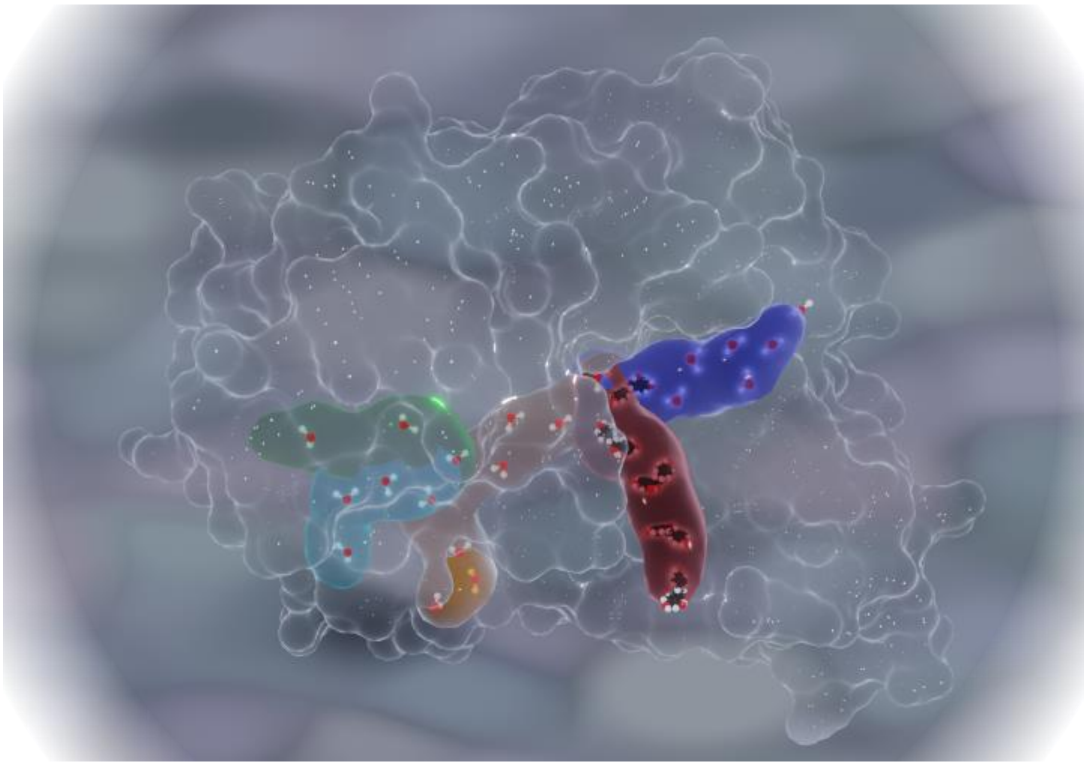

#### USER GUIDE

### 1. *TransportTools* library

*TransportTools* library builds on and extends results of CAVER and AQUA-DUCT analyses to provide comprehensive insights into molecular details of transport processes related to biomolecules.

Using the *TransportTools* engine, end-users can get access to

- efficient analyses of molecular tunnels in extensive molecular dynamics (MD) simulations, including those originating from massively parallel calculations or very long simulations,
- information on molecular voids in the biomolecules with their actual utilization by small molecules,
- rigorous comparison of transport processes in different simulation sets, e.g., contrasting transport in the original system and system perturbed by mutations, different solvents, presence of ligands, etc.

From a programmatic point of view, the library will also help to

- simplify custom-made analyses of transport processes, application of various filters on explored molecular voids,
- facilitate further development of tools focusing on the study of transport processes by providing a rich environment and interface to the popular packages in the field.

***Availability:***
*TransportTools* is licensed under the GNU GPL v3 license, and the source code is available at github.com. Currently, it is fully supported for Linux(-mimicking) operating system and initially also on Windows.

***Minimal requirements:***

- python 3.8
- mdtraj 1.9.4
- biopython 1.78
- fastcluster 1.1.25
- hdbscan 0.8.24
- numpy 1.17.3
- scikit-learn 0.24.0
- scipy 1.5.2

***Suggested installation steps via CONDA:***

1. if you do not have already, get CONDA management system (Please note that such installation does not require superuser privileges on most of the standard Linux-based distributions):
2. create a new environment for transport tools:
   *conda create --name transport_tools python=3.8*
3. activate the environment:
   *conda activate transport_tools*
4. install *TransportTools* and all required dependencies
   *conda install transport_tools -c labbit -c conda-forge*

***Suggested installation steps via PyPi for Linux-based OS*** *(please note that it is not possible to install TransportTools on Windows OS in this straightforward way due to installation problems with mdtraj – one of the dependencies)* ***:***

1. pip install transport_tools

Additionally, you might want to install the following two packages to enhance your experience:

1. AmberTools to speed-up trajectory processing with *pytraj* engine; by following these instructions
2. PyMCubes to enable supercluster visualization with the surface: *pip install --upgrade PyMCubes*

***Troubleshooting:***
If you encounter any problems with installation or the use of *TransportTools* please contact us at github.com.

### 2. *TransportTools* engine

#### 2.1. Overview of *TransportTools* operation

Currently, *TransportTools* engine can perform its action in 10 consecutive stages:

1. Preparatory stage including defining transformations needed for unified analyses
2. Processing of input datasets of tunnel networks
3. Layering tunnel clusters to get their simplified representation
4. Computing distances among the layered clusters
5. Clustering the layered clusters into superclusters and creating initial outputs
6. Filtering superclusters and creating filtered outputs
7. Processing datasets of transport events of ligands
8. Layering transport events to get their simplified representation
9. Assigning transport events to tunnel networks in superclusters and creating initial outputs
10. Filtering supercluster with events and creating filtered outputs with events

Note that stages 7-10 depend on the availability of data on transport events from AQUA-DUCT and hence are in fact optional.

***Usage:***

***Linux-based OS:*** **tt_engine.py [-h] [-c CONFIG_FILENAME] [-w] [-a] [-v] [-l] [--overwrite]**

***Windows OS:*** **tt_engine.exe [-h] [-c CONFIG_FILENAME] [-w] [-a] [-v] [-l] [--overwrite]**

Engine to perform massive analyses of internal voids in biomolecules and ligand transport through them with TransportTool lib.

optional arguments:

| | |
|---|---|
| -h, --help | show this help message and exit |
| -c CONFIG_FILENAME, | File with job configuration; runs TransportTool job with the specified configuration. |
| --config CONFIG_FILENAME | |
| -w, --write_template_config | Writes a template job configuration to file 'tmp_config.ini' and exits. |
| -a, --advanced | Enables extension of a configuration file ('tmp_config.ini') by advanced section. This parameter should be used together with '-w' flag. |
| -v, --version | Prints versions and exits. |
| -l, --license | Prints short license info and exits. |
| --overwrite | Enables cleaning of a non-empty folder with outputs and overwriting of checkpoints files |

***Basic general parameters controlling overall calculations:***

- *start_from_stage*
- *stop_after_stage*
- *num_cpus*
- *output_path*

#### 2.2. Considerations on input preparation

***Minimal required inputs:***

- configuration file
- results of tunnel networks' calculations by CAVER for all analyzed simulations

***Optional inputs:***

- results of tracking ligand movements in simulations by AQUA-DUCT (same or different from those used in tunnel network analyses)
- original simulation trajectories and topologies used to calculate the other inputs

##### 2.2.1. Analysis of tunnel networks across individual MD simulations

The data on tunnel networks are calculated with CAVER3 package that can be obtained from http://caver.cz. In this chapter, we further discuss the essential settings of CAVER3 that provide optimal inputs for *TransportTools* analyses, including the related CAVER3 parameters (*in italics*). For more details on running CAVER3 analyses please consult its user guide.

- protein structures in each simulation MUST be aligned and the simulation boxes imaged
- all simulations MUST use the same/equivalent starting point definition and number of frames. Ideally, we advise using the most stationary residues (or atoms) across all analyzed MD simulations that have their center of mass close to the intended starting point
  - *starting_point_atom*
  - *starting_point_residue*
- tunnel profiles data generation MUST be enabled
  - *generate_tunnel_profiles yes*
- optionally enable identification of bottleneck residues if those are to be analyzed by *TransportTools*
  - *compute_bottleneck_residues yes*
- the smaller probe provides more flexibility to *TransportTools* analyses as well as more insights into "closed" conformation of tunnels. The biological relevance and accuracy of the analyses can be controlled by restricting the supercluster generation to relevant tunnels via *TransportTools* input filters (see chapter 3.2)
  - *probe_radius 0.7*
- *TransportTools* does not use visualizations, histograms, and heatmaps from CAVER. Hence we advise turning those time-demanding options off
  - *visualize_tunnels_per_cluster 1*
  - *save_error_profiles no*
  - *save_dynamics_visualization no*
  - *generate_tunnel_characteristics no*
  - *generate_profile_heat_map no*
  - *generate_histograms no*
  - *generate_bottleneck_heat_map no*
  - *compute_errors no*
- also, do not ask for detection of all tunnel-lining residues as this is extremely slow and of limited precision
  - *compute_tunnel_residues no*

- do NOT use approximate clustering that frequently results in a tunnel cluster of lower quality. Should the CAVER clustering phase represent a considerable analysis hurdle, it is preferable to split your trajectories into several parts and analyze them separately, leaving their combination to *TransportTools* instead of relying on simplified approximate clustering.
    - *do_approximate_clustering no*
- smaller CAVER clustering cutoff provides a more refined view on tunnel clusters leaving the control over overall clustering for *TransportTools*. Hence we suggest using the values between 2 and 4 here.
    - *clustering_threshold 3*

***TransportTools utilizes the following results of CAVER calculations:***

- *bottlenecks.csv* (optional data on tunnel bottlenecks residues)
- *tunnel_profiles.csv* (required data on tunnel networks)
- *reference PDB file* (required, its naming depends on the user-defined naming of trajectory frames when analyzed by CAVER, currently the same naming across analyzed trajectories is required)
- *v_origins.pdb* (required data on an ensemble of starting points of tunnel network)

The location of these files (highlighted in red) within the standard structure of the output folder generated by CAVER is following:

```
├── analysis
│   ├── bottlenecks.csv
│   ├── ids_and_clusters.csv
│   └── tunnel_profiles.csv
├── data
│   ├── clustering.txt
│   ├── clusters_timeless
│   ├── end_zone.pdb
│   ├── origins.pdb
│   ├── start_zone.pdb
│   ├── stripped_system.000500.pdb
│   ├── times.txt
│   ├── tree_.txt
│   ├── tree.txt
│   ├── tunnel_edges
│   ├── tunnels
│   └── v_origins.pdb
├── log.txt
├── pymol
│   ├── modules
│   ├── view_plugin.py
│   ├── view.py
│   ├── view_timeless.py
│   ├── view_trajectory.py
│   └── zones.py
├── vmd
│   ├── scripts
│   ├── vmd.bat
│   ├── vmd.sh
│   ├── vmd_timeless.bat
│   └── vmd_timeless.sh
├── warnings.txt
└── whole_clustering.csv
```

##### 2.2.2. Analysis of ligand movements across individual MD simulations

The data on transport events connected with ligand entry/release from the site of interest can be calculated with ligand tracking capabilities of AQUA-DUCT package that can be obtained from www.aquaduct.pl. In this section, we further discuss the essential settings of AQUA-DUCT that provide optimal inputs for *TransportTools* analyses, including the related AQUA-DUCT parameters (*in italics*). For more details on running AQUA-DUCT analyses, please consult its user guide.

- protein structures in each simulation MUST be aligned and the simulation boxes imaged
- define *Object* of AQUA-DUCT calculation using same starting point definition as for CAVER, for example:
    - *object = (resname YOUR_LIG) and (sphzone DIST (resnum RID1 or resnum RID2 or … ))*
        - *YOUR_LIG* - residue name analyzed ligand
        - RID1, RID2, … - residue IDs matching those used in *starting_point_residue* CAVER parameters
        - DIST – distance from the center of mass of specified residues to the AQUA-DUCT *Object*'s boundary. Note that this distance should be used to define *TransportTools* parameter *event_min_distance* (see chapter 3.7)
- use biomolecule's backbone as the *Scope* of AQUA-DUCT calculation to better agree with standard ends of tunnels calculated by CAVER3 in sections *[traceable_residues]* and *[raw_paths]*
    - *scope_convexhull = True* - in sections *[traceable_residues]* and *[raw_paths]*
    - *scope = backbone* - use protein backbone to define the convex hull
- avoid trimming of separate paths with autobarber, since *TransportTools* processing algorithm for these paths benefits from their extended forms
    - *auto_barber = False*
- enable storing of traces of transport events as AQUA-DUCT raw paths in section *[visualize]*
    - *paths_raw = True*
    - *paths_states = True*
    - *show_molecule = protein*
    - *show_molecule_frames = 0*

***TransportTools utilizes the following results of AQUA-DUCT calculations:***

- *5_analysis_results.txt* (required general summary on transport events)
- *6_visualize_results.tar.gz* (required data on traces of transport events)

The location of these files (highlighted in red) within the standard structure of the output folder generated by AQUA-DUCT is following:

```
├── 1_traceable_residues_data.dump
├── 2_raw_paths_data.dump
├── 3_separate_paths_data.dump
├── 4_inlets_clusterization_data.dump
├── 5_analysis_results.txt
├── 5_analysis_results.txt.csv
├── 6_visualize_results.py
├── 6_visualize_results.tar.gz
└── AQUA-DUCT_config.txt
```

#### 2.3. Basic tutorial on *TransportTools* engine usage

This chapter will cover the basic functionalities of *TransportTools* engine. The commands assume **Linux-based OS**, when using Windows OS you might need to modify your approach accordingly, e.g., using tt_engine.exe in place of tt_engine.py. The tutorial is based on two trajectories of 1,000 frames from MD simulations with only several preselected water molecules that exhibit interesting transport events. Note that both MD trajectories are already aligned and imaged as required by both CAVER and AQUA-DUCT.

First, we provide a guideline on how to obtain optimal CAVER and AQUA-DUCT results, including a short description of the crucial settings as well as generated data. Further, we briefly explain the preparation of the configuration file for *TransportTools* run. This is followed by several exemplary runs utilizing different settings (i.e., limiting the number of calculation stages, applying filtering based on the tunnels' properties or the number of events, etc.), which is aimed to provide the User with a basic understanding of the calculation process. Finally, we describe the results in terms of basic statistics and visualization in order to supplement the overall view of *TransportTools* engine operation.

##### 2.3.1. Input data generation

To continue with the tutorial, please download *tutorial.tar.gz* archive and extract the content (**Figure 1**) in a location on your drive which will be suitable for further calculations and analyses with CAVER, AQUA-DUCT and *TransportTools* – you will need roughly ~1GB of empty space. We strongly recommend installing AQUA-DUCT in the same Python environment as *TransportTools* and using this environment for CAVER, AQUA-DUCT and *TransportTools* calculations to avoid any troubles related to dependencies. The tutorial archive contains the following folders and files:

```
tutorial
├── custom_analysis
│   └── custom_analysis.py
├── precomputed_inputs
│   ├── md1
│   │   ├── aquaduct
│   │   └── caver
│   └── md2
│       ├── aquaduct
│       └── caver
├── simulations
│   ├── md1
│   │   ├── structure.parm7
│   │   └── trajectory.nc
│   └── md2
│       ├── structure.parm7
│       └── trajectory.nc
└── templates
    ├── AQUA-DUCT_config.txt
    └── run_CAVER.sh
```

```
tutorial
├── custom_analysis
│   └── custom_analysis.py
├── precomputed_inputs
│   ├── md1
│   │   ├── aquaduct
│   │   └── caver
│   └── md2
│       ├── aquaduct
│       └── caver
├── simulations
│   ├── md1
│   │   ├── aquaduct
│   │   ├── caver
│   │   ├── structure.parm7
│   │   └── trajectory.nc
│   └── md2
│       ├── aquaduct
│       ├── caver
│       ├── structure.parm7
│       └── trajectory.nc
└── templates
    ├── AQUA-DUCT_config.txt
    └── run_CAVER.sh
```

**Figure 1 Tree of tutorial archive content (left) and folder structure needed for the further steps in case of following the entire tutorial and running AQUA-DUCT and CAVER calculations (right).**

Before we start, please note, that in case one would like to avoid running CAVER and AQUA-DUCT calculations, we provide *precomputed_inputs* folder in the tutorial archive. In this case, you can skip to chapter 2.3.2. Otherwise, in each folder with MD simulation replica we need to create subfolders for CAVER and AQUA-DUCT calculations - let's call them *caver* and *aquaduct*, respectively (**Figure 1**). In *templates* two input files are provided that will enable to set up and run CAVER and AQUA-DUCT calculations producing results compatible with the *TransportTools* workflow. Before moving these files to MD folders the *CAVER_HOME* parameter in *run_CAVER.sh* script has to be adjusted accordingly to your caver folder location, e.g., if you have downloaded and extracted CAVER software into /home/user/software folder, *CAVER_HOME* should point to *caver_3.0/caver* subfolders unpacked in this folder:

*CAVER_HOME=/home/user/software/caver_3.0/caver*

Next, copy *run_CAVER.sh* and *AQUA-DUCT_config.txt* files to corresponding *caver* and *aquaduct* folders in md1 and md2. Before running calculations, one important action has to be done on both files. *STARTING_POINT_ATOMS* parameter in *run_CAVER.sh* script defines the starting point for the tunnel searches (**Figure 2**) and special attention should be paid to the proper selection of those. In the current system, we select atoms that are the least mobile in the set of MD simulations and define starting point located in the active site cavity of the enzyme. Analogously, the *object* parameter in *AQUA-DUCT_config.txt* should correspond to the same region of the protein (**Figure 3**). In current settings, it is defined by the same enzyme atoms as a *STARTING_POINT_ATOMS* in CAVER and the active site is specified as 6.0 Å around this point.

***CAVER:***

```
STARTING_POINT_ATOMS="1608 1634 2602 3867"

# those are following atoms:
# ATOM   1608  CG  ASP   103
# ATOM   1634  CD2 TRP   104
# ATOM   2602  N   PHE   165
# ATOM   3867  N   LEU   243
```

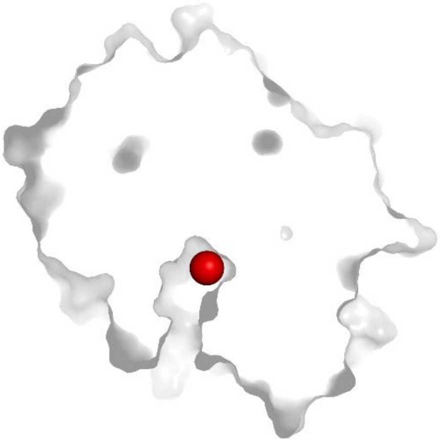

**Figure 2 The definition of the starting point (red sphere) for tunnel calculations in CAVER.**

***AQUA-DUCT:***

```
scope = backbone

object = (resname WAT) and (sphzone 6.0 (
(resnum 103 and name CG) or
(resnum 104 and name CD2) or
(resnum 243 and name N) or
(resnum 165 and name N)))

# those are following atoms:
# ATOM   1608  CG  ASP   103
# ATOM   1634  CD2 TRP   104
# ATOM   2602  N   PHE   165
# ATOM   3867  N   LEU   243
```

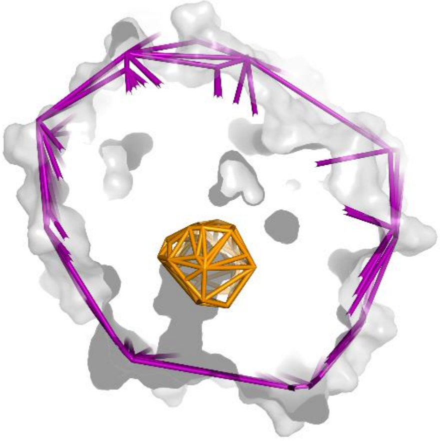

**Figure 3 The definition of *scope* (purple) and *object* (gold) for transport event tracking by AQUA-DUCT.**

With template files moved to appropriate folders, we are prepared to generate input data for *TransportTools*. To execute CAVER calculations, enter *simulations/md1/caver* folder and run the following command in the terminal:

*./run_CAVER.sh*

Once the calculations are finished, execute the same command in *simulations/md2/caver* folder.

Tip: the amount of memory taken for the calculation can be reduced/increased according to available resources. To do so, simply modify -Xmx8000m in run_CAVER.sh script. Currently set up -Xmx8000m is equivalent to ~8GB of memory used. Running CAVER calculations in parallel for both MD simulations may consume whole memory on your system, hence we recommend completing this stage in a step-wise manner. A single calculation may take roughly ~30 minutes.

The desired completion of CAVER calculation will be indicated by the message "*Finished successfully*". Next, enter *simulations/md1/aquaduct* folder and run AQUA-DUCT calculations using the pre-prepared configuration file. Use the following command:

```
valve_run -c AQUA-DUCT_config.txt --debug-file aq.out
```

Current settings enable the generation of desired results rapidly. However, please note that analyzed trajectories contain only a few selected water molecules to reduce the size of files, time of calculations, and the complexity of the analysis. In standard settings, we would most likely take all water/ligand molecules present in the system into account, hence making analysis more time-consuming and computationally demanding. Simplified visualization of the CAVER and AQUA-DUCT input data is presented in **Figure 4**.

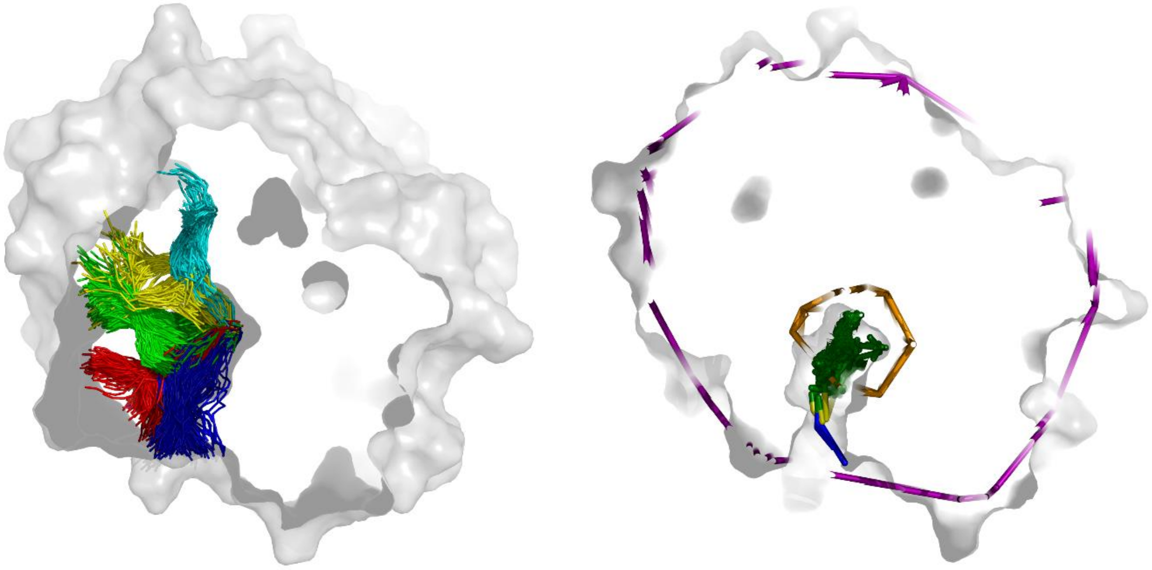

**Figure 4 Visualization of input data used by *TransportTools*.** First five best CAVER clusters (colored sets of lines) selected for visualization generated from timeless clusters (left). AQUA-DUCT transport events detected for analyzed water molecules (green lines emerging from the golden *object*) present in the system (right).

#### 2.3.2. Preparing configuration file

All necessary input data from CAVER and AQUA-DUCT have been generated and we can continue with the preparation of *TransportTools* run. We will start with the generation of the configuration file. In the *tutorial* folder, run the following command:

```
tt_engine.py -w
```

The configuration file has to be adjusted accordingly to our needs. We will proceed with the modification gradually. First, **[INPUT_PATHS]** to the CAVER and AQUA-DUCT results, as well as the analyzed trajectories, have to be specified (see **Figure 5** representing the tree of current *simulations* folder):

- In our case, all input data are located in the *simulations* folder. Based on those, we set up *caver_results_path, aquaduct_results_path,* and *trajectory_path* parameters as *simulations.* Note that if you have skipped CAVER and AQUA-DUCT calculations and used precomputed inputs instead, please set *caver_results_path, aquaduct_results_path* to *precomputed_inputs*, and *trajectory_path* parameter as *simulations.*

- *caver_results_folder_pattern*, *aquaduct_results_folder_pattern*, and *trajectory_folder_pattern* are designed to match particular folder patterns and work on the basis of UNIX-like patterns – set up these to '*' matching everything or more specifically use 'md*' to match md1 and md2 subfolders.
- Assign *caver_results_relative_subfolder_path* to *caver* according to the naming of the CAVER results folder.
- Assign *aquaduct_results_relative_tarfile* and *aquaduct_results_relative_summaryfile* to *aquaduct/6_visualize_results.tar.gz* and *aquaduct/5_analysis_results.txt* for AQUA-DUCT results, respectively.
- Define *trajectory_relative_file* and *topology_relative_file* as *trajectory.nc* and *structure.parm7* correspondingly to the names of the trajectory and topology files used in the tutorial.

Properly defined parameters related to paths of input folders and files are presented in **Figure 6**.

```
simulations/
├── md1
│   ├── aquaduct
│   │   ├── 1_traceable_residues_data.dump
│   │   ├── 2_raw_paths_data.dump
│   │   ├── 3_separate_paths_data.dump
│   │   ├── 4_inlets_clustering_data.dump
│   │   ├── 5_analysis_results.txt
│   │   ├── 5_analysis_results.txt.csv
│   │   ├── 6_visualize_results.py
│   │   ├── 6_visualize_results.tar.gz
│   │   ├── aq.out
│   │   └── AQUA-DUCT_config.txt
│   ├── caver
│   │   ├── analysis
│   │   ├── data
│   │   ├── log.txt
│   │   ├── run_CAVER.sh
│   │   ├── warnings.txt
│   │   └── whole_clustering.csv
│   ├── structure.parm7
│   └── trajectory.nc
└── md2
    ├── aquaduct
    │   ├── 1_traceable_residues_data.dump
    │   ├── 2_raw_paths_data.dump
    │   ├── 3_separate_paths_data.dump
    │   ├── 4_inlets_clustering_data.dump
    │   ├── 5_analysis_results.txt
    │   ├── 5_analysis_results.txt.csv
    │   ├── 6_visualize_results.py
    │   ├── 6_visualize_results.tar.gz
    │   ├── aq.out
    │   └── AQUA-DUCT_config.txt
    ├── caver
    │   ├── analysis
    │   ├── data
    │   ├── log.txt
    │   ├── run_CAVER.sh
    │   ├── warnings.txt
    │   └── whole_clustering.csv
    ├── structure.parm7
    └── trajectory.nc
```

**Figure 5 Tree of *simulations* folder representing files and catalogs necessary for *TransportTools* calculations.**

```
[INPUT_PATHS]
# CAVER results
caver_results_path = simulations
caver_results_folder_pattern = *
caver_results_relative_subfolder_path = caver
# AQUA-DUCT results
aquaduct_results_path = simulations
aquaduct_results_folder_pattern = *
aquaduct_results_relative_tarfile = aquaduct/6_visualize_results.tar.gz
aquaduct_results_relative_summaryfile = aquaduct/5_analysis_results.txt
# Source MD trajectories
trajectory_path = simulations
trajectory_folder_pattern = *
trajectory_relative_file = trajectory.nc
topology_relative_file = structure.parm7
```

**Figure 6 Definition of parameters related to input paths in *TransportTools* configuration file.**

Following the settings shown in **Figure 5** and **Figure 6**, assign paths to input folders and files in your tmp_config.ini file and rename it:

| *mv tmp_config.ini config.ini* |
|---|

Before the first execution of *TransportTools*, several basic **[CALCULATIONS_SETTINGS]** have to be adjusted:

- We will start calculations from default stage 1 (*start_from_stage = 1*) but as an initial run, we aim to finish just after this stage to see whether the program starts correctly. Hence, set up the desired stage on which we expect to stop calculations (for now *stop_after_stage = 1*).
- It is possible to adjust the number of CPU threads depending on the resources we have, we will use half of the threads available to our machines (e.g., *num_cpus = 4*).
- Optionally, reference structure can be specified, in case it is not, CAVER representative will be automatically taken from a *caver* folder (*pdb_reference_structure = None*).
- Regarding settings related to parsing of tunnel clusters from CAVER results, we want to read bottlenecks information from CAVER outputs (*process_bottleneck_residues = True*).
- A number of snapshots per simulation has to be specified (*snapshots_per_simulation = 1000*) unless it can be detected from provided simulations (*snapshots_per_simulation = None)*, together with the difference in IDs of *mdtraj/pytraj* frames and caver snapshots (*caver_traj_offset = 1*).
- Finally, we define which part of the snapshot filename contains its ID (*snapshot_id_position = 1*) and what is the delimiter to split the snapshot name (*snapshot_delimiter = .*): Example: in our case the snapshot name is *'stripped_system.000500.pdb'* in folder *precomputed_inputs/md2/caver/data*. If the file name is split using dot (.) as a delimiter we get ['*stripped_system*', *'000500'*, 'pdb']. Here, the snapshot ID (500) is located in the 1st element (counting from 0, according to Python convention).
- Last but not least, an important parameter defining the way of AQUA-DUCT transport events processing has to be specified. Namely, we have to set up the closest distance of the event molecule from the starting point to be processed, and this should be compatible with *sphzone* (here 6.0) in the object definition for AQUA-DUCT calculations (*event_min_distance = 6.00*).

At the moment, we configured all important parameters, and we can proceed with a test run of the first tt_engine.py stage. Enter the *tutorial* folder and execute the following command:

| *tt_engine.py -c config.ini* |
|---|

A successfully configured program utilizing a correctly prepared configuration file should report a message as presented in **Figure 7**.

```
bart@BIT3:tutorial$ tt_engine.py -c config.ini
12:13:05 - transport_tools.libs.ui - INFO - ======== ********************************************* ========
12:13:05 - transport_tools.libs.ui - INFO - ========       TransportTools execution started        ========
12:13:05 - transport_tools.libs.ui - INFO - ======== ********************************************* ========
12:13:06 - __main__ - INFO - ------------------------------------------------------------------------------
12:13:06 - __main__ - INFO - STAGE 1 - Preparatory stage for unified analyses
12:13:06 - __main__ - INFO - ------------------------------------------------------------------------------
12:13:06 - transport_tools.libs.tools - INFO - Computing transformation to unify the simulations coordinate system
s.
12:13:06 - transport_tools.libs.tools - INFO - Aligning to reference file simulations/md1/caver/data/stripped_syst
em.000500.pdb.
12:13:06 - transport_tools.libs.ui - INFO - |████████████████████████████████████████████████████████████| 100.0%
12:13:07 - transport_tools.libs.ui - INFO - Computation took: 0:00:00.
12:13:07 - transport_tools.libs.tools - INFO - CHECKPOINT saved to: results/_internal/checkpoints/stage001.dump
12:13:07 - __main__ - INFO - Requested to stop after this STAGE
12:13:07 - transport_tools.libs.ui - INFO - ======== ********************************************* ========
12:13:07 - transport_tools.libs.ui - INFO - ========         Overall elapsed time: 0:00:01         ========
12:13:07 - transport_tools.libs.ui - INFO - ======== ********************************************* ========
```

**Figure 7 Testing tt_engine.py execution with analysis stopped after the first stage.**

Note that we will be using PyMOL package to visualize the results. When PyMOL based on Python2 is applied, it is necessary to have the support of legacy versions enabled in **[ADVANCED_SETTINGS]** section of the configuration file to generate compatible data used by the visualization scripts (see chapter 2.3.7 for details).

##### 2.3.3. Understanding calculation progress

Proper settings of the program and *config.ini* file were already confirmed, and we can continue with a complete analysis. Reset *stop_after_stage* parameter to *None,* and run full analysis. By running the same command as previously, an error will be reported that checkpoint file already exists, and *TransportTools* will suggest specifying a different name or enable to overwrite files by using '*--overwrite*' option. Another solution is to remove the *result* folder which will solve this problem as well. We will adjust the command according to the reported suggestion:

| *tt_engine.py -c config.ini --overwrite* |
| --- |

Analyzed trajectories are reduced to be computationally as cheap as possible for the purposes of this tutorial, hence the program finished analysis after ~1 minute. With larger trajectories containing an enormous amount of tunnels and ligands to analyze, the execution might take hours. Note, that at the initial stage of the analysis, *TransportTools* detected 100% coverage of source MD trajectories and CAVER tunnels data (**Figure 8**), and suggested using a more accurate protocol for *ambiguous_event_assignment_resolution*.

```
bart@BIT3:tutorial$ tt_engine.py -c config.ini --overwrite
12:15:55 - transport_tools.libs.ui - INFO - ======== ********************************************* ========
12:15:55 - transport_tools.libs.ui - INFO - ========       TransportTools execution started        ========
12:15:55 - transport_tools.libs.ui - INFO - ======== ********************************************* ========
12:15:56 - transport_tools.libs.config - WARNING -
Since it seems that source MD trajectories as well as tunnel data (tunnel data coverage = 100%) are available,
 consider using 'exact_matching' for 'ambiguous_event_assignment_resolution' parameter to provide the most reliabl
e assignment of events to supercluster at a cost of only mild increase in computation time.
```

**Figure 8 Warning message regarding the suboptimal setting of *ambiguous_event_assignment_resolution*.**

Therefore, we will update *ambiguous_event_assignment_resolution* to '*exact_matching*' and rerun the calculations.

Note: As we can see, the overall time of the calculation was not affected significantly by the utilization of a more accurate protocol. Importantly, this behavior is also valid for larger datasets – only a mild increase in computational cost should be expected.

With progressing analysis, the information is printed to the User stating the current stage of the process and most important activities performed. For our run, we should see the following stages and activities:

- stage 1 - Preparatory stage for unified analyses
  - Computing transformation to unify the simulations coordinate systems
  - Aligning to reference file *simulations/md1/caver/data/stripped_system.000500.pdb*
- stage 2 - Processing datasets of tunnel networks from CAVER results
  - Processing 2 tunnel networks
- stage 3 - Layering tunnel clusters to get their simplified representation
  - Computing layered representation for 170 tunnel clusters
- stage 4 - Computing distances among the layered clusters
  - Computing distances for 170 tunnel clusters
- stage 5 - Clustering the layered clusters into superclusters and creating initial outputs
  - Clustering tunnel clusters into superclusters using complete-linkage agglomerative clustering with distance cutoff 2.00 Å
  - Identified 45 superclusters
  - Aggregating supercluster data for 45 superclusters
- stage 6 - Filtering superclusters and creating filtered outputs
  - Filtering 45 supercluster profiles
  - 45 superclusters kept after filtering
- stage 7 - Processing datasets of transport events from AQUA-DUCT results
  - Processing 2 AQUA-DUCT networks
- stage 8 - Layering transport events to get their simplified representation
  - Computing layered representation for 11 transport events
- stage 9 - Assigning layered events to tunnel networks in superclusters and creating initial outputs with events
  - Assigning 11 transport events to 45 superclusters
  - 1 transport event was not assigned to superclusters
- stage 10 - Filtering supercluster with events and creating filtered outputs with events
  - Filtering 45 supercluster profiles
  - 45 superclusters kept after filtering

At each stage of the calculations, *TransportTools* generate a checkpoint dump file in *results/_internal/checkpoints* folder. These files enable the restart of the process from any stage and will be discussed later. Now, we will briefly describe actions reported in the progress of calculations. In stage 1, systems were prepared for unified analyses including alignment to reference structure.

Next, stage 2 covered the processing of two CAVER tunnel networks, where each network corresponds to a particular MD trajectory from our dataset. Stage 3 and stage 4 were concentrated on the processing of CAVER clusters, note that 170 tunnel clusters were detected. They were further layered to simplify their representation and distances among them were computed, which were necessary for the following clustering (stage 5). At this stage, the complete-linkage agglomerative clustering was used with a cutoff of 2 Å (could be adjusted in the configuration file), resulting in the generation of 45 superclusters and followed by the generation of initial outputs. *TransportTools* enable filtering of the superclusters, which is performed in stage 6. Nevertheless, we have not specified any filtering parameters for these calculations. Hence all initially reported superclusters were retained.

Stage 7 corresponds to the processing of AQUA-DUCT transport event datasets – two networks, analogously to CAVER networks, one for each MD trajectory. Next, transport events were layered to simplify the representation and assigned to superclusters (stage 8 and stage 9). For our dataset, 11 transport events were detected and most of these events were successfully assigned to some supercluster with the exception of one of them. Stage 10 provides an opportunity of filtering superclusters based on the events. For example, we can ask for a supercluster with the desired number of entries, releases or the total number of events. We have not specified any filtering criteria in the configuration file, hence again all detected superclusters were retained. All this information is also reported in the logfile (by default, '*transport_tools.log*' in the *results* folder), together with the parameters used. Please keep in mind that option '*--overwrite*' does not delete/overwrite logfile, which is extended with the following runs as long as it is not deleted manually.

##### 2.3.4. Analyzing superclusters

At this point, as the calculation process should be already sufficiently understandable, we can focus on the primary analysis of the results generated during executed tt_engine.py run. First, a short description of the folders and files produced (**Figure 9**). In the *data* folder, we will find detailed information about initial and filtered superclusters, supercluster events, and bottlenecks. *_internal* folder holds all the intermediate data generated during a run, which are also crucial for the rerunning calculations from a particular stage. However, we will not focus on this part of the outputs in this tutorial. Further, *visualization* folder with scripts to visualize results in PyMOL, and finally *statistics* folder where the results are comprehensively summarized.

```
results
├── data
│   └── super_clusters
├── _internal
│   ├── checkpoints
│   ├── clustering
│   ├── layered_data
│   ├── network_data
│   ├── super_cluster_pathsets
│   ├── super_cluster_profiles
│   └── transformations
├── statistics
│   ├── 1-initial_tunnels_statistics_bottleneck_residues.txt
│   ├── 1-initial_tunnels_statistics.txt
│   ├── 2-filtered_tunnels_statistics_bottleneck_residues.txt
│   ├── 2-filtered_tunnels_statistics.txt
│   ├── 3-initial_events_statistics_bottleneck_residues.txt
│   ├── 3-initial_events_statistics.txt
│   ├── 4-filtered_events_statistics_bottleneck_residues.txt
│   └── 4-filtered_events_statistics.txt
├── transport_tools.log
└── visualization
    ├── 1-visualize_initial_tunnels.py
    ├── 2-visualize_filtered_tunnels.py
    ├── 3-visualize_initial_events.py
    ├── 4-visualize_filtered_events.py
    └── sources
```

**Figure 9 Tree of files and folders generated during the whole calculation pipeline.**

As mentioned in the previous section, without applying additional filtering options, we have obtained 45 superclusters, which are presented in **Figure 10** using a *visualization/1-visualize_initial_tunnels.py* script. Visualization of the entire superclusters set results in a relatively complex picture. Hence in the following steps, we will perform a more focused analysis using filtering criteria. In order to visualize all superclusters use:

| |
|---|
| graphical user interface: *PyMOL > File > Run script… > 1-visualize_initial_tunnels.py*<br>terminal: *pymol 1-visualize_initial_tunnels.py* |

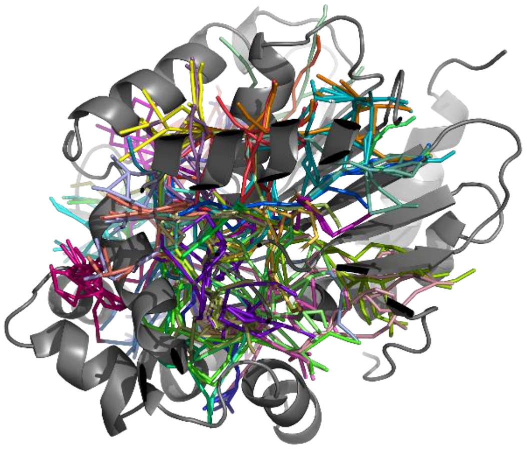

**Figure 10 Initial visualization of detected 45 superclusters generated based on 170 CAVER clusters from 2 datasets.**

Note: When PyMOL based on Python2 is applied, it is necessary to have the support of legacy versions enabled in **[ADVANCED_SETTINGS]** section of the configuration file to generate compatible data used by the visualization scripts (see chapter 2.3.7 for details).

To analyze details of all superclusters reported, open the 1-*initial_tunnels_statistics.txt* file from the *statistics* folder in any text editor. As shown in **Figure 11** upper part of the file informs the User regarding the source data (statistics for overall, number of simulations – 2, and amount of snapshots per simulation – 1000). Further, parameters used for pre-selection of tunnels for clustering are defined (in this case, default from the configuration file) and applied active filters (turned off for these calculations). The remaining part of the file is occupied by a statistics table containing all 45 superclusters. The first column corresponds to supercluster ID (SC_ID), further, the number of the simulations in which particular supercluster was detected (No_Sims), the total number of frames containing supercluster (Total_No_Frames), followed by more general properties such as average bottleneck radius (Avg_BR), average length (Avg_Len), together with standard deviations (StDev columns), maximum bottleneck radius (Max_BR), and other properties. Please note that the table is not sorted by the supercluster IDs. Hence the order in the first column is not always consecutive.

We requested to process the bottleneck residues in our calculations (*process_bottleneck_residues* = *True*), hence tt_engine also produced a *1-initial_tunnels_statistics_bottleneck_residues.txt* summary file containing information about bottleneck forming residues together with frequencies, which connects the description of each supercluster with determining residues in protein structures. Finally, transport events (sum of entries and releases) and entries or releases separately, which are assigned to particular superclusters are reported in a *3-initial_events_statistics.txt* file as Num_Events, Num_entries, and Num_releases, respectively. Each of these files also has its '*filtered*' equivalent, but as the filtering was turned off, they include precisely the same information.

```
Statistics for: overall
Total simulations = 2
No snapshots per simulation = 1000

Parameters used for pre-selection of input tunnels for clustering:
length >= 5.00 A
radius >= 0.75 A
curvature <= 2.00

Active tunnel filters:
length = (0.00, 999.00) A
radius = (0.00, 999.00) A
curvature = (1.00, 999.00)
occurred in at least 0 simulations
occurred in at least 0 snapshots
occurred in at least 0.000 snapshots on average per simulation

SC_ID, No_Sims, Total_No_Frames, Avg_No_Frames, Avg_BR, StDev, Max_BR, Avg_Len,  StDev, Avg_Cur, StDev, Avg_throug,   StDev, Priority
   1,       2,            1982,         991.0, 1.108, 0.186, 1.950, 14.139,  1.604,   1.249, 0.090,    0.52990, 0.08014,  1.05026
   2,       2,            1387,         693.5, 0.860, 0.118, 1.423, 16.617,  2.812,   1.415, 0.208,    0.40020, 0.09212,  0.55508
   3,       2,             965,         482.5, 0.953, 0.170, 1.515, 16.324,  4.271,   1.301, 0.206,    0.41384, 0.13795,  0.39935
   4,       2,             829,         414.5, 0.790, 0.076, 1.157, 14.239,  2.060,   1.256, 0.142,    0.33249, 0.07567,  0.27564
   5,       2,             524,         262.0, 0.757, 0.056, 1.030, 21.403,  2.700,   1.286, 0.138,    0.20701, 0.05517,  0.10847
  11,       2,             239,         119.5, 0.843, 0.137, 1.239, 18.531,  2.515,   1.297, 0.139,    0.28304, 0.10718,  0.06765
   7,       2,             395,         197.5, 0.758, 0.053, 0.970, 25.757,  4.175,   1.833, 0.340,    0.16593, 0.05178,  0.06554
   9,       2,             431,         215.5, 0.749, 0.046, 0.979, 29.232,  7.798,   1.603, 0.296,    0.14550, 0.05576,  0.06271
   6,       2,             377,         188.5, 0.750, 0.043, 1.029, 26.692,  4.609,   1.916, 0.283,    0.16543, 0.05013,  0.06237
   8,       2,             197,          98.5, 0.783, 0.139, 1.599, 16.920,  2.982,   1.488, 0.200,    0.30586, 0.10438,  0.06025
  10,       2,             424,         212.0, 0.747, 0.043, 0.952, 32.572,  7.788,   1.649, 0.443,    0.13197, 0.05970,  0.05596
  13,       2,             277,         138.5, 0.740, 0.041, 0.952, 25.258,  2.774,   1.381, 0.149,    0.13084, 0.04061,  0.03624
  12,       2,             200,         100.0, 0.739, 0.041, 0.951, 21.090,  2.802,   1.527, 0.192,    0.18074, 0.05708,  0.03615
  18,       2,             362,         181.0, 0.733, 0.032, 0.909, 40.726, 10.791,   1.766, 0.336,    0.06689, 0.04090,  0.02421
  17,       2,             226,         113.0, 0.740, 0.036, 0.856, 35.778,  9.694,   1.802, 0.743,    0.10157, 0.05029,  0.02295
  14,       2,             155,          77.5, 0.747, 0.039, 0.895, 27.926,  3.638,   1.518, 0.162,    0.14491, 0.04191,  0.02246
  15,       2,             217,         108.5, 0.748, 0.047, 0.945, 36.641,  4.069,   1.787, 0.290,    0.10063, 0.04049,  0.02184
  16,       2,             283,         141.5, 0.729, 0.025, 0.823, 38.636,  7.543,   1.491, 0.207,    0.05745, 0.02959,  0.01626
  19,       2,             164,          82.0, 0.729, 0.028, 0.838, 32.562,  4.905,   1.972, 0.268,    0.08077, 0.03107,  0.01325
  20,       2,             108,          54.0, 0.744, 0.040, 0.893, 36.646,  4.892,   1.838, 0.212,    0.08648, 0.02999,  0.00934
  23,       2,             190,          95.0, 0.724, 0.024, 0.837, 42.942,  4.479,   1.496, 0.149,    0.03574, 0.01548,  0.00679
  22,       2,              70,          35.0, 0.733, 0.032, 0.846, 32.916,  7.495,   1.658, 0.328,    0.09300, 0.04359,  0.00651
  25,       2,             126,          63.0, 0.729, 0.030, 0.845, 38.104,  5.418,   1.688, 0.412,    0.05001, 0.02465,  0.00630
  27,       2,              78,          39.0, 0.729, 0.028, 0.879, 38.699,  6.946,   1.706, 0.295,    0.07039, 0.03606,  0.00549
  21,       2,             124,          62.0, 0.723, 0.025, 0.839, 39.654,  5.799,   1.778, 0.445,    0.04381, 0.01844,  0.00543
  24,       2,              62,          31.0, 0.722, 0.021, 0.785, 30.308,  5.795,   1.793, 0.328,    0.08374, 0.04041,  0.00519
  28,       2,             167,          83.5, 0.721, 0.021, 0.807, 48.407,  7.357,   1.775, 0.365,    0.02777, 0.01459,  0.00464
  26,       2,              56,          28.0, 0.729, 0.024, 0.794, 37.675,  5.939,   1.920, 0.499,    0.06736, 0.02963,  0.00377
  29,       1,             107,          53.5, 0.740, 0.041, 0.869, 47.846,  7.581,   2.493, 0.582,    0.05826, 0.02884,  0.00312
  30,       2,              81,          40.5, 0.723, 0.024, 0.833, 42.941,  6.380,   1.638, 0.270,    0.03731, 0.01698,  0.00302
  32,       2,              24,          12.0, 0.731, 0.034, 0.859, 31.815,  5.463,   1.936, 0.279,    0.07723, 0.04639,  0.00185
  33,       1,              51,          25.5, 0.727, 0.027, 0.830, 35.091,  5.239,   1.888, 0.248,    0.06655, 0.02568,  0.00170
  31,       2,              25,          12.5, 0.726, 0.027, 0.791, 36.648,  7.667,   1.798, 0.282,    0.05442, 0.03124,  0.00136
  34,       2,              42,          21.0, 0.733, 0.032, 0.809, 51.968,  5.530,   1.897, 0.241,    0.02908, 0.01559,  0.00122
  35,       1,              71,          35.5, 0.727, 0.029, 0.851, 44.984,  4.667,   1.729, 0.241,    0.03072, 0.01398,  0.00109
  36,       2,              53,          26.5, 0.730, 0.022, 0.791, 56.496,  5.362,   1.978, 0.187,    0.01429, 0.00678,  0.00076
  37,       1,               7,           3.5, 0.740, 0.037, 0.817, 24.602,  2.893,   1.925, 0.229,    0.12791, 0.03646,  0.00045
  39,       1,              20,          10.0, 0.729, 0.030, 0.829, 57.450,  3.802,   2.015, 0.152,    0.01265, 0.00556,  0.00013
  40,       1,              10,           5.0, 0.722, 0.023, 0.781, 44.834,  4.217,   1.663, 0.181,    0.02496, 0.01209,  0.00012
  38,       1,               7,           3.5, 0.738, 0.035, 0.811, 43.389,  8.049,   1.594, 0.304,    0.03417, 0.02751,  0.00012
  41,       1,               2,           1.0, 0.786, 0.029, 0.815, 34.220,  0.749,   1.950, 0.151,    0.09245, 0.01844,  0.00009
  42,       1,               1,           0.5, 0.822, 0.000, 0.822, 36.017,  0.000,   1.599, 0.000,    0.11911, 0.00000,  0.00006
  43,       1,               3,           1.5, 0.744, 0.033, 0.791, 53.515,  2.099,   1.949, 0.072,    0.02013, 0.00376,  0.00003
  44,       1,               5,           2.5, 0.712, 0.020, 0.752, 57.384,  7.469,   2.004, 0.235,    0.01200, 0.00752,  0.00003
  45,       1,               2,           1.0, 0.727, 0.024, 0.751, 54.447,  4.805,   1.974, 0.185,    0.01396, 0.00760,  0.00001
```

**Figure 11 Initial supercluster statistics without additional geometry-based filtering applied.**

Importantly, membership of the particular tunnel from the particular simulation to the corresponding supercluster can be obtained from the *data/super_clusters/details/initial_super_cluster_details.txt* file. In the same folder *initial_super_cluster_events_details.txt* file also contains detailed information about the same membership of tunnels to superclusters extended by the information on a particular event assigned to superclusters (**Figure 12 – top**) as well as unassigned events (**Figure 12 – bottom**) which can be found in *data/super_cluster/details/outlier_transport_events_details.txt* file. We will describe the possible treatment of the unassigned event later in the next section. Analogously to the *statistics* folder, *'filtered'* equivalents to each file are available, nevertheless due to filtering turned off, containing exactly the same information.

```
Tunnel clusters and transport events assignment to superclusters (ordered by their priority):
-----------------------------------------------------------------------------------------------
Supercluster ID 1

Details on tunnel network:
Number of MD simulations = 2
Number of tunnel clusters = 4
Tunnel clusters:
from md1: 1, 5,
from md2: 1, 3,

Details on transport events:
Number of MD simulations = 1
Number of entry events = 2
Number of release events = 2
entry: (from Simulation: AQUA-DUCT ID, (Resname:Residue), start_frame->end_frame; ... )
from md2: 2, (WAT:294), 689->695; 3, (WAT:295), 714->715;
release: (from Simulation: AQUA-DUCT ID, (Resname:Residue), start_frame->end_frame; ... )
from md2: 1, (WAT:293), 722->726; 4, (WAT:296), 71->87;
-----------------------------------------------------------------------------------------------
Supercluster ID 2

Details on tunnel network:
Number of MD simulations = 2
Number of tunnel clusters = 6
Tunnel clusters:
from md1: 3, 22, 70, 93,
from md2: 2, 16,

Details on transport events:
Number of MD simulations = 0
Number of entry events = 0
Number of release events = 0
-----------------------------------------------------------------------------------------------
Supercluster ID 3

Details on tunnel network:
Number of MD simulations = 2
Number of tunnel clusters = 7
Tunnel clusters:
from md1: 2, 8, 30, 62,
from md2: 5, 22, 25,

Details on transport events:
Number of MD simulations = 1
Number of entry events = 2
Number of release events = 2
entry: (from Simulation: AQUA-DUCT ID, (Resname:Residue), start_frame->end_frame; ... )
from md1: 1, (WAT:326), 144->893; 4, (WAT:329), 880->902;
release: (from Simulation: AQUA-DUCT ID, (Resname:Residue), start_frame->end_frame; ... )
from md1: 1, (WAT:326), 908->965; 4, (WAT:329), 902->941;
-----------------------------------------------------------------------------------------------
.
.
.
-----------------------------------------------------------------------------------------------
Supercluster ID 8

Details on tunnel network:
Number of MD simulations = 2
Number of tunnel clusters = 7
Tunnel clusters:
from md1: 12, 21, 25,
from md2: 15, 23, 29, 34,

Details on transport events:
Number of MD simulations = 1
Number of entry events = 1
Number of release events = 1
entry: (from Simulation: AQUA-DUCT ID, (Resname:Residue), start_frame->end_frame; ... )
from md2: 5, (WAT:297), 943->957;
release: (from Simulation: AQUA-DUCT ID, (Resname:Residue), start_frame->end_frame; ... )
from md2: 2, (WAT:294), 698->704;
-----------------------------------------------------------------------------------------------
```

```
Unassigned transport events:
Number of entry events = 1
Number of release events = 0
entry: (from Simulation: AQUA-DUCT ID, (Resname:Residue), start_frame->end_frame; ... )
from md1: 6, (WAT:331), 497->588;
```

**Figure 12 Membership of particular tunnel clusters to superclusters together with detailed information on transport events assigned to the corresponding supercluster (top). Remaining unassigned events (bottom).**

##### 2.3.5. Filtering superclusters

One of the considerable advantages of tt_engine.py is its ability to restart the calculation from any desired stage. For filtering purposes, there is no need to start the calculation from scratch, we will directly start from the filtering stage (*start_from_stage = 6*). As we have already seen, analysis of a complete set of superclusters is complicated, and we may have difficulties spotting the most important details in the system, hence we will now employ a set of filters to focus the analysis on particularly interesting superclusters. Initially, change the settings of tunnel filtering based on minimum bottleneck radius – let it be at least 1.0 Å (*min_bottleneck_radius = 1.00*) and rerun the calculations. First of all, while running the calculations started from stage 6, you will notice information that stages 1-5 were skipped as presented in **Figure 13**.

During this stage, 45 initial superclusters are filtered applying minimum bottleneck radius filtering of tunnels, which retain only eight superclusters. This is followed by an unaltered procedure of processing AQUA-DUCT transport networks, their layering, and assignment to, in this case, eight filtered superclusters (**Figure 14** and **Figure 15**). Interestingly, all ten previously assigned events were assigned to a reduced set of superclusters (eight out of the initial 45), which indicates that while filtering (in this case focusing on wider superclusters), we have not lost any of the superclusters important for transport events. Additionally, one can visualize superclusters' volumes by changing *visualize_super_cluster_volumes* parameter to *True* from [**OUTPUT_SETTINGS**] section. To enable this kind of visualization you may need to install PyMCubes package first (simply run '*pip install --upgrade PyMCubes*' command in transport_tools conda environment). The result of such volume representation is shown in **Figure 14**.

```
bart@BIT3:tutorial$ tt_engine.py -c config.ini --overwrite
14:34:46 - transport_tools.libs.ui - INFO - ======== ********************************************* ========
14:34:46 - transport_tools.libs.ui - INFO - ========       TransportTools execution started        ========
14:34:46 - transport_tools.libs.ui - INFO - ======== ********************************************* ========
14:34:47 - __main__ - INFO - ------------------------------------------------------------------------------
14:34:47 - __main__ - INFO - STAGE 1 - Preparatory stage for unified analyses
14:34:47 - __main__ - INFO - ------------------------------------------------------------------------------
14:34:47 - __main__ - INFO - Requested to start from STAGE 6 => skipping STAGE 1
14:34:47 - __main__ - INFO - ------------------------------------------------------------------------------
14:34:47 - __main__ - INFO - STAGE 2 - Processing datasets of tunnel networks from CAVER results
14:34:47 - __main__ - INFO - ------------------------------------------------------------------------------
14:34:47 - __main__ - INFO - Requested to start from STAGE 6 => skipping STAGE 2
14:34:47 - __main__ - INFO - ------------------------------------------------------------------------------
14:34:47 - __main__ - INFO - STAGE 3 - Layering tunnel clusters to get their simplified representation
14:34:47 - __main__ - INFO - ------------------------------------------------------------------------------
14:34:47 - __main__ - INFO - Requested to start from STAGE 6 => skipping STAGE 3
14:34:47 - __main__ - INFO - ------------------------------------------------------------------------------
14:34:47 - __main__ - INFO - STAGE 4 - Computing distances among the layered clusters
14:34:47 - __main__ - INFO - ------------------------------------------------------------------------------
14:34:47 - __main__ - INFO - Requested to start from STAGE 6 => skipping STAGE 4
14:34:47 - __main__ - INFO - ------------------------------------------------------------------------------
14:34:47 - __main__ - INFO - STAGE 5 - Clustering the layered clusters into superclusters and creating initial out
puts
14:34:47 - __main__ - INFO - ------------------------------------------------------------------------------
14:34:47 - __main__ - INFO - Requested to start from STAGE 6 => skipping STAGE 5
```

**Figure 13 *TransportTools* reports skipping 1-5 stages and starts the calculations from stage 6 to not recompute initial stages unnecessarily.**

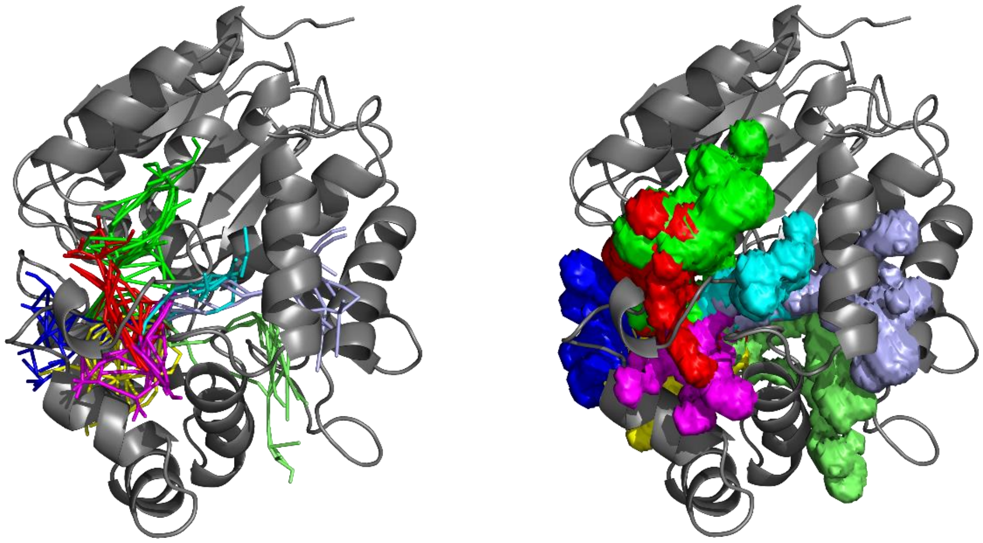

**Figure 14 Visualization of the filtered superclusters based on the minimum bottleneck radius ≥ 1.0 Å. Superclusters represented as lines (left) and as volumes (right).**

```
Statistics for: overall
Total simulations = 2
No snapshots per simulation = 1000

Parameters used for pre-selection of input tunnels for clustering:
length >= 5.00 A
radius >= 0.75 A
curvature <= 2.00

Active tunnel filters:
length = (0.00, 999.00) A
radius = (1.00, 999.00) A
curvature = (1.00, 999.00)
occurred in at least 0 simulations
occurred in at least 0 snapshots
occurred in at least 0.000 snapshots on average per simulation

SC_ID, No_Sims, Total_No_Frames, Avg_No_Frames, Avg_BR, StDev, Max_BR, Avg_Len, StDev, Avg_Cur, StDev, Avg_throug,   StDev, Priority
    1,       2,            1390,         695.0,  1.199, 0.141,  1.950,  13.911, 1.564,   1.236, 0.082,    0.56138, 0.06320,  0.78031
    3,       2,             392,         196.0,  1.124, 0.089,  1.515,  14.316, 1.918,   1.193, 0.071,    0.52584, 0.05406,  0.20613
    2,       2,             186,          93.0,  1.075, 0.067,  1.423,  15.416, 2.187,   1.342, 0.121,    0.51091, 0.05911,  0.09503
   11,       1,              36,          18.0,  1.095, 0.056,  1.239,  16.829, 1.505,   1.210, 0.054,    0.43010, 0.04071,  0.00774
    8,       2,              12,           6.0,  1.238, 0.224,  1.599,  13.510, 2.341,   1.276, 0.126,    0.56932, 0.10994,  0.00683
    4,       1,              12,           6.0,  1.043, 0.039,  1.157,  13.265, 1.053,   1.206, 0.089,    0.47947, 0.04534,  0.00288
    5,       1,               2,           1.0,  1.019, 0.012,  1.030,  21.187, 1.637,   1.253, 0.072,    0.34981, 0.04406,  0.00035
    6,       1,               1,           0.5,  1.029, 0.000,  1.029,  24.347, 0.000,   1.652, 0.000,    0.33560, 0.00000,  0.00017
```

**Figure 15 Statistics of the filtered superclusters based on the tunnel minimum bottleneck radius ≥ 1.0 Å.**

##### 2.3.6. Analyzing transport through superclusters

We had learned how to 'by property' filter input tunnels taken for *TransportTools* clustering before event assignment, which in the case of our system resulted in a reduced overall number of generated superclusters. Furthermore, an additional set of filters can be applied already after the event assignment, which enables to filter superclusters by a minimum number of entries, releases, or a total number of events. Analogously to filtering before event assignment, there is no need to rerun the whole procedure from scratch. As noted in chapter 2.3.3, the filtering of events is covered by stage 10, and we will rerun the analysis from this stage. Set *start_from_stage = 10,* apply filter defining a minimum number of total events in superclusters (*min_total_events* = *1*), and rerun calculations using the same command as previously (from the level of the *tutorial* folder).

*tt_engine.py -c config.ini --overwrite*

In the terminal, we could notice that stages 1-9 were skipped, as we requested to start from stage 10. As a report related to stage 10, *TransportTools* informed us that only two superclusters were kept after filtering performed after event assignment (**Figure 16**).

```
15:44:20 - __main__ - INFO - ------------------------------------------------------------------------------
15:44:20 - __main__ - INFO - STAGE 10 - Filtering supercluster with events and creating filtered outputs with even
ts
15:44:20 - __main__ - INFO - ------------------------------------------------------------------------------
15:44:20 - transport_tools.libs.tools - INFO - CHECKPOINT loaded from: results/_internal/checkpoints/stage009.dump
15:44:20 - transport_tools.libs.tools - INFO - Filtering 8 supercluster profiles using 4 parallel processes:
15:44:21 - transport_tools.libs.ui - INFO - |████████████████████████████████████████████████████████████| 100.0%
15:44:21 - transport_tools.libs.tools - INFO - 3 superclusters kept after filtering.
15:44:21 - transport_tools.libs.ui - INFO - Prioritization took: 0:00:00.
15:44:21 - transport_tools.libs.ui - INFO - Filtering took: 0:00:00.
15:44:21 - transport_tools.libs.tools - INFO - Saving visualization of superclusters.
15:44:29 - transport_tools.libs.ui - INFO - |████████████████████████████████████████████████████████████| 100.0%
15:44:29 - transport_tools.libs.ui - INFO - Visualization generation took: 0:00:08.
15:44:29 - transport_tools.libs.tools - INFO - Generating statistics for superclusters.
15:44:29 - transport_tools.libs.ui - INFO - Statistics generation took: 0:00:00.
15:44:29 - transport_tools.libs.tools - INFO - CHECKPOINT saved to: results/_internal/checkpoints/stage010.dump
15:44:29 - transport_tools.libs.ui - INFO - ======== ********************************************* ========
15:44:29 - transport_tools.libs.ui - INFO - ========        Overall elapsed time: 0:00:09         ========
15:44:29 - transport_tools.libs.ui - INFO - ======== ********************************************* ========
```

**Figure 16 Filtering of superclusters after events assignment (minimum number of total events ≥ 1, calculations started from stage 10).**

Entire *TransportTools* analysis applying an additional two-step filtering process (filtering before and after event assignment) reduces the number of superclusters taken into consideration gradually. The calculation process started with 170 original tunnels from two CAVER tunnel networks. These tunnels were merged into 45 superclusters (*statistics/1-initial_tunnels_statistics.txt*). Further, based on the minimum tunnel bottleneck radius, we reduced the total number of superclusters to eight (*statistics/2-filtered_tunnels_statistics.txt,* *statistics/3-initial_events_statistics.txt*). Finally, by defining additional criteria to search for the superclusters which feature at least one transport event, we have obtained three superclusters (*statistics/4-filtered_events_statistics.txt*). Now, we will have a look at the visualization and statistics of all three clusters. Go to the *results/visualization* folder and run the *4-visualize_filtered_events.py* script in PyMOL:

| *pymol 4-visualize_filtered_events.py* |
|---|

This should result in a visualization similar to **Figure 17**, if using volume visualization turned on, you will see also objects responsible for their representation (*cluster_001_vol*, *cluster_003_vol and cluster_008_vol*). Generated PyMOL session, besides volume-related objects, contains another objects (right site of the PyMOL window) – protein structure (*protein_structure*), first supercluster (blue, *cluster_001*) together with its events of entry and release (*entry_001, release_001*), third supercluster (green, *cluster_003*) with events (*entry_003, release_003*), and eighth supercluster (red, cluster_008) together with events visualizations (*entry_008, release_008*). Furthermore, an additional object is generated to represent an unassigned entry event (*entry_outlier)*. Based on statistics available in the *statistics/4-filtered_events_statistics.txt* file, we can learn the details about corresponding superclusters and correspondingly assigned events (**Figure 18**). Supercluster 1 was detected in both simulations, all together in 1390 frames. Its average bottleneck radius and length equals 1.199 Å and 13.911 Å, respectively. Four transport events were assigned to supercluster 1, accounting for two entries and two releases. Similarly, supercluster 3 was also detected in both MD trajectories but a significantly lower number of frames – 392. The average bottleneck is slightly narrower reaching 1.124 Å, while the average length is a little higher and equals 14.316 Å. In the case of supercluster 3, a total of four transport events were assigned, similarly to supercluster 1, composed of two entries and two releases. Finally, supercluster 8 occurred rarely (12 frames only) but was still detected in both simulations. It reached the average bottleneck radius and the average length of 1.238 and 13.510 Å, respectively. Two water transport events were assigned to this supercluster and accounted for one entry and one release.

Interestingly, we can observe that the *entry_outlier* event object visually overlaps with the volume of supercluster 3, besides not being assigned to any cluster. This is conditioned by the *event_assignment_cutoff* which defines the minimal buriedness of transport event along its trace in a supercluster to be assigned to that supercluster (by default set up to 0.85). By reducing the value of this parameter, *TransportTools* will be less strict during the assignment of events to particular superclusters and the chance of a particular event to be assigned will be increased. In our case, as the *entry_outlier* event substantially overlaps with supercluster 3, reducing the *event_assignment_cutoff* to 0.80 results in the successful assignment of all events to three superclusters (001, 003, and 008). Note, that in such a case it is sufficient to rerun the analysis just from stage 9, in which events assignment is performed.

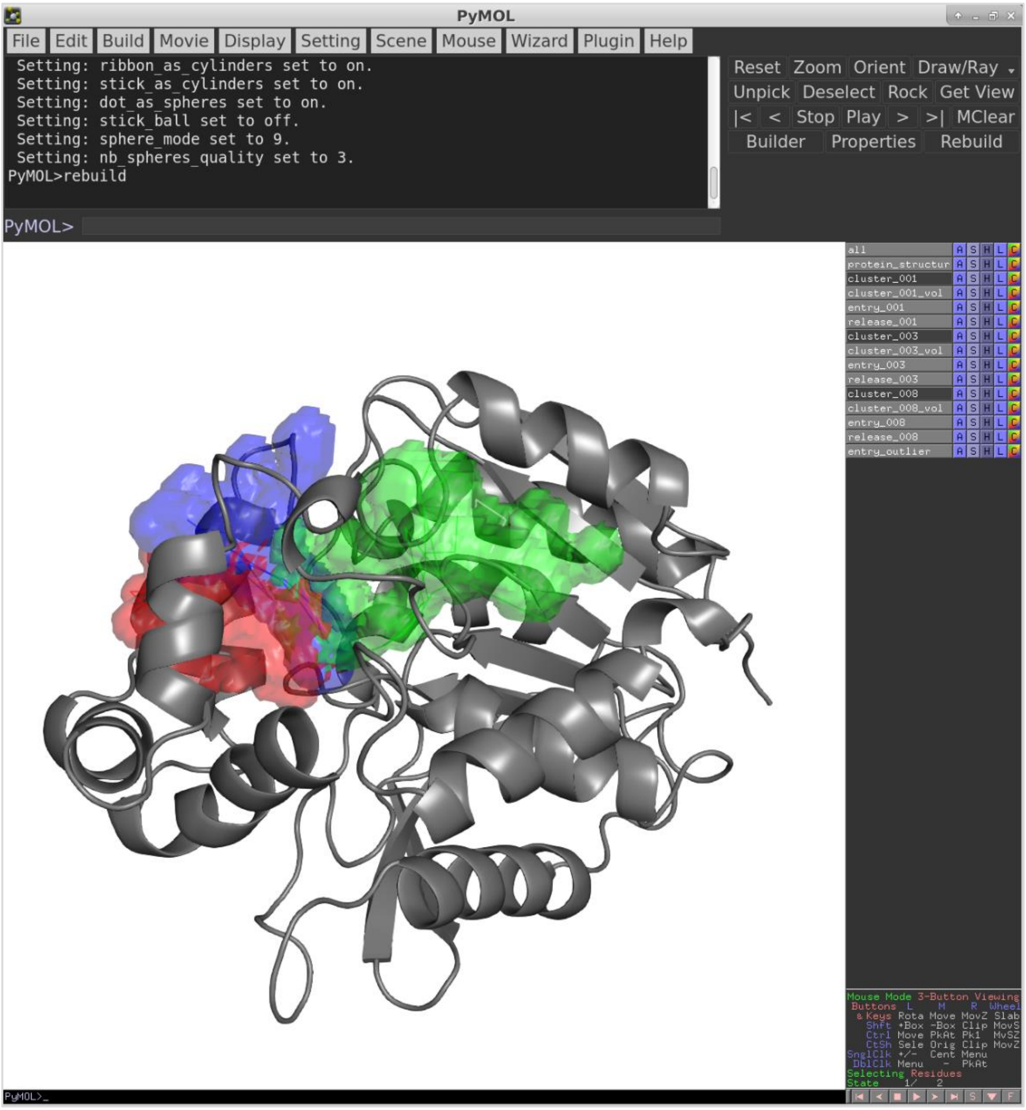

**Figure 17 Visualization of the superclusters and transport events filtered after event assignment based on the minimum number of total events ≥ 1.**

```
Statistics for: overall
Total simulations = 2
No snapshots per simulation = 1000

Parameters used for pre-selection of input tunnels for clustering:
length >= 5.00 A
radius >= 0.75 A
curvature <= 2.00

Active tunnel filters:
length = (0.00, 999.00) A
radius = (1.00, 999.00) A
curvature = (1.00, 999.00)
occurred in at least 0 simulations
occurred in at least 0 snapshots
occurred in at least 0.000 snapshots on average per simulation
has at least 1 transport events
has at least 0 entry events
has at least 0 release events

SC_ID, No_Sims, Total_No_Frames, Avg_No_Frames, Avg_BR, StDev, Max_BR, Avg_Len, StDev, Avg_Cur, StDev, Avg_throug,   StDev, Priority, Num_Events, Num_entries, Num_releases
    1,       2,            1390,         695.0,  1.199, 0.141,  1.950,  13.911, 1.564,   1.236, 0.082,    0.56138, 0.06320,  0.78031,          4,           2,            2
    3,       2,             392,         196.0,  1.124, 0.089,  1.515,  14.316, 1.918,   1.193, 0.071,    0.52584, 0.05406,  0.20613,          4,           2,            2
    8,       2,              12,           6.0,  1.238, 0.224,  1.599,  13.510, 2.341,   1.276, 0.126,    0.56932, 0.10994,  0.00683,          2,           1,            1
Total number of unassigned events:                                                                                                                    1,           1,            0
```

**Figure 18 Filtered superclusters statistics based on the minimum number of total events ≥ 1.**

#### 2.3.7. Advanced settings and comparative analysis

This chapter represents the last part of the tutorial and will cover more advanced functionalities of *TransportTools* engine. For clarity, we will run new tt_engine.py calculations and we will save the results in the output folder named differently not to overwrite currently generated data so that we can optionally explore and compare the differences between the standard and comparative analyses. We will start with the generation of the configuration file extended with the **[ADVANCED_SETTINGS]** section. Enter the *tutorial* folder and run the following command to produce such a configuration file:

```
tt_engine.py -wa
```

This will generate new *tmp_config.ini*. Now, using terminal, please rename it to *config_advanced.ini:*

```
mv tmp_config.ini config_advanced.ini
```

As we can notice, the newly created file has default settings for all parameters, so we have to update ones that are interesting for planned comparative analysis. The **[INPUT_PATHS]** parameters have to be specified analogously to the previous run. Further, we use the same setting for parsing of CAVER tunnel clusters:

- *snapshots_per_simulation = 1000, caver_traj_offset = 1, snapshot_id_position = 1, snapshot_delimiter = .* and *process_bottleneck_residues = True*
- We will apply a filter of minimum tunnel bottleneck radius to 1 Å (*min_bottleneck_radius = 1.00*) from the beginning.
- We will use '*exact_matching*' for *ambiguous_event_assignment_resolution* parameter, and we will not apply any filters after the event assignment.
- Additionally, in **[OUTPUT_SETTINGS]** section, we change the name of the output folder to *'results_comparative' (output_path = results_comparative)*.

Finally, in the newly created **[ADVANCED_SETTINGS]** section, we will ask *TransportTools* to perform comparative analysis (*perform_comparative_analysis = True*), which requires a definition of groups for comparison (*comparative_groups_definition = traj1: [md1]; traj2: [md2]*) – the first group will be called *traj1* and will cover md1 trajectory, while the second will be called *traj2* and will include remaining md2 trajectory data. Note that the names of groups cannot overlap with the names of particular simulation folders. Analogously to standard analysis, the User can use *visualize_comparative_super_cluster_volumes* parameter to represent superclusters as volumes. At this stage of analysis, we will leave it as default (*False*).

Optionally, we can ask *TransportTools* to overwrite output automatically, so an additional '*--overwrite*' flag will not be necessary during program execution (*overwrite = True*). Another parameter that might be helpful to adjust relates to the PyMOL version we are currently using. For older PyMOL based on Python2, it is necessary to have the support of legacy versions enabled (*legacy_pymol_support = True*) to generate compatible data used by the visualization scripts. Furthermore, *TransportTools* enables the selection of engines for the processing of trajectories. This is controlled by *trajectory_engine* in [**ADVANCED SETTINGS**] section and two possible options are available at present: *mdtraj* and *pytraj*. *mdtraj* is the default option in case *pytraj* is not detected in the currently used environment. Otherwise, *pytraj* is automatically selected. Selection of the engine does not affect the *TransportTools* results *per se*, nevertheless please note that it might affect the runtimes, being faster for *pytraj* in particular with some trajectory formats like netcdf. On the other hand, *mdtraj* although slightly slower*,* has the advantage of being automatically installed with *TransportTools,* without additional system requirements necessary for *pytraj* (to install follow instructions). For the purpose of this tutorial, we will use default *mdtraj* for simplicity. The complete configuration file settings are presented in **Figure 19**.

With a properly prepared advanced configuration file, we can proceed with the next *tt_engine.py* run. Analogously to previous executions, from the *tutorial* folder (we should have the *config_advanced.ini* file in this location), run the following command (note that the '*--overwrite*' option is not necessary anymore due to changes in the **[ADVANCED_SETTINGS]** section as mentioned above):

tt_engine.py -c config_advanced.ini

```
[INPUT_PATHS]
# CAVER results
caver_results_path = precomputed_inputs
caver_results_folder_pattern = *
caver_results_relative_subfolder_path = caver
# AQUA-DUCT results
aquaduct_results_path = precomputed_inputs
aquaduct_results_folder_pattern = *
aquaduct_results_relative_tarfile = aquaduct/6_visualize_results.tar.gz
aquaduct_results_relative_summaryfile = aquaduct/5_analysis_results.txt
# Source MD trajectories
trajectory_path = simulations
trajectory_folder_pattern = *
trajectory_relative_file = trajectory.nc
topology_relative_file = structure.parm7

[CALCULATIONS_SETTINGS]
start_from_stage = 1
stop_after_stage = None
num_cpus = 4
pdb_reference_structure = None
layer_thickness = 1.50
# Parsing of tunnel clusters from CAVER results
snapshots_per_simulation = 1000
caver_traj_offset = 1
snapshot_id_position = 1
snapshot_delimiter = .
process_bottleneck_residues = True
# Clustering of tunnel clusters into superclusters
min_tunnel_radius4clustering = 0.75
min_tunnel_length4clustering = 5.00
max_tunnel_curvature4clustering = 2.00
clustering_linkage = complete
clustering_cutoff = 2.00
# Filters applied on superclusters before event assignment (-1 => inactive filter)
min_length = -1.00
max_length = -1.00
min_bottleneck_radius = 1.00
max_bottleneck_radius = -1.00
min_curvature = -1.00
max_curvature = -1.00
min_sims_num = -1
min_snapshots_num = -1
min_avg_snapshots_num = -1.00
# Processing of transport events from AQUA-DUCT results, and their assignment to superclusters
event_min_distance = 6.00
event_assignment_cutoff = 0.85
ambiguous_event_assignment_resolution = exact_matching
# Additional filters applied on superclusters after event assignment (-1 => inactive filter)
min_total_events = -1
min_entry_events = -1
min_release_events = -1

[OUTPUT_SETTINGS]
output_path = results_comparative
# Optional data generation
save_super_cluster_profiles_csvs = False
save_distance_matrix_csv = False
# Optional visualization
visualize_super_cluster_volumes = True
visualize_transformed_tunnels = False
visualize_transformed_transport_events = False

[ADVANCED_SETTINGS]
# Calculations
random_seed = 4
directional_cutoff = 1.57
aqauduct_ligand_effective_radius = 1.00
perform_exact_matching_analysis = False
perform_comparative_analysis = True
visualize_comparative_super_cluster_volumes = False
comparative_groups_definition = traj1: [md1]; traj2: [md2]
overwrite = True
# Finer control of outputs & logging
visualize_exact_matching_outcomes = False
visualize_layered_clusters = False
visualize_layered_events = False
legacy_pymol_support = True
trajectory_engine = mdtraj
logfilename = transport_tools.log
log_level = info
max_events_per_cluster4visualization = 1000
```

**Figure 19 Adjusted parameters of the advanced configuration file.**

Comparative analysis performed by *TransportTools* does not differ in terms of the calculations process or the information printed in the terminal. Nevertheless, this functionality enables generating additional statistics files and visualization scripts containing information for each of the specified groups independently, which provides an opportunity to compare/contrast these groups. **Figure 20** illustrates the newly created *results_comparative* folder.

As we can see, *statistics* and *visualization* folders contain an additional subfolder called *comparative_analysis*. In the *statistics/comparative_analysis* folder, we can find merged statistics files containing statistics for each group in a separate section. Subfolders *traj1* and *traj2* contain the same type of file, although covering information about a particular group only. Analogously, the *visualization/comparative_analysis* subfolder contains separate subfolders for *traj1* and *traj2* groups with visualization scripts to visualize only a particular group.

```
results_comparative
├── data
│   └── super_clusters
├── _internal
│   ├── checkpoints
│   ├── clustering
│   ├── layered_data
│   ├── network_data
│   ├── super_cluster_pathsets
│   ├── super_cluster_profiles
│   └── transformations
├── statistics
│   ├── 1-initial_tunnels_statistics_bottleneck_residues.txt
│   ├── 1-initial_tunnels_statistics.txt
│   ├── 2-filtered_tunnels_statistics_bottleneck_residues.txt
│   ├── 2-filtered_tunnels_statistics.txt
│   ├── 3-initial_events_statistics_bottleneck_residues.txt
│   ├── 3-initial_events_statistics.txt
│   ├── 4-filtered_events_statistics_bottleneck_residues.txt
│   ├── 4-filtered_events_statistics.txt
│   └── comparative_analysis
├── transport_tools.log
└── visualization
    ├── 1-visualize_initial_tunnels.py
    ├── 2-visualize_filtered_tunnels.py
    ├── 3-visualize_initial_events.py
    ├── 4-visualize_filtered_events.py
    ├── comparative_analysis
    └── sources
```

```
results_comparative/statistics/comparative_analysis/
├── 1-initial_tunnels_statistics_bottleneck_residues.txt
├── 1-initial_tunnels_statistics.txt
├── 2-filtered_tunnels_statistics_bottleneck_residues.txt
├── 2-filtered_tunnels_statistics.txt
├── 3-initial_events_statistics_bottleneck_residues.txt
├── 3-initial_events_statistics.txt
├── 4-filtered_events_statistics_bottleneck_residues.txt
├── 4-filtered_events_statistics.txt
├── traj1
│   ├── 1-initial_tunnels_statistics_bottleneck_residues.txt
│   ├── 1-initial_tunnels_statistics.txt
│   ├── 2-filtered_tunnels_statistics_bottleneck_residues.txt
│   ├── 2-filtered_tunnels_statistics.txt
│   ├── 3-initial_events_statistics_bottleneck_residues.txt
│   ├── 3-initial_events_statistics.txt
│   ├── 4-filtered_events_statistics_bottleneck_residues.txt
│   └── 4-filtered_events_statistics.txt
└── traj2

results_comparative/visualization/comparative_analysis/
├── traj1
│   ├── 1-visualize_initial_tunnels.py
│   ├── 2-visualize_filtered_tunnels.py
│   ├── 3-visualize_initial_events.py
│   └── 4-visualize_filtered_events.py
└── traj2
```

**Figure 20 Tree of *results_comparative* folder representing the structure of outputs generated by comparative analysis performed by *TransportTools*.**

Interestingly, such a comparative study for two trajectories analyzed in this tutorial elucidates differences between superclusters occurrence among them. The first trajectory (md1) contains representatives for seven out of eight superclusters (obtained with filtering of tunnels clusters by the minimum bottleneck radius), while the second trajectory contains only five out of them.

Based on the *results_comparative/statistics/comparative_analysis/2-filtered_tunnels_statistics.txt* file (**Figure 21**), one can notice that *md1* trajectory is missing representative of supercluster 11, while *md2* is missing representatives for superclusters 4, 5, and 6 but has a representative of cluster 11 missing in md1. On the other hand, both simulations contain representative for superclusters 1, 2, 3, and 8 which was already seen in **Figure 15**, indicated by standard (non-comparative) analysis – both superclusters were observed in two simulations (No_sims).

Further, using scripts from newly generated *visualization/comparative_analysis/traj1* and *visualization/comparative_analysis/traj2* folders (**Figure 22**) we can visualize supercluster for each group separately – seven superclusters for *traj1*, five superclusters for *traj2*.

```
Statistics for: traj1 ['md1']
Total simulations = 1
No snapshots per simulation = 1000

Parameters used for pre-selection of input tunnels for clustering:
length >= 5.00 A
radius >= 0.75 A
curvature <= 2.00

Active tunnel filters:
length = (0.00, 999.00) A
radius = (1.00, 999.00) A
curvature = (1.00, 999.00)
occurred in at least 0 simulations
occurred in at least 0 snapshots
occurred in at least 0.000 snapshots on average per simulation

SC_ID, No_Sims, Total_No_Frames, Avg_No_Frames, Avg_BR, StDev, Max_BR, Avg_Len, StDev, Avg_Cur, StDev, Avg_throug,   StDev, Priority
    1,       1,             639,         639.0,  1.176, 0.126,  1.720,  14.096, 1.475,   1.242, 0.090,    0.54233, 0.06062,  0.34655
    3,       1,             387,         387.0,  1.125, 0.090,  1.515,  14.281, 1.903,   1.191, 0.067,    0.52741, 0.05240,  0.20411
    2,       1,             116,         116.0,  1.084, 0.074,  1.423,  15.434, 2.415,   1.345, 0.126,    0.52655, 0.05930,  0.06108
   11,      -,               -,             -,      -,     -,      -,       -,     -,       -,     -,          -,       -,        -
    8,       1,              11,          11.0,  1.251, 0.229,  1.599,  13.330, 2.364,   1.274, 0.131,    0.57271, 0.11423,  0.00630
    4,       1,              12,          12.0,  1.043, 0.039,  1.157,  13.265, 1.053,   1.206, 0.089,    0.47947, 0.04534,  0.00575
    5,       1,               2,           2.0,  1.019, 0.012,  1.030,  21.187, 1.637,   1.253, 0.072,    0.34981, 0.04406,  0.00070
    6,       1,               1,           1.0,  1.029, 0.000,  1.029,  24.347, 0.000,   1.652, 0.000,    0.33560, 0.00000,  0.00034
-------------------------------------------------------------------------------------------------------------------------------------------

Statistics for: traj2 ['md2']
Total simulations = 1
No snapshots per simulation = 1000

Parameters used for pre-selection of input tunnels for clustering:
length >= 5.00 A
radius >= 0.75 A
curvature <= 2.00

Active tunnel filters:
length = (0.00, 999.00) A
radius = (1.00, 999.00) A
curvature = (1.00, 999.00)
occurred in at least 0 simulations
occurred in at least 0 snapshots
occurred in at least 0.000 snapshots on average per simulation

SC_ID, No_Sims, Total_No_Frames, Avg_No_Frames, Avg_BR, StDev, Max_BR, Avg_Len, StDev, Avg_Cur, StDev, Avg_throug,   StDev, Priority
    1,       1,             751,         751.0,  1.219, 0.149,  1.950,  13.753, 1.620,   1.232, 0.074,    0.57758, 0.06079,  0.43376
    3,       1,               5,           5.0,  1.078, 0.041,  1.136,  17.059, 0.695,   1.377, 0.101,    0.40423, 0.04027,  0.00202
    2,       1,              70,          70.0,  1.060, 0.048,  1.201,  15.386, 1.745,   1.336, 0.110,    0.48500, 0.04878,  0.03395
   11,       1,              36,          36.0,  1.095, 0.056,  1.239,  16.829, 1.505,   1.210, 0.054,    0.43010, 0.04071,  0.01548
    8,       1,               1,           1.0,  1.088, 0.000,  1.088,  15.492, 0.000,   1.297, 0.000,    0.53213, 0.00000,  0.00053
    4,      -,               -,             -,      -,     -,      -,       -,     -,       -,     -,          -,       -,        -
    5,      -,               -,             -,      -,     -,      -,       -,     -,       -,     -,          -,       -,        -
    6,      -,               -,             -,      -,     -,      -,       -,     -,       -,     -,          -,       -,        -
-------------------------------------------------------------------------------------------------------------------------------------------
```

**Figure 21 Comparative statistics generated for each group separately.**

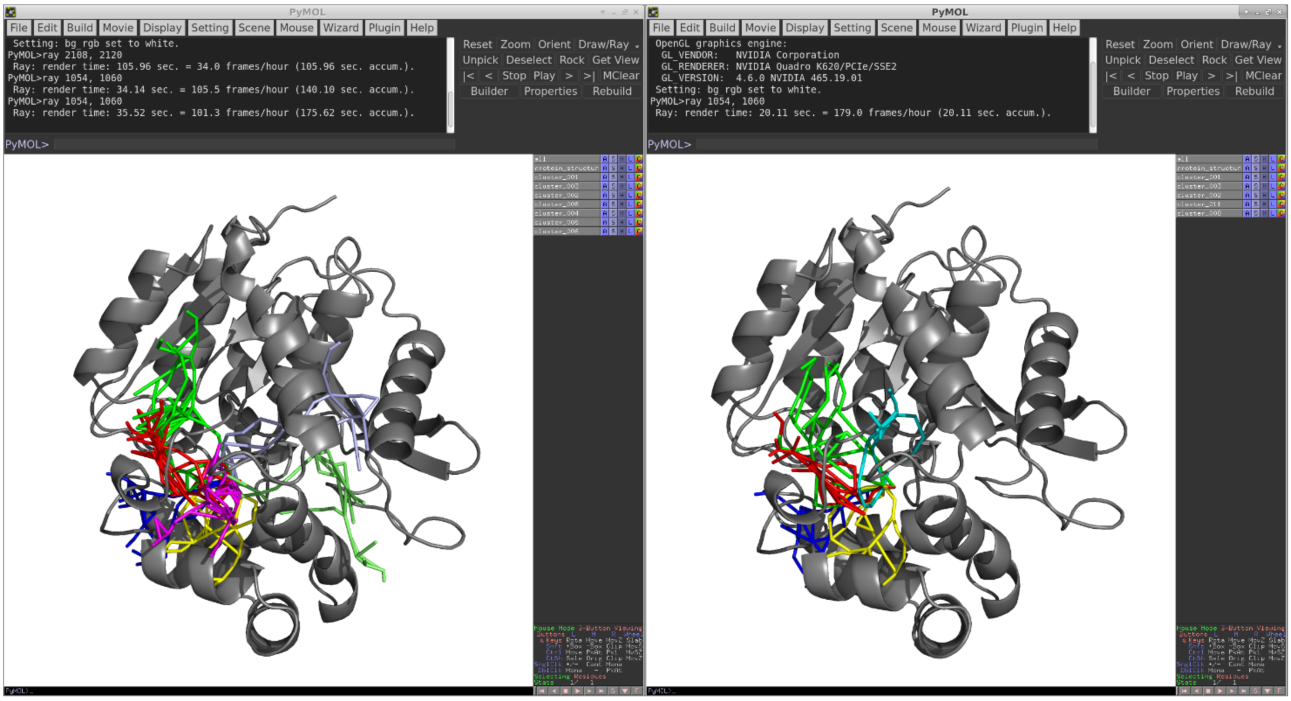

**Figure 22 Visualization of superclusters for *traj1* (left) and *traj2* (right) groups defined for comparative analysis.**

##### 2.3.8. Introduction to custom analysis with *TransportTools library*

Aside from the access to general statistics and visualization of entire superclusters from the massive analysis of transport pathways, it is often desirable to inspect and visualize different details. Hence, among standard outputs including statistics, visualization scripts and other data, we provide an additional script to help develop your own customized analyzes based on *TransportTools* results. This chapter will focus on the description of several steps which will enrich standard *TransportTools* workflow outcomes.

We will start with copying *custom_analysis.py* script from *custom_analysis* folder to the location of the *config.ini*. From the level of *tutorial* folder type in terminal:

```
cp custom_analysis/custom_analysis.py .
```

Next, open the script using the preferred text editor and have a look at the content of the file. In line 36, one can find the configuration parameter which has to correspond to the actual configuration file of the *TransportTools* calculations to be analyzed. For the purpose of the tutorial, we will perform a custom analysis of the results generated in chapters 2.3.1 - 2.3.6, hence the configuration file of interest is *config.ini*, as shown in **Figure 23**. This will enable us to load the output files from the previous calculations and initialize the environment for the custom analysis.

```
36  config = AnalysisConfig("config.ini")
37
38  # load outcomes from tt_engine run
39  mol_system = tools.load_checkpoint(os.path.join(config.get_parameter("output_path"), "_internal", "checkpoints",
40                                                  "stage010.dump"))
41
```

**Figure 23 Initialization of the *custom_analysis.py* script with previously calculated data.**

The following part of the script (lines 42-71) is responsible for the enumeration of the tunnels within specified length (line 44) and tunnel radius (line 63) criteria across selected superclusters (line 45). As presented in **Figure 24**, we will consider the tunnels in all superclusters which meet the criteria of minimum bottleneck radius of 1.4 Å and the minimum length of 10 Å. Tunnels specified by mentioned filters will be counted during the execution of the script and returned in a form of a simplified per-supercluster summary in the terminal while the time-evolution data for all tunnels longer than 10 Å will be stored as separate text files in the *custom_analysis/time_evolution_data* folder. We will come back to this aspect later.

```
42  # how many open tunnels (radius > 1.4 A)  with minimal length of 10 A
43  os.makedirs(os.path.join("custom_analysis", "time_evolution_data"), exist_ok=True)
44  filters = tools.define_filters(min_length=10)
45  super_cluster_id = None  # analyze all superclusters
46  parameter = "bottleneck_radius"
47  dataset = mol_system.get_property_time_evolution_data(parameter, active_filters=filters,
48                                                        sc_id=super_cluster_id)
49  for sc_id, sc_data in dataset.items():
50      sc_overall_num_frames = len(sc_data.keys()) * config.get_parameter("snapshots_per_simulation")
51      sc_overall_data = None
52      for md_label, md_data in sc_data.items():  # per source MD simulation data
53          if sc_overall_data is None:
54              sc_overall_data = md_data
55          else:
56              sc_overall_data = np.concatenate((sc_overall_data, md_data))
57          # saving data separately per MD simulation, can be later concatenated if a single set is desired
58          with open(os.path.join("custom_analysis", "time_evolution_data",
59                                 "{}_{}_sc{}.txt".format(parameter, md_label, sc_id)), "w") as out_stream:
60              for i, radii in enumerate(md_data):
61                  out_stream.write("{:6d} {:6.2f}\n".format(i + 1, radii))
62
63      open_tunnels = (sc_overall_data >= 1.4)  # tunnels with min radius of 1.4 A
64      max_value = np.max(sc_overall_data)
65      how_many_open_tunnels = np.count_nonzero(open_tunnels)
66      percent_open = how_many_open_tunnels * 100 / sc_overall_num_frames
67      if how_many_open_tunnels > 0:
68          logger.info("In supercluster {}:".format(sc_id))
69          logger.info("{} of open tunnels {:.3f}%".format(how_many_open_tunnels, percent_open))
70          logger.info("{:.3f} max radius".format(max_value))
71          logger.info("-----------------------------------")
```

**Figure 24 Search of tunnels in all superclusters within specified minimum bottleneck radius and tunnel length.**

Remaining part of the script consist of three subsections responsible for:

1. (Lines 73-82) – Visualization of all open tunnels from particular parts of the trajectory satisfying specified geometry criteria, with single reference PDB structure. Parameters controlling visualization at this stage are (**Figure 25**):
   - bottleneck radius and minimum tunnel length (line 76), use 1.4 Å and 10 Å, respectively
   - supercluster (line 77), limit the visualization to supercluster 1 only
   - MD trajectories (line 78), take both simulations into account
   - range of snapshots of interest (lines 79-80), for current analysis, limit the visualization to part of the simulations from 100$^{th}$ to 200$^{th}$ frame

```
73  # visualizing open tunnels (radius > 1.4 A)  with minimal length of 10 A from supercluster 1, snapshots 100-200,
74  # with single reference pdb structure
75  visualization_output_folder = os.path.join("custom_analysis", "vis_details", "static")
76  filters = tools.define_filters(min_length=10, min_bottleneck_radius=1.4)
77  super_cluster_id = 1
78  md_labels = None  # all trajectories
79  start_snapshot = 100
80  end_snapshot = 200
81  mol_system.show_tunnels_passing_filter(super_cluster_id, filters, visualization_output_folder, md_labels=md_labels,
82                                         start_snapshot=start_snapshot, end_snapshot=end_snapshot, trajectory=False)
83
```

**Figure 25 Visualization of open tunnels from the particular part of a trajectory with a single reference PDB structure.**

2. (Lines 84-93) – Visualization of all open tunnels from particular parts of the trajectory satisfying specified geometry criteria, with corresponding protein ensemble. Parameters controlling visualization at this stage are analogous to previous functionality and cover (**Figure 26**):
   - bottleneck radius and minimum tunnel length (line 87), use 1.4 Å and 10 Å, respectively
   - supercluster (line 88), limit the visualization to supercluster 1 only
   - MD trajectories (line 89), select only md1 trajectory for visualization
   - range of snapshots of interest (lines 90-91), for current analysis, limit the visualization to part of the simulations from 100$^{th}$ to 200$^{th}$ frame

```
84  # visualizing open tunnels (radius > 1.4 A)  with minimal length of 10 A from supercluster 1,  MD simulation
85  # md1 snapshots 100-200, including corresponding protein ensembles in pdb files
86  visualization_output_folder = os.path.join("custom_analysis", "vis_details", "dynamics")
87  filters = tools.define_filters(min_length=10, min_bottleneck_radius=1.4)
88  super_cluster_id = 1
89  md_labels = ["md1"]
90  start_snapshot = 100
91  end_snapshot = 200
92  mol_system.show_tunnels_passing_filter(super_cluster_id, filters, visualization_output_folder, md_labels=md_labels,
93                                         start_snapshot=start_snapshot, end_snapshot=end_snapshot, trajectory=True)
94
```

**Figure 26 Visualization of open tunnels from the particular part of a trajectory including a corresponding ensemble of protein structures.**

3. (Lines 95-102) – Visualization of the particular tunnel from selected supercluster and trajectory. Parameters controlling visualization at this stage are (**Figure 27**):
   - bottleneck radius and minimum tunnel length (line 97), use 1.4 Å and 10 Å, respectively
   - supercluster (line 98), select supercluster 1 for visualization
   - MD trajectories (line 99), select only md1 trajectory for visualization
   - snapshot (lines 100), for current analysis select tunnels from snapshot #120 for visualization

```
 95  # visualize particular tunnel from supercluster 1, MD simulation md1, snapshot 120
 96  visualization_output_folder = os.path.join("custom_analysis", "vis_details", "1snapshot")
 97  filters = tools.define_filters(min_length=10, min_bottleneck_radius=1.4)
 98  super_cluster_id = 1
 99  md_labels = ["md1"]
100  start_snapshot = end_snapshot = 120
101  mol_system.show_tunnels_passing_filter(super_cluster_id, filters, visualization_output_folder, md_labels=md_labels,
102                                         start_snapshot=start_snapshot, end_snapshot=end_snapshot, trajectory=True)
103
```

**Figure 27 Visualization of the particular tunnel from the selected supercluster and trajectory frame with the corresponding PDB structure.**

All parameters for the custom analysis are adjusted, hence we continue and execute the script from *tutorial* folder using the command:

| *python custom_analysis.py* |
|---|

This will result in the following prompt in the terminal (**Figure 28**):

```
bart@BIT3:tutorial$ ./custom_analysis.py
16:52:20 - transport_tools.libs.ui - INFO - ======== ********************************************* ========
16:52:20 - transport_tools.libs.ui - INFO - ========        TransportTools execution started         ========
16:52:20 - transport_tools.libs.ui - INFO - ======== ********************************************* ========
16:52:20 - transport_tools.libs.tools - INFO - CHECKPOINT loaded from: results/_internal/checkpoints/stage010.dump
16:52:21 - __main__ - INFO - In supercluster 2:
16:52:21 - __main__ - INFO - 1 of open tunnels 0.050%
16:52:21 - __main__ - INFO - 1.423 max radius
16:52:21 - __main__ - INFO - -----------------------------------
16:52:21 - __main__ - INFO - In supercluster 3:
16:52:21 - __main__ - INFO - 6 of open tunnels 0.300%
16:52:21 - __main__ - INFO - 1.515 max radius
16:52:21 - __main__ - INFO - -----------------------------------
16:52:21 - __main__ - INFO - In supercluster 8:
16:52:21 - __main__ - INFO - 4 of open tunnels 0.200%
16:52:21 - __main__ - INFO - 1.599 max radius
16:52:21 - __main__ - INFO - -----------------------------------
16:52:21 - __main__ - INFO - In supercluster 1:
16:52:21 - __main__ - INFO - 115 of open tunnels 5.750%
16:52:21 - __main__ - INFO - 1.746 max radius
16:52:21 - __main__ - INFO - -----------------------------------
16:52:21 - transport_tools.libs.tools - INFO - Preparing visualization of tunnels from supercluster 1 present betw
een snapshots 100 and 200.
16:52:21 - transport_tools.libs.tools - INFO - Prepared visualization will be available in custom_analysis/vis_det
ails/static.
16:52:21 - transport_tools.libs.tools - INFO - Saving visualization for 3 tunnel(s) from md1 Cluster_1.
16:52:21 - transport_tools.libs.tools - INFO - Saving visualization for 7 tunnel(s) from md2 Cluster_1.
16:52:21 - transport_tools.libs.tools - INFO - Preparing visualization of tunnels from supercluster 1 present betw
een snapshots 100 and 200.
16:52:21 - transport_tools.libs.tools - INFO - Prepared visualization will be available in custom_analysis/vis_det
ails/dynamics.
16:52:21 - transport_tools.libs.tools - INFO - Saving visualization for 3 tunnel(s) from md1 Cluster_1.
16:52:33 - transport_tools.libs.tools - INFO - Preparing visualization of tunnels from supercluster 1 present betw
een snapshots 120 and 120.
16:52:33 - transport_tools.libs.tools - INFO - Prepared visualization will be available in custom_analysis/vis_det
ails/1snapshot.
16:52:34 - transport_tools.libs.tools - INFO - Saving visualization for 1 tunnel(s) from md1 Cluster_1.
16:52:34 - transport_tools.libs.ui - INFO - ======== ********************************************* ========
16:52:34 - transport_tools.libs.ui - INFO - ========         Overall elapsed time: 0:00:14           ========
16:52:34 - transport_tools.libs.ui - INFO - ======== ********************************************* ========
```

**Figure 28 Execution of *custom_analysis.py* run and its standard output.**

The pre-supercluster summary describing the count and percentage of open tunnels is provided in the standard output according to specified geometry filters. In supercluster 2, only 1 tunnel met the 1.4 Å and 10 Å criteria for minimum bottleneck radius and minimum length, respectively, which represents 0.05 % of tunnels in entire supercluster 2. 6 tunnels were found in supercluster 3, representing 0.3 % of tunnels in that supercluster. Further, 4 tunnels were found in supercluster 8, corresponding to 0.2 % of all tunnels in this supercluster and finally, in supercluster 1, using specified criteria, 115 tunnels were found representing 5.75 % of tunnels this supercluster.

Additionally, in *custom_analysis* folder, two subfolders were created: (i) *time_evolution_data* containing a change of the bottleneck radius in time for each supercluster in all simulations, and (ii) *vis_details* with folders *static* (tunnels + reference structure), *dynamics* (tunnels + protein ensemble) and *1snapshot* (tunnels from particular snapshot) which contain scripts for visualization of requested details in PyMOL. To better illustrate the usefulness of such customized analysis, we can use the generated data to plot (using your favorite tool) the time-evolution of bottleneck radius in the supercluster 1 in simulation md1 (**Figure 29** left panel) together with the visualization of the tunnel and protein structure in snapshot #120 from the same supercluster and simulation (**Figure 29** right panel), hence one can observe that in this particular snapshot the tunnel opens above-requested cutoff and benefiting from *custom_analysis.py* we have an opportunity to visualize and better understand the molecular grounds of such behavior (residues flips, backbone motions, etc.). Moreover, in **Figure 30** two PyMOL sessions are shown, which were prepared using visualization scripts from *static* (left) and *dynamics* (right) folders containing details requested according to the code in *custom_analysis.py* shown in **Figure 25** and **Figure 26**.

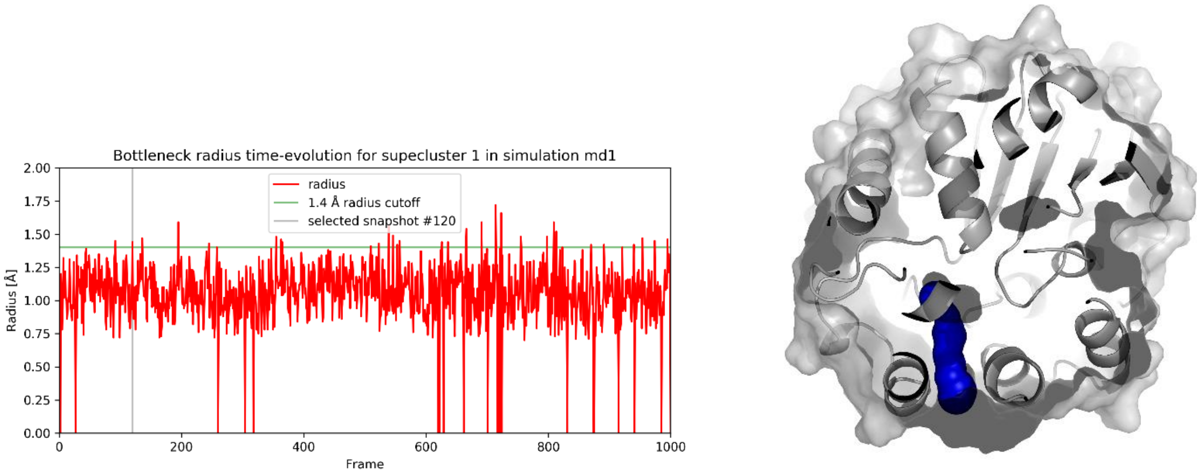

**Figure 29 Time evolution of the bottleneck radius for supercluster 1 in simulation md1 (left), horizontal green line depicts the bottleneck radius of 1.4 Å filtering criteria, while gray vertical line indicates snapshot #120 selected for visualization (right).**

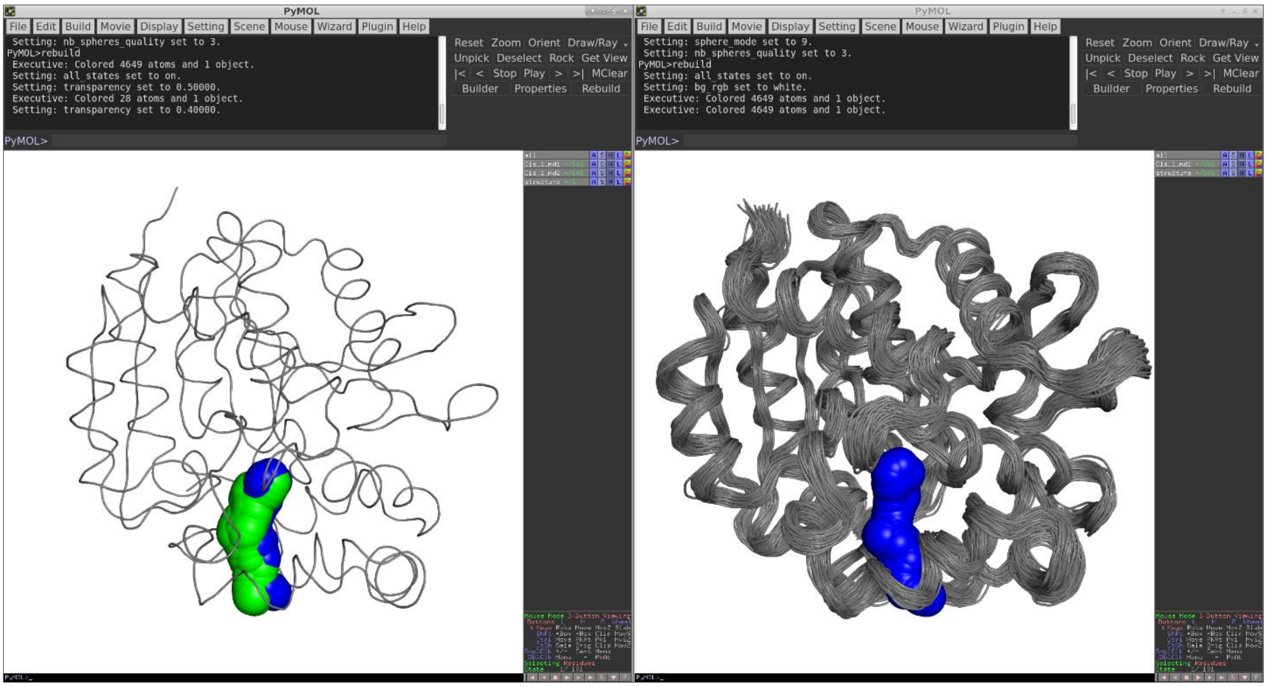

**Figure 30 PyMOL sessions obtained using visualization scripts from *static* (left) and *dynamics* (right) folders.** In the static representation, PyMOL session contains the ensemble of tunnels for supercluster 1 in md1 (blue spheres) and md2 (green spheres) with a single representative protein reference structure (gray ribbon). In the dynamics representation, the ensemble of tunnels for supercluster 1 in md1 trajectory only (blue spheres) is presented according to the definition in the *custom_analysis.py* script, together with the corresponding ensemble of protein structures (superimposed gray ribbons).

### 3. Standard *TransportTools* workflow

#### 3.1. Preparatory stage defining transformations needed for unified analyses

In stage 1, the output folder for the analyses' results is prepared at the location specified by the *output_path* parameter. Next, the biomolecular structures representing individual systems with both tunnel networks and/or transport events [if those are to be analyzed] are aligned on the reference PDB file (can be specified by *pdb_reference_structure* parameter), and the whole system is translated to have their global origin in coordinates of the mean starting point of all tunnel networks, producing respective transformation matrices.

***Inputs required:***

- reference PDB files from *data* folders of CAVER results
- *6_visualize_results.tar.gz* files from AQUA-DUCT [optional if transport events are to be analyzed]

***Outputs produced:***

- *output_path/*_internal/transformations - transformation matrices and the reference PDB file

***Basic relevant parameters:***

- *caver_results_path*
- *caver_results_folder_pattern*
- *caver_results_relative_subfolder_path*
- *aquaduct_results_path* [optional if transport events are to be analyzed]
- *aquaduct_results_folder_pattern* [optional if transport events are to be analyzed]
- *aquaduct_results_relative_tarfile* [optional if transport events are to be analyzed]
- *trajectory_engine* [optional, is autodetected]
- *snapshots_per_simulation* [optional, can be detected from trajectory files if available]
- *pdb_reference_structure* [optional, can be automatically selected from CAVER results]

#### 3.2. Processing of input datasets of tunnel networks

In stage 2, tunnel networks are parsed from the CAVER results, transformed into a uniform coordinate system, and stored to enable their further processing. Only tunnels with their bottleneck radii >= *min_tunnel_radius4clustering*, lengths >= *min_tunnel_length4clustering*, and curvature below *max_tunnel_curvature4clustering* are considered at this stage.

***Inputs required:***

- *tunnel_profiles.csv* files from *analysis* folder of CAVER results
- *bottlenecks.csv* files from *analysis* folder of CAVER results [optional if *process_bottleneck_residues* = True]
- transformation matrices produced in stage 1

***Outputs produced:***

- *output_path*/_internal/network_data/caver – folder with binary files with transformed tunnel networks
- *output_path*/visualization/sources/network_data/caver – folder with files for visualization of transformed networks [optional if *visualize_transformed_tunnels* = True]

***Basic relevant parameters:***

- *snapshot_id_position*
- *snapshot_delimiter*
- *process_bottleneck_residues*
- *min_tunnel_radius4clustering*
- *min_tunnel_length4clustering*
- *max_tunnel_curvature4clustering*
- *visualize_transformed_tunnels*

#### 3.3. Layering tunnel clusters to get their simplified representation

In stage 3, points forming tunnel clusters processed in the previous stage (**Figure 31A**) are divided into layers with given *layer_thickness* (**Figure 31B**). Next, all points in the same layers are then clustered into representative nodes, while filtering the outliers (**Figure 31C**). Finally, layered paths connecting those nodes from the starting point to all nodes corresponding to the ends of original tunnels in a manner similar to the original tunnels are identified (**Figure 31D**). A set of these paths can then serve to efficiently represent the geometry of all tunnel cluster's within the analyzed MD simulation (**Figure 31E**).

***Inputs required:***

- files with transformed tunnel networks produced in stage 2

***Outputs produced:***

- *output_path*/_internal/layered_data/caver – folder with binary files with layered representations of tunnel clusters

***Basic relevant parameters:***

- *layer_thickness*

#### 3.4. Computing distances among the layered clusters

In stage 4, the average distances between surfaces of layered paths representing tunnel clusters are computed across all analyzed MD simulations (**Figure 32A,B**). Note that distances between tunnel clusters that are seriously directionally misaligned are not directly evaluated, and their distance is assumed to be 999. *Hint*: Since the calculations of distances between two tunnel clusters are stopped when those are clearly too distant (defined as 3 * *clustering_cutoff* parameter * a number of layered paths in firsts tunnel cluster * a number of layered paths in the second tunnel cluster) and hence cannot belong to the same supercluster and are assumed to be 999 apart. Therefore, these distance calculations will take progressively longer when larger values are used for the *clustering_cutoff* parameter due to later termination.

***Inputs required:***

- files with layered representations of tunnel clusters produced in stage 3

***Outputs produced:***

- *output_path/*_internal/clustering – folder with binary files containing distance matrix and the corresponding definition of tunnel clusters
- *output_path*/data/clustering/tunnel_clusters_distance_matrix.csv – CSV formatted distance matrix for tunnel clusters [optional if *save_distance_matrix_csv* = True]

***Basic relevant parameters:***

- *clustering_cutoff*
- *save_distance_matrix_csv*

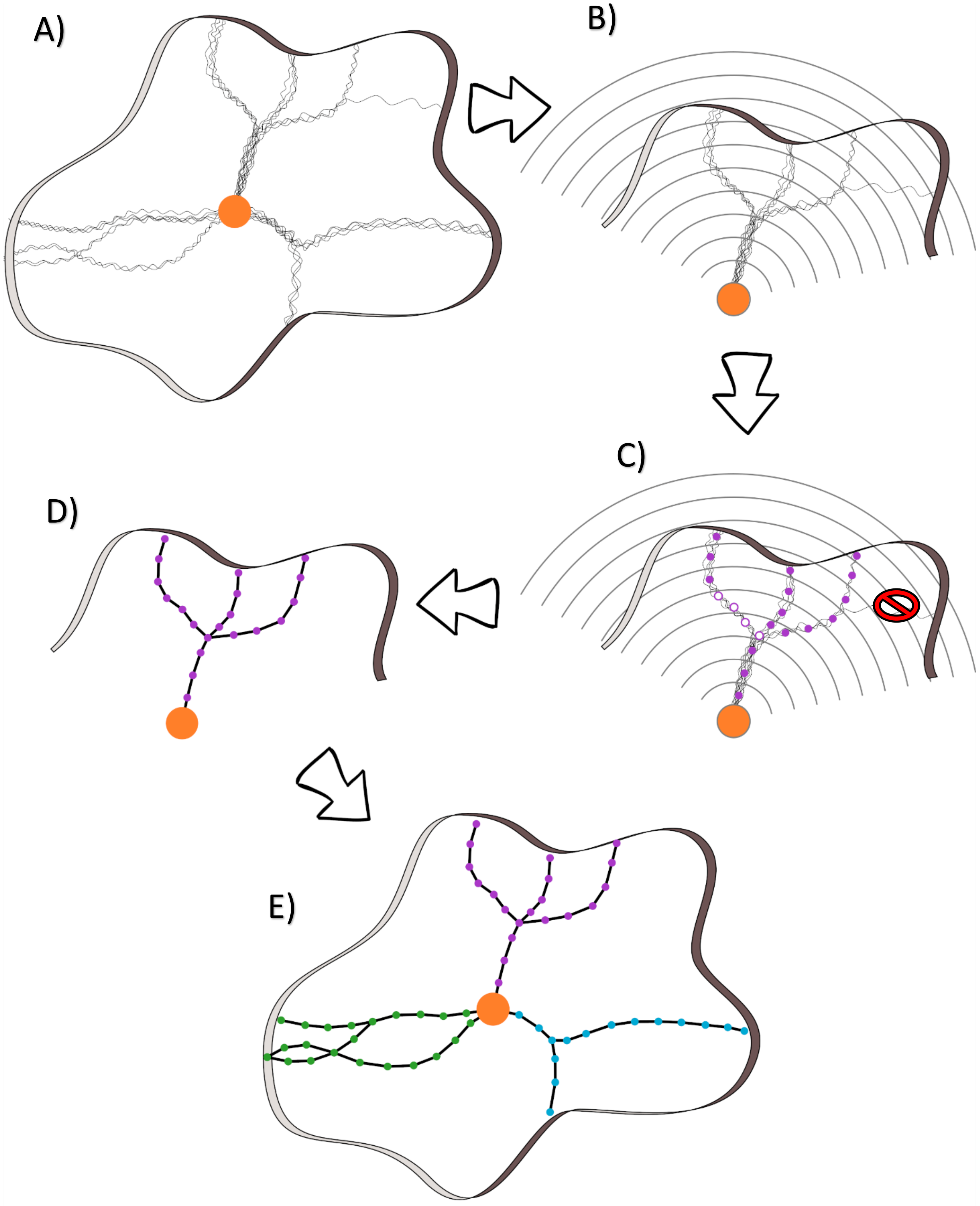

**Figure 31. A schema of tunnel clusters transformation into their simplified representations. A**) Multiple tunnel clusters found in a single MD simulation formed by tunnels leading from the starting point (orange dot). **B**) Division of tunnels forming the clusters into layers of given thickness (*layer_thickness*). **C**) After layering, nodes are defined for each layer by clustering the original tunnel spheres assigned to this layer. Please note that there can be more than one node in the same layer (empty circles). Also, some very rare tunnels could be omitted as outliers. **D**) The nodes are joined and the representation of the geometry of the whole tunnel cluster is obtained. **E**) Finally, the simplified representations of all tunnel clusters are obtained.

#### 3.5. Clustering the layered clusters into superclusters and creating initial outputs

In stage 5, layered tunnel clusters are clustered into superclusters using a hierarchical clustering algorithm with linkage defined by *clustering_linkage* parameter and using distance cutoff for merging of layered clusters into superclusters defined by *clustering_cutoff* parameter (**Figure 32C**). Subsequently, by processing complete data on tunnel networks belonging to each supercluster, overall statistics on the superclusters, their cumulative tunnel profiles, bottleneck residues, and details about membership of tunnel clusters in superclusters are generated together with respective visualization files (**Figure 32D**). Note that only one tunnel with the highest throughput is assigned to a single supercluster in a context of a single protein structure.

***Inputs required:***

- files containing distance matrix and the corresponding definition of tunnel clusters produced in stage 4
- binary files with transformed tunnel networks produced in stage 2

***Outputs produced:***

- *output_path/*_internal/super_cluster_pathsets – folder with binary files containing the layered representation of overall volumes occupied by superclusters
- *output_path/*_internal/super_cluster_profiles/initial – folder with binary files containing tunnel profiles and optionally also bottleneck data for each supercluster
- *output_path*/data/super_clusters/details/initial_super_cluster_details.txt – file with detailed information on composition of superclusters
- *output_path*/visualization/sources/super_cluster_CGOs – folder with CGO formatted data for visualization of superclusters (*_pathset1.dump.gz)
- *output_path*/visualization/1-visualize_initial_tunnels.py – PyMOL script for visualization of superclusters
- *output_path*/statistic/1-initial_tunnels_statistics.txt – file with overall statistics on tunnel ensembles forming each supercluster
- *output_path*/data/super_clusters/CSV_profiles/initial – folder with CSV formatted files containing tunnel profiles for each supercluster [optional if *save_super_cluster_profiles_csvs* = True]
- *output_path*/data/super_clusters/bottlenecks/initial – folder with CSV formatted files carrying information on bottlenecks and their constitutive residues for each supercluster [optional if *process_bottleneck_residues* = True]
- *output_path*/statistics/1-initial_tunnels_statistics_bottleneck_residues.txt – file with overall statistics on bottleneck residues of each supercluster [optional if *process_bottleneck_residues* = True]
- *output_path*/visualization/sources/super_cluster_CGOs/*_volume1.dump.gz – files data for visualization of supercluster volumes [optional if *visualize_super_cluster_volumes* = True]

***Basic relevant parameters:***

- *clustering_linkage*
- *clustering_cutoff*
- *save_super_cluster_profiles_csvs*
- *process_bottleneck_residues*
- *visualize_super_cluster_volumes*

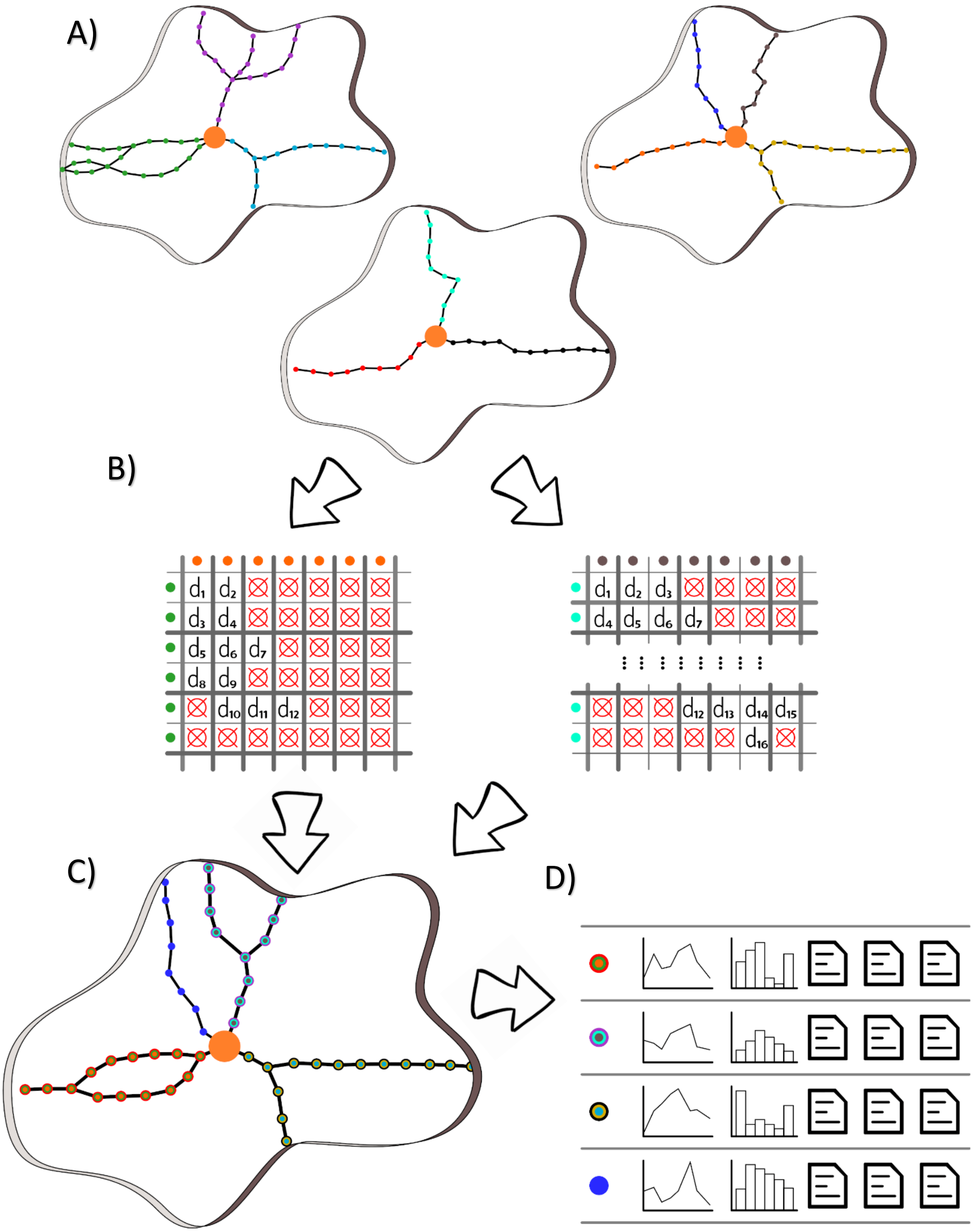

**Figure 32. Workflow for superclusters generation. A**) Simplified representations of tunnel clusters geometries from different MD simulations. **B**) For each layered path representing a tunnel cluster, distances to all paths in other clusters are calculated by averaging the shortest distances between nodes forming those paths located in the adjacent layers (distances above the *clustering_cutoff* are denoted as ⌖). **C**) tunnel clusters that are within *clustering_cutoff* are then merged into superclusters. **D**) Finally, each supercluster is analyzed, and outputs are generated for each one.

#### 3.6. Filtering superclusters and creating filtered outputs

In stage 6, superclusters profiles are filtered using defined filters on tunnel geometry and their numbers in superclusters. Then, overall statistics on the filtered superclusters, their cumulative tunnel profiles, bottleneck residues, and details about the membership of tunnel clusters in superclusters are generated together with respective visualization files.

***Inputs required:***

- binary files containing tunnel profiles and optionally also bottleneck data for each supercluster generated in stage 5

***Outputs produced:***

- *output_path/*_internal/super_cluster_profiles/filtered01 – folder with binary files containing tunnel profiles and optionally also filtered bottleneck data for each supercluster filtered based on tunnels
- *output_path*/data/super_clusters/details/filtered_super_cluster_details1.txt – file with detailed information on composition of superclusters filtered based on tunnels
- *output_path*/visualization/sources/super_cluster_CGOs – folder with CGO formatted data for visualization of superclusters filtered based on tunnels (*_pathset2.dump.gz)
- *output_path*/visualization/2-visualize_filtered_tunnels.py – PyMOL script for visualization of superclusters filtered based on tunnels
- *output_path*/statistics/2-filtered_tunnels_statistics.txt – file with overall statistics on tunnel ensembles forming each supercluster filtered based on tunnels
- *output_path*/data/super_clusters/CSV_profiles/filtered01 – folder with CSV formatted files containing tunnel profiles for each supercluster filtered based on tunnels [optional if *save_super_cluster_profiles_csvs* = True]
- *output_path*/data/super_clusters/bottlenecks/filtered01 – folder with CSV formatted files carrying information on bottlenecks and their constitutive residues for each supercluster filtered based on tunnels [optional if *process_bottleneck_residues* = True]
- *output_path*/statistics/2-filtered_tunnels_statistics_bottleneck_residues.txt – file with overall statistics on bottleneck residues of each supercluster filtered based on tunnels [optional if *process_bottleneck_residues* = True]
- *output_path*/visualization/sources/super_cluster_CGOs/*_volume2.dump.gz – files data for visualization of supercluster volumes [optional if *visualize_super_cluster_volumes* = True]

***Basic relevant parameters:***

- *min_length*
- *max_length*
- *min_bottleneck_radius*
- *max_bottleneck_radius*
- *min_curvature*
- *max_curvature*
- *min_sims_num*
- *min_snapshots_num*
- *min_avg_snapshots_num*
- *save_super_cluster_profiles_csvs*
- *process_bottleneck_residues*
- *visualize_super_cluster_volumes*

#### 3.7. Processing datasets of transport events of ligands

In stage 7, networks of transport events are parsed from the AQUA-DUCT results, transformed to a uniform coordinate system, and stored to enable their further processing. Only events that get closer than *event_min_distance* to the starting point of tunnel networks are considered at this stage (**Figure 33A**).

***Inputs required:***

- *5_analysis_results.txt* and *6_visualize_results.tar.gz* files from AQUA-DUCT analyses
- transformation matrices produced in stage 1

***Outputs produced:***

- *output_path*/_internal/network_data/aquaduct – folder with binary files with transformed networks of transport events
- *output_path*/visualization/sources/network_data/aquaduct – folder with files for visualization of transformed networks [optional if *visualize_transformed_transport_events* = True]

***Basic relevant parameters:***

- *aquaduct_results_path*
- *aquaduct_results_folder_pattern*
- *aquaduct_results_relative_tarfile*
- *aquaduct_results_relative_summaryfile*
- *event_min_distance*
- *visualize_transformed_transport_events*

#### 3.8. Layering transport events to get their simplified representation

In stage 8, points forming traces of transport events processed in the previous stage are divided into layers with given *layer_thickness.* Next, all points in the same layers are then clustered into representative nodes while filtering the outliers (**Figure 33B**). Finally, layered paths connecting those nodes in a manner similar to the original event's trace are identified to represent the geometry of the event's trace efficiently (**Figure 33C**).

***Inputs required:***

- files with transformed networks of events produced in stage 7

***Outputs produced:***

- *output_path*/_internal/layered_data/aquaduct – folder with binary files with layered representations of transport events
- *output_path*/visualization/sources/layered_data/aquaduct – folder with CGO formatted data for visualization of layered events

***Basic relevant parameters:***

- *layer_thickness*

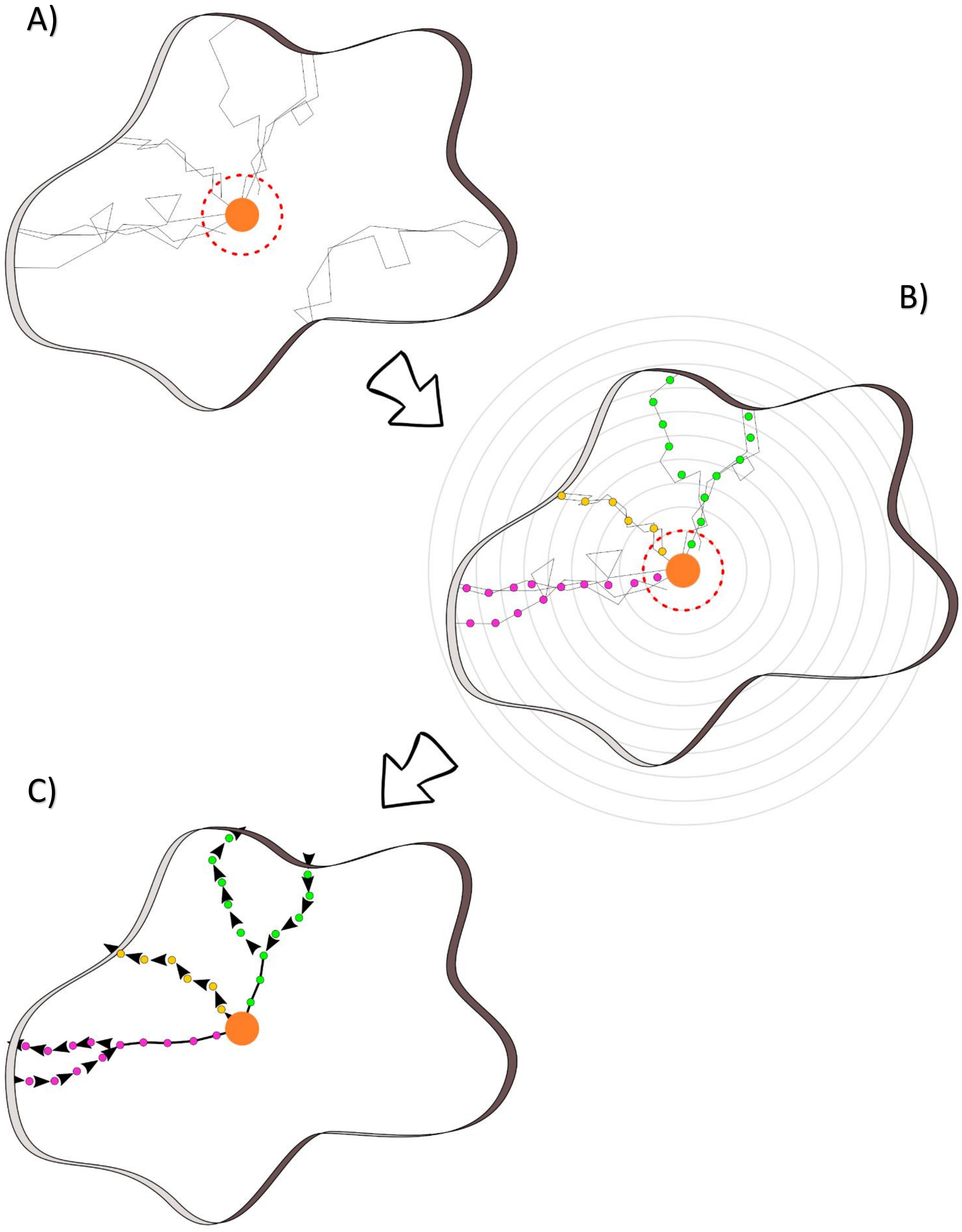

**Figure 33 A schema of transformation of transport events into their simplified representations. A)** Transport events are considered for analysis only if they reach the internal region (dotted red circle). **B)** The layering of the clusters follows the same procedure as for tunnel clusters (see chapter 3.3). **C)** Layered transport events are separated into those representing ligand entry and release, shown here with arrows.

#### 3.9. Assigning layered events to tunnel networks in superclusters and creating initial outputs

In stage 9, layered transport events are assigned to superclusters when the fraction of their traces buried inside the supercluster volumes are larger than *event_assignment_cutoff* (**Figure 34**). In case an event is equally buried in multiple superclusters, the final assignment is resolved according to the *ambiguous_event_assignment_resolution* method.

***Inputs required:***

- binary files with layered representations of transport events generated in stage 8
- binary files containing a layered representation of overall volumes occupied by superclusters generated in stage 5
- binary files containing tunnel profiles and optionally also bottleneck data for each supercluster generated in stage 5

***Outputs produced:***

- *output_path*/data/super_clusters/details/initial_super_cluster_events_details.txt – file with detailed information on the composition of superclusters filtered based on tunnels and data on events assigned to those superclusters
- *output_path*/data/super_clusters/details/outlier_transport_events_details.txt – file with detailed information outlier events that could not be assigned to any supercluster filtered based on tunnels
- *output_path*/visualization/sources/super_cluster_CGOs – folder with CGO formatted data for visualization of superclusters filtered based on tunnels (*_pathset3.dump.gz)
- *output_path*/visualization/3-visualize_initial_events.py – PyMOL script for visualization of superclusters filtered based on tunnels as well as visualization of assigned and outlying events
- *output_path*/statistics/3-initial_events_statistics.txt – file with overall statistics on tunnel ensembles forming each supercluster filtered based on tunnels with assigned events
- *output_path*/statistics/3-initial_events_statistics_bottleneck_residues.txt – file with overall statistics on bottleneck residues of each supercluster with assigned events filtered based on tunnels [optional if *process_bottleneck_residues* = True]
- *output_path*/visualization/sources/super_cluster_CGOs/*_volume3.dump.gz – files data for visualization of supercluster volumes [optional if *visualize_super_cluster_volumes* = True]

***Basic relevant parameters:***

- *event_assignment_cutoff*
- *ambiguous_event_assignment_resolution*
- *process_bottleneck_residues*
- *trajectory_engine*
- *visualize_super_cluster_volumes*

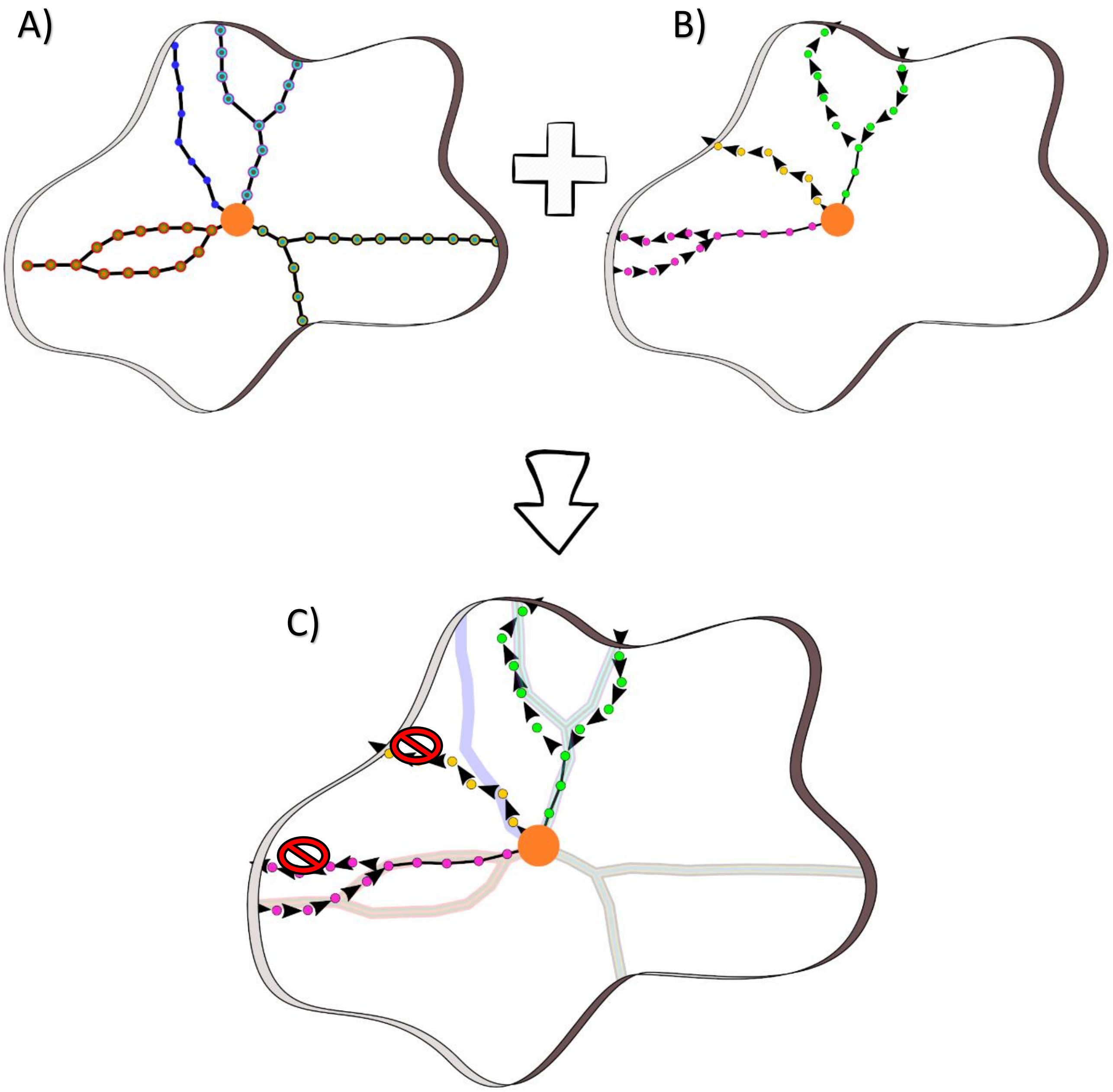

**Figure 34 Workflow for assignment of transport events into superclusters. A)** Superclusters and **B)** information from layered transport events are joined to assign transport events to superclusters. **C)** A transport event (colored discs) is assigned to a supercluster only if a fraction of their trajectory is buried in the volume formed by a given supercluster (shaded regions) is larger than *event_assignment_cutoff.*

#### 3.10. Filtering superclusters with events and creating filtered outputs with events

In stage 10, superclusters profiles are filtered using additional filters on a number of transport events assigned to them. Then, overall statistics on the superclusters, their cumulative tunnel profiles, bottleneck residues, and details about the membership of tunnel clusters in superclusters are generated together with respective visualization files.

***Inputs required:***

- binary files containing tunnel profiles and optionally also bottleneck data for each supercluster generated in stage 5

***Outputs produced:***

- *output_path/*_internal/super_cluster_profiles/filtered02 – folder with binary files containing filtered tunnel profiles and optionally also bottleneck data for each supercluster filtered based on both tunnels and events
- *output_path*/data/super_clusters/details/filtered_super_cluster_details2.txt – file with detailed information on the composition of superclusters with assigned events filtered based on both tunnels and events
- *output_path*/visualization/sources/super_cluster_CGOs – folder with CGO formatted data for visualization of superclusters filtered based on both tunnels and events (*_pathset4.dump.gz)
- *output_path*/visualization/4-visualize_filtered_events.py – PyMOL script for visualization of superclusters with assigned as well as outlying events filtered based on both tunnels and events
- *output_path*/statistics/4-filtered_events_statistics.txt – file with overall statistics on tunnel ensembles forming each supercluster filtered based on both tunnels and events
- *output_path*/data/super_clusters/CSV_profiles/filtered02 – folder with CSV formatted files containing tunnel profiles for each supercluster filtered based on both tunnels and events [optional if *save_super_cluster_profiles_csvs* = True]
- *output_path*/data/super_clusters/bottlenecks/filtered02 – folder with CSV formatted files carrying information on bottlenecks and their constitutive residues for each supercluster filtered based on both tunnels and events [optional if *process_bottleneck_residues* = True]
- *output_path*/statistics/4-filtered_events_statistics_bottleneck_residues.txt – file with overall statistics on bottleneck residues of each supercluster filtered based on both tunnels and events [optional if *process_bottleneck_residues* = True]
- *output_path*/visualization/sources/super_cluster_CGOs/*_volume4.dump.gz – files data for visualization of supercluster volumes [optional if *visualize_super_cluster_volumes* = True]

***Basic relevant parameters:***

- *min_total_events*
- *min_entry_events*
- *min_release_events*
- *save_super_cluster_profiles_csvs*
- *process_bottleneck_residues*
- *visualize_super_cluster_volumes*

### 4. Parameters in the configuration file

The configuration file uses INI-like formatting following the *configparser* Python module (see here for more details on general formatting and parsing rules). There are three standard sections defining **[INPUT_PATHS]**, **[CALCULATIONS_SETTINGS]**, and **[OUTPUT_SETTINGS]**, as well as the optional **[ADVANCED_SETTINGS]** section. The three compulsory parameters are underlined, the remaining parameters are optional and can be omitted from the configuration file.

#### 4.1. [INPUT_PATHS] section

| **Parameter** | **Values [default]** | **Relevant Stage(s)** | **Description** |
|---|---|---|---|
| *caver_results_path* | [None] | 1, 2, 3, 9 | A path to the folder with results from CAVER analyses per individual MD simulation. It cannot include UNIX-like patterns. |
| *caver_results_folder_pattern* | * | 1, 2, 3, 9 | UNIX-like pattern to recognize which subfolders located in *caver_results_path* should be analyzed for CAVER results per individual MD simulation. By default, all subfolders will be considered. |
| *caver_results_relative_subfolder_path* | [None] | 1, 2, 3, 9 | Path to a subfolder with CAVER results located in *caver_results_path/caver_results_folder_pattern* that contains standard outputs of CAVER, in particular *analysis* and *data* folders. Can include UNIX-like patterns. |
| *aquaduct_results_path* | [None] | 1, 7, 8, 9 | A path to the folder with results from AQUA-DUCT analyses per individual MD simulation. It cannot include UNIX-like patterns. |
| *aquaduct_results_folder_pattern* | * | 1, 7, 8, 9 | UNIX-like pattern to recognize which subfolders located in *aquaduct_results_path* should be analyzed for AQUA-DUCT results per individual MD simulation. By default, all subfolders will be considered. Can include UNIX-like patterns. |
| *aquaduct_results_relative_tarfile* | [None] | 1, 7, 8, 9 | Path to *6_visualize_results.tar.gz* files from AQUA-DUCT located in *aquaduct_results_path/aquaduct_results_folder_pattern.* Can include UNIX-like patterns. |
| *aquaduct_results_relative_summaryfile* | [None] | 7, 8, 9 | Path to *5_analysis_results.txt* files from AQUA-DUCT located in *aquaduct_results_path/aquaduct_results_folder_pattern.* Can include UNIX-like patterns. |
| *trajectory_path* | [None] | 1, 9 | A path to the folder with trajectories from individual MD simulations. It cannot include UNIX-like patterns. |
| *trajectory_folder_pattern* | * | 1, 9 | UNIX-like pattern to recognize which subfolders located in *trajectory_path* should be analyzed for trajectories from individual MD simulation. By default, all subfolders will be considered. |
| *trajectory_relative_file* | [None] | 1, 9 | Path to trajectory file located in *trajectory_path/trajectory_folder_pattern.* Can include UNIX-like patterns. Accepted file formats are dependent on support in *mdtraj/pytraj* package. For details see here and here, respectively. |
| *topology_relative_file* | [None] | 1, 9 | Path to topology file located in *trajectory_path/trajectory_folder_pattern.* Can include UNIX-like patterns. Accepted file formats are dependent on support in *mdtraj/pytraj* package. For details see here and here, respectively. |

#### 4.2. [CALCULATIONS_SETTINGS] section

| **Parameter** | **Values [default]** | **Relevant Stage(s)** | **Description** |
|---|---|---|---|
| *start_from_stage* | 0-10, [1] | all | From which stage, the analysis starts. |
| *stop_after_stage* | 1-10 / [None] | all | After which stage, the analysis stops. |
| *num_cpus* | >=1 / [None] | all | The number of CPU cores to be used during parallel processing phases. By default, the number is decided based on the number of CPU cores detected in the system while leaving 1 or 2 cores free in systems with 2-4 cores and more than 4 cores, respectively. |
| *pdb_reference_structure* | [None] | 1 | The reference structure to align all systems to. |
| *layer_thickness* | >1.0, [1.5] | 3, 8 | The thickness of a concentrically layered grid. |
| *snapshots_per_simulation* | >=1 / [None] | 5, 6, 9, 10 | A number of snapshots used for CAVER calculation. By default, the number of snapshots is read from MD trajectories. If the trajectories are not available, this becomes a compulsory parameter. |

| | | | |
|---|---|---|---|
| *caver_traj_offset* | >=0 / [1] | 9 | The difference in IDs of *mdtraj*/*pytraj* frames (from 0) and caver snapshots (often from 1). |
| *snapshot_id_position* | 1 | 2 | The location of IDs in snapshot filenames from CAVER when split by *snapshot_delimiter.* |
| *snapshot_delimiter* | . | 2 | The delimiter used for splitting snapshot filenames from CAVER to get IDs at position *snapshot_id_position.* |
| *process_bottleneck_residues* | True / [False] | 2, 5, 6, 9, 10 | Process bottlenecks.csv files from the *analysis* folder of standard CAVER results to get info about tunnels' bottleneck residues and controls if the bottleneck-related outputs will be produced in later stages. |
| *min_tunnel_radius4clustering* | >=0 [0.75] | 2 | Minimum bottleneck radius filter applied during tunnel processing that is relevant for layering, supercluster definition & visualization but does not affect the superclusters statistics. |
| *min_tunnel_length4clustering* | >=0 [5.0] | 2 | Minimum length filter applied during tunnel processing that is relevant for layering, supercluster definition & visualization but does not affect the superclusters statistics. |
| *max_tunnel_curvature4clustering* | >=1 [2.0] | 2 | Maximum curvature filter applied during tunnel processing that is relevant for layering, supercluster definition & visualization but does not affect the superclusters statistics. |
| *clustering_linkage* | single / [complete] / average / weighted / ward / centroid / median | 5 | Linkage used during agglomerative clustering of tunnel clusters into superclusters. See the documentation of *fastcluster* package for more details see here. |
| *clustering_cutoff* | >=0 [2.0] | 4, 5 | A cutoff for surface distances among tunnel clusters to be considered a single supercluster. |
| *min_length* | >=0 [-1] | 6, 10 | Output filter on minimum tunnel length. By default, the filter is inactive. |
| *max_length* | >=0 [-1] | 6, 10 | Output filter on maximum tunnel length. By default, the filter is inactive. |
| *min_bottleneck_radius* | >=0 [-1] | 6, 10 | Output filter on minimum tunnel bottleneck radius. By default, the filter is inactive. |
| *max_bottleneck_radius* | >=0 [-1] | 6, 10 | Output filter on maximum tunnel bottleneck radius. By default, the filter is inactive. |
| *min_curvature* | >=1 [-1] | 6, 10 | Output filter on minimum tunnel curvature. By default, the filter is inactive. |
| *max_curvature* | >1 [-1] | 6, 10 | Output filter on maximum tunnel curvature. By default, the filter is inactive. |
| *min_sims_num* | >=0 [-1] | 6, 10 | Output filter on supercluster presence in a minimum number of MD simulations. By default, the filter is inactive. |
| *min_snapshots_num* | >=0 [-1] | 6, 10 | Output filter on supercluster presence in a minimum number of snapshots. By default, the filter is inactive. |
| *min_avg_snapshots_num* | >=0 [-1] | 6, 10 | Output filter on supercluster presence in a minimum number of snapshots on average. By default, the filter is inactive. |
| *event_min_distance* | >0 [6.0] | 7 | The closest distance of the event molecule from the starting point to be processed. |
| *event_assignment_cutoff* | 0.0-1.0 [0.85] | 9 | The minimal buriedness of transport event in a supercluster to be assigned to that supercluster. |
| *ambiguous_event_assignment_resolution* | assign2all / [penetration_depth] / exact_matching | 9 | A method for selecting assigned supercluster from among all those that the event is buried in:<br>• *penetration_depth* – assign based on the depth the event penetrates the supercluster,<br>• *assign2all* - assign the event to all superclusters in which it is buried,<br>• *exact_matching* - matching of actual ligand transport event (all atoms from MD simulation) and real tunnels existing in a given simulation during the event occurrence. |
| *min_total_events* | >=0 [-1] | 10 | Additional output filter on a supercluster having minimum events assigned. By default, the filter is inactive. |
| *min_entry_events* | >=0 [-1] | 10 | Additional output filter on a supercluster having minimum entry events assigned. By default, the filter is inactive. |
| *min_release_events* | >=0 [-1] | 10 | Additional output filter on a supercluster having minimum release events assigned. By default, the filter is inactive. |

#### 4.3. [OUTPUT_SETTINGS] section

| **Parameter** | **Values [default]** | **Relevant Stage(s)** | **Description** |
|---|---|---|---|
| *output_path* | [results] | 1 | The name of the output folder. |
| *save_super_cluster_profiles_csvs* | True / [False] | 5, 6, 10 | If the cumulative supercluster tunnel profile is to be generated in CSV format akin to CAVER tunnel profiles in: *output_path*/data/super_clusters/CSV_profiles Subfolders *initial, filtered01* and *filtered02* correspond to superclusters at stages 5, 6, and 10, respectively. |
| *save_distance_matrix_csv* | True / [False] | 4 | If the distances among tunnel clusters are to be saved to CSV formatted file in: *output_path*/data/clustering/tunnel_clusters_distance_matrix.csv |
| *visualize_super_cluster_volumes* | True / [False] | 5, 6, 9, 10 | If the volume representation of superclusters is to be calculated for visualization in: *output_path/*visualization/sources/super_cluster_CGOs |
| *visualize_transformed_tunnels* | True / [False] | 2 | If the visualizations of transformed tunnel networks are to be generated in: *output_path*/visualization/sources/network_data/caver |
| *visualize_transformed_transport_events* | True / [False] | 7 | If the visualizations of transformed events are to be generated in: *output_path*/visualization/sources/network_data/aquaduct |

#### 4.4. [ADVANCED_SETTINGS] section

| **Parameter** | **Values [default]** | **Relevant Stage(s)** | **Description** |
|---|---|---|---|
| *random_seed* | [4] | 3, 4, 5, 6, 8, 9, 10 | A value to initiate the random number generator to have reproducible results. |
| *directional_cutoff* | [np.pi / 2] | 4, 9 | A cutoff on the angle between the layered representation of events and clusters used for the detection of severe misalignments during tunnel clusters clustering into superclusters and assignment of events to superclusters. |
| *aqauduct_ligand_effective_radius* | >0 [1.0] | 7, 9 | Approximate radius for AQUA-DUCT events for calculation of overlaps. |
| *perform_exact_matching_analysis* | True / [False] | 9 | Performs exact analyses of the event concerning tunnel clusters in frames during the event occurrence originating directly from the simulation sampling the event. See chapter 6.2 for more details. |
| *perform_comparative_analysis* | True / [False] | 1, 5, 6, 9, 10 | Separates supercluster statistics and visualization per input MD simulation and groups defined by parameter *comparative_groups_definition*. See chapter 6.1 for more details. |
| *visualize_comparative_super_cluster_volumes* | True / [False] | 5, 6, 9, 10 | If the volume representation of superclusters is to be calculated for visualization of groups defined during comparative analysis in: *output_path/*visualization/sources/super_cluster_CGOs |
| *comparative_groups_definition* | [None] | 1, 5, 6, 9, 10 | Defines which input MD simulations are to be considered as a single group when *perform_comparative_analysis* = True. See chapter 6.1 for more details. |
| *overwrite* | True / [False] | All | Controls if the cleaning of non-empty *output_path* folder is performed and overwriting of existing checkpoint files. |
| *visualize_exact_matching_outcomes* | True / [False] | 9 | Prepare complete visualization of matched events analyzed when *perform_exact_matching_analysis* = True. See chapter 6.2 for more details. Note that activating this option could be very demanding on computing resources in terms of I/O operations, used memory. |

| | | | |
|---|---|---|---|
| *visualize_layered_clusters* | True / [False] | 3 | Produces folders with files for visualization of layered representations of tunnel clusters from individual MD simulations in *output_path*/visualization/sources/layered_data/caver<br><br>Each such folder contains the following data:<br>• *nodes* – folder with PDB files of nodes representing a cluster of tunnel points in a particular layer, shown with at maximum 500 original tunnel points,<br>• *paths* – folder with layered representations of tunnel clusters in CGO format,<br>• *origin.pdb* – PDB file with a starting point,<br>• *_C_trans_rot.pdb – PDB file of structure representing that particular biomolecule aligned to reference structure,<br>*Cluster_*.py* - scripts for PyMOL to visualize layered tunnel cluster including nodes. |
| *visualize_layered_events* | True / [False] | 8 | Produces folders with files for visualization of layered representations of events from individual MD simulations in *output_path*/visualization/sources/layered_data/aquaduct<br><br>Each such folder contains the following data:<br>• *nodes* – subfolder with PDB files of nodes representing an event in a particular layer, shown with at maximum 500 original event points,<br>• *_[entry, release].py - scripts for PyMOL to visualize layered tunnel cluster including nodes. |
| *legacy_pymol_support* | True / [False] | 5, 6, 9, 10 | Visualization data are prepared in a format compliant with Python2-based PyMOL distributions. |
| *trajectory_engine* | pytraj / [mdtraj] | 1, 9 | Defines engine to be used for loading and processing trajectories. Default is *mdtraj* but in case *pytraj* is detected in the environment, it is automatically selected unless overridden in this parameter. |
| *logfilename* | [transport_tools.log] | all | Filename to which the *TransportTools* log will be stored. This file will be located in the *output_path* folder. |
| *log_level* | debug / warn / error / critical / [info] | all | Controls the level of details printed to the console. See here for more details. |
| *max_events_per_cluster4visualization* | 100 – 5000, [1000] | 9, 10 | Limit on the number of visualized events assigned to a single supercluster to avoid too long load times and potential memory troubles when visualizing in PyMOL. |

### 5. Description of standard *TransportTools* outputs

The results of *TransportTools* analyses employing its standard workflow are stored in the *output_path* folder that contains calculation *logfile* and the following structure of subfolders (optionally generated folders are shown in gray):

```
├── transport_tools.log
├── _internal
│   ├── transformations
│   │   ├── caver
│   │   └── aquaduct
│   ├── network_data
│   │   ├── caver
│   │   └── aquaduct
│   ├── layered_data
│   │   ├── caver
│   │   └── aquaduct
│   ├── clustering
│   ├── super_cluster_pathsets
│   ├── super_cluster_profiles
│   │   ├── initial
│   │   ├── filtered01
│   │   └── filtered02
│   └── checkpoints
├── data
│   ├── clustering
│   └── super_clusters
│       ├── details
│       ├── CSV_profiles
│       │   ├── initial
│       │   ├── filtered01
│       │   └── filtered02
│       └── bottlenecks
│           ├── initial
│           ├── filtered01
│           └── filtered02
├── statistics
└── visualization
    └── sources
        ├── network_data
        │   ├── caver
        │   │   ├── md1
        │   │   └── md2
        │   └── aquaduct
        │       ├── md1
        │       └── md2
        ├── layered_data
        │   └── aquaduct
        │       ├── md1
        │       └── md2
        └── super_cluster_CGOs
```

#### 5.1. _*internal* folder

This folder contains various interim data on tunnel networks and transport events used during the subsequent calculations, including the following subfolders:

- *transformations* – transformation matrices and the reference PDB files for networks of tunnels and transport events input data
- *network_data* - binary files with transformed networks of tunnels and transport events
- *layered_data* - binary files with layered representations of tunnel clusters and transport events
- *clustering* - binary files containing distance matrix and the corresponding definition of tunnel clusters
- *super_cluster_pathsets* - binary files containing a layered representation of overall volumes occupied by superclusters
- *super_cluster_profiles* - binary files containing tunnel profiles and optionally also bottleneck data for each supercluster
- *checkpoints* – binary restart files to start from particular stages

#### 5.2. *data* folder

This folder contains detailed data on superclusters, and <u>optionally</u> their clustering from original tunnel clusters [if *save_distance_matrix_csv* = True] in human-readable files with either plain text or CSV formats.

##### 5.2.1. *data/clustering* folder

In this <u>optional</u> location [if *save_distance_matrix_csv* = True], we can find a CSV-formatted distance matrix that is used for merging individual tunnel clusters (initially produced by CAVER) into the superclusters.

***Example:***

| | md1: cls1 | md1: cls2 | md1: cls3 | md1: cls4 | md1: cls5 | md1: cls6 | md1: cls7 | md1: cls8 | md1: cls9 | md1: cls10 | md1: cls11 | md1: cls12 | md1: cls13 | md1: cls14 |
|---|---|---|---|---|---|---|---|---|---|---|---|---|---|---|
| md1: cls1 | 0 | 999 | 2.22 | 3.538 | 0.618 | 999 | 999 | 999 | 999 | 999 | 999 | 999 | 0.424 | 999 |
| md1: cls2 | 999 | 0 | 0.949 | 3.315 | 3.627 | 3.343 | 999 | 0.492 | 999 | 999 | 999 | 999 | 999 | 999 |
| md1: cls3 | 2.22 | 0.949 | 0 | 3.162 | 1.448 | 999 | 999 | 1.924 | 999 | 999 | 999 | 999 | 999 | 999 |
| md1: cls4 | 3.538 | 3.315 | 3.162 | 0 | 3.827 | 999 | 999 | 999 | 999 | 999 | 999 | 999 | 3.204 | 999 |
| md1: cls5 | 0.618 | 3.627 | 1.448 | 3.827 | 0 | 999 | 999 | 999 | 999 | 999 | 999 | 999 | 2.067 | 999 |
| md1: cls6 | 999 | 3.343 | 999 | 999 | 999 | 0 | 999 | 4.498 | 999 | 2.933 | 999 | 3.896 | 999 | 2.033 |
| md1: cls7 | 999 | 999 | 999 | 999 | 999 | 999 | 0 | 999 | 1.131 | 3.456 | 0.786 | 3.486 | 999 | 999 |
| md1: cls8 | 999 | 0.492 | 1.924 | 999 | 999 | 4.498 | 999 | 0 | 999 | 999 | 999 | 999 | 999 | 999 |
| md1: cls9 | 999 | 999 | 999 | 999 | 999 | 999 | 1.131 | 999 | 0 | 3.867 | 2.744 | 2.921 | 999 | 4.155 |
| md1: cls10 | 999 | 999 | 999 | 999 | 999 | 2.933 | 3.456 | 999 | 3.867 | 0 | 2.786 | 0.777 | 999 | 1.792 |
| md1: cls11 | 999 | 999 | 999 | 999 | 999 | 999 | 0.786 | 999 | 2.744 | 2.786 | 0 | 3.517 | 999 | 999 |
| md1: cls12 | 999 | 999 | 999 | 999 | 999 | 3.896 | 3.486 | 999 | 2.921 | 0.777 | 3.517 | 0 | 999 | 1.256 |
| md1: cls13 | 0.424 | 999 | 999 | 3.204 | 2.067 | 999 | 999 | 999 | 999 | 999 | 999 | 999 | 0 | 999 |
| md1: cls14 | 999 | 999 | 999 | 999 | 999 | 2.033 | 999 | 999 | 4.155 | 1.792 | 999 | 1.256 | 999 | 0 |

***Legend:***
The first column and row contain the name of the input MD simulation and the original ID of the tunnel cluster. For example: *md1: cls1* -> tunnel cluster with ID 1 from simulation labeled as *md1.*

The distances between the clusters are the average distances between surfaces of layered paths representing tunnel clusters in Å. Clusters with overlapping volumes will have the distance of 0 Å, while clusters whose directions are severely misaligned or clearly too distant (defined as 3 * *clustering_cutoff* parameter * a number of layered paths in firsts tunnel cluster * a number of layered paths in the second tunnel cluster) have the distance shown as 999 Å.

##### 5.2.2. data/super_clusters folder

In this location, we can find details about superclusters' composition in *details* subfolder, and <u>optionally</u> also CSV-formatted data on tunnel profiles in *CSV_profiles* [if *save_super_cluster_profiles_csvs* = True], and tunnel bottlenecks in *bottlenecks* [if *process_bottleneck_residues* = True] subfolders.

###### 5.2.2.1. *data/super_clusters/details* folder

In this location, we can find the following five files with details about the composition of individual superclusters:

- *initial_super_cluster_details.txt* – initial composition of unfiltered superclusters
- *filtered_super_cluster_details1.txt* – composition of superclusters after filtering based on tunnels
- *initial_super_cluster_events_details.txt* – composition of superclusters after filtering based on tunnels including also information on assigned events
- *filtered_super_cluster_details2.txt* – composition of superclusters with assigned events after filtering based on both tunnels and events
- *outlier_transport_events_details.txt* – information about outlier events that could not be assigned to any supercluster after filtering based on tunnels

***Example:***

```
Tunnel clusters and transport events assignment to super clusters (ordered by their priority):
-------------------------------------------------------------------------------------------
Super cluster ID 1

Details on tunnel network:
Number of MD simulations = 2
Number of tunnel clusters = 4
Tunnel clusters:
from md1: 1, 5,
from md2: 1, 3,

Details on transport events:
Number of MD simulations = 1
Number of entry events = 3
Number of release events = 3
entry: (from Simulation: AQUA-DUCT ID, (Resname:Residue), start_frame->end_frame; ... )
from md2: 2, (WAT:294), 689->695; 3, (WAT:295), 714->715; 5, (WAT:297), 943->957;
release: (from Simulation: AQUA-DUCT ID, (Resname:Residue), start_frame->end_frame; ... )
from md2: 1, (WAT:293), 722->726; 2, (WAT:294), 698->704; 4, (WAT:296), 71->87;
-------------------------------------------------------------------------------------------
```

***Legend:***

There is a section for each supercluster providing information on

1. Details on tunnel network forming this supercluster
   a) Number of different MD simulations contributing some tunnel cluster
   b) Number of such tunnel cluster
   c) Identity of tunnel clusters forming this supercluster:
      *from (MD simulation name): a list of the original tunnel cluster IDs*
2. Details on transport events assigned to this supercluster
   a) Number of different MD simulations contributing some event
   b) Number of entry events assigned
   c) Number of release events assigned
   d) Identity of events assigned to this supercluster:
      *from (MD simulation name): a list of data on events separated by ';' where each event is defined by its original ID, (resname:residue), starting frame → end frame*

##### 5.2.2.2. *data/super_clusters/CSV_profiles* folder

In this optional location [if *save_super_cluster_profiles_csvs* = True], we can find CSV-formatted files containing tunnel profiles for each supercluster. Subfolder *initial* contains complete profile files, while subfolders *filtered01* and *filtered02* contain profile files after filtering based on tunnels and events, respectively.

***Example:***

| *MD_traj* | *Snapshot* | *Tunnel cluster* | *Tunnel* | *Through put* | *Cost* | *Bottleneck radius* | *Average R error bound* | *Max. R error bound* | *Bottleneck R error bound* | *Curvature* | *Length* | *Axis* | *Values ...* | | | | |
|---|---|---|---|---|---|---|---|---|---|---|---|---|---|---|---|---|---|
| md1 | system.1.pdb | 1 | 2 | 0.40 | 0.92 | 0.90 | - | - | - | 1.26 | 13.99 | *X* | -0.33 | -0.21 | 0.09 | 0.26 | ... |
| md1 | system.1.pdb | 1 | 2 | 0.40 | 0.92 | 0.90 | - | - | - | 1.26 | 13.99 | *Y* | 0.80 | 0.73 | 0.42 | 0.05 | ... |
| md1 | system.1.pdb | 1 | 2 | 0.40 | 0.92 | 0.90 | - | - | - | 1.26 | 13.99 | *Z* | -0.51 | -0.92 | -1.16 | -1.44 | ... |
| md1 | system.1.pdb | 1 | 2 | 0.40 | 0.92 | 0.90 | - | - | - | 1.26 | 13.99 | *distance* | 1.00 | 1.19 | 1.24 | 1.47 | ... |
| md1 | system.1.pdb | 1 | 2 | 0.40 | 0.92 | 0.90 | - | - | - | 1.26 | 13.99 | *length* | 1.00 | 1.50 | 2.00 | 2.50 | ... |
| md1 | system.1.pdb | 1 | 2 | 0.40 | 0.92 | 0.90 | - | - | - | 1.26 | 13.99 | *R* | 1.84 | 1.77 | 1.65 | 1.48 | ... |
| md1 | system.1.pdb | 1 | 2 | 0.40 | 0.92 | 0.90 | - | - | - | 1.26 | 13.99 | *Upper limit of R overestimation* | - | - | - | - | ... |
| md1 | system.2.pdb | 1 | 1 | 0.46 | 0.78 | 0.88 | - | - | - | 1.31 | 14.92 | *X* | 0.42 | 0.61 | 0.85 | 1.05 | ... |
| md1 | system.2.pdb | 1 | 1 | 0.46 | 0.78 | 0.88 | - | - | - | 1.31 | 14.92 | *Y* | -0.44 | -0.82 | -1.22 | -1.51 | ... |
| md1 | system.2.pdb | 1 | 1 | 0.46 | 0.78 | 0.88 | - | - | - | 1.31 | 14.92 | *Z* | 0.08 | -0.17 | -0.29 | -0.61 | ... |
| md1 | system.2.pdb | 1 | 1 | 0.46 | 0.78 | 0.88 | - | - | - | 1.31 | 14.92 | *distance* | 0.62 | 1.04 | 1.52 | 1.93 | ... |
| md1 | system.2.pdb | 1 | 1 | 0.46 | 0.78 | 0.88 | - | - | - | 1.31 | 14.92 | *length* | 0.62 | 1.12 | 1.62 | 2.12 | ... |
| md1 | system.2.pdb | 1 | 1 | 0.46 | 0.78 | 0.88 | - | - | - | 1.31 | 14.92 | *R* | 2.02 | 1.97 | 1.90 | 1.80 | ... |
| md1 | system.2.pdb | 1 | 1 | 0.46 | 0.78 | 0.88 | - | - | - | 1.31 | 14.92 | *Upper limit of R overestimation* | - | - | - | - | ... |

***Legend:***

*MD_Traj* – the name of the input MD simulation.
*Snapshot* – the name of the input PDB formatted file used for tunnel calculation by CAVER.
*Tunnel cluster* – the ID of a tunnel cluster to which a given tunnel belongs.
*Tunnel* – the tunnel ID in a given snapshot.
*Throughput* – the throughput of a given tunnel ($e^{-cost}$).
*Cost* – the cost of traversing a given tunnel according to the CAVER cost function.
*Bottleneck radius* – the radius of the narrowest part of a given tunnel (the bottleneck).
*Average R error bound* – average upper error bound for tunnel radii calculated by CAVER.
*Max. R error bound* – the maximal upper error bound for tunnel radii calculated by CAVER.
*Bottleneck R error bound* – the maximal possible overestimation of the bottleneck radius by CAVER.
*Curvature* – the tunnel curvature (tunnel length/distance between the tunnel start and end spheres).
*Length* – the tunnel length.
*Axis* – characteristics reported for each sphere forming the tunnel.
*Axis X* – the x coordinate of a given sphere.
*Axis Y* – the y coordinate of a given sphere.
*Axis Z* – the z coordinate of a given sphere.
*Axis distance* – the shortest possible distance between a given sphere and the tunnel starting point.
*Axis length* – length of the tunnel at a given sphere.
*Axis R* – tunnel radius at a given sphere.
*Axis Upper limit of R overestimation* – the upper error bound for estimation of the sphere radius.

###### 5.2.2.3. *data/super_clusters/bottlenecks* folder

In this <u>optional</u> location [if *process_bottleneck_residues* = True], we can find CSV-formatted files containing information on tunnel bottlenecks and their constitutive residues for each supercluster. Subfolder *initial* contains complete profile files, while subfolders *filtered01* and *filtered02* contain profile files after filtering based on tunnels and events, respectively.

***Example:***

| *MD_traj* | *Snapshot* | *Tunnel cluster* | *Tunnel* | *Throughput* | *Cost* | *Bottleneck X* | *Bottleneck Y* | *Bottleneck Z* | *Bottleneck R* | *Bottleneck residues* | | | | | | | | | |
|---|---|---|---|---|---|---|---|---|---|---|---|---|---|---|---|---|---|---|---|
| md1 | system.4.pdb | 1 | 1 | 0.518 | 0.657 | -1.138 | -5.071 | -4.763 | 1.199 | 142 | 146 | 173 | 169 | 141 | 138 | 172 | 165 | 242 | |
| md1 | system.5.pdb | 1 | 1 | 0.474 | 0.747 | -1.919 | -6.013 | -4.503 | 1.048 | 169 | 142 | 173 | 146 | 145 | 138 | 141 | 172 | 165 | 149 |
| md1 | system.8.pdb | 1 | 1 | 0.582 | 0.540 | -2.349 | -7.026 | -4.594 | 1.270 | 169 | 142 | 146 | 141 | 172 | 173 | 149 | 145 | 168 | 170 |
| md1 | system.10.pdb | 1 | 1 | 0.583 | 0.540 | -1.419 | -5.214 | -4.765 | 1.118 | 142 | 146 | 141 | 173 | 169 | 138 | 145 | 242 | | |
| md1 | system.11.pdb | 1 | 2 | 0.528 | 0.639 | -1.297 | -4.914 | -5.355 | 1.222 | 146 | 142 | 173 | 141 | 169 | 138 | 242 | 172 | 145 | |
| md1 | system.12.pdb | 1 | 1 | 0.471 | 0.752 | -1.493 | -5.257 | -5.035 | 1.015 | 146 | 142 | 173 | 138 | 169 | 145 | 141 | 242 | 172 | |

***Legend:***

*MD_Traj* – the name of the input MD simulation.
*Snapshot* – the name of the input PDB formatted file used for tunnel calculation by CAVER.
*Tunnel cluster* – the ID of a tunnel cluster to which a given tunnel belongs.
*Tunnel* – the tunnel ID in a given snapshot.
*Throughput* – the throughput of a given tunnel ($e^{-cost}$).
*Cost* – the cost of traversing a given tunnel according to the CAVER cost function.
*Bottleneck X* – x-coordinate of the tunnel's bottleneck sphere.
*Bottleneck Y* – y-coordinate of the tunnel's bottleneck sphere.
*Bottleneck Z* – z-coordinate of the tunnel's bottleneck sphere.
*Bottleneck R* – radius of the tunnel's bottleneck sphere.
*Bottleneck residues* – residue IDs of bottleneck residues surrounding the sphere.

#### 5.3. *statistics* folder

This folder contains four files with statistics on tunnels forming superclusters and assigned events, and <u>optionally</u> also four files with statistics on their bottleneck residues [if *process_bottleneck_residues* = True].

**Statistics on tunnel ensembles forming superclusters:**

- *1-initial_tunnels_statistics.txt* – the initial one
- *2-filtered_tunnels_statistics.txt* – after filtering based on tunnels
- *3-initial_events_statistics.txt* – with assigned events & after filtering based on tunnels
- *4-filtered_events_statistics.txt* – with assigned events & after filtering based on both tunnels and events

**Statistics on bottlenecks of tunnels forming superclusters:**

- *1-initial_tunnels_statistics_bottleneck_residues.txt* – the initial one
- *2-filtered_tunnels_statistics_bottleneck_residues.txt* – after filtering based on tunnels
- *3-initial_events_statistics_bottleneck_residues.txt* – with assigned events & after filtering based on tunnels
- *4-filtered_events_statistics_bottleneck_residues.txt* – with assigned events & after filtering based on both tunnels and events

***Example of statistics on tunnel ensembles forming superclusters:***

```
Statistics for: overall
Total simulations = 1
No snapshots per simulation = 1000

Parameters used for pre-selection of input tunnels for clustering:
length >= 5.00 A
radius >= 0.75 A
curvature <= 5.00

Active tunnel filters:
length = (5.00, 999.00) A
radius = (1.00, 999.00) A
curvature = (1.00, 999.00)
occurred in at least 0 simulations
occurred in at least 1 snapshots
occurred in at least 0.000 snapshots on average per simulation
has at least 0 transport events
has at least 0 entry events
has at least 0 release events

SC_ID, No_Sims, Total_No_Frames, Avg_No_Frames, Avg_BR, StDev, Max_BR, Avg_Len, StDev, Avg_Cur, StDev, Avg_throug,   StDev, Priority, Num_Events, Num_entries, Num_releases
    1,       1,             646,         646.0,  1.178, 0.129,  1.720,  14.094, 1.420,   1.243, 0.091,    0.54384, 0.06088,  0.35132,          0,           0,            0
    2,       1,             384,         384.0,  1.125, 0.090,  1.565,  14.274, 1.863,   1.189, 0.064,    0.52771, 0.05154,  0.20264,          2,           1,            1
    3,       1,             117,         117.0,  1.082, 0.077,  1.461,  15.505, 2.302,   1.341, 0.125,    0.53594, 0.04716,  0.06271,          0,           0,            0
    4,       1,              10,          10.0,  1.061, 0.056,  1.169,  13.042, 1.045,   1.212, 0.086,    0.49458, 0.02408,  0.00495,          0,           0,            0
    5,       1,               1,           1.0,  1.007, 0.000,  1.007,  18.971, 0.000,   1.180, 0.000,    0.40153, 0.00000,  0.00040,          0,           0,            0
    7,       1,               1,           1.0,  1.041, 0.000,  1.041,  24.420, 0.000,   1.653, 0.000,    0.32229, 0.00000,  0.00032,          0,           0,            0
Total number of unassigned events:                                                                                                                     3,           2,            1
```

***Legend:***

*SC_ID* – a label of a given supercluster, which to some extend follows their relevance.
*No_Sims* – the number of simulations contributing some tunnels to a supercluster.
*Total_No_Frames* – the number of snapshots with at least one tunnel satisfying the active filters.
*Avg_No_Frames* – the number of snapshots with at least one tunnel satisfying the active filters averaged across the contributing MD simulations.
*Avg_BR* – the average bottleneck radius of tunnels in a supercluster.
*Max_BR* – the maximum bottleneck radius of tunnels in a supercluster.
*Avg_Len* – the average length of tunnels in a supercluster.
*Avg_Cur* – the average curvature (tunnel length/distance between the tunnel start and end spheres) of tunnels in a supercluster.
*Priority* – the priority of a given supercluster calculated by averaging the sum of tunnel throughputs over all snapshots and fraction of contributing simulations.
*StDev* – the standard deviation of property in the preceding column.

***Example of statistics on bottlenecks of tunnels forming superclusters:***

```
Bottleneck residues frequency for: overall
Total simulations = 2
No snapshots per simulation = 1000

Parameters used for pre-selection of input tunnels for clustering:
length >= 5.00 A
radius >= 0.75 A
curvature <= 2.00

Active tunnel filters:
length = (0.00, 999.00) A
radius = (0.00, 999.00) A
curvature = (1.00, 999.00)
occurred in at least 0 simulations
occurred in at least 0 snapshots
occurred in at least 0.000 snapshots on average per simulation
has at least 1 transport events
has at least 0 entry events
has at least 0 release events

SC_ID, Total_No_Frames, Bottleneck residues: frequencies
    1,            1986, 142:0.977, 173:0.969, 146:0.952, 169:0.946, 141:0.901, 138:0.856,
    2,            1812, 243:0.991, 138:0.976, 132:0.668, 242:0.664, 139:0.610, 206:0.471,
```

***Legend:***

*SC_ID* – a label of a given supercluster, which to some extend follows their relevance.

*Total_No_Frames* – the number of snapshots with at least one tunnel satisfying the active filters.

*Bottleneck residues: frequencies* – a list of residue IDs forming tunnel bottlenecks and the respective frequency with which these residues contribute to such a bottleneck

#### 5.4. *visualization* folder

This folder contains python scripts to visualize analyzed superclusters in PyMOL and all the necessary data stored in the *sources* folder. The PyMOL scripts can be executed either directly from the command line: pymol script_filename or from PyMOL GUI by typing: run script_filename.

**PyMOL scripts for visualization:**

- *1-visualize_initial_tunnels.py* – shows initial superclusters
- *2-visualize_filtered_tunnels.py* – shows superclusters filtered based on tunnels
- *3-visualize_initial_events.py* – shows superclusters filtered based on tunnels as well as visualization of assigned and outlaying events
- *4-visualize_filtered_events.py* – shows superclusters with assigned as well as outlaying events filtered based on both tunnels and events

**Folder with sources for visualization**

- *super_cluster_CGOs* – subfolder with CGO formatted data for visualization of:
  - initial superclusters (*_pathset1.dump.gz)
  - superclusters filtered based on tunnels (*_pathset2.dump.gz)
  - superclusters with assigned events & filtered based on tunnels (*_pathset3.dump.gz)
  - superclusters with assigned events & filtered based on both tunnels and events (*_pathset4.dump.gz)
- corresponding supercluster volume visualizations (*_volume[1-4].dump.gz) [optional if *visualize_super_cluster_volumesvisualize_super_cluster_volumes* = True]
- *layered_data/aquaduct* – subfolder with CGO formatted data for visualization of layered events
- *network_data/caver* – subfolder with files for visualization of transformed networks for each input simulation, including view_network.py scripts for PyMOL [optional if *visualize_transformed_tunnels* = True]

- *network_data/aquaduct* – subfolder with files for visualization of transformed networks for each input simulation, including view_network.py scripts for PyMOL [optional if *visualize_transformed_transport_events* = True]

### 6. Specialized analysis modes

#### 6.1. Comparative analysis

In this analysis mode, the supercluster statistics, as well as their visualization, are provided separately for each input MD simulation. Possibly user-defined groups of simulations can be used besides as specified by the *comparative_groups_definition* parameter. In such a case, separate data for simulations assigned to the groups will not be reported. The typical application scenario for this mode is when comparing the transport processes in different simulation sets, e.g., when contrasting transport in the original system and system perturbed by mutations, different solvents, presence of ligands, simulations under different conditions (T, p, pH, etc.) or different force fields.

***Relevant parameters:***

- *perform_comparative_analysis*
- *comparative_groups_definition*
- *visualize_comparative_super_cluster_volumes*

The format of the group definition follows, also UNIX-like patterns are accepted here:

group_name1: [folder1_1, folder1_2, folder_pattern1_1, folder_pattern1_2]; group_name2: [folder2_1, folder2_2, folder_pattern2]; group_name3 [folder3]

For example:
*comparative_groups_definition* = mutants1: [md*_mut1, md_mut2]; wild_type: [mdwt*]

where mutants1 is the first analyzed group formed by input folders matching pattern *md*_mut1* and folder *md_mut2*, and the second analyzed group wild_type is formed by input folders matching pattern *mdwt**

***Description of outputs:***
The results of comparative analysis are located in *output_path/*statistics/comparative_analysis and *output_path/*visualization/comparative_analysis folders.

```
results_comparative/statistics/comparative_analysis/
├── 1-initial_tunnels_statistics_bottleneck_residues.txt
├── 1-initial_tunnels_statistics.txt
├── 2-filtered_tunnels_statistics_bottleneck_residues.txt
├── 2-filtered_tunnels_statistics.txt
├── 3-initial_events_statistics_bottleneck_residues.txt
├── 3-initial_events_statistics.txt
├── 4-filtered_events_statistics_bottleneck_residues.txt
├── 4-filtered_events_statistics.txt
├── traj1
│   ├── 1-initial_tunnels_statistics_bottleneck_residues.txt
│   ├── 1-initial_tunnels_statistics.txt
│   ├── 2-filtered_tunnels_statistics_bottleneck_residues.txt
│   ├── 2-filtered_tunnels_statistics.txt
│   ├── 3-initial_events_statistics_bottleneck_residues.txt
│   ├── 3-initial_events_statistics.txt
│   ├── 4-filtered_events_statistics_bottleneck_residues.txt
│   └── 4-filtered_events_statistics.txt
└── traj2
```

```
results_comparative/visualization/comparative_analysis/
├── traj1
│   ├── 1-visualize_initial_tunnels.py
│   ├── 2-visualize_filtered_tunnels.py
│   ├── 3-visualize_initial_events.py
│   └── 4-visualize_filtered_events.py
└── traj2
```

In the *output_path/*statistics/comparative_analysis folder tunnels statistics (initial and filtered) and events statistics (initial and filtered) files are generated and contain merged data of statistics for each of the groups. Additionally, a separate folder for each group is created with statistics files only for the particular group. All these files are structured as standard files generated in *statistics* folder (as described in chapter 5.3) but differ in the data used for their generation (by group).

In the *output_path/*visualization/comparative_analysis folder, the User can find separate folders for each of the defined groups. These folders contain visualization scripts equivalent to standard visualization scripts present in *visualization* folder (as described in chapter 5.4) but their content is limited to the data for a particular group only. Optionally [if *visualize_comparative_super_cluster_volumes* = True], volumes of superclusters in each analyzed groups can be visualized too.

#### 6.2. Exact matching of transport events and molecular tunnels

Here, exact analyses of an event with respect to tunnel clusters in frames during the event occurrence originating directly from the simulation sampling the event is performed to learn which tunnels were utilized by transported molecule during the transport event and obtain the data on the property of those tunnels. Optionally, all matched events could be visualized depending on *visualize_exact_matching_outcomes* settings. Note that enabling the visualization will result in markedly increased demands on memory as well as storage space.

***Relevant parameters:***

- *perform_exact_matching_analysis*
- *visualize_exact_matching_outcomes*

***Description of outputs:***

The results of the exact matching analysis are located in *output_path/*data/exact_matching_analysis and [if *visualize_exact_matching_outcomes* = True] also in *output_path/*visualization/exact_matching_analysis folders.

```
├── _internal
├── data
│   ├── clustering
│   ├── super_clusters
│   └── exact_matching_analysis
├── statistics
└── visualization
    ├── exact_matching_analysis
    └── sources
```

In the *output_path/*data/exact_matching_analysis folder, for each analyzed MD simulation, plain-text files with assignment details, named as (EventID)_(Event_type)_sc(SuperclusterID).txt, for example, *1_entry_sc2.txt*, are generated.

***Example:***

```
Exact matching of transport event '('md1', '1_entry', ('WAT:326', (144, 893)))' to tunnels from Super cluster 2
fraction of frames in which ligand is inside SC tunnels 0.86
fraction of frames in which SC tunnels exist and the ligand is inside of them 0.91
Data on the tunnel spheres closest to the ligand:
-----------------------------------------------
     Frame,X-coordLig,Y-coordLig,Z-coordLig,  Dist2lig,CaverClsID,X-coordSph,Y-coordSph,Z-coordSph,   Dist2SP,    Radius, TunLength,
       144,    -1.586,     6.456,   -14.271,     2.627,         3,    -1.415,     3.583,   -10.855,    11.468,     1.840,    15.339
       145,    -2.164,     5.239,   -14.729,     2.791,         3,    -2.193,     2.369,   -11.776,    12.166,     1.327,    18.621
       146,    -1.279,     5.770,   -11.684,     0.375,         3,    -1.018,     4.149,   -10.643,    11.416,     1.569,    18.340
       147,    -1.354,     4.712,   -11.891,    -0.080,         3,    -1.046,     4.015,   -10.478,    11.217,     1.686,    14.969
       148,    -1.170,     5.064,   -11.949,    -0.755,         3,    -1.406,     4.052,   -11.523,    12.244,     1.879,    17.292
       149,    -1.289,     5.351,   -11.683,     0.101,         3,    -1.175,     4.215,   -10.789,    11.589,     1.350,    22.930
       151,    -1.222,     4.235,   -10.128,    -0.742,         2,    -1.558,     4.172,   -10.303,    11.168,     1.127,    21.514
```

***Legend:***
After a header specifying the assignment of which event and supercluster are reported, information on the overall buriedness of the event in the supercluster is provided, followed by the details on the closest sphere of tunnels belonging in the supercluster and the full trajectory of ligand responsible for the event.
*Frame* – a current trajectory frame belonging to the analyzed transport event.
*X-coordLig* – x-coordinate of any ligand atom closest to any tunnels' sphere.
*Y-coordLig* – y-coordinate of any ligand atom closest to any tunnels' sphere.
*Z-coordLig* – z-coordinate of any ligand atom closest to any tunnels' sphere.
*Dist2lig* – the shortest distance between surfaces of any ligand atom and tunnels' sphere. Note that negative values mean overlap in surfaces. As a rule of thumb, the distances < 1 Å might be indicative of the ligand's burial in a given tunnel.
*CaverClsID* – the ID of a tunnel cluster to which the tunnel's sphere closest to any ligand atom belongs.
*X-coordSph* – x-coordinate of the tunnel's sphere closest to any ligand atom.
*Y-coordSph* – y-coordinate of the tunnel's sphere closest to any ligand atom.
*Z-coordSph* – z-coordinate of the tunnel's sphere closest to any ligand atom.
*Dist2SP* – the shortest possible distance between the sphere closest to any ligand atom and the tunnel starting point.
*Radius* – radius of the tunnel's sphere closest to any ligand atom.
*TunLength* – length of the tunnel at the sphere closest to any ligand atom.

In the *output_path/* visualization /exact_matching_analysis folder, for each analyzed MD simulation, a set of subfolders for each analyzed event will be generated. These subfolders are named as (EventID)_(Event_type)_sc(SuperclusterID); for example, *1_entry_sc2,* and contain a python script for visualization of assignments in PyMOL (*show_matched_event.py*) and all the necessary source data.
