## Supplementary File 4 for "TransportTools: a library for high-throughput analyses of internal voids in biomolecules and ligand transport through them"

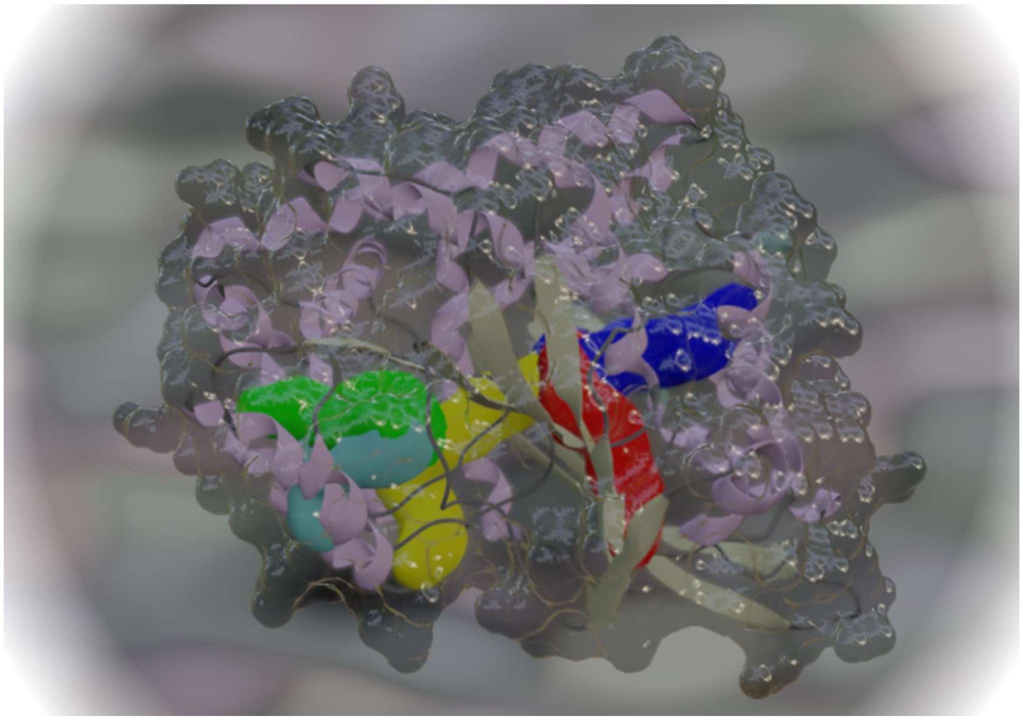

### TransportTools 0.9.0

***A library for massive analyses of internal voids in biomolecules and ligand transport through them***

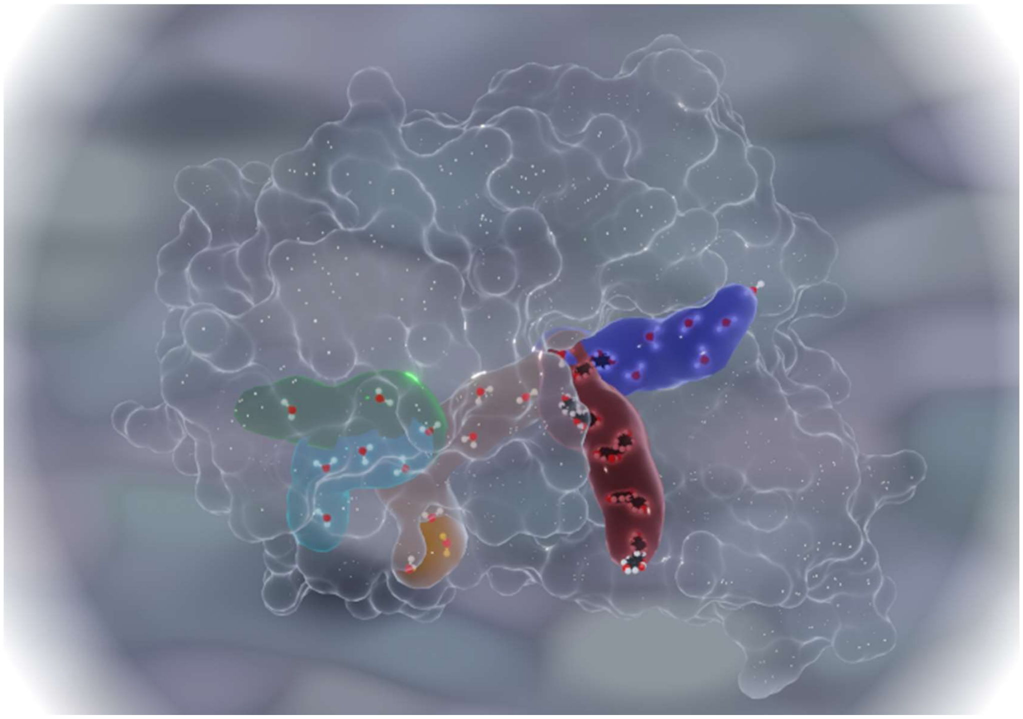

#### TECHNICAL DOCUMENTATION

### Namespace transport_tools

#### Sub-modules

- transport_tools.libs
- transport_tools.scripts

### Module transport_tools.libs

#### Sub-modules

- transport_tools.libs.config
- transport_tools.libs.geometry
- transport_tools.libs.networks
- transport_tools.libs.protein_files
- transport_tools.libs.tools
- transport_tools.libs.ui
- transport_tools.libs.utils

### Module transport_tools.libs.config

#### Classes

##### Class AnalysisConfig

```
class AnalysisConfig(
    file2load_from: Optional[str] = None,
    logging: bool = True
)
```

Class storing the job configuration, also enables some parameter evaluation & completion
:param file2load_from: INI file with configuration to load from
:param logging: if logging should be set up

###### Methods

###### Method get_filters

```
def get_filters(
    self
) -> List[Union[float, int]]
```

Collect filter values properly ordered to act as argument for filter_super_cluster_profiles and define_filters
:return: list of filter values

###### Method get_input_folders

```
def get_input_folders(
    self
) -> (typing.List[str], typing.List[str], typing.List[str])
```

Enumerates source folders with CAVER data, MD trajectories and AQUA-DUCT data separately
:return: list of source folders with CAVER data, MD trajectories, and AQUA-DUCT data, respectively

###### Method get_parameter

```
def get_parameter(
    self,
    par_name: str
)
```

Return values for query parameter
:param par_name: parameter name

###### Method get_parameters

```
def get_parameters(
    self
) -> dict
```

Return compiled parameters of the job

###### Method get_reference_pdb_file

```
def get_reference_pdb_file(
    self
) -> str
```

Returns path to the reference PDB file that will define transformations for the job,
unless user specified, it is the file from first CAVER folder

###### Method `report_updates`

```
def report_updates(
    self,
    old_parameters: dict
)
```

Logs information on modified parameters upon restart
:param old_parameters: previous parameters to compare with

###### Method `set_parameter`

```
def set_parameter(
    self,
    par_name: str,
    value: Any
)
```

Set new value for query parameter
:param par_name: parameter name
:param value: new value of the par_name parameter

###### Method `write_template_file`

```
def write_template_file(
    self,
    filepath: str,
    advanced: bool = False
)
```

Save job configuration template with default values to a file
:param filepath: path to INI file to which the configuration template will be saved
:param advanced: write also advanced section

### Module `transport_tools.libs.geometry`

#### Functions

###### Function `assign_layer_from_distances`

```
def assign_layer_from_distances(
    distances: np.array,
    layer_thickness: float
) -> (np.array, np.array)
```

Calculate membership of points in layers based on point distance form starting point and the layer thickness
:param distances: distance of points from starting point
:param layer_thickness: layer thickness
:return: unique IDs of layers and membership of points in layers

###### Function `average_starting_point`

```
def average_starting_point(
    tunnel_origin_file: str,
```

```
    md_label: str = ''
) -> (np.array, str)
```

Computes the average coordinates of starting points from origin_file
:param tunnel_origin_file: file with caver starting points
:param md_label: name of folder with the source MD simulation data
:return: array containing the average starting point coordinates & md_label

**Function** `cart2spherical`

```
def cart2spherical(
    xyz: np.array
) -> <built-in function array>
```

Converts cartesian coordinates to spherical ones
:param xyz: cartesian coords
:return: spherical coords

**Function** `einsum_dist`

```
def einsum_dist(
    xyz1: np.array,
    xyz2: np.array
)
```

Calculates distance between two points in computationally rather efficient manner
:param xyz1: coordinates of the first point
:param xyz2: coordinates of the second point
:return:

**Function** `get_coarse_grained_path`

```
def get_coarse_grained_path(
    point2cluster_map: Dict[int, Dict[int, str]],
    path_id: int
) -> List[str]
```

Node path formed by nodes labels along the original path
:param point2cluster_map:
:param path_id: ID of original path that is being coarse-grained
:return: path consisting of node labels

**Function** `get_layer_id_from_distance`

```
def get_layer_id_from_distance(
    distances: np.array,
    layer_thickness: float
) -> <built-in function array>
```

Calculate layers for points based on their distances form starting point and the layer thickness
:param distances: distance of points from starting point
:param layer_thickness: layer thickness
:return: layerIDs

**Function** `get_redundant_path_ids`

```
def get_redundant_path_ids(
    all_paths: Dict[int, List[str]]
) -> Set[int]
```

Detects node paths that are subset of another node paths from all_paths
:param all_paths: paths to analyze
:return: set of IDs of redundant paths that are subsets of others

**Function** `remove_loops_from_path`

```
def remove_loops_from_path(
    node_path: List[str]
) -> Optional[List[str]]
```

Removes loops (repetitively visited nodes) from the node path while guaranteeing the largest span of layers
:param node_path: analyzed node path = list of node labels
:return: node path without loops

**Function** `vector_angle`

```
def vector_angle(
    v1: np.array,
    v2: np.array
) -> float
```

Returns angle between two vectors
:param v1: input vector
:param v2: input vector
:return: angle

#### Classes

**Class** `ClusterInLayer`

```
class ClusterInLayer(
    points_mat: np.array,
    thickness: float,
    quantile: float,
    end_point: bool = False,
    cls_id: int = -1,
    layer_id: int = -1
)
```

Clusters representing original points in layers
:param cls_id: ID of this cluster in a given layer
:param layer_id: ID of layer to which this cluster belongs
:param points_mat: data of included points
:param thickness: layer thickness
:param quantile: to use for calculation of representative radius of this cluster
:param end_point: does cluster represent end point of original tunnel or transport event

**Methods**

**Method** `compute_averages`

```
def compute_averages(
    self
)
```

Compute average coordinates, rmsf, radius for this cluster, determine if it contains starting or end points

**Method** `get_coords`

```
def get_coords(
    self
) -> <built-in function array>
```

Return coordinates of all points in this cluster

**Method** `get_node_label`

```
def get_node_label(
    self
) -> str
```

Create label node representing this cluster from its layer and cls IDs
:return: node label

**Method** `is_representative`

```
def is_representative(
    self,
    tolerance: float = 0
) -> bool
```

Test if cluster average represents well the original tunnel points
:param tolerance: additional tolerance

**Method** `merge_with_cluster`

```
def merge_with_cluster(
    self,
    other_cluster: ClusterInLayer
)
```

Join other cluster to this cluster
:param other_cluster: Cluster to add

**Method** `resolve_avg_failures`

```
def resolve_avg_failures(
    self,
    random_seed: int,
    num_new_clusters: int = 0,
    max_new_clusters: Optional[int] = None,
    tolerance: float = 0
) -> List[transport_tools.libs.geometry.ClusterInLayer]
```

Keep splitting this cluster to two, until all created clusters represents well their points or the maximum number of new clusters is created
:param random_seed: value to initiate the random number generator
:param num_new_clusters: number of newly created clusters so far during the resolution process
:param max_new_clusters: limit on the maximum of newly created clusters

:param tolerance: additional tolerance for testing the representativeness
:return: list of representative clusters

###### Class `Layer`

```
class Layer(
    layer_id: int,
    layer_thickness: float,
    parameters: dict,
    entity_label: str,
    md_label: str
)
```

Class storing layered points; enable clustering of these points to Clusters
:param layer_id: ID of this layer
:param layer_thickness: layer thickness
:param parameters: job configuration parameters
:param entity_label: name of the entity (tunnel cluster or transport event) to be layered

###### Descendants

- transport_tools.libs.geometry.Layer4Events
- transport_tools.libs.geometry.Layer4Tunnels

###### Methods

###### Method `add_cluster`

```
def add_cluster(
    self,
    cluster: ClusterInLayer
)
```

Adds new cluster to this layer, making sure it adheres to the layer specification
:param cluster: cluster to add
:return: id of the new cluster

###### Method `cluster_data`

```
def cluster_data(
    self,
    points_mat: np.array
)
```

Perform clustering of data of points assigned to this layer, separately for common points and end points, finally representativeness of formed clusters is enforced
:param points_mat: points to cluster

###### Method `contains_data`

```
def contains_data(
    self
) -> bool
```

Test if layer contain any data (has been assigned any point)

###### Method pop_cluster

```
def pop_cluster(
    self,
    cls_id: int
) -> transport_tools.libs.geometry.ClusterInLayer
```

Pops a required cluster with given id from this layer
:param cls_id: id of the required cluster
:return: the required cluster

###### Method save_points

```
def save_points(
    self,
    out_folder: str,
    save_points_prefix: str,
    transform_mat: np.array
) -> List[str]
```

Save clusters (nodes) and their constituent points for visualization
:param out_folder: folder to which PDB files with nodes will be saved
:param save_points_prefix: prefix for names of PDB file defining nature of layered entity
:param transform_mat: transformation matrix to transform output points
:return: list of names of created PDB files

##### Class Layer4Events

```
class Layer4Events(
    layer_id: int,
    layer_thickness: float,
    parameters: dict,
    entity_label: str,
    md_label: str
)
```

Class storing layered points; enable clustering of these points to Clusters
:param layer_id: ID of this layer
:param layer_thickness: layer thickness
:param parameters: job configuration parameters
:param entity_label: name of the entity (tunnel cluster or transport event) to be layered

###### Ancestors (in MRO)

- transport_tools.libs.geometry.Layer

##### Class Layer4Tunnels

```
class Layer4Tunnels(
    layer_id: int,
    layer_thickness: float,
    parameters: dict,
    entity_label: str,
    md_label: str
)
```

Class storing layered points; enable clustering of these points to Clusters
:param layer_id: ID of this layer
:param layer_thickness: layer thickness

:param parameters: job configuration parameters
:param entity_label: name of the entity (tunnel cluster or transport event) to be layered

###### Ancestors (in MRO)

- transport_tools.libs.geometry.Layer

##### Class `LayeredPathSet`

```
class LayeredPathSet(
    entity_label: str,
    md_label: str,
    parameters: dict,
    starting_point_coords: np.matrix = None
)
```

Class for manipulation and analyses of simplified set of paths designed to represent original tunnel clusters or transport events
:param entity_label: name of the entity, e.g., Cluster_1, Path_2 ...
:param md_label: name of folder with the source MD simulation data
:param parameters: job configuration parameters
:param starting_point_coords: coordinates of average starting point for this simulation

###### Methods

###### Method `add_node_path`

```
def add_node_path(
    self,
    node_path: List[str],
    layers: Dict[int, Layer]
)
```

Add path of node labels representing the original entity (tunnel cluster or transport event) and the data for clusters visited along the path to this pathset
:param node_path: path formed by node labels to add
:param layers: layers containing processed clusters

###### Method `avg_distance2path_set`

```
def avg_distance2path_set(
    self,
    other_set: LayeredPathSet,
    distance_cutoff: float = 999,
    dist_type: int = 1
) -> float
```

Compute mean closest surface-to-surface distance of all paths from two pathsets
Note that if during calculation the mean distance is projected to surpass distance cutoff, the two patsets are deemed faraway with 999 distance. The same distance is also assumed for directionally misaligned patset pairs.
:param other_set: other set to which the distance is calculated
:param distance_cutoff: cutoff on accurate distance calculation, anything beyond this value is far (999)
:param dist_type: 0 - center distances, 1 - surface distances, 2 - surface and rmsf distances
:return: mean distance (surface-to-surface by default) between pathsets

**Method** `compute_node_depths`

```
def compute_node_depths(
    self
)
```

Assign average depth along the paths to each node

**Method** `get_fragmented_paths`

```
def get_fragmented_paths(
    self
) -> Tuple[int, List[List[List[str]]]]
```

Compile path fragment's for all paths forming this pathset
:return: number of fragmented paths, list of fragmented paths

**Method** `how_much_is_inside`

```
def how_much_is_inside(
    self,
    other_set: LayeredPathSet
) -> (float, float)
```

Computes the fraction of nodes from this set buried inside the nodes of the other set, and maximal depth (counted towards starting point (SP) along shortest path)
:param other_set: other set in which the buriedness is calculated
:return: buriedness, and maximal depth towards SP

**Method** `is_empty`

```
def is_empty(
    self
) -> bool
```

If this pathset contain no valid node_paths

**Method** `is_same`

```
def is_same(
    self,
    other: LayeredPathSet
) -> bool
```

Test if the pathset is same as the other pathset
:param other: other pathset to compare with
:return: are two pathset same?

**Method** `remove_duplicates`

```
def remove_duplicates(
    self
)
```

Remove duplicate nodes and paths from a merged pathset to speed-up the assignment of other pathsets

**Method** `remove_unnecessary_paths`

```
def remove_unnecessary_paths(
    self
)
```

Remove paths that do lead to already explored terminal nodes, and do not include any additional unexplored node

**Method** `set_traced_event`

```
def set_traced_event(
    self,
    traced_residue: Tuple[str, int, Tuple[int, int], Tuple[int, int]]
)
```

Sets information on residue and frames traced by AQUA-DUCT that is source for this layered event
:param traced_residue: tuple containing resname & resid of ligand responsible for this path,
and beginning and last frames for entry and release events

**Method** `transform_coordinates`

```
def transform_coordinates(
    self,
    transform_mat: np.array
)
```

Transform nodes coordinates forming this pathset according to the transformation matrix
:param transform_mat: transformation matrix to be applied on the nodes coordinates

**Method** `visualize_cgo`

```
def visualize_cgo(
    self,
    output_folder: str,
    entity_label: str,
    color_id: int = 0,
    merged: bool = False,
    flag: str = '',
    surface_cgo: bool = False
)
```

Save this pathset as Pymol compiled graphics object(CGO) for visualization
:param output_folder: folder to which CGO will be saved
:param entity_label: name of the layered entity (tunnel cluster or transport event) to visualize
:param color_id: Pymol ID of color to use for this pathset
:param merged: if all paths should be in single CGO
:param surface_cgo: if to generate also surface visualization
:param flag: additional description enabling differentiation of cgo files among various results after filtering

**Class** `LayeredRepresentation`

```
class LayeredRepresentation(
    parameters: dict,
    entity_label: str
)
```

Class responsible for splitting of original data to layers and defining of layered (coarse-grained) paths
representing the original entity (tunnel cluster or transport event)
:param parameters: job configuration parameters
:param entity_label: name of the entity to be layered

###### Descendants

- transport_tools.libs.geometry.LayeredRepresentationOfEvents
- transport_tools.libs.geometry.LayeredRepresentationOfTunnels

###### Methods

###### Method `find_representative_paths`

```
def find_representative_paths(
    self,
    transform_mat: np.array,
    starting_point_coord: np.array,
    visualize: bool = False
) -> transport_tools.libs.geometry.LayeredPathSet
```

###### Method `load_points`

```
def load_points(
    self,
    source_entity: Union[Tunnel, TransportEvent]
)
```

Load data of points from the original entity (tunnel cluster or transport event)
:param source_entity: the original entity from which data is to be extracted

###### Method `prep_visualization`

```
def prep_visualization(
    self,
    layered_paths: List[str],
    transform_mat: np.array,
    show_original_data: bool = False
)
```

Prepare visualization of the layered representation - clusters (nodes) and representative paths
:param layered_paths: analyzed node path = list of node labels
:param transform_mat: transformation matrix to transform output pathset
:param show_original_data: should original data be visualized too

##### Class `LayeredRepresentationOfEvents`

```
class LayeredRepresentationOfEvents(
    parameters: dict,
    entity_label: str
)
```

Class responsible for splitting of transport event data to layers and defining of their representative paths
:param parameters: job configuration parameters
:param entity_label: name of the entity (transport event) to be layered

###### Ancestors (in MRO)

- transport_tools.libs.geometry.LayeredRepresentation

###### Methods

###### Method find_representative_paths

```
def find_representative_paths(
    self,
    transform_mat: np.array,
    starting_point_coords: np.array = None,
    visualize: bool = False
) -> transport_tools.libs.geometry.LayeredPathSet
```

Find representative paths leading from starting point to terminal clusters (nodes)
:param transform_mat: transformation matrix to transform output pathset
:param starting_point_coords: NOT USED
:param visualize: should the layered representation be prepared for visualization
:return: set of representative paths

###### Method split_points_to_layers

```
def split_points_to_layers(
    self
)
```

Splits points to Layers and execute their per layer clustering

##### Class LayeredRepresentationOfTunnels

```
class LayeredRepresentationOfTunnels(
    parameters: dict,
    entity_label: str
)
```

Class responsible for splitting of tunnel cluster data to layers and defining of their representative paths
:param parameters: job configuration parameters
:param entity_label: name of the entity (tunnel cluster) to be layered

###### Ancestors (in MRO)

- transport_tools.libs.geometry.LayeredRepresentation

###### Methods

###### Method find_representative_paths

```
def find_representative_paths(
    self,
    transform_mat: np.array,
    starting_point_coords: np.array,
    visualize: bool = False
) -> transport_tools.libs.geometry.LayeredPathSet
```

Find representative paths leading from starting point to terminal clusters (nodes)
:param transform_mat: transformation matrix to transform output pathset
:param starting_point_coords: coordinates of average starting point for this simulation
:param visualize: should the layered representation be prepared for visualization
:return: set of representative paths

**Method** `split_points_to_layers`

```
def split_points_to_layers(
    self
)
```

Splits points to Layers and execute their per layer clustering

**Class** `Point`

```
class Point(
    xyz,
    distance=-1,
    radius=0
)
```

Class for point storing and manipulation
:param xyz: coordinates
:param distance: distance to arbitrary point
:param radius: radius of allocated space

**Methods**

**Method** `convert2viz_atom`

```
def convert2viz_atom(
    self,
    atom_id: int,
    res_id: int,
    resname: str = 'UNK'
) -> transport_tools.libs.protein_files.VizAtom
```

Convert points to VizAtom object for saving PDB lines
:param atom_id: ID of atom in PDB record
:param res_id: ID of residue in PDB record
:param resname: name of residue in PDB record
:return: VizAtom object

**Method** `distance2point`

```
def distance2point(
    self,
    other: Point
) -> float
```

Calculate distance to another point
:param other: point to which we calculated the distance
:return: distance between the points

**Method** `save_point`

```
def save_point(
    self,
    filename
)
```

Save point as PDB line to file
:param filename: file to which we save the point

##### Class `PointMatrix`

```
class PointMatrix(
    points_mat: np.array
)
```

Class to handle points data
:param points_mat: input points data

###### Methods

###### Method `alter_coords`

```
def alter_coords(
    self,
    new_xyz: np.array
) -> <built-in function array>
```

Updates current coordinates of points with new set of coordinates
:param new_xyz: new coordinates use

###### Method `get_coords`

```
def get_coords(
    self
) -> <built-in function array>
```

Return coordinates of all points in Matrix

###### Method `get_end_points_indexing`

```
def get_end_points_indexing(
    self
) -> <built-in function array>
```

Return indices of points representing end of tunnels or events

###### Method `get_num_columns`

```
def get_num_columns(
    self
) -> int
```

Compute number of columns this matrix has

###### Method `get_num_points`

```
def get_num_points(
    self
) -> int
```

Compute number of points this matrix contain

###### Method `get_points_ids4tunnel`

```
def get_points_ids4tunnel(
    self,
    tunnel_id: int
) -> <built-in function array>
```

Return IDs of points for particular tunnel(event)
:param tunnel_id: ID of query tunnel

**Method** `get_radii`

```
def get_radii(
    self
) -> <built-in function array>
```

Return radii of all points in Matrix

**Method** `get_start_points_indexing`

```
def get_start_points_indexing(
    self
) -> <built-in function array>
```

Return indices of points representing start of tunnels or events

**Method** `get_tunnels_ids`

```
def get_tunnels_ids(
    self
) -> <built-in function array>
```

Return IDs of all tunnels(events) in Matrix

**Method** `get_whole_matrix`

```
def get_whole_matrix(
    self
) -> <built-in function array>
```

Return numpy representation of this matrix

**Method** `is_empty`

```
def is_empty(
    self
) -> bool
```

Test if this matrix is empty

### Module `transport_tools.libs.networks`

#### Functions

**Function** `define_filters`

```
def define_filters(
    min_length: float = -1,
    max_length: float = -1,
    min_bottleneck_radius: float = -1,
    max_bottleneck_radius: float = -1,
    min_curvature: float = -1,
    max_curvature: float = -1,
    min_sims_num: int = -1,
    min_snapshots_num: int = -1,
```

```
        min_avg_snapshots_num: float = -1,
        min_total_events: int = -1,
        min_entry_events: int = -1,
        min_release_events: int = -1
    )
```

Defines filters to be used for filtering; keeping them in acceptable ranges
NOTE: -1 => filter is not active
:param min_length: minimum tunnel length
:param max_length: maximum tunnel length
:param min_bottleneck_radius: minimum tunnel bottleneck radius
:param max_bottleneck_radius: maximum tunnel bottleneck radius
:param min_curvature: minimum tunnel curvature
:param max_curvature: maximum tunnel curvature
:param min_sims_num: present in minimum number of MD simulations
:param min_snapshots_num: present in minimum number of snapshots
:param min_avg_snapshots_num: present in minimum number of snapshots on average
:param min_total_events: having minimum transport events
:param min_entry_events: having minimum entry events
:param min_release_events: having minimum release events

###### **Function** `get_md_membership4groups`

```
def get_md_membership4groups(
    comparative_groups_definition: Dict[str, List[str]]
) -> Dict[str, str]
```

Converts group definitions to membership of MD simulations to groups
:param comparative_groups_definition: definition of groups and belonging MD simulations for comparative analyses
:return: membership of MD simulations to groups

###### **Function** `subsample_events`

```
def subsample_events(
    transport_events: Dict[str, List[Tuple[str, Tuple[str, Tuple[int, int]]]]],
    random_seed: int,
    max_events: int,
    md_label: str = 'overall',
    comparative_groups_definition: Optional[Dict[str, List[str]]] = None
) -> List[Tuple[str, str]]
```

Randomly selects limited number of transport events for visualization
:param transport_events: evaluated information about transport events
:param random_seed: value to initiate the random number generator
:param max_events: maximum number of events to keep in supercluster
:param md_label: subsampling from which simulations to prepare; by default ‘overall’ from all
:param comparative_groups_definition: definition of groups and belonging MD simulations for comparative analyses
:return: retained information on transport events

##### Classes

###### **Class** `AquaductNetwork`

```
class AquaductNetwork(
    parameters: dict,
    md_label: str,
    load_only: bool = False
```

```
)
```

Class for processing of AQUA-DUCT results, containing AquaductPath objects
:param parameters: job configuration parameters
:param md_label: name of folder with the source MD simulation data
:param load_only: object will be used to load already processed network, no need for pdb_file

###### Ancestors (in MRO)

- transport_tools.libs.networks.Network

###### Methods

###### Method `clean_tempfile`

```
def clean_tempfile(
    self
)
```

Removes tempfile

###### Method `get_events4layering`

```
def get_events4layering(
    self
) -> List[transport_tools.libs.networks.TransportEvent]
```

Get events in this aqua network for layering
:return: list of events for processing

###### Method `get_pdb_file`

```
def get_pdb_file(
    self
)
```

Create temporary PDB file with protein structure from AQUA-DUCT tarfile

###### Method `read_raw_paths_data`

```
def read_raw_paths_data(
    self,
    parallel_processing: bool = True
)
```

Parallel processing of all raw_paths present in AQUA-DUCT tarfile to create corresponding AquaductPath object
:param parallel_processing: if we process the raw_paths in parallel

##### Class `AquaductPath`

```
class AquaductPath(
    path_label: str,
    parameters: dict,
    traced_residue: Tuple[str, int, Tuple[int, int], Tuple[int, int]],
    transform_mat: np.array,
    md_label: str = ''
)
```

Class for processing of transport paths produced by AQUA-DUCT, consisting of transport events
:param path_label: the name of this path derived from AQUA-DUCT raw paths names
:param parameters: job configuration parameters
:param traced_residue: tuple containing resname & resid of ligand responsible for this path,
and beginning and last frames for entry and release event
:param transform_mat: transformation matrix to be applied on the input coordinates
:param md_label: name of folder with the source MD simulation data

**Methods**

**Method `get_events4layering`**

```
def get_events4layering(
    self
) -> List[transport_tools.libs.networks.TransportEvent]
```

Get events in this path for layering
:return: list of events for processing

**Method `has_transport_event`**

```
def has_transport_event(
    self
) -> bool
```

Test if entry or release type event is among events forming this path

**Method `is_same`**

```
def is_same(
    self,
    other: AquaductPath
) -> bool
```

Test if the path is same as the other path
:param other: other path to compare with
:return: are two paths same?

**Method `process_path`**

```
def process_path(
    self,
    cgo_object: list
)
```

Process Compiled Graphics Object (CGO) representing a raw path from AQUA-DUCT to get initial transport events;
these events are then processed as follows:
1) Merge continuous sequences of inside events => max 3 events (inside, entry and release) in a single path
2) Exclude events that does not reach close enough to the site of interest (starting point (SP))
3) Extend transit events (entry and release) by points from inside events that are continuously closer to SP
4) Extend transit events by overlapping inside points that form a shortest direct path to SP or close to it
:param cgo_object: AQUA-DUCT CGO containing info on a raw path

**Method `save_cgo`**

```
def save_cgo(
    self,
    out_path: str
)
```

Dump all events as single Pymol compiled graphics object(CGO) to gzipped files for visualization of this path
:param out_path: folder where the CGO should be dumped to

###### Class `CumulativeTunnelProfile4SuperCluster`

```
class CumulativeTunnelProfile4SuperCluster(
    sc_id: int,
    tunnel_clusters: Dict[str, List[int]],
    parameters: dict
)
```

Class to store and process the cumulative tunnel profile of all tunnels belonging to caver clusters
from parent supercluster (SC), there is a single such profile per SC
:param sc_id: initial/original ID of parent SC
:param tunnel_clusters: mapping of clusters belonging to particular MD simulation
:param parameters: job configuration parameters

###### Methods

###### Method `filter_clusters`

```
def filter_clusters(
    self,
    active_filters: dict,
    total_num_md_sims: int = 1
) -> Dict[str, List[int]]
```

Filtering of tunnels and their clusters and respective MD simulations; those clusters and MDs that do not
retain any tunnel are removed from this cumulative tunnel profile; possibly flagging whole SC as
failed (self.passed_filter = False)
:param active_filters: filters to be applied (created by define_filters() function )
:param total_num_md_sims: number of all input MD simulation
:return: mapping of tunnel clusters from MD simulations that are valid after filtering

###### Method `get_bottleneck_residues_frequency`

```
def get_bottleneck_residues_frequency(
    self
) -> Dict[str, Dict[str, float]]
```

Compute overall bottleneck residues frequency of supercluster (SC) based on its cumulative bottleneck data
NOTE that this assumes residue numbering equivalency across analyzed simulations
:return: bottleneck residues frequency of SC

###### Method `get_properties`

```
def get_properties(
    self,
    total_num_md_sims: int = 1
) -> Dict[str, Dict[str, Union[float, int]]]
```

Compute overall properties of parent supercluster (SC) based on the cumulative tunnel profiles
:param total_num_md_sims: number of all input MD simulation, not only in this SC
:return: overall properties of parent SC

**Method** `get_property_time_evolution_data`

```
def get_property_time_evolution_data(
    self,
    property_name: str,
    md_labels: List[str],
    missing_value_default: float = 0
) -> Dict[str, <built-in function array>]
```

For each MD simulation return array containing values of given tunnel property for each simulation frame
:param property_name: name of property to extract
:param md_labels: list of all MD simulations contributing to the original supercluster
:param missing_value_default: value to be used for frames where tunnels are missing or invalid in this cluster
:return: mapping of tunnel property values for each MD simulation

**Method** `has_no_tunnels`

```
def has_no_tunnels(
    self
)
```

Test if this profile has no valid tunnels

**Method** `load_networks`

```
def load_networks(
    self
) -> Dict[str, List[int]]
```

Load all tunnel profiles for all MD simulations that contribute to the parent supercluster(SC)
:return: mapping of tunnel clusters from MD simulations that are valid after keeping only single tunnel per snapshot

**Method** `write_csv`

```
def write_csv(
    self,
    filename: str
)
```

Saves the actual cumulative tunnel profile to CSV file formatted akin to CAVER tunnel profiles
:param filename: path to the CSV formatted file to save the cumulative tunnel profile to

**Method** `write_residues`

```
def write_residues(
    self,
    filename: str
)
```

Saves the actual cumulative bottleneck data to CSV file formatted akin to CAVER bottlenecks
:param filename: path to the CSV formatted file to save the cumulative bottleneck data to

**Class** `Network`

```
class Network(
    parameters: dict,
    md_label: str
)
```

Generic class for handling transport tunnels and paths
:param parameters: job configuration parameters
:param md_label: name of folder with the source MD simulation data

###### Descendants

- transport_tools.libs.networks.AquaductNetwork
- transport_tools.libs.networks.TunnelNetwork

###### Methods

###### Method `add_layered_entity`

```
def add_layered_entity(
    self,
    entity_id: Union[int, str],
    layered_pathset: LayeredPathSet
)
```

Add layered entity to the network
:param entity_id: ID of layered entity
:param layered_pathset: layered entity to add

###### Method `is_layering_complete`

```
def is_layering_complete(
    self,
    entity_ids: List[Union[int, str]]
) -> bool
```

Tests is all original entities to verify existence of their layered counterparts
:param entity_ids: list of IDs of original entities
:return: if all original entities have been layered

###### Method `load_layered_network`

```
def load_layered_network(
    self
)
```

Load pre-computed layered entities (LayeredPathSet representing transport events or tunnel clusters)

###### Method `load_orig_network`

```
def load_orig_network(
    self
)
```

Load pre-computed original entities (AquaductPath, TunnelCluster)

###### Method `save_layered_network`

```
def save_layered_network(
    self
)
```

Dump layered entities (LayeredPathSet representing transport events or tunnel clusters) to files
for later processing

**Method** `save_layered_visualization`

```
def save_layered_visualization(
    self,
    save_pdb_files: bool = False
)
```

Saves CGO files with layered entities (LayeredPathSet representing transport events or tunnel clusters) and optionally also PDBs of transformed protein structure and tunnel starting point
:param save_pdb_files: if the PDB files are to be saved

**Method** `save_orig_network`

```
def save_orig_network(
    self
)
```

Dump original entities (AquaductPath, TunnelCluster) to files for later processing

**Method** `save_orig_network_visualization`

```
def save_orig_network_visualization(
    self
)
```

Saves CGO files with original entities AquaDuctPath, TunnelCluster) and PDBs of transformed protein structure for visualization

**Class** `SuperCluster`

```
class SuperCluster(
    sc_id: int,
    parameters: dict,
    total_num_md_sims: int
)
```

Class for storing and operation on supercluster (SC) created from caver clusters across various MD simulation
:param sc_id: initial/original supercluster ID
:param parameters: job configuration parameters
:param total_num_md_sims: number of all input MD simulation

**Methods**

**Method** `add_caver_cluster`

```
def add_caver_cluster(
    self,
    md_label: str,
    cls_id: int,
    path_set: LayeredPathSet
)
```

Store information on tunnel cluster forming this supercluster (SC)
:param md_label: name of folder with the MD simulation data that contain this cluster
:param cls_id: ID of this cluster
:param path_set: Layered paths representing this cluster

**Method** `add_transport_event`

```
def add_transport_event(
    self,
    md_label: str,
    path_id: str,
    event_type: str,
    traced_event: Tuple[str, Tuple[int, int]]
) -> bool
```

Store information on transport event assigned to this supercluster (SC)
:param md_label: name of folder with the MD simulation data that contain this transport event
:param path_id: AQUA-DUCT path ID of this event
:param event_type: type(release or entry) of this event
:param traced_event: tuple containing identity of ligand responsible for this event,
and beginning and last frames of the event
:return: if the event was locally assigned also to the group/particular md_label

**Method** `compute_distance2transport_event`

```
def compute_distance2transport_event(
    self,
    transport_event: LayeredPathSet
) -> (float, float)
```

Computes the fraction of nodes from Layered path that are buried inside the supercluster, and their maximal depth (counted towards starting point (SP) along shortest path)
:param transport_event: Layered path representing the transport event
:return: path buriedness, max depth towards SP

**Method** `compute_space_descriptors`

```
def compute_space_descriptors(
    self
) -> (int, np.array)
```

Collect all unique nodes and paths from all caver clusters to a single PathSet and compute overall direction in which the end points of supercluster lay
:return: id of supercluster and its average direction

**Method** `count_md_labels4events`

```
def count_md_labels4events(
    self
)
```

Counts how many simulations contributed some event in this supercluster
:return: number of simulations

**Method** `filter_super_cluster`

```
def filter_super_cluster(
    self,
    consider_transport_events: bool,
    active_filters: dict,
    flag: int
) -> (int, Dict[str, Dict[str, Union[float, int]]], Dict[str, Dict[str, float]], Dict[str, List
```

Filtering of tunnels loaded from supercluster (SC) profile dumps generated by process_cluster_profile, recalculating the SC properties, and saving the filtered cumulative profile

Note: due to parallel processing, these SC properties and valid clusters after filtering are assigned to this SC via set_properties() and update_caver_clusters_validity() methods called by TransportProcess.filter_super_clusters() method.
:param consider_transport_events: if filters related to transport events should be used
:param active_filters: filters to be applied (created by define_filters() function)
:param flag: filtering ID for subfolder name to differentiate among various results after different steps
:return: (ID of this SC to to identify SC in TP.filter_super_clusters(), SC properties after tunnel filtering, bottleneck residues frequency, and tunnel clusters that are valid after filtering)

**Method** `get_caver_cluster_ids4md_label`

```
def get_caver_cluster_ids4md_label(
    self,
    md_label: str
) -> List[int]
```

Enumerate cluster IDs for given name of the source MD simulation folder that are valid and belong to this SC
:param md_label: name of folder with the source MD simulation data
:return: list of IDs of valid tunnel clusters for this md_label

**Method** `get_caver_clusters`

```
def get_caver_clusters(
    self,
    md_labels: List[str] = None,
    snap_ids: Optional[List[int]] = None
) -> Dict[str, List[transport_tools.libs.networks.TunnelCluster]]
```

Get tunnel clusters from this SC, possibly filtered for specified Snapshot IDs and particular MD simulations
:param md_labels:
:param snap_ids:
:return: dictionary with list of requested tunnel clusters for each md_label

**Method** `get_caver_clusters_full_labels`

```
def get_caver_clusters_full_labels(
    self
) -> List[str]
```

Generate full names of valid tunnel clusters consisting of foldername of their source MD simulation and their ID
:return: list of full names of valid clusters

**Method** `get_md_labels`

```
def get_md_labels(
    self,
    only_with_transport_events: bool = False
) -> List[str]
```

Enumerate names of folders with the MD simulations (md_labels) contributing with at least one valid tunnel cluster to this supercluster
:consider_transport_events: if the md_labels should be listed also considering assigned events
:return: list of md_labels with valid tunnel clusters

**Method** `get_property_time_evolution_data`

```
def get_property_time_evolution_data(
    self,
    property_name: str,
```

```
        active_filters: dict,
        missing_value_default: float = 0
    ) -> Dict[str, <built-in function array>]
```

For each MD simulation return array containing values of given tunnel property for each simulation frame
:param property_name: name of property to extract
:param active_filters: filters to be applied (created by define_filters() function)
:param missing_value_default: value to be used for frames where tunnels are missing or invalid in this cluster
:return: mapping of tunnel property values for each MD simulation

**Method** `get_summary_line_data`

```
    def get_summary_line_data(
        self,
        print_transport_events: bool = False,
        md_label: str = 'overall'
    ) -> List[str]
```

Generates data for creation of line summarizing overall properties of this supercluster (SC)
:param print_transport_events: if properties related to transport events should be reported
:param md_label: summary of which simulations to report; by default report 'overall' stats
:return: list of items for the summary line

**Method** `has_passed_filter`

```
    def has_passed_filter(
        self,
        consider_transport_events: bool = False,
        active_filters: dict = None
    ) -> bool
```

Test if the supercluster (SC) is valid under conditions defined by the active filters
:param consider_transport_events: if related filters related to transport events should be considered
:param active_filters: active filters (created by define_filters() function) to be evaluated
for transport events

**Method** `is_directionally_aligned`

```
    def is_directionally_aligned(
        self,
        other_direction: np.array
    ) -> bool
```

Test if the supercluster direction is aligned to other_direction within directional_cutoff
:param other_direction: other evaluated direction

**Method** `load_path_sets`

```
    def load_path_sets(
        self
    )
```

**Method** `prepare_visualization`

```
    def prepare_visualization(
        self,
        md_label: str = 'overall',
        flag: str = ''
    ) -> Tuple[List[str], Optional[Tuple[transport_tools.libs.geometry.LayeredPathSet, Tuple[str, s
```

Prepare overall CGO files for visualization of paths representing this supercluster (SC) and generate lines for Pymol visualization script
:param md_label: visualization of which simulations to prepare; by default 'overall' visualization
:param flag: additional description enabling differentiation of cgo files among various results after filtering
:return: lines to load visualization of this SC into Pymol, LayeredPathSet and parameters to generate CGO file

**Method** `process_cluster_profile`

```
def process_cluster_profile(
    self
) -> (int, Dict[str, Dict[str, Union[float, int]]], Dict[str, Dict[str, float]], Dict[str, List
```

Merging of individual caver tunnel profiles into the new one for a single supercluster (SC) and computes SC properties.
Note: due to parallel processing, these SC properties are assigned to this SC via set_properties() method called by TransportProcess.create_super_cluster_profiles() method.
:return: ID of this SC to identify SC in TP.create_super_cluster_profiles(), overall properties of this SC, bottleneck residues frequency, and tunnel clusters that are valid after keeping only single tunnel per snapshot

**Method** `report_details`

```
def report_details(
    self,
    events_assigned: bool
)
```

Prints detailed information on the content of supercluster (SC)
:param events_assigned: were transport events already assigned to decide if to report them
:return: string with SC details

**Method** `set_bottleneck_residue_freq`

```
def set_bottleneck_residue_freq(
    self,
    new_bottleneck_residue_freq: dict
)
```

**Method** `set_properties`

```
def set_properties(
    self,
    new_properties: dict
)
```

**Method** `update_caver_clusters_validity`

```
def update_caver_clusters_validity(
    self,
    retained_tunnel_clusters: Dict[str, List[int]]
)
```

Update info on validity of clusters forming this SC, which is stored in self.tunnel_clusters_valid
:param retained_tunnel_clusters: which clusters from which source MD simulations are valid

**Class** `TransportEvent`

```
class TransportEvent(
    event_type: str,
```

```
        path_label: str,
        parameters: dict,
        md_label: str,
        traced_residue: Tuple[str, int, Tuple[int, int], Tuple[int, int]],
        transform_mat: np.array
    )
```

Creates TransportEvent object - an elemental unit carrying info on a single transport event
:param event_type: AQUA-DUCT type of events: “inside”, “entry”, “release”, “outside”
:param path_label: the name of the path containing this event derived from AQUA-DUCT raw paths names
:param parameters: job configuration parameters
:param md_label: name of folder with the source MD simulation data
:param traced_residue: tuple containing resname & resid of ligand responsible for this path,
and beginning and last frames for entry and release event

**Methods**

**Method** `add_point`

```
    def add_point(
        self,
        point: Point
    )
```

Adds point to the event
:param point: point to add

**Method** `assign_distances2point`

```
    def assign_distances2point(
        self,
        dist_point: Point
    )
```

For all points in this event, calculate distance to dist_point and store it in point.data
:param dist_point: reference point for calculations

**Method** `create_layered_event`

```
    def create_layered_event(
        self
    ) -> Tuple[str, str, transport_tools.libs.geometry.LayeredPathSet]
```

Loads event points into LayeredRepresentation and process them to get the
actual set of Layered paths
:return: set of layered paths representing this event

**Method** `extend_points_back`

```
    def extend_points_back(
        self,
        points2add: List[Point]
    )
```

Add points to the end of this transport event
:param points2add: a list of points to add

**Method** **extend_points_front**

```
def extend_points_front(
    self,
    points2add: List[Point]
)
```

Add points to the beginning of this transport event
:param points2add: a list of points to add

**Method** **get_min_distance**

```
def get_min_distance(
    self
) -> float
```

Find smallest distance to other point stored for all points in this event
:return: the smallest distance

**Method** **get_min_distance2starting_point**

```
def get_min_distance2starting_point(
    self
) -> float
```

Compute minimal distance of points in the event to the overall starting point [0, 0, 0]
:return: minimal distance to starting point

**Method** **get_num_points**

```
def get_num_points(
    self
) -> int
```

Count number of points in this transport event
:return: number of points

**Method** **get_points_data**

```
def get_points_data(
    self
) -> <built-in function array>
```

Convert data of points forming this event, adding info on order of points and identifying the tunnel end point
:return: augmented data for this event suitable for LayeredRepresentation.load_points() method

**Method** **get_visualization_cgo**

```
def get_visualization_cgo(
    self
) -> List[float]
```

Converts event points to Pymol compiled graphics object(CGO) for visualization
:return: CGO of event points

**Method** **has_transition**

```
def has_transition(
    self
) -> bool
```

Test if this event has transition between bulk solvent and active site

**Method** `is_same`

```
def is_same(
    self,
    other: TransportEvent
) -> bool
```

Test if the event is same as the other event
:param other: other event to compare with
:return: are two events same?

**Method** `merge_event`

```
def merge_event(
    self,
    other_event: TransportEvent
)
```

Add points from the other event to the beginning of this transport event
:param other_event: event with points to merge

**Class** `Tunnel`

```
class Tunnel(
    parameters: dict,
    transform_mat: np.array
)
```

Class for processing of tunnels from CAVER - an elemental unit carrying info on a tunnel
:param parameters: job configuration parameters
:param transform_mat: transformation matrix to be applied on the input coordinates

**Methods**

**Method** `create_layered_tunnel`

```
def create_layered_tunnel(
    self,
    entity_label: str = ''
) -> transport_tools.libs.geometry.LayeredRepresentationOfTunnels
```

Loads tunnel spheres(points) into LayeredRepresentation and forwards to the cluster layering
:param entity_label: name of the entity (a cluster to which this tunnel belong) to be layered
:return: raw, unprocessed LayeredRepresentation of the tunnel

**Method** `does_tunnel_pass_filters`

```
def does_tunnel_pass_filters(
    self,
    active_filters: dict
) -> bool
```

Check if this tunnel adhere to the applied filters
:param active_filters: filters to be applied (created by define_filters() function)

**Method** `fill_bottleneck_data`

```
def fill_bottleneck_data(
    self,
    bottleneck_data: List[str],
    transform_mat: np.array = None
)
```

Process line from bottlenecks.csv produced by CAVER to fill in data about Tunnel bottleneck
:param bottleneck_data: line of data from bottlenecks.csv corresponding to bottleneck residues of this tunnel
:param transform_mat: transformation matrix to be applied on the input coordinates

**Method** `fill_data`

```
def fill_data(
    self,
    data_section: List[str]
)
```

Processes seven lines of data from tunnel_profiles.csv produced by CAVER to create Tunnel object
:param data_section: seven lines of data from tunnel_profiles.csv corresponding to a single tunnel

**Method** `get_bottleneck_line`

```
def get_bottleneck_line(
    self,
    md_label: str
) -> str
```

Produces this tunnel's data in CVS format analogous to CAVER bottlenecks.csv files, adding info on its md_label
:param md_label: name of folder with the source MD simulation data
:return: line in CVS format

**Method** `get_center_line`

```
def get_center_line(
    self
) -> <built-in function array>
```

Returns coordinates of spheres centers forming the tunnel

**Method** `get_closest_sphere2coords`

```
def get_closest_sphere2coords(
    self,
    xyz: np.array
) -> (float, np.array) or (None, None)
```

Identifies closest sphere from this tunnel to given coordinates, using spherical grid for efficiency.
:param xyz: coordinates of point to which the distance is computed
:return: distance to the closest sphere and the closest sphere data

**Method** `get_csv_lines`

```
def get_csv_lines(
    self,
    md_label: str
) -> str
```

Produces this tunnel's data in CVS format analogous to caver tunnel_profile files, adding info on its md_label
:param md_label: name of folder with the source MD simulation data
:return: seven lines in CVS format

**Method** `get_parameters`

```
def get_parameters(
    self
) -> (float, float, float, float)
```

Get basic parameters of this tunnel
:return: length, bottleneck_radius, curvature, and throughput of this tunnel

**Method** `get_pdb_file_format`

```
def get_pdb_file_format(
    self
) -> List[str]
```

Saves tunnel points as PDB file according to CAVER format
:return

**Method** `get_points_data`

```
def get_points_data(
    self
) -> <built-in function array>
```

Convert data of points forming this tunnel, adding info on order of points and identifying the tunnel end point
:return: augmented data for this tunnel suitable for LayeredRepresentation.load_points() method

**Method** `get_snapshot_id`

```
def get_snapshot_id(
    self,
    id_position: int = 1,
    delimiter: str = '.'
) -> int
```

Returns numerical position of snapshot to which this tunnel belongs based on snapshot name from CAVER
:param id_position: which field will contain snapshot ID after splitting with the delimiter
:param delimiter: delimiter to find the snapshot ID at the specified position
:return: snapshot ID

**Method** `get_visualization_cgo`

```
def get_visualization_cgo(
    self
) -> List[float]
```

Converts tunnel points to Pymol compiled graphics object(CGO) for visualization
:return: CGO of tunnel points

**Method** `has_better_throughput`

```
def has_better_throughput(
    self,
    other_tunnel: Tunnel
) -> bool
```

Check if this tunnel has better throughput than the other one
:param other_tunnel: other tunnel to compare with

###### Method `is_same`

```
def is_same(
    self,
    other: Tunnel
) -> bool
```

Test if the tunnel is same as the other tunnel
:param other: other tunnel to compare with
:return: are two tunnels same?

##### Class `TunnelCluster`

```
class TunnelCluster(
    cluster_id: int,
    parameters: dict,
    transform_mat: np.array,
    starting_point_coords: np.array
)
```

Class for processing of tunnel clusters from CAVER, containing tunnels
:param cluster_id: ID of the cluster from CAVER
:param parameters: job configuration parameters
:param transform_mat: transformation matrix to be applied on the nodes coordinates

###### Methods

###### Method `add_tunnel`

```
def add_tunnel(
    self,
    tunnel: Tunnel
)
```

Adds a tunnel to this cluster
:param tunnel: Tunnel object to add

###### Method `count_tunnels`

```
def count_tunnels(
    self
) -> int
```

Counts number of tunnels in this cluster
:return: number of tunnels

###### Method `count_valid_tunnels`

```
def count_valid_tunnels(
    self
) -> int
```

Counts number of tunnels that passed the initial filters for creation of cluster representative
:return: number of valid tunnels

**Method** `create_layered_cluster`

```
def create_layered_cluster(
    self
) -> Tuple[int, str, transport_tools.libs.geometry.LayeredPathSet]
```

Combines LayeredRepresentation of all valid tunnels within the cluster, executes the actual layering, and identifies the representative set of Layered paths
:return: cluster ID, md_label and a set of layered paths representing this cluster

**Method** `get_characteristics`

```
def get_characteristics(
    self
)
```

Get information on the average throughput and the number of tunnels in this cluster
:return: the average throughput and the number of tunnels

**Method** `get_closest_tunnel_sphere_in_frame2coords`

```
def get_closest_tunnel_sphere_in_frame2coords(
    self,
    xyz: np.array,
    snap_id: int
) -> (float, np.array) or (None, None)
```

Identifies closest sphere from a tunnel in the investigated snapshot to given coordinates
:param xyz: coordinates of point to which the distance is computed
:param snap_id: ID of investigated snapshot
:return: distance to the closest sphere and the closest sphere data, or None if no tunnel exists in the snapshot

**Method** `get_property`

```
def get_property(
    self,
    property_name: str,
    active_filters: dict
) -> <built-in function array>
```

Returns array containing all values of given property for all tunnels in cluster that fulfill active filters
:param property_name: name of property to extract
:param active_filters: filters to be applied (created by define_filters() function)
:return: array of tunnel property values adhering to filters

**Method** `get_subcluster`

```
def get_subcluster(
    self,
    snap_ids: Optional[List[int]] = None,
    active_filters: Optional[dict] = None
) -> transport_tools.libs.networks.TunnelCluster
```

Returns subcluster with tunnels from the frames and/or all tunnels in cluster fulfilling the active filters
:param snap_ids: list of snapshot IDs to create the subcluster from
:param active_filters: filters to be applied (created by define_filters() function)
:return: subcluster with tunnels from given frames and/or fulfilling filters

**Method is_same**

```
def is_same(
    self,
    other: TunnelCluster
) -> bool
```

Test if the cluser is same as the other cluster
:param other: other cluster to compare with
:return: are two clusters same?

**Method remove_tunnel**

```
def remove_tunnel(
    self,
    snap_id: int
)
```

Remove tunnel with a given snapshot ID from this cluster
:param snap_id: snapshot ID

**Method save_cgo**

```
def save_cgo(
    self,
    out_path: str
)
```

Dump all tunnel as single Pymol compiled graphics object(CGO) to gzipped files for visualization of this cluster
:param out_path: folder where the CGO should be dumped to

**Method save_pdb_files**

```
def save_pdb_files(
    self,
    snap_ids: List[int],
    out_file: str
) -> bool
```

Writes tunnels with specified snapshot IDs to MULTIMODEL PDB file
:param snap_ids: list of snapshot IDs to save
:param out_file: file to save to
:return: if saved successfully

**Class TunnelNetwork**

```
class TunnelNetwork(
    parameters: dict,
    md_label: str
)
```

Class for processing of CAVER results for tunnels, contains TunnelCluster objects
:param parameters: job configuration parameters
:param md_label: name of folder with the source MD simulation data

**Ancestors (in MRO)**

- transport_tools.libs.networks.Network

###### Methods

###### Method `cluster_exists`

```
def cluster_exists(
    self,
    query_id: int
) -> bool
```

Verifies if cluster with given id exists
:param query_id: ID of evaluated cluster

###### Method `get_cluster`

```
def get_cluster(
    self,
    cluster_id: int
) -> transport_tools.libs.networks.TunnelCluster
```

Retrieve cluster with given ID
:param cluster_id: ID of retrieved cluster
:return: the cluster with given ID

###### Method `get_clusters4layering`

```
def get_clusters4layering(
    self
) -> List[transport_tools.libs.networks.TunnelCluster]
```

Get cluster in this tunnel network for layering
:return: list of clusters for processing

###### Method `read_tunnels_data`

```
def read_tunnels_data(
    self
)
```

Process tunnel_profile_file from CAVER to create TunnelClusters and their Tunnels

##### Class `TunnelProfile4MD`

```
class TunnelProfile4MD(
    md_label: str,
    caver_clusters: List[int],
    dump_file: str,
    parameters: dict
)
```

Class to store and process the tunnel profile of tunnels belonging to caver clusters of given MD simulation; there can be multiple such profiles per one cumulative tunnel profile of parent supercluster (SC)
:param md_label: name of folder with the source MD simulation data
:param caver_clusters: list of valid clusters from this MD simulation belonging to parent SC
:param dump_file: dump file of original TunnelCluster from TunnelNetwork of this MD simulation
:param parameters: job configuration parameters

###### Methods

**Method** `count_tunnels`

```
def count_tunnels(
    self
)
```

Count number of tunnels in this tunnel profile

**Method** `enumerate_caver_cluster_ids`

```
def enumerate_caver_cluster_ids(
    self
) -> List[int]
```

Report tunnel cluster IDs that are present (valid) in tunnel profile from this MD simulation
:return:

**Method** `filter_tunnels`

```
def filter_tunnels(
    self,
    active_filters: dict
)
```

Filtering of tunnels in this tunnel profiles; tunnels that failed to pass are removed for this profile
:param active_filters: filters to be applied (created by define_filters() function)

**Method** `get_bottleneck_residues_frequency`

```
def get_bottleneck_residues_frequency(
    self
) -> Dict[str, float]
```

Collect bottleneck residues frequency for all tunnels present in tunnel profile from this MD simulation
:return: frequency of bottleneck residues occurrence

**Method** `get_parameters`

```
def get_parameters(
    self
) -> (List[float], List[float], List[float], List[float])
```

Collect tunnel parameters (length, radius, curvature, throughput) for all tunnels present in tunnel profile from this MD simulation
:return: lists of tunnel parameters

**Method** `get_property_time_evolution_data`

```
def get_property_time_evolution_data(
    self,
    property_name: str,
    missing_value_default: float = 0
) -> <built-in function array>
```

Returns array containing values of given tunnel property for each simulation frame
:param property_name: name of property to extract
:param missing_value_default: value to be used for frames where tunnels are missing or invalid in this cluster
:return: array of tunnel property values adhering to filters

**Method** `load_network`

```
def load_network(
    self
)
```

Extract all clusters from this MD simulation belonging to parent SC and create tunnel records single (or none) tunnel per snapshot to represent parent SC in given snapshot, here also distances of tunnel spheres are adjusted to general starting point at the origin [0,0,0]

**Method** `write_csv_section`

```
def write_csv_section(
    self,
    file_handler: TextIO
)
```

Writes tunnel data for tunnels from this profile to CSV file
:param file_handler: file object for opened CSV file to write tunnel profile data to

**Method** `write_residues`

```
def write_residues(
    self,
    file_handler: TextIO
)
```

Writes bottleneck residues data for tunnels from this profile to CSV file
:param file_handler: file object for opened CSV file to write bottleneck residues data to

### Module `transport_tools.libs.protein_files`

#### Functions

**Function** `TrajectoryFactory`

```
def TrajectoryFactory(
    parameters: dict,
    md_label: str,
    superpose_mask: str = None
)
```

**Function** `get_general_rot_mat_from_2_ca_atoms`

```
def get_general_rot_mat_from_2_ca_atoms(
    in_pdb_file: str
) -> <built-in function array>
```

Arbitrary selects Calpha atoms from 1/4 and 3/4 of sequence and orients the 1st one along z-axis, and 2nd one into yz-plane
:param in_pdb_file: input PDB file
:return: 4x4 transformation matrix describing the re-orientation

**Function** `get_transform_matrix`

```
def get_transform_matrix(
    moved_protein: str,
```

```
    target_protein: str,
    md_label: str = '',
    max_iter: int = 5,
    rmsd_cutoff: float = 0.1
) -> (<built-in function array>, <class 'str'>)
```

Performs a sequence alignment of the target and moved proteins, then tries to reduce the RMSD by a series of iterations removing the atoms with the higher difference before and after alignment.
:param moved_protein: input PDB file, to be moved
:param target_protein: reference PDB file to which we align
:param md_label: name of folder with the source MD simulation data
:param max_iter: maximum number of iterations
:param rmsd_cutoff: rmsd convergence cutoff to stop the iterations
:return: 4x4 transformation matrix describing the alignment, similar to default Pymol align command & md_label

**Function** `transform_aquaduct`

```
def transform_aquaduct(
    md_label: str,
    tar_file: str,
    aquaduct_results_pdb_filename: str,
    reference_pdb_file: str
) -> (<built-in function array>, <class 'str'>, <class 'int'>)
```

Prepares temporary files from AQUA-DUCT data and gets transformation matrix
:param md_label: name of folder with the source MD simulation data
:param tar_file: tarfile with AQUA-DUCT results
:param aquaduct_results_pdb_filename: name of PDB file with protein structure in the AQUA-DUCT tarfile
:param reference_pdb_file: reference PDB file to which we align
:return: transformation matrix, md_label, number_of_raw_paths in tar_file

**Function** `transform_pdb_file`

```
def transform_pdb_file(
    in_pdb_file: str,
    out_pdb_file: str,
    transform_mat: <built-in function array>
)
```

Translates and rotates atoms in PDB file by transformation matrix
:param in_pdb_file: input pdb file path
:param out_pdb_file: path to the transformed pdb file
:param transform_mat: 4x4 transformation matrix to be applied on the input coordinates

#### Classes

**Class** `AtomFromPDB`

```
class AtomFromPDB(
    line: str
)
```

Generating Atoms from pdb lines
:param line: line from PDB file starting with “ATOM” field

**Methods**

###### Method `isprotein`

```
def isprotein(
    self
)
```

Test if the atom is from protein residues

##### Class `TrajectoryMdtraj`

```
class TrajectoryMdtraj(
    parameters: dict,
    md_label: str,
    superpose_mask: str = 'name CA'
)
```

Class for handling MD trajectories with MDtraj package
:param parameters: job configuration parameters
:param md_label: name of folder with the source MD simulation data
:param superpose_mask: MDtraj mask for selection to get same system used in CAVER, if None, all atoms are used

###### Ancestors (in MRO)

- transport_tools.libs.protein_files.TrajectoryTT

###### Methods

###### Method `get_coords`

```
def get_coords(
    self,
    start_frame: int,
    end_frame: int,
    keep_mask: Optional[str] = None,
    out_file: Optional[str] = None
) -> <built-in function array>
```

Get coordinates of system across specified frames, potentially after removing some of its parts
:param start_frame: start frame to consider
:param end_frame: end frame to consider
:param keep_mask: MDtraj mask to select part of system to keep
:param out_file: file where to save specified frames as MULTIMODEL PDB file
:return: coordinates of the kept system in selected frames

##### Class `TrajectoryPytraj`

```
class TrajectoryPytraj(
    parameters: dict,
    md_label: str,
    superpose_mask: str = '@CA'
)
```

Class for handling MD trajectories with Pytraj package
:param parameters: job configuration parameters
:param md_label: name of folder with the source MD simulation data
:param superpose_mask: Pytraj mask for selection to get same system used in CAVER, if None, all atoms are used

###### Ancestors (in MRO)

- transport_tools.libs.protein_files.TrajectoryTT

###### Methods

###### Method `get_coords`

```
def get_coords(
    self,
    start_frame: int,
    end_frame: int,
    keep_mask: Optional[str] = None,
    out_file: Optional[str] = None
) -> <built-in function array>
```

Get coordinates of system across specified frames, potentially after removing some of its parts
:param start_frame: start frame to consider
:param end_frame: end frame to consider
:param keep_mask: inverted AMBER mask to select part of system to keep
:param out_file: file where to save specified frames as MULTIMODEL PDB file
:return: coordinates of the kept system in selected frames

##### Class `TrajectoryTT`

```
class TrajectoryTT(
    parameters: dict,
    md_label: str,
    superpose_mask: str
)
```

Generic class for handling MD trajectories
:param parameters: job configuration parameters
:param md_label: name of folder with the source MD simulation data
:param superpose_mask: mask for selection to get same system used in CAVER, if None, all atoms are used

###### Descendants

- transport_tools.libs.protein_files.TrajectoryMdtraj
- transport_tools.libs.protein_files.TrajectoryPytraj

###### Methods

###### Method `get_coords`

```
def get_coords(
    self,
    start_frame: int,
    end_frame: int,
    keep_mask: Optional[str] = None,
    out_file: Optional[str] = None
) -> <built-in function array>
```

Get coordinates of system across specified frames, potentially after removing some of its parts. And if out_file is provided, the resulting structure is saved too
:param start_frame: start frame to consider
:param end_frame: end frame to consider
:param keep_mask: mask to select part of system to keep

:param out_file: file where to save specified frames as MULTIMODEL PDB file
:return: coordinates of the kept system in selected frames

###### Method `inputs_exists`

```
def inputs_exists(
    self
) -> bool
```

Tests if inputs exists

###### Method `write_frames`

```
def write_frames(
    self,
    start_frame: int,
    end_frame: int,
    out_file: str,
    keep_mask: Optional[str] = None
)
```

Write specified frames to MULTIMODEL PDB file
:param start_frame: start frame to consider
:param end_frame: end frame to consider
:param out_file: file where to save specified frames
:param keep_mask: mask to select part of system to keep

##### Class `VizAtom`

```
class VizAtom(
    array: list,
    use_hetatm: bool = True
)
```

Creation of PDB line from variables
:param array: list of variables to create PDB line
:param use_hetatm: if the PDB line should use HETATM

### Module `transport_tools.libs.tools`

#### Functions

##### Function `load_checkpoint`

```
def load_checkpoint(
    filename: str = 'checkpoints/transport_processes.dump',
    update_config: Optional[transport_tools.libs.config.AnalysisConfig] = None
) -> transport_tools.libs.tools.TransportProcesses
```

Loads previously saved TransportProcess object, possibly updating new parameters from provided update_config object
:param filename: input file with saved TransportProcess object
:param update_config: configuration object with job parameters
:return: loaded TransportProcess object

**Function** `save_checkpoint`

```
def save_checkpoint(
    object_to_save: transport_tools.libs.tools.TransportProcesses,
    filename: str = 'checkpoints/transport_processes.dump',
    overwrite: bool = False
)
```

Save TransportProcess object for later used
:param object_to_save: TransportProcess object to save
:param filename: filename where to save the object
:param overwrite: if replace the content of the file in case the file with a given filename exists

**Function** `visualize_transport_details`

```
def visualize_transport_details(
    out_folder_path: str,
    trajectory: transport_tools.libs.protein_files.TrajectoryTT,
    start_frame: int,
    end_frame: int,
    caver_traj_offset: int,
    caver_clusters: Optional[List[transport_tools.libs.networks.TunnelCluster]] = None,
    start_snapshot: Optional[int] = None,
    end_snapshot: Optional[int] = None,
    resids: Optional[List[int]] = None
)
```

Visualize dynamics of tunnels and/or events in given set of frames together with biomolecule
:param out_folder_path: output folder path
:param trajectory: MD simulation trajectory to process
:param start_frame: start frame for visualization
:param end_frame: end frame for visualization
:param caver_traj_offset: difference in IDs of MD frames (from 0) and caver snapshots (often from 1)
:param caver_clusters: list of tunnel clusters for visualization
:param start_snapshot: start snapshot for visualization
:param end_snapshot: end snapshot for visualization
:param resids: residue ID(s) to show as events

#### Classes

**Class** `EventAssigner`

```
class EventAssigner(
    parameters: dict,
    event_specification: Tuple[str, str, Tuple[str, Tuple[int, int]]],
    event: transport_tools.libs.geometry.LayeredPathSet,
    superclusters: Dict[int, transport_tools.libs.networks.SuperCluster],
    active_filters: dict
)
```

Class to perform assignment of single transport event to matching superclusters
:param parameters: job configuration parameters
:param event_specification: folder name of source MD simulation, event label and tuple containing resname&resid of ligand responsible for this path, supplemented with info on beginning and last frames for the transport event
:param event: pathset containing representative path for evaluated event
:param superclusters: dictionary with SuperCluster objects to assign to

###### Methods

###### Method `perform_assignment`

```
def perform_assignment(
    self
) -> (typing.Tuple[str, str, typing.Tuple[str, typing.Tuple[int, int]]], <built-in function arra
```

Identifies most likely supercluster (SC) through which a single evaluated transport event occurred
:return: event specification, array with IDs of SC to which the event is assigned, maximal buriedness and penetration depth of these SCs

##### Class `OutlierTransportEvents`

```
class OutlierTransportEvents(
    parameters: dict
)
```

Class for storing information on transport events that cannot be assigned to any supercluster, & their reporting
:param parameters: job configuration parameters

###### Methods

###### Method `add_transport_event`

```
def add_transport_event(
    self,
    md_label: str,
    path_id: str,
    event_type: str,
    traced_event: Tuple[str, Tuple[int, int]],
    globally_unassigned: bool = True
)
```

Store information on unassigned transport event
:param md_label: name of folder with the MD simulation data that contain this transport event
:param path_id: AQUA-DUCT path ID of this event
:param event_type: type(release or entry) of this event
:param traced_event: tuple containing identity of ligand responsible for this event,
and beginning and last frames of the event
:param globally_unassigned: if the event is outlier for all simulations or just md_label

###### Method `count_events`

```
def count_events(
    self,
    md_label: str = 'overall'
) -> (<class 'int'>, <class 'int'>, <class 'int'>)
```

Counts number of all unassigned events (total, entries, releases)
:param md_label: name of folder with the MD simulation data that contain this transport event

###### Method `exist`

```
def exist(
    self,
    md_label: str = 'overall'
) -> bool
```

Tests if some events were unassigned hence considered as outlying
:param md_label: name of folder with the MD simulation data that contain this transport event

**Method** `prepare_visualization`

```
def prepare_visualization(
    self,
    md_label: str = 'overall'
) -> List[str]
```

Generates lines for Pymol visualization script
:param md_label: visualization of which simulations to prepare; by default 'overall' visualization
:return: lines to load visualization of this SC into Pymol

**Method** `report_events_details`

```
def report_events_details(
    self,
    filename: str
)
```

Save file with detailed information about unassigned transport events
:param filename: output filename

**Method** `report_summary_line`

```
def report_summary_line(
    self,
    widths: List[int],
    md_label: str = 'overall'
) -> str
```

Prepares information on transport events flagged as outliers for generation of summary of superclusters
:param widths: column widths to match the format of the rest of the summary file
:param md_label: summary of which simulations to report; by default report 'overall' stats
:return: info on outlier events

**Class** `TransportProcesses`

```
class TransportProcesses(
    config: transport_tools.libs.config.AnalysisConfig
)
```

Class for analysis of transport processes and tunnels, contains 'global' work-flows using methods from SuperClusters (SC)
:param config: configuration object with job parameters

**Methods**

**Method** `assign_transport_events`

```
def assign_transport_events(
    self,
    md_labels: Optional[List[str]] = None
)
```

Finds superclusters (SCs) through evaluated transport event happened, and assigns remaining events as outliers; this changes self._events_assigned = True, enabling consideration of transport events during filtering
:param md_labels: list to restrict assignment to particular Networks (simulations) only

**Method** `clear_results`

```
def clear_results(
    self,
    overwrite: bool = False
)
```

Removes output folder
:param overwrite: if to perform the cleaning of non empty folder

**Method** `compute_transformations`

```
def compute_transformations(
    self
)
```

Compute transformation matrices that are used to align all data

**Method** `compute_tunnel_clusters_distances`

```
def compute_tunnel_clusters_distances(
    self
)
```

Compute pairwise cluster-cluster distances, and save their matrix

**Method** `create_layered_description4aquaduct_networks`

```
def create_layered_description4aquaduct_networks(
    self
)
```

Creates layered representation of AquaDuct networks for all MD simulations

**Method** `create_layered_description4tunnel_networks`

```
def create_layered_description4tunnel_networks(
    self
)
```

Creates layered representation of tunnel networks for all MD simulations

**Method** `create_super_cluster_profiles`

```
def create_super_cluster_profiles(
    self
)
```

Parallel merging of caver tunnel profiles into new ones for superclusters (SCs), and saving SC details

**Method** `enumerate_valid_super_clusters`

```
def enumerate_valid_super_clusters(
    self,
    consider_transport_events: bool = False
) -> int
```

Counts how many superclusters (SCs) are valid
:param consider_transport_events: if related filters related to transport events should be considered
:return: number of valid SCs

**Method `filter_super_cluster_profiles`**

```
def filter_super_cluster_profiles(
    self,
    min_length: float = -1,
    max_length: float = -1,
    min_bottleneck_radius: float = -1,
    max_bottleneck_radius: float = -1,
    min_curvature: float = -1,
    max_curvature: float = -1,
    min_sims_num: int = 1,
    min_snapshots_num: int = 1,
    min_avg_snapshots_num: float = -1,
    min_total_events: int = -1,
    min_entry_events: int = -1,
    min_release_events: int = -1
)
```

Performs parallel filtering of superclusters (SC) profiles using defined filters, and saves SC details
NOTE: -1 => filter is not active
:param min_length: minimum tunnel length
:param max_length: maximum tunnel length
:param min_bottleneck_radius: minimum tunnel bottleneck radius
:param max_bottleneck_radius: maximum tunnel bottleneck radius
:param min_curvature: minimum tunnel curvature
:param max_curvature: maximum tunnel curvature
:param min_sims_num: present in minimum number of MD simulations
:param min_snapshots_num: present in minimum number of snapshots
:param min_avg_snapshots_num: present in minimum number of snapshots on average
:param min_total_events: having minimum transport events
:param min_entry_events: having minimum entry events
:param min_release_events: having minimum release events

**Method `generate_super_cluster_summary`**

```
def generate_super_cluster_summary(
    self,
    out_filename: str = 'super_cluster_statistics.csv'
)
```

Generates summary file with superclusters statistics.
:param out_filename: name of file to save the summary to

**Method `get_property_time_evolution_data`**

```
def get_property_time_evolution_data(
    self,
    property_name: str,
    active_filters: dict,
    sc_id: Optional[int] = None,
    missing_value_default: float = 0
) -> Dict[int, Dict[str, <built-in function array>]]
```

For each MD simulation in specified supercluster, return array containing values of given tunnel property
for each simulation frame
:param property_name: name of property to extract
:param active_filters: filters to be applied (created by define_filters() function)
:param sc_id: supercluster ID after prioritization if we want to focus on particular tunnel only
:param missing_value_default: value to be used for frames where tunnels are missing or invalid in given cluster
:return: for each supercluster ID we have mapping of tunnel property values per MD simulation

**Method** `merge_tunnel_clusters2super_clusters`

```
def merge_tunnel_clusters2super_clusters(
    self
)
```

Performs clustering of tunnel clusters and creates of their superclusters (SCs)

**Method** `process_aquaduct_networks`

```
def process_aquaduct_networks(
    self
)
```

Process AQUA-DUCT networks for all MD simulations

**Method** `process_tunnel_networks`

```
def process_tunnel_networks(
    self
)
```

Process tunnel networks for all MD simulations

**Method** `save_super_clusters_visualization`

```
def save_super_clusters_visualization(
    self,
    script_name: str = 'view_super_clusters.py'
)
```

Save visualization of superclusters (SCs) for Pymol
:param script_name: filename of the generated visualization script

**Method** `show_tunnels_passing_filter`

```
def show_tunnels_passing_filter(
    self,
    sc_id: int,
    active_filters: dict,
    out_folder_path: str,
    md_labels: Optional[List[str]] = None,
    start_snapshot: Optional[int] = None,
    end_snapshot: Optional[int] = None,
    trajectory: bool = False
)
```

Visualize tunnels from particular supercluster that fulfill active_filters, possibly showing only particular snapshots, selected MD simulations and providing pdb file with corresponding protein ensemble
:param sc_id: ID of source supercluster to visualize its tunnels
:param active_filters: filters to be applied (created by define_filters() function)
:param out_folder_path: path to folder into which we save the visualization
:param md_labels: list to restrict visualization to tunnel networks from particular MD simulations only
:param start_snapshot: start snapshot for tunnel visualization
:param end_snapshot: end snapshot for tunnel visualization
:param trajectory: if we should visualize tunnels per MD simulation trajectory with protein ensembles

**Method** `update_configuration`

```
def update_configuration(
    self,
    new_config: transport_tools.libs.config.AnalysisConfig
)
```

Updates parameters based on new job parameters
:param new_config: object with job parameters

### Module `transport_tools.libs.ui`

#### Functions

**Function** `bye_bye`

```
def bye_bye(
    process_start
)
```

**Function** `greetings`

```
def greetings()
```

**Function** `init_logging`

```
def init_logging(
    verbose_logging: bool = False,
    logfile: str = 'transport_tools.log'
)
```

Initiates and sets logging, also defines logging filtering
:param verbose_logging: if more details should be provided on debug level
:param logfile: file to log into

**Function** `init_parser`

```
def init_parser() -> argparse.ArgumentParser
```

Initiates command line parser
:return: the parser

**Function** `initiate_tools`

```
def initiate_tools(
    level: str = 'info',
    verbose_logging: bool = False,
    logfile: str = 'transport_tools.log'
)
```

Starts logging, enables initial and terminal messages
:param level: logging level to be used
:param verbose_logging: if more details should be provided on debug level
:param logfile: file to log into

**Function** `license_printer`

```
def license_printer()
```

Prints info about the license

**Function** `process_count`

```
def process_count(
    num_processes: int
) -> str
```

**Function** `progressbar`

```
def progressbar(
    iteration: int,
    total: int
)
```

Generates progress bar for processes
:param iteration: current iteration of the process
:param total: total number of iterations

**Function** `set_logging_level`

```
def set_logging_level(
    level: str
)
```

Sets currently used level of logging
:param level: logging level to be used

#### Classes

**Class** `SuppressMsg`

```
class SuppressMsg
```

A context manager to suppress all messages even those originating from external programs but not suppressing exceptions.

**Class** `TimeProcess`

```
class TimeProcess(
    prefix_msg: str = ''
)
```

Monitors and reports duration of the process(es)
:param prefix_msg: text preceding the report on the process duration

### Module `transport_tools.libs.utils`

#### Functions

##### Function `convert_coords2cgo`

```
def convert_coords2cgo(
    coords: <built-in function array>,
    color_id: int
) -> List[float]
```

Converts xyz-coordinates of sequence of points such as trace and tunnel to Pymol compiled graphics object(CGO)
:param coords: xyz-coordinates of points
:param color_id: Pymol color id
:return: CGO of the points

##### Function `convert_spheres2cgo_surface`

```
def convert_spheres2cgo_surface(
    spheres: List[Tuple[<built-in function array>, float]],
    color_id: int,
    resolution: float = 0.5
) -> List[float]
```

Converts spheres (tuples that contain XYZ coords and radius) to surface complied graphics object for Pymol.
:param spheres: list of spheres to generate their approximate surface
:param color_id: Pymol color id
:param resolution: surface grid resolution in Angstroms
:return: CGO of spheres surface

##### Function `get_caver_color`

```
def get_caver_color(
    color_id: int
) -> List[float]
```

Converts Pymol color IDs to RGB format, keeping within the set of ‘reasonable’ colors from CAVER rgb.py
:param color_id: Pymol color ID
:return: RBG colors in Pymol format Pymol

##### Function `get_filepath`

```
def get_filepath(
    root_folder: str = '.',
    pattern: str = '*'
) -> str
```

Get full and unique path to file in root folder specified by file pattern (may include subfolders)
:param root_folder: folder to search in
:param pattern: search patter
:return: file path

##### Function `node_labels_split`

```
def node_labels_split(
    node_label: str
) -> (<class 'int'>, <class 'int'>)
```

Returns layer ID and cluster ID, useful for sorting of nodes labels
:param node_label: node label formatted as {layer}_{cluster_id} or SP for starting point
:return: sortable tuple for ordering by layer and then by cluster id

###### Function `path_loader_string`

```
def path_loader_string(
    path: str
) -> str
```

Converts a string with a path to a string with an os.path.join loader for a cross-platform compatibility.
:param path: path to convert
:return: os path loader

###### Function `reweighting_file_parser`

```
def reweighting_file_parser(
    engine: str = 'gamd'
)
```

###### Function `splitall`

```
def splitall(
    path2split: str
) -> List[str]
```

Get all sub-parts of a file or directory path. Credited to Trent Mick.
:param path2split: path to process
:return: list of subpart of the path

###### Function `test_file`

```
def test_file(
    filename: str
)
```

Tests if file exists
:param filename: path to the file to test

### Module `transport_tools.scripts`

#### Sub-modules

- transport_tools.scripts.tt_engine

### Module `transport_tools.scripts.tt_engine`

#### Functions

###### Function `test_checkpoint`

```
def test_checkpoint(
    checkpoint_file: str
```

)

Test that we can save checkpoint before running expensive calculations

---

Generated by pdoc 0.10.0 (https://pdoc3.github.io).
