## Supplementary File 5 for "TransportTools: a library for high-throughput analyses of internal voids in biomolecules and ligand transport through them"

---

#### Supplementary File 5 – TransportTools performance

### 1. Description of molecular dynamics simulation datasets

The three enzymes which were selected for benchmarking the performance of TransportTools belong to the same subfamily of hydrolases: (i) *Rhodococcus rhodochrous* haloalkane dehalogenase DhaA composed of 296 residues (Stepankova *et al.*, 2013), (ii) potato *Solanum tuberosum* epoxide hydrolase 1 StEH1 composed of 328 residues (Mowbray *et al.*, 2006), and (iii) *Candida rugosa* lipase CRL composed of 534 residues (Grochulski *et al.*, 1993, 1994). The accessibility of the active sites to surrounding solvent differs notably among these three enzymes. In case of DhaA, the deeply buried site is connected to surface by rather narrow transient tunnels enabling only limited exchange, while the StEH1 features several well-defined tunnels enabling massive transport of water moleculs. Finally, the accessibility of the CRL active site is controlled by a lid-like gate, making it considerably exposed to the surrounding environment in its open form.

The following high-quality structures were retrieved from the Protein Data Bank: 4E46 for DhaA, 2CJP for StEH1, as well as 1CRL and 1TRH for open and closed conformations of CRL, respectively. These systems were prepared in their biologically active pH and solvated using 4-point OPC water model (Izadi *et al.*, 2014) and then neutralized with counterions ($Na^+$ and $Cl^-$) to reach 0.1 M concentration. Using PMEMD and PMEMD.CUDA modules (Salomon-Ferrer *et al.*, 2013) of AMBER18 (D.A. Case *et al.*) with ff14SB force field (Maier *et al.*, 2015), the systems were first energy minimized in several rounds with decreasing harmonic restrains. The minimization procedure was followed by equilibration MD simulations with gradual heating to 310 K under a constant volume using the Langevin thermostat (Zwanzig, 1973), periodic boundary conditions with the particle mesh Ewald method (Darden *et al.*, 1993), and 4 fs time-step enabled by SHAKE and hydrogen mass repartitioning algorithms (Hopkins *et al.*, 2015; Ryckaert *et al.*, 1977). Finally, these simulations were continued with an unrestrained 200 ns simulation at constant pressure and temperature using the weak-coupling barostat and thermostat (Berendsen *et al.*, 1984).

The last snapshot served to initiate an adaptive sampling approach using High-Throughput Molecular Dynamics package (Doerr *et al.*, 2016). During the sampling, the dihedral angles of residues around the active site were employed as the guiding metric. The adaptive sampling scheme performed for each protein system was the following: ten epochs, each featuring five parallel simulations of 100 ns, reaching the overall sampling of 5 µs per protein. Next, the generated MD trajectories were analyzed by CAVER 3.0.1 (Pavelka *et al.*, 2016; Chovancova *et al.*, 2012) to study the tunnel dynamics. Furthermore, AQUA-DUCT 1.0 software (Magdziarz *et al.*, 2020) was used to track the movement of water molecules. Overall, data on tunnel ensembles and water movement from 150 trajectories capturing 15 µs of aggregate simulation data were used for benchmarking of TransportTools.

### 2. Overview of transport-related properties of the analyzed datasets

During the analyses across 50 simulations for each protein system, we identified 985,219 distinct tunnels in the simulations of haloalkane dehalogenase, 5,008,440 distinct tunnels in the simulations of epoxide hydrolase, and 4,207,210 distinct tunnels in the simulations of lipase (Fig. 1). Regarding the number of events describing the exchange of water molecules between active sites and bulk solvent, we could detect 2,650 (haloalkane dehalogenase), 49,100 (epoxide hydrolase), and 47,916 (lipase) of such events in the simulations. The majority of these events could be assigned to one of the transport pathways (Fig. 2). Identified tunnels and events were then integrated to form 71, 164, and 209 superclusters for haloalkane dehalogenase, epoxide hydrolase, and lipase, respectively, out of which about 1/3 were involved in the transport of at least one water molecule (Fig. 3).

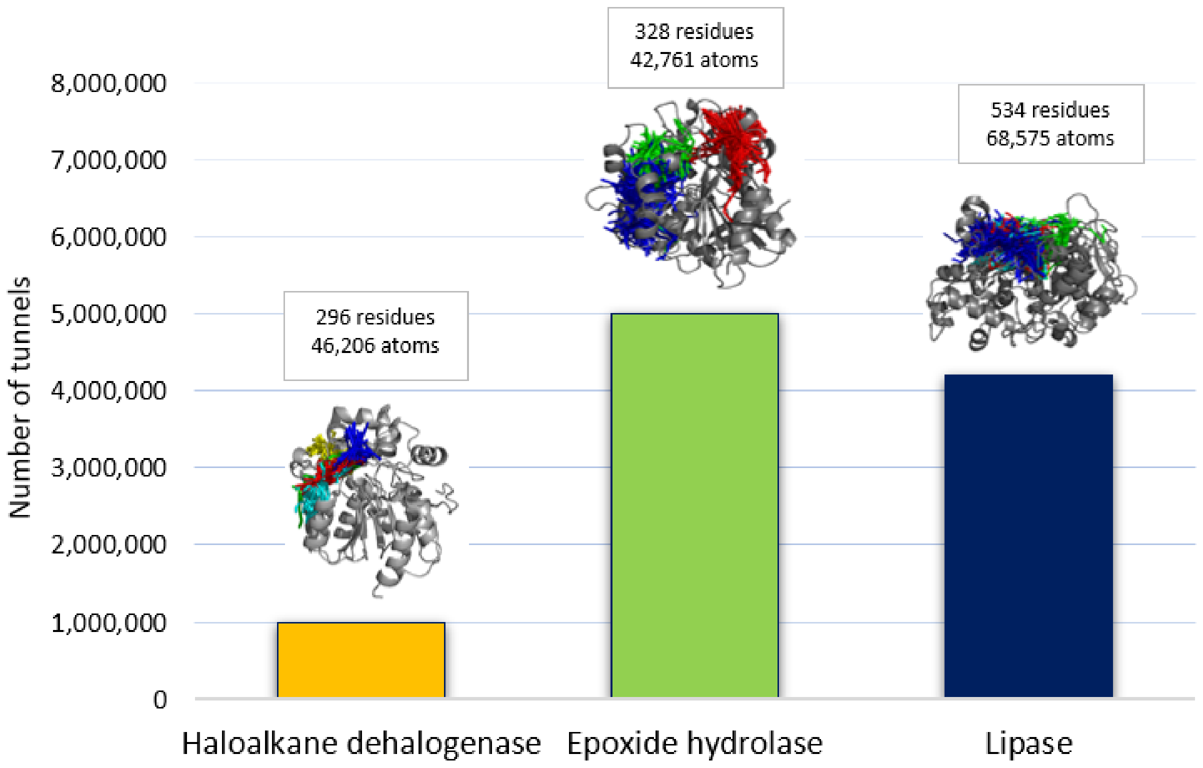


**Fig. 1.** Tunnels detected by CAVER across 50 simulations for each protein system. The representative protein is shown as cartoon and major tunnel clusters.

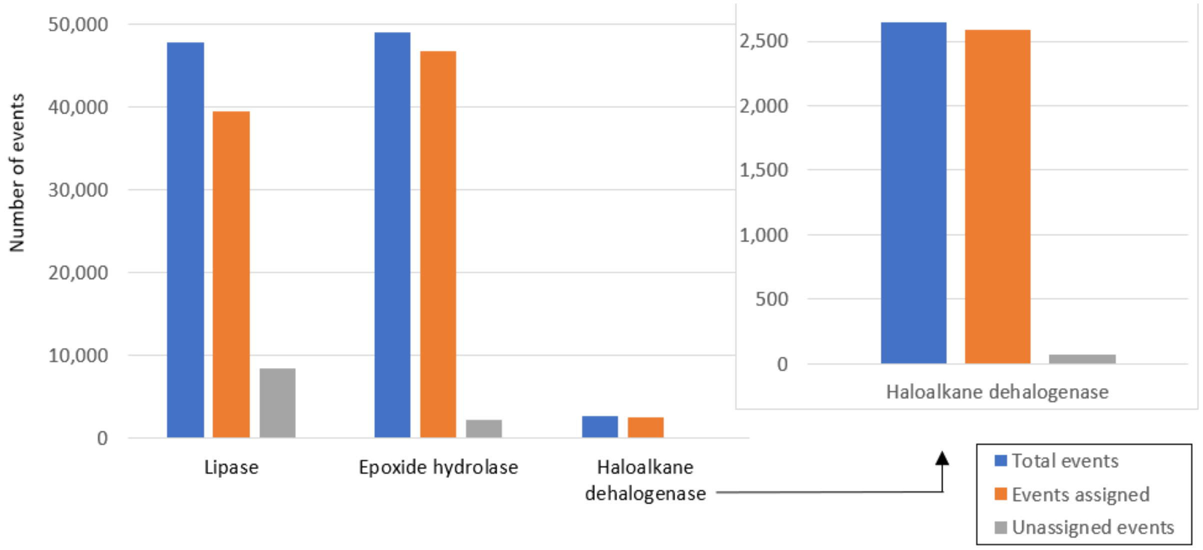


**Fig. 2.** Transport events of water molecules detected by AQUA-DUCT and assigned to transport pathways by TransportTools across 50 simulations for each protein system.

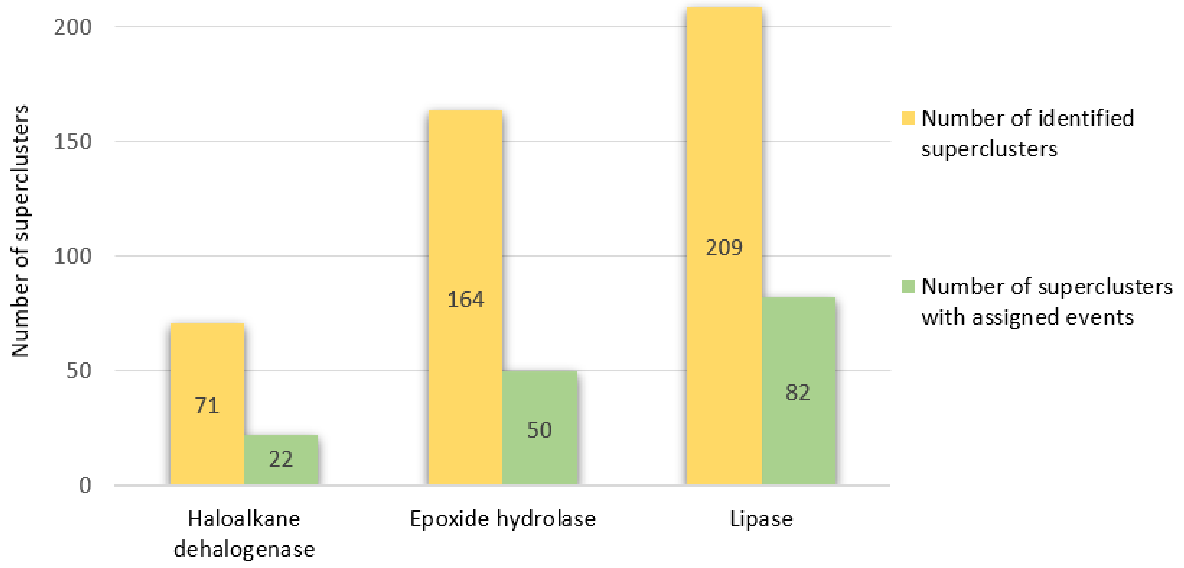


**Fig. 3.** Superclusters created by TransportTools across 50 simulations for each protein system.

#### 3. Runtimes and parallel scalability

We have evaluated the resources required to run the standard TransportTools workflow on all three systems using a moderately powerful workstation (HP Z2 TWR G4). The overall runtime did not exceed 21 hours for the most complex system consisting of millions of tunnels, forming over hundreds of distinct superclusters and tens of thousands of transport events (Fig. 4). Considering the most demanding calculation stages, the calculation of pairwise distances among the tunnel clusters (stage 4) and assignment of transport events to superclusters (stage 9) represents the most time-consuming steps, which are also the most affected by the number of input tunnel clusters and events, as well as their complexity.

Considering the effectiveness of parallel implementation, the TransportTools workflow scales favorably up to eight utilized CPU cores (Fig. 5). Further increases in allocated CPUs results in only minor improvements since the major rate limitations are shifted to I/O operations. Also note that in our hands the effectivness of I/O processing of trajectories in netcdf format depended significantly on the trajectory analysis engine used, in particular for stage 9 of the analyses (Table 1). Regarding the memory requirements, the 1 GB RAM per utilized CPU core was sufficient to perform even the most expensive analysis on the lipase system.

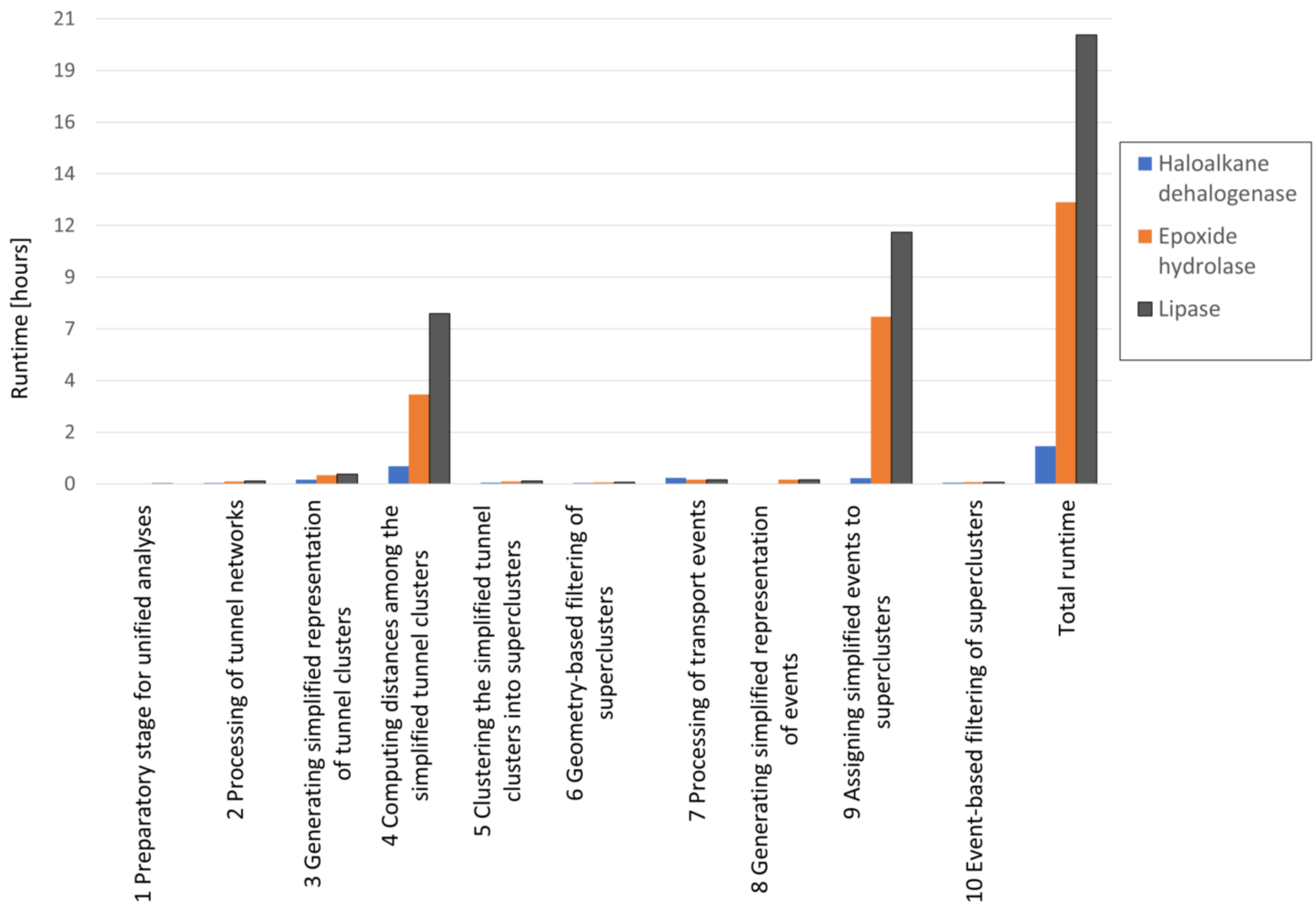


**Fig. 4.** Runtime required for each calculation stage of the standard TransportTools workflow for the three protein systems. The calculation was carried out using a HP Z2 TWR G4 workstation equipped with four Intel Xeon E-2124G CPUs.

A)

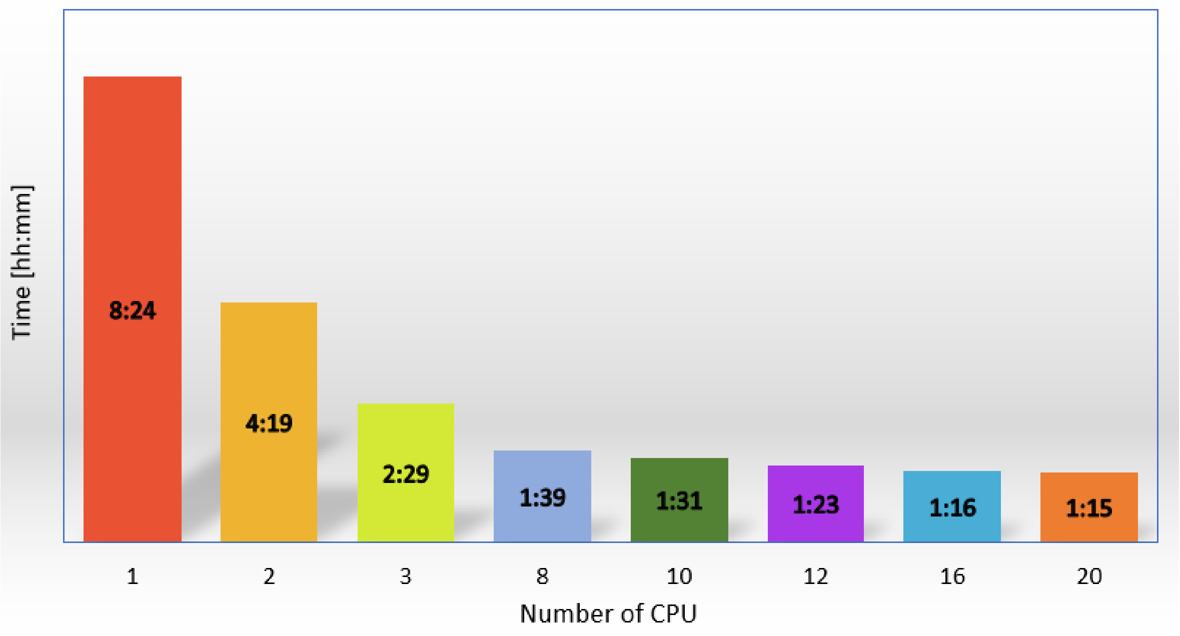


B)

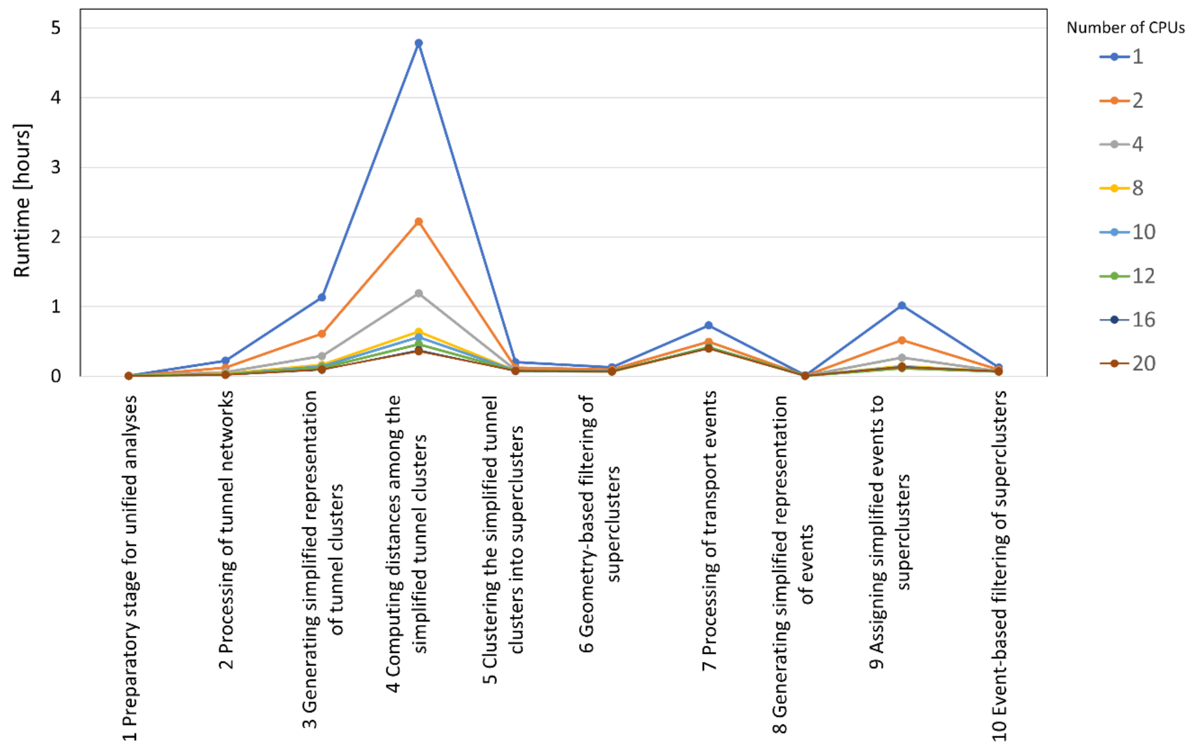


**Fig. 5.** The parallel scalability of TransportTools standard workflow with haloalkane dehalogenase as the benchmarking system. **A)** Scaling of the overall runtime for all ten calculation stages. **B)** Individual runtime for each stage with different CPUs used. For this evaluation, an ETERIO 200TE1 server equipped with Intel Xeon Silver 4114 2.2GHz CPUs was used.

**Table 1. Comparison of runtimes using MDtraj versus pytraj for processing subset of 10 trajectories from haloalkane dehalogenase simulations.**

| Stage / engine | MDtraj | | | pytraj | | |
|---|---|---|---|---|---|---|
| Number of CPUs | 2 | 4 | 8 | 2 | 4 | 8 |
| 1 Preparatory stage for unified analyses | 00:00:03 | 00:00:02 | 00:00:01 | 00:00:03 | 00:00:02 | 00:00:02 |
| 2 Processing of tunnel networks | 00:03:20 | 00:02:01 | 00:01:23 | 00:04:03 | 00:02:01 | 00:01:23 |
| 3 Generating simplified representation of tunnel clusters | 00:05:36 | 00:03:13 | 00:02:10 | 00:06:14 | 00:03:16 | 00:02:09 |
| 4 Computing distances among the simplified tunnel clusters | 00:06:19 | 00:03:23 | 00:02:06 | 00:06:19 | 00:03:29 | 00:01:57 |
| 5 Clustering the simplified tunnel clusters into superclusters | 00:01:12 | 00:00:59 | 00:00:55 | 00:01:20 | 00:01:00 | 00:00:55 |
| 6 Geometry-based filtering of superclusters | 00:00:56 | 00:00:50 | 00:00:48 | 00:01:04 | 00:00:50 | 00:00:48 |
| 7 Processing of transport events | 00:05:31 | 00:04:29 | 00:03:56 | 00:06:09 | 00:04:31 | 00:03:54 |
| 8 Generating simplified representation of events | 00:00:03 | 00:00:03 | 00:00:03 | 00:00:04 | 00:00:03 | 00:00:03 |
| 9 Assigning simplified events to superclusters | 01:49:38 | 01:01:25 | 00:43:42 | 00:17:38 | 00:09:55 | 00:07:09 |
| 10 Event-based filtering of superclusters | 00:00:54 | 00:00:49 | 00:00:48 | 00:01:02 | 00:00:48 | 00:00:48 |
| Total runtime | 02:31:40 | 01:28:01 | 01:02:28 | 01:04:03 | 00:36:31 | 00:25:45 |

#### 4. REFERENCES


Berendsen,H.J.C. *et al.* (1984) Molecular dynamics with coupling to an external bath. *J. Chem. Phys.*, **81**, 3684–3690.

Chovancova,E. *et al.* (2012) CAVER 3.0: A Tool for the Analysis of Transport Pathways in Dynamic Protein Structures. *PLoS Comput. Biol.*, **8**, e1002708.

D.A. Case *et al.* AMBER 18 University of California, San Francisco.

Darden,T. *et al.* (1993) Particle mesh Ewald: An N·log(N) method for Ewald sums in large systems. *J. Chem. Phys.*, **98**, 10089–10092.

Doerr,S. *et al.* (2016) HTMD: High-Throughput Molecular Dynamics for Molecular Discovery. *J. Chem. Theory Comput.*, **12**, 1845–1852.

Grochulski,P. *et al.* (1993) Insights into interfacial activation from an open structure of Candida rugosa lipase. *J. Biol. Chem.*, **268**, 12843–12847.

Grochulski,P. *et al.* (1994) Two conformational states of Candida rugosa lipase. *Protein Sci. Publ. Protein Soc.*, **3**, 82–91.

Hopkins,C.W. *et al.* (2015) Long-Time-Step Molecular Dynamics through Hydrogen Mass Repartitioning. *J. Chem. Theory Comput.*, **11**, 1864–1874.

Izadi,S. *et al.* (2014) Building Water Models: A Different Approach. *J. Phys. Chem. Lett.*, **5**, 3863–3871.

Magdziarz,T. *et al.* (2020) AQUA-DUCT 1.0: structural and functional analysis of macromolecules from an intramolecular voids perspective. *Bioinformatics*, **36**, 2599–2601.

Maier,J.A. *et al.* (2015) ff14SB: Improving the Accuracy of Protein Side Chain and Backbone Parameters from ff99SB. *J. Chem. Theory Comput.*, **11**, 3696–3713.

Mowbray,S.L. *et al.* (2006) X-ray structure of potato epoxide hydrolase sheds light on substrate specificity in plant enzymes. *Protein Sci. Publ. Protein Soc.*, **15**, 1628–1637.

Pavelka,A. *et al.* (2016) CAVER: Algorithms for Analyzing Dynamics of Tunnels in Macromolecules. *IEEE/ACM Trans. Comput. Biol. Bioinform.*, **13**, 505–517.

Ryckaert,J.-P. *et al.* (1977) Numerical integration of the cartesian equations of motion of a system with constraints: molecular dynamics of n-alkanes. *J. Comput. Phys.*, **23**, 327–341.

Salomon-Ferrer,R. *et al.* (2013) Routine Microsecond Molecular Dynamics Simulations with AMBER on GPUs. 2. Explicit Solvent Particle Mesh Ewald. *J. Chem. Theory Comput.*, **9**, 3878–3888.

Stepankova,V. *et al.* (2013) Expansion of Access Tunnels and Active-Site Cavities Influence Activity of Haloalkane Dehalogenases in Organic Cosolvents. *ChemBioChem*, **14**, 890–897.

Zwanzig,R. (1973) Nonlinear generalized Langevin equations. *J. Stat. Phys.*, **9**, 215–220.
