## Supplementary File 6 for "TransportTools: a library for high-throughput analyses of internal voids in biomolecules and ligand transport through them"

---

### Supplementary File 6 – Use Case I:

### Disclosing rare transient tunnels and their usage by water molecules in DhaA dehalogenase

### 1. INTRODUCTION

Rare events such as the opening of transient transport pathways in biomolecules and their usage for the migration of small molecules represent a vital aspect of the function of many biological processes (Zhou and McCammon, 2010; Gora *et al.*, 2013; Marques *et al.*, 2017). However, their detection and analysis constitute a nontrivial task considering the need to compromise between sufficient sampling to visit the desired conformational space and maintaining the adequate resolution and sensitivity of the method for their detection (Klepeis *et al.*, 2009). In many cases, transport pathways can be equipped with multiple gates, e.g., ones located at the beginning and end of such a pathway. The stochastic opening and closing of these gates render the detection of the whole length of the pathway rather unlikely (McCammon, 2011; Gora *et al.*, 2013). In this study, we illustrate the advantages of TransportTools for the investigation of rare pathways from a large dataset of simulations.

### 2. MATERIALS AND METHODS

The initial coordinates for DhaA enzyme were obtained from the PDB database (PDB code: 4E46) and protonated using the H++ webserver at pH 8.5 (Gordon *et al.*, 2005; Anandakrishnan *et al.*, 2012). The system was solvated using the 4-point OPC water model (Izadi *et al.*, 2014) and then neutralized with counterions ($Na^+$ and $Cl^-$) to reach 0.1 M concentration. Using PMEMD and PMEMD.CUDA modules (Salomon-Ferrer *et al.*, 2013) of AMBER18 (D.A. Case *et al.*) with ff14SB force field (Maier *et al.*, 2015), the systems were first energy minimized in several rounds with decreasing harmonic restrains. The minimization procedure was followed by equilibration MD simulations with gradual heating to 310 K under a constant volume using the Langevin thermostat (Zwanzig, 1973), periodic boundary conditions with the particle mesh Ewald method (Darden *et al.*, 1993), and 4 fs time-step enabled by SHAKE and hydrogen mass repartitioning algorithms (Hopkins *et al.*, 2015; Ryckaert *et al.*, 1977). Finally, these simulations were continued with an unrestrained 200 ns simulation at constant pressure and temperature using the weak-coupling barostat and thermostat (Berendsen *et al.*, 1984), storing the frames every 10 ps. The last snapshot served to initiate an adaptive sampling approach using the High-Throughput Molecular Dynamics package (Doerr *et al.*, 2016). During the sampling, the dihedral angles of residues around the active site were employed as the guiding metric. The adaptive sampling was performed for ten epochs, each featuring five parallel simulations of 100 ns, reaching an overall sampling of 5 µs.

For this use case, a subset of ten simulations was analyzed (five simulations from initial and five from final epochs). Tunnel dynamics in each simulation was studied by CAVER 3.0.1 (Pavelka *et al.*, 2016; Chovancova *et al.*, 2012). The starting point for tunnel detection was specified as a center of mass of CG atom of residue 103, CD2 of residue 104, and N atoms of residues 165 and 243. The tunnels were searched in 10,000 snapshots from each MD simulation using a *probe_radius* of 0.7 Å and the default settings otherwise. Tunnels were clustered with agglomerative Murtagh's algorithm using *clustering_threshold* of 3 and *weighting_coefficient* of 2 to increase the importance of tunnel ends during the calculation of pairwise distances among tunnels. Furthermore, AQUA-DUCT 1.0 software (Magdziarz *et al.*, 2020) was used to track the movement of water molecules in each simulation, defining *scope* as the convex hull of backbone atoms and *object* as a sphere of 6 Å from the center of mass of atoms used for the definition of the CAVER starting point. Finally, TransportTools was applied to acquire a comprehensive view of transiently opened tunnels and their utilization for rare events of water migration.

The package (~320 MB) with results generated by CAVER and AQUA-DUCT analyses (inputs for TransportTools) for all ten simulations of the DhaA, as well as the configuration file for TransportTools analysis are available at: https://zenodo.org/record/5642954/files/use_case_1.tar.gz?download=1

### 3. RESULTS AND DISCUSSION

Standard workflow of TransportTools library calculations on selected 10MD simulations resulted in the identification of 47 superclusters of transport pathways in DhaA enzyme, 18 of which had detectable transport of water molecules through them (Fig. 1). Based on these statistics, distinct properties of superclusters in terms of their opening and water transport can be analyzed. While not permanently open, the five most prominent superclusters were readily detected in all investigated simulations, with 10-68% frequency of the open state, and were often massively utilized by water molecules. Those superclusters match the well-known ***p1***, ***p2***, and ***p3*** tunnels and their branches (Klvana *et al.*, 2009). On the other hand, our analysis also elucidates highly transient tunnels which occur very rarely but still can contribute to water transport. Some of these rarely occurring tunnels forming, for example, supercluster 44, represent yet uncharacterized pathways.

```
Total simulations = 10
No snapshots per simulation = 10000

Parameters used for pre-selection of input tunnels for clustering:
length >= 5.00 A
radius >= 0.75 A
curvature <= 2.00

Active tunnel filters:
length = (0.00, 999.00) A
radius = (0.00, 999.00) A
curvature = (1.00, 999.00)
occurred in at least 0 simulations
occurred in at least 0 snapshots
occurred in at least 0.000 snapshots on average per simulation
has at least 1 transport events
has at least 0 entry events
has at least 0 release events

SC_ID, No_Sims, Total_No_Frames, Avg_No_Frames, Avg_BR, StDev, Max_BR, Avg_Len, StDev, Avg_Cur, StDev, Avg_throug,   StDev, Priority, Num_Events, Num_entries, Num_releases
    1,      10,           79411,        7941.1,  1.152, 0.177,  2.937,  14.389, 1.738,   1.257, 0.098,    0.53160, 0.08166,  4.22153,        326,         167,          159
    2,      10,           19081,        1908.1,  0.996, 0.080,  1.605,  15.646, 1.930,   1.319, 0.122,    0.46394, 0.05955,  0.88524,         19,          10,            9
    3,      10,           12910,        1291.0,  1.057, 0.122,  1.670,  15.880, 2.409,   1.297, 0.154,    0.45466, 0.09267,  0.58697,         62,          28,           34
    4,      10,            7019,         701.9,  1.047, 0.096,  1.420,  17.183, 1.570,   1.210, 0.055,    0.40914, 0.04960,  0.28718,          7,           4,            3
    5,      10,            5622,         562.2,  0.964, 0.058,  1.329,  13.945, 1.711,   1.231, 0.118,    0.42897, 0.05657,  0.24117,          4,           2,            2
    6,      10,            1737,         173.7,  0.965, 0.057,  1.307,  17.344, 2.380,   1.259, 0.145,    0.36426, 0.06443,  0.06327,          2,           1,            1
    9,      10,             709,          70.9,  0.997, 0.108,  1.733,  14.863, 2.204,   1.369, 0.127,    0.45033, 0.08448,  0.03193,        169,          88,           81
   10,       6,            1419,         141.9,  0.982, 0.101,  1.665,  24.203, 2.444,   1.243, 0.094,    0.31936, 0.07156,  0.02719,          3,           2,            1
   11,      10,             700,          70.0,  0.965, 0.058,  1.257,  20.363, 2.425,   1.634, 0.173,    0.31430, 0.06522,  0.02200,         13,           8,            5
   13,       1,            1703,         170.3,  1.137, 0.205,  2.468,  14.751, 1.952,   1.236, 0.097,    0.51777, 0.08833,  0.00882,        187,          94,           93
   15,       5,             582,          58.2,  0.985, 0.102,  1.508,  28.889, 6.849,   1.829, 0.623,    0.28246, 0.09226,  0.00822,          1,           1,            0
   20,      10,             103,          10.3,  0.944, 0.048,  1.186,  22.073, 3.200,   1.401, 0.126,    0.26676, 0.07011,  0.00275,          3,           1,            2
   22,       4,             193,          19.3,  0.969, 0.073,  1.305,  29.299, 4.934,   1.993, 0.246,    0.25883, 0.06218,  0.00200,          1,           1,            0
   21,       5,             138,          13.8,  0.949, 0.045,  1.128,  29.449, 5.293,   1.752, 0.197,    0.25429, 0.07104,  0.00175,          1,           1,            0
   31,       6,              22,           2.2,  0.979, 0.071,  1.118,  18.309, 6.452,   1.434, 0.339,    0.38242, 0.12232,  0.00050,          4,           2,            2
   32,       8,              24,           2.4,  0.934, 0.033,  1.030,  26.156, 2.128,   1.658, 0.183,    0.25000, 0.04824,  0.00048,          2,           1,            1
   34,       4,              50,           5.0,  0.936, 0.036,  1.097,  39.320, 6.299,   1.845, 0.316,    0.12147, 0.04260,  0.00024,          1,           0,            1
   44,       2,               2,           0.2,  0.919, 0.004,  0.923,  28.607, 1.298,   1.719, 0.020,    0.18938, 0.02018,  0.00001,          1,           0,            1
Total number of unassigned events:                                                                                                        30,          15,           15
```

**Fig. 1**. **The statistics for pathway superclusters in DhaA enzyme with at least one transport event assigned.**

Focusing on this novel supercluster 44, we can see this pathway is on average over 28 Å long, which provides ample space to accommodate multiple gates, resulting in the sporadic opening of this path in its entire length to be detected by the CAVER package. Hence, an assignment of transport events of any ligands via such pathway is much less likely in a single simulation. To overcome those obstacles, we can benefit from the integrative nature of TransportTools analyses and map ligand migration onto tunnels originating from either identical or different simulations. Inspecting the composition of the supercluster 44 (Fig. 2), we have revealed an example of these analytical obstacles. Here, this supercluster is composed of tunnels originating from two simulations (e10s3_e9s3p0f1650 and e10s5_e9s3p0f1590 from the final epoch), different from the one (e1s1_conf1), in which the water transport event was sampled without detecting the corresponding tunnel (Fig. 3).

```
-------------------------------------------------------------------------------------------------------------------
Supercluster ID 44

Details on tunnel network:
Number of MD simulations = 2
Number of tunnel clusters = 2
Tunnel clusters:
from e10s3_e9s3p0f1650: 53,
from e10s5_e9s3p0f1590: 65,

Details on transport events:
Number of MD simulations = 1
Number of entry events = 0
Number of release events = 1
release: (from Simulation: AQUA-DUCT ID, (Resname:Residue), start_frame->end_frame; ... )
from e1s1_conf1: 36, (WAT:9966), 1329->1335;
-------------------------------------------------------------------------------------------------------------------
```

**Fig. 2. Missing corresponding tunnel cluster in e1s1_conf simulation, which is responsible for the detection of a ligand-transport event in supercluster 44.**

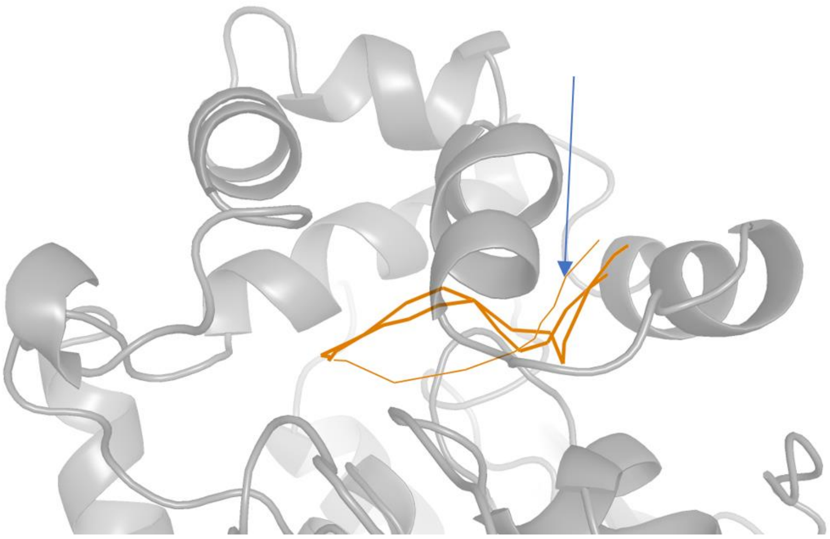

**Fig. 3. An integrated view on the novel transient pathways forming supercluster44.** The DhaA protein is shown as the gray cartoon; the constituent tunnels are visualized as solid orange lines, while the transport event assigned to this supercluster is represented as a thin orange line and highlighted by the blue arrow.

In summary, the integrative analysis of massive MD simulation datasets with TransportTools can enhance the sensitivity of the detection of rare events and their assignment to massively gated transient tunnels, which could otherwise be overlooked during their separate investigation. By combining information on pathways networks and transport events, our method can detect occurrences of water transport from simulations in which the corresponding pathways were not observed due to the closure of one of their gates. This obstacle is alleviated by mapping the events to the pathways detected within superclusters unified across the entire simulation dataset. In this way, the high-throughput analysis with TransportTools prevents the omission of rare and challenging to capture tunnel openings and their utilization, improving the overall process of transient pathway exploration. Knowledge of these functional ligand-transport pathways contributes to the understanding of the molecular mechanism behind the protein function or, conversely, dysfunction (Haberland *et al.*, 2009; Silman and Sussman, 2005; Marques *et al.*, 2016). Additionally, such pathways constitute relevant yet novel targets for drug discovery (Marques *et al.*, 2017) as well as protein engineering efforts (Kokkonen *et al.*, 2019).

#### 4. REFERENCES


Anandakrishnan,R. *et al.* (2012) H++ 3.0: automating pK prediction and the preparation of biomolecular structures for atomistic molecular modeling and simulations. *Nucleic Acids Res.*, **40**, W537–W541.

Berendsen,H.J.C. *et al.* (1984) Molecular dynamics with coupling to an external bath. *J. Chem. Phys.*, **81**, 3684–3690.

Chovancova,E. *et al.* (2012) CAVER 3.0: A Tool for the Analysis of Transport Pathways in Dynamic Protein Structures. *PLoS Comput. Biol.*, **8**, e1002708.

D.A. Case *et al.* AMBER 18 University of California, San Francisco.

Darden,T. *et al.* (1993) Particle mesh Ewald: An N·log(N) method for Ewald sums in large systems. *J. Chem. Phys.*, **98**, 10089–10092.

Doerr,S. *et al.* (2016) HTMD: High-Throughput Molecular Dynamics for Molecular Discovery. *J. Chem. Theory Comput.*, **12**, 1845–1852.

Gora,A. *et al.* (2013) Gates of Enzymes. *Chem. Rev.*, **113**, 5871–5923.

Gordon,J.C. *et al.* (2005) H++: a server for estimating p Ka s and adding missing hydrogens to macromolecules. *Nucleic Acids Res.*, **33**, W368–W371.

Haberland,M. *et al.* (2009) The many roles of histone deacetylases in development and physiology: implications for disease and therapy. *Nat. Rev. Genet.*, **10**, 32–42.

Hopkins,C.W. *et al.* (2015) Long-Time-Step Molecular Dynamics through Hydrogen Mass Repartitioning. *J. Chem. Theory Comput.*, **11**, 1864–1874.

Izadi,S. *et al.* (2014) Building Water Models: A Different Approach. *J. Phys. Chem. Lett.*, **5**, 3863–3871.

Klepeis,J.L. *et al.* (2009) Long-timescale molecular dynamics simulations of protein structure and function. *Curr. Opin. Struct. Biol.*, **19**, 120–127.

Klvana,M. *et al.* (2009) Pathways and mechanisms for product release in the engineered haloalkane dehalogenases explored using classical and random acceleration molecular dynamics simulations. *J. Mol. Biol.*, **392**, 1339–1356.

Kokkonen,P. *et al.* (2019) Engineering enzyme access tunnels. *Biotechnol. Adv.*, **37**, 107386.

Magdziarz,T. *et al.* (2020) AQUA-DUCT 1.0: structural and functional analysis of macromolecules from an intramolecular voids perspective. *Bioinformatics*, **36**, 2599–2601.

Maier,J.A. *et al.* (2015) ff14SB: Improving the Accuracy of Protein Side Chain and Backbone Parameters from ff99SB. *J. Chem. Theory Comput.*, **11**, 3696–3713.

Marques,S. *et al.* (2016) Role of tunnels and gates in enzymatic catalysis. In, Svendsen,A. (ed), *Understanding Enzymes: Function, Design, Engineering, and Analysis*. CRC Press, Boca Raton, pp. 421–463.

Marques,S.M. *et al.* (2017) Enzyme Tunnels and Gates As Relevant Targets in Drug Design. *Med. Res. Rev.*, **37**, 1095–1139.

McCammon,J.A. (2011) Gated Diffusion-controlled Reactions. *BMC Biophys.*, **4**, 4.

Pavelka,A. *et al.* (2016) CAVER: Algorithms for Analyzing Dynamics of Tunnels in Macromolecules. *IEEE/ACM Trans. Comput. Biol. Bioinform.*, **13**, 505–517.

Ryckaert,J.-P. *et al.* (1977) Numerical integration of the cartesian equations of motion of a system with constraints: molecular dynamics of n-alkanes. *J. Comput. Phys.*, **23**, 327–341.

Salomon-Ferrer,R. *et al.* (2013) Routine Microsecond Molecular Dynamics Simulations with AMBER on GPUs. 2. Explicit Solvent Particle Mesh Ewald. *J. Chem. Theory Comput.*, **9**, 3878–3888.

Silman,I. and Sussman,J.L. (2005) Acetylcholinesterase: ‘classical’ and ‘non-classical’ functions and pharmacology. *Curr. Opin. Pharmacol.*, **5**, 293–302.

Zhou,H.-X. and McCammon,J.A. (2010) The gates of ion channels and enzymes. *Trends Biochem. Sci.*, **35**, 179–185.

Zwanzig,R. (1973) Nonlinear generalized Langevin equations. *J. Stat. Phys.*, **9**, 215–220.
