## Supplementary File 7 for "TransportTools: a library for high-throughput analyses of internal voids in biomolecules and ligand transport through them"

### Supplementary File 7 – Use Case II:

### Understanding the effect of mutations by contrasting simulations of three different variants of LinB dehalogenase

### 1. INTRODUCTION

The structure and dynamics of tunnel networks in enzymes can be markedly affected by mutations, leading to an increase or decrease in the efficacy of the enzyme (Kokkonen *et al.*, 2018). To understand the effect of mutations on the tunnel networks and transport through them, the consistent comparison between independent datasets obtained from multiple MD trajectories is critical and represents a considerable challenge (Marques *et al.*, 2016). In this study, we illustrate the ability of TransportTools to combine such complex information about tunnel networks and transport events from independent MD simulations of haloalkane dehalogenase LinB wild-type and two of its mutants—LinB32 with a closed primary tunnel and LinB86 with the same primary tunnel closed and newly opened a *de novo* tunnel (Brezovsky *et al.*, 2016). Taking advantage of the comparative analysis feature of TransportTools, we gathered information on the individual tunnels in LinB variants and studied the effect of introduced mutations on the tunnel dynamics and transport process in detail.

### 2. MATERIALS AND METHODS

MD simulations on three LinB enzyme variants we performed: wild-type (PDB code: 1MJ5), LinB32 (PDB code: 4WDQ), and LinB86 (PDB code: 5LKA) (Brezovsky *et al.*, 2016). The systems were protonated using the H++ webserver at pH 8.5 (Gordon *et al.*, 2005; Anandakrishnan *et al.*, 2012). The molecular systems were solvated using 4-point OPC water models (Izadi *et al.*, 2014) and then neutralized with counter ions ($Na^+$ and $Cl^-$) to reach 0.1 M concentration. Using PMEMD and PMEMD.CUDA modules (Salomon-Ferrer *et al.*, 2013) of AMBER18 (D.A. Case *et al.*) with ff14SB force field (Maier *et al.*, 2015), the systems were first energy minimized in several rounds with decreasing harmonic restrains. The minimization procedure was followed by equilibration MD simulations with gradual heating to 310 K under a constant volume using the Langevin thermostat (Zwanzig, 1973), periodic boundary conditions with the particle mesh Ewald method (Darden *et al.*, 1993), and 4 fs time-step enabled by SHAKE and hydrogen mass repartitioning algorithms (Hopkins *et al.*, 2015; Ryckaert *et al.*, 1977). Finally, these simulations were continued with an unrestrained 100 ns production simulation at constant pressure and temperature, storing the frames every 20 ps.

Next, the generated MD trajectories were analyzed by CAVER 3.0.1 (Pavelka *et al.*, 2016; Chovancova *et al.*, 2012) to study the mutational effects on tunnel dynamics. The starting point for tunnel detection was specified as a center of mass of residues 37, 108, and 271. The tunnels were searched in 5,000 snapshots from each MD simulation using a *probe_radius* of 0.7 Å and the default settings otherwise. Tunnels were clustered with agglomerative Murtagh's algorithm using *clustering_threshold* of 3 and *weighting_coefficient* of 2 to increase the importance of tunnel ends during the calculation of pairwise distances among tunnels. Also, the identification of tunnel bottleneck residues was enabled. Furthermore, AQUA-DUCT 1.0 software (Magdziarz *et al.*, 2020) was used to track the movement of water molecules in each simulation, defining *scope* as the convex hull of backbone atoms and *object* as a sphere of 6 Å from the center of mass of residues 37, 108, and 271. Finally, TransportTools library was applied to combine the information gathered from CAVER and AQUA-DUCT, taking advantage of the comparative analysis feature to consistently investigate the effect of mutations on both the tunnel network and its utilization for the transport of water molecules.

The package (~840 MB) with results generated by CAVER and AQUA-DUCT analyses (inputs for Transport tools), the configuration file for TransportTools as well as analyzed trajectories from which the irrelevant water molecules were stripped away are available for all three investigated LinB dehalogenase variants at: https://zenodo.org/record/5642954/files/use_case_2.tar.gz?download=1

### 3. RESULTS AND DISCUSSION

Based on the comprehensive TransportTools analysis, the eight most relevant tunnel superclusters detected in the analyzed set of three MD simulations of three LinB variants were visualized (Fig. 1). Importantly, each visualized supercluster corresponds well with already identified tunnels (Brezovsky *et al.*, 2016; Klvana *et al.*, 2009): ***p1*** (***a*** – lime green, and ***b*** – blue), ***p2*** (***a*** – green, ***b*** – cyan and yellow, ***c*** – red, and ***c**** – light purple) and ***p3*** (pink). Regarding the quantitative data on tunnels, the generated TransportTools summary (Table 1) clearly highlights the differences in tunnel dynamics caused by the introduced mutations. In the LinB wild-type (Table 1), ***p1b*** tunnel (supercluster 1) represents the primary conduit either in terms of its opening (4655/5000 snapshots) or the use for water transport (273 events). In contrast to LinB32 (closed) and LinB86 (open) variants, the L177W mutations were introduced to the ***p1*** tunnel to limit its utilization (Brezovsky *et al.*, 2016). This outcome can be systematically observed in the summary data (Table 1) – ***p1b*** tunnel opens in 402 and 368 snapshots only for LinB32 and LinB86, respectively, with no corresponding transport event detected. Furthermore, LinB86 was engineered to support the use of ***p3*** tunnel by additional three mutations (W140A+F143L+I211L), resulting in a more open geometry and is known to be utilized by water more frequently (Brezovsky *et al.*, 2016). TransportTools guided analysis elucidates that indeed the ***p3*** tunnel (supercluster 6) is approximately four times more frequently open in the LinB86 variant compared to the remaining two proteins (Table 1). Additionally, these relatively short simulations enabled the detection of water transport via the ***p3*** tunnel only in the case of the open LinB86 variant.

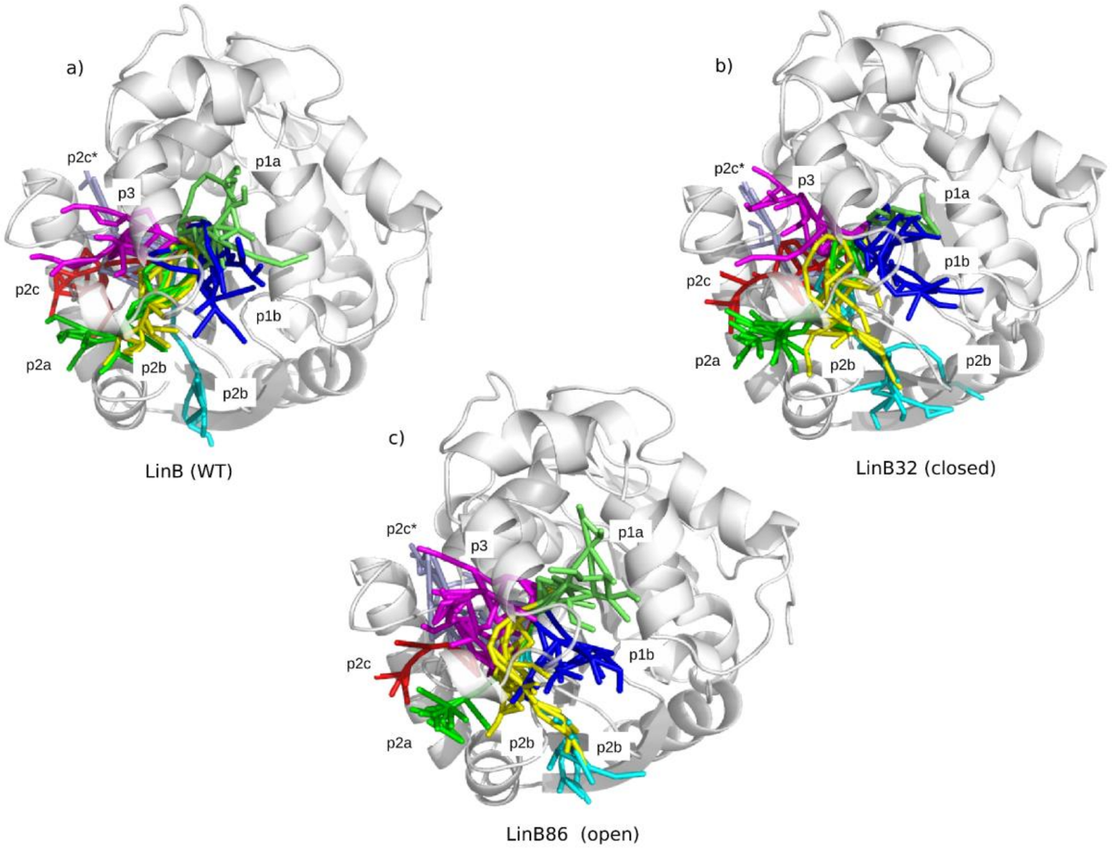


**Fig. 1**. **Representation of the volume occupied by the seven most relevant superclusters detected during the comparative analysis**. **a)** LinB wild-type, **b)** LinB32 with the narrowed ***p1*** tunnels, and **c)** LinB86 (open) with the narrowed ***p1*** tunnels and additionally a *de novo* opened ***p3*** tunnel

**Statistics for: LinB wild-type ['md1_1mj5']**

No snapshots per simulation = 5000

| SC_ID, | Total_No_Frames, | BottleRadius, | Max_BottleRadius, | Length, | Curvature, | Throughput, | Num_Events, | Num_entries, | Num_releases, | Tunnel_name |
|---|---|---|---|---|---|---|---|---|---|---|
| 1, | 4655, | 1.40±0.28, | 2.38, | 12±2, | 1.2±0.1, | 0.66±0.09, | 273, | 136, | 137, | *p1b* |
| 2, | 1700, | 1.18±0.20, | 1.79, | 19±2, | 1.3±0.1, | 0.49±0.09, | 16, | 10, | 6, | *p2a* |
| 3, | 201, | 0.97±0.06, | 1.22, | 22±2, | 1.3±0.1, | 0.33±0.05, | -, | -, | - | *p2c* |
| 5, | 174, | 0.95±0.05, | 1.11, | 29±2, | 1.6±0.2, | 0.26±0.04, | 1, | 0, | 1, | *p2b* |
| 4, | 516, | 1.11±0.19, | 1.78, | 17±2, | 1.4±0.1, | 0.50±0.08, | 6, | 2, | 4, | *p2b* |
| 6, | 163, | 0.96±0.07, | 1.25, | 16±2, | 1.3±0.1, | 0.42±0.07, | -, | -, | - | *p3* |
| 7, | 137, | 0.95±0.05, | 1.14, | 25±4, | 1.6±0.2, | 0.27±0.06, | -, | -, | - | *p2c** |
| 8, | 103, | 1.04±0.13, | 1.47, | 19±3, | 1.5±0.2, | 0.45±0.09, | 3, | 1, | 2, | *p1a* |

---

**Statistics for: LinB32 (closed) ['md2_4wdq']**

No snapshots per simulation = 5000

| SC_ID, | Total_No_Frames, | BottleRadius, | Max_BottleRadius, | Length, | Curvature, | Throughput, | Num_Events, | Num_entries, | Num_releases, | Tunnel_name |
|---|---|---|---|---|---|---|---|---|---|---|
| 1, | 402, | 1.00±0.10, | 1.43, | 13±2, | 1.3±0.1, | 0.49±0.07, | -, | -, | - | *p1b* |
| 2, | 2606, | 1.17±0.20, | 1.91, | 20±2, | 1.3±0.1, | 0.45±0.09, | 41, | 22, | 19, | *p2a* |
| 3, | 2312, | 1.03±0.10, | 1.52, | 21±2, | 1.3±0.1, | 0.38±0.06, | 3, | 1, | 2, | *p2c* |
| 5, | 1808, | 1.00±0.08, | 1.33, | 26±2, | 1.8±0.3, | 0.30±0.04, | 2, | 1, | 1, | *p2b* |
| 4, | 1252, | 1.05±0.14, | 1.56, | 18±3, | 1.5±0.2, | 0.42±0.07, | 4, | 2, | 2, | *p2b* |
| 6, | 160, | 0.96±0.05, | 1.12, | 17±2, | 1.3±0.1, | 0.36±0.05, | -, | -, | - | *p3* |
| 7, | 423, | 0.97±0.06, | 1.21, | 27±3, | 1.7±0.2, | 0.26±0.05, | -, | -, | - | *p2c** |
| 8, | 83, | 1.00±0.12, | 1.46, | 14±1, | 1.3±0.1, | 0.48±0.06, | -, | -, | - | *p1a* |

---

**Statistics for: LinB86 (open) ['md3_5lka']**

No snapshots per simulation = 5000

| SC_ID, | Total_No_Frames, | BottleRadius, | Max_BottleRadius, | Length, | Curvature, | Throughput, | Num_Events, | Num_entries, | Num_releases, | Tunnel_name |
|---|---|---|---|---|---|---|---|---|---|---|
| 1, | 368, | 0.99±0.09, | 1.58, | 13±2, | 1.3±0.1, | 0.47±0.07, | -, | -, | - | *p1b* |
| 2, | 2170, | 1.22±0.22, | 1.99, | 19±2, | 1.3±0.1, | 0.50±0.08, | 40, | 20, | 20, | *p2a* |
| 3, | 1918, | 1.03±0.10, | 1.50, | 23±2, | 1.4±0.1, | 0.37±0.05, | 2, | 1, | 1, | *p2c* |
| 5, | 1986, | 1.04±0.09, | 1.43, | 25±2, | 1.7±0.2, | 0.33±0.05, | 3, | 2, | 1, | *p2b* |
| 4, | 806, | 1.03±0.11, | 1.48, | 18±2, | 1.5±0.2, | 0.43±0.07, | 4, | 3, | 1, | *p2b* |
| 6, | 684, | 0.99±0.08, | 1.33, | 16±2, | 1.3±0.1, | 0.42±0.07, | 1, | 1, | 0, | *p3* |
| 7, | 701, | 0.97±0.06, | 1.30, | 29±3, | 2.0±0.2, | 0.24±0.04, | -, | -, | - | *p2c** |
| 8, | 493, | 0.99±0.08, | 1.37, | 15±2, | 1.4±0.2, | 0.44±0.06, | -, | -, | - | *p1a* |

---

**Table 1. Quantitative data for the eight most relevant tunnel superclusters for three LinB variants adopted from TransportTools statistics.**

To further study the effect of mutations, we have extracted the information about the bottleneck residues for each supercluster corresponding to modified tunnels (supercluster 1 for ***p1b*** and supercluster 6 for ***p3***) and focused on these residues that were contributing to tunnel bottlenecks most frequently in the analyzed simulations. Upon introducing the L177W mutation in the ***p1b*** tunnel, the residues most frequently forming the tunnel bottleneck changed to 177, 247, 248, and 272 in LinB32 and LinB86 variants compared to 147, 151, 173, and 177 in the wild-type (Fig. 2a). Similarly, by analyzing the effects of mutation in the ***p3*** tunnel (Fig. 2b), the shift in the bottleneck residues could be observed. Here, we emphasize tryptophan at position 140, as it significantly affects the bottleneck of the ***p3*** tunnel in wild-type and LinB32 variants, while upon its mutation to alanine in LinB86, its contribution is negligible.

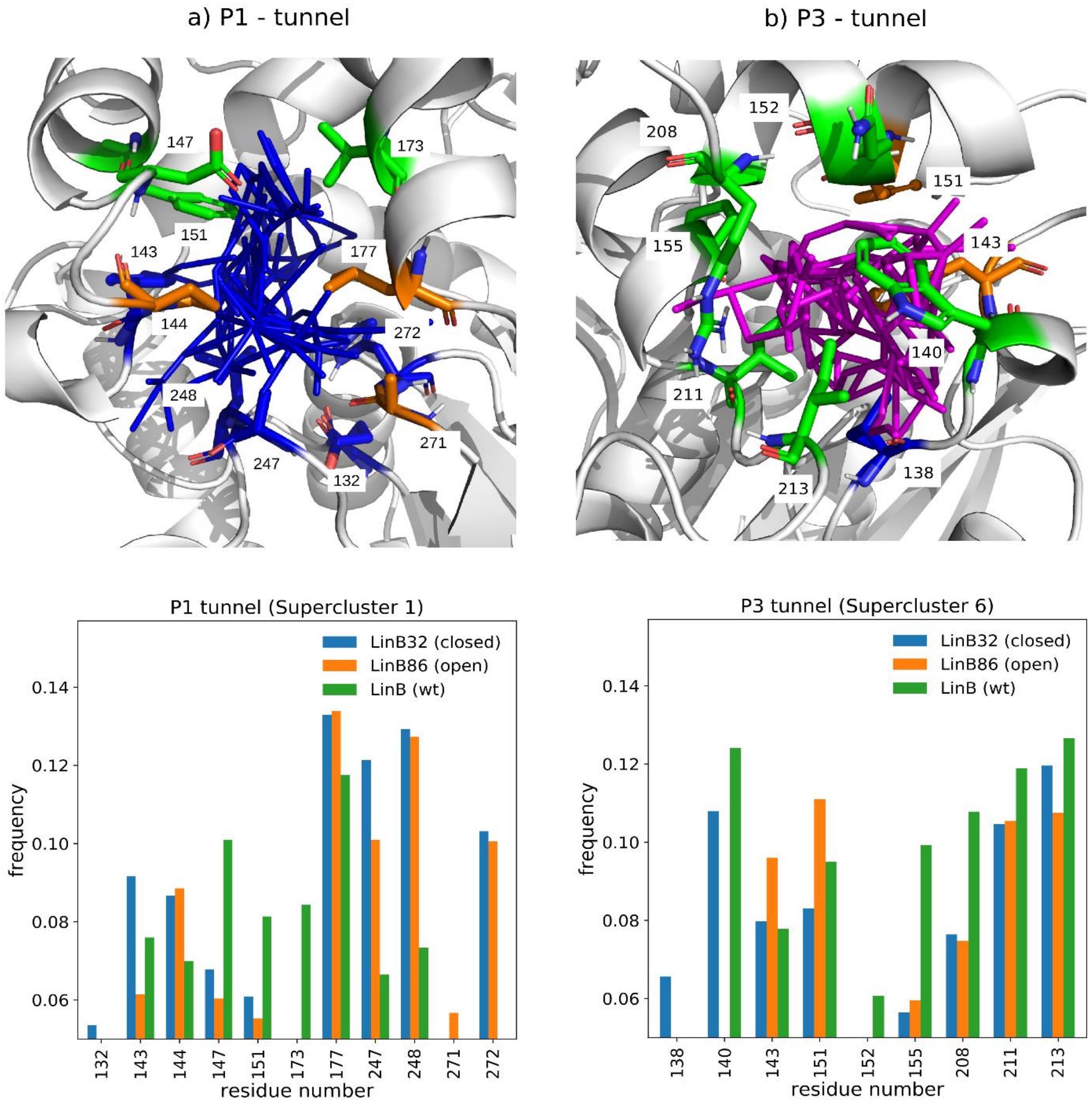


**Fig. 2**. **Analysis of the bottleneck residues of a) *p1* and b) *p3* tunnels modulated upon mutations.** The upper panel represents the most contributing bottleneck residues as sticks for ***p1b*** (supercluster 1, blue lines) and ***p3*** (supercluster 6, pink lines) tunnels in LinB wild-type (green), LinB32 (blue), and LinB86 (orange). The bottom panel presents the residues forming the tunnel bottleneck (only residues with frequency > 5 % are shown).

In summary, TransportTools supported analysis enabled a comparative investigation of three independent simulations of LinB protein variants, uncovering all critical changes in the tunnel dynamics, geometry, and water transport upon introduced perturbations. The obtained insights explain molecular mechanisms behind the effect of mutations on transport components governing the function of these enzymes, which are consistent with the current knowledge (Brezovsky *et al.*, 2016; Kokkonen *et al.*, 2018).
