## Supplementary File 8 for "TransportTools: a library for high-throughput analyses of internal voids in biomolecules and ligand transport through them"

#### Supplementary File 8 – Use Case III:

#### Inferring selectivity of transport pathways in LinB86 dehalogenase for a substrate molecule from almost 600 simulations

### 1. INTRODUCTION

Proteins interact with various small molecules to perform their biological activities. Notably, many proteins have their active sites buried in a deep hydrophobic core linked to the surface by transport paths known as tunnels or channels. The molecular mechanisms behind the utilization of transport tunnels by small molecules are of growing interest in structure-based drug design and protein engineering (Marques *et al.*, 2017). Hence, it is crucial to understand how small molecules traverse through transport paths and determine the ones preferred by particular ligands (Kingsley and Lill, 2015). In this study, we analyzed the utilization of transport tunnels for entry and release of the 1,2-dibromoethane substrate to and from the active site of the mutant haloalkane dehalogenase LinB86, which features a complex network of tunnels (Brezovsky *et al.*, 2016). By integrating outcomes from almost 600 adaptive sampling simulations designed to facilitate substrate transport (Doerr and De Fabritiis, 2014) with the data on the tunnel network using TransportTools library, details on the selectivity of the tunnels for the substrate molecule readily become available.

### 2. MATERIALS AND METHODS

The input model was based on the crystallographic structure of the haloalkane dehalogenase mutant enzyme LinB86 (PDB code: 5LKA). The protein was modeled using the ff14SB (Maier *et al.*, 2015) force field and the structure of 1,2-dibromoethane (DBE) substrate was retrieved from PubChem and modeled using the General Amber Force Field – GAFF (Wang *et al.*, 2004). The protein structure was protonated at pH 8.5 using H++ webserver (Anandakrishnan *et al.*, 2012; Gordon *et al.*, 2005). The DBE substrate was initially docked to LinB86 active site using Autodock4 program (Morris *et al.*, 2009) and the 30 modes with the lowest binding energies were selected. The 30 complexes together with one structure of free LinB86 without DBE molecule were solvated using 4-point OPC water model (Izadi *et al.*, 2014) and then neutralized with counterions ($Na^+$ and $Cl^-$) to reach 0.1 M concentration. Using PMEMD and PMEMD.CUDA modules (Salomon-Ferrer *et al.*, 2013) of AMBER18 (D.A. Case *et al.*) with ff14SB force field (Maier *et al.*, 2015), the systems were first energy minimized in several rounds with decreasing harmonic restrains. The minimization procedure was followed by equilibration MD simulations with gradual heating to 310 K under a constant volume using the Langevin thermostat (Zwanzig, 1973), periodic boundary conditions with the particle mesh Ewald method (Darden *et al.*, 1993), and 4 fs time-step enabled by SHAKE and hydrogen mass repartitioning algorithms (Hopkins *et al.*, 2015; Ryckaert *et al.*, 1977). Finally, these simulations were continued with an unrestrained 100 ns simulation at constant pressure and temperature using the weak-coupling barostat and thermostat (Berendsen *et al.*, 1984), storing the frames every 200 ps. The last snapshot from each simulation served to initiate an adaptive sampling approach using High-Throughput Molecular Dynamics package (Doerr *et al.*, 2016). During the sampling, the distances between all Cα carbon atoms of protein residues and four heavy atoms of the substrate were employed as the guiding metric. The adaptive sampling was performed for 30 epochs, each featuring 30 parallel simulations of 50 ns length, resulting in a total sampling of 45 µs.

Next, the simulation of free LinB86 was analyzed by CAVER 3.0.1 (Pavelka *et al.*, 2016; Chovancova *et al.*, 2012) to study the tunnel dynamics. The starting point for tunnel detection was specified as a center of mass of residues 37, 107, 108, and 271. The tunnels were searched in 500 snapshots from each MD simulation using a *probe_radius* of 0.5 Å and the default settings otherwise. Tunnels were clustered with agglomerative Murtagh's algorithm using *clustering_threshold* of 4 and *weighting_coefficient* of 2 to increase the importance of tunnel ends during the calculation of pairwise distances among tunnels. Furthermore, AQUA-DUCT 1.0 software (Magdziarz *et al.*, 2020) was used to track the movement of DBE molecules in each simulation, defining *scope* as the convex hull of backbone atoms deflated by 1 Å (*scope_convexhull_inflate* = -1) and *object* as a sphere of 6 Å from the center of mass of atoms used for the definition of the CAVER starting point. Finally, 593 trajectories that exhibited transport events detected by AQUA-DUCT and CAVER tunnel network were processed by TransportTools to reveal the preference of transport pathways for the substrate molecules.

The package (~70 MB) with the results from CAVER analysis of 100 ns simulation and from tracking the DBE substrate transport in 593 simulations by AQUA-DUCT (inputs for TransportTools), and the configuration file for TransportTools are available at:
https://zenodo.org/record/5642954/files/use_case_3.tar.gz?download=1

#### 3. RESULTS AND DISCUSSION

Pathways identified by CAVER cover the ***p1a***, ***p1b***, ***p2,*** and ***p3*** tunnels and their branches (Fig. 1). Identified pathways correspond well with the literature data on LinB86 protein variant (Brezovsky *et al.*, 2016), confirming that the 100 ns simulation was sufficient to sample the primary transport pathways in the investigated system.

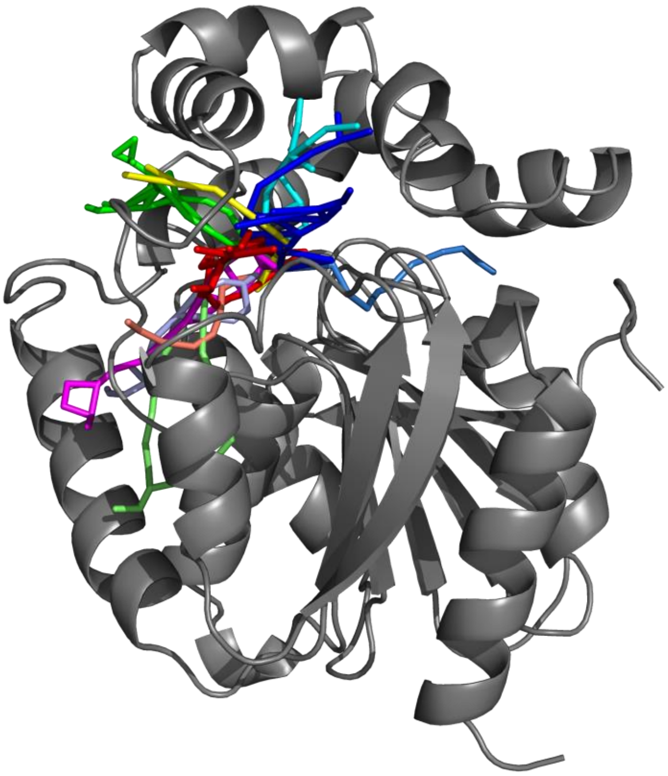

**Fig. 1. Visualization volume occupied by pathway superclusters in LinB86.** Superclusters corresponding to ***p1a*** and ***p1b*** tunnels are shown in cyan and blue. The branches of ***p2*** tunnel are shown in red, orange, pink and light green, while ***p3*** tunnel's branches are in green and yellow.

With 900 trajectories from adaptive sampling, AQUA-DUCT elucidated 593 trajectories with DBE molecules traced inside the protein body. Hence, based on this extensive set of relevant simulations with transport events detected, we performed a massive analysis with TransportTools to combine tunnel network data with the information on transport events. This assignment revealed 40 events of complete substrate transport between the active site of LinB86 and the bulk solvent. 33 events represented the release of DBE molecule from the active site, which is consistent with starting the initial simulations from complexes of DBE bound in the active site. However, we have also observed six instances of substrate entry to the active site corresponding to rebinding events. 33 transport events were assigned to five pathway superclusters, while seven events could not be unambiguously assigned to any supercluster (Fig. 2 and 3). From the properly assigned events, supercluster 1 (***p1b*** pathway) and supercluster 4 (***p1a*** pathway) were more frequently utilized for the transport of DBE when compared to supercluster 2 and 5 (***p3*** pathway) and supercluster 3 (***p2*** pathway). Inferred preferences of pathways for DBE substrates are well aligned with the previous computational and experimental reports on LinB86 protein (Brezovsky *et al.*, 2016).

| SC_ID | No_Sims | Total_No_Frames | Avg_No_Frames | Avg_BR | StDev | Max_BR | Avg_Len | StDev | Avg_Cur | StDev | Avg_throug | StDev | Priority | Num_Events | Num_entries | Num_releases |
|---|---|---|---|---|---|---|---|---|---|---|---|---|---|---|---|---|
| 1 | 1 | 292 | 292.0 | 1.258 | 0.209 | 1.795 | 12.622 | 1.824 | 1.236 | 0.116 | 0.61424 | 0.08206 | 0.35872 | 14 | 2 | 12 |
| 2 | 1 | 204 | 204.0 | 1.042 | 0.110 | 1.370 | 15.412 | 2.258 | 1.283 | 0.121 | 0.47286 | 0.06976 | 0.19293 | 6 | 2 | 4 |
| 3 | 1 | 125 | 125.0 | 1.172 | 0.230 | 1.804 | 16.384 | 1.922 | 1.309 | 0.108 | 0.50129 | 0.10573 | 0.12532 | 1 | 0 | 1 |
| 4 | 1 | 45 | 45.0 | 0.999 | 0.087 | 1.321 | 14.290 | 2.017 | 1.361 | 0.144 | 0.46566 | 0.06640 | 0.04191 | 11 | 3 | 8 |
| 5 | 1 | 17 | 17.0 | 0.965 | 0.067 | 1.200 | 18.290 | 2.414 | 1.618 | 0.158 | 0.42695 | 0.06587 | 0.01452 | 1 | 0 | 1 |
| 6 | 1 | 8 | 8.0 | 0.954 | 0.037 | 0.998 | 23.959 | 3.343 | 1.455 | 0.196 | 0.29030 | 0.05081 | 0.00464 | 0 | 0 | 0 |
| 7 | 1 | 3 | 3.0 | 0.953 | 0.051 | 1.022 | 27.286 | 0.795 | 1.855 | 0.146 | 0.24393 | 0.04250 | 0.00146 | 0 | 0 | 0 |
| 8 | 1 | 2 | 2.0 | 0.944 | 0.005 | 0.949 | 29.246 | 2.363 | 1.537 | 0.026 | 0.23446 | 0.02134 | 0.00094 | 0 | 0 | 0 |
| 9 | 1 | 1 | 1.0 | 0.952 | 0.000 | 0.952 | 18.294 | 0.000 | 1.293 | 0.000 | 0.26785 | 0.00000 | 0.00054 | 0 | 0 | 0 |
| 10 | 1 | 1 | 1.0 | 0.931 | 0.000 | 0.931 | 22.814 | 0.000 | 1.437 | 0.000 | 0.20132 | 0.00000 | 0.00040 | 0 | 0 | 0 |
| Total number of unassigned events: | | | | | | | | | | | | | | 7 | 1 | 6 |

**Fig. 2**. **Assignment of 40 transport events obtained from 593 simulations with traced DBE ligand to tunnel superclusters.**

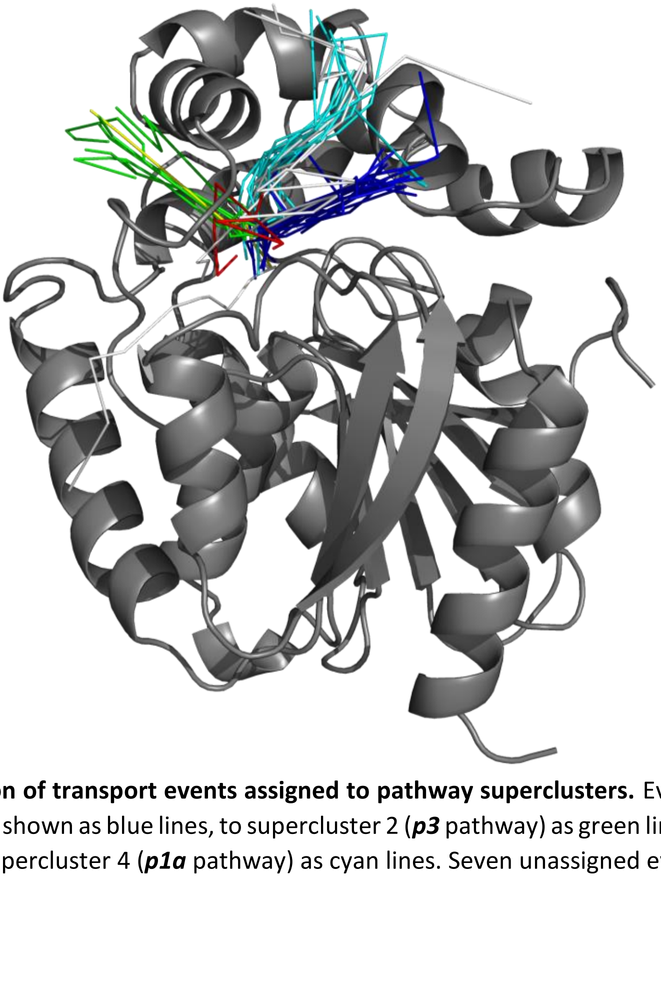

**Fig. 3**. **Visualization of transport events assigned to pathway superclusters.** Events assigned to supercluster 1 (***p1b*** pathway) are shown as blue lines, to supercluster 2 (***p3*** pathway) as green lines, supercluster 3 (***p2*** pathway) as red lines and supercluster 4 (***p1a*** pathway) as cyan lines. Seven unassigned events are shown as white lines.

To conclude, TransportTools ability to combine massive MD simulations sampling ligand transport with information on tunnel networks enabled the recognition of pathways in LinB86 enzyme, which were preferred by the DBE substrate. These pathways represent promising targets for the design of inhibitors (Marques *et al.*, 2017) and protein engineering efforts targeting enzyme selectivity (Kokkonen *et al.*, 2019; Brezovsky *et al.*, 2016).

#### 4. REFERENCES


Anandakrishnan,R. *et al.* (2012) H++ 3.0: automating pK prediction and the preparation of biomolecular structures for atomistic molecular modeling and simulations. *Nucleic Acids Res.*, **40**, W537–W541.

Berendsen,H.J.C. *et al.* (1984) Molecular dynamics with coupling to an external bath. *J. Chem. Phys.*, **81**, 3684–3690.

Brezovsky,J. *et al.* (2016) Engineering a de novo transport tunnel. *ACS Catal.*, **6**, 7597–7610.

Chovancova,E. *et al.* (2012) CAVER 3.0: A Tool for the Analysis of Transport Pathways in Dynamic Protein Structures. *PLoS Comput. Biol.*, **8**, e1002708.

D.A. Case *et al.* AMBER 18 University of California, San Francisco.

Darden,T. *et al.* (1993) Particle mesh Ewald: An N·log(N) method for Ewald sums in large systems. *J. Chem. Phys.*, **98**, 10089–10092.

Doerr,S. *et al.* (2016) HTMD: High-Throughput Molecular Dynamics for Molecular Discovery. *J. Chem. Theory Comput.*, **12**, 1845–1852.

Doerr,S. and De Fabritiis,G. (2014) On-the-Fly Learning and Sampling of Ligand Binding by High-Throughput Molecular Simulations. *J. Chem. Theory Comput.*, **10**, 2064–2069.

Gordon,J.C. *et al.* (2005) H++: a server for estimating p Ka s and adding missing hydrogens to macromolecules. *Nucleic Acids Res.*, **33**, W368–W371.

Hopkins,C.W. *et al.* (2015) Long-Time-Step Molecular Dynamics through Hydrogen Mass Repartitioning. *J. Chem. Theory Comput.*, **11**, 1864–1874.

Izadi,S. *et al.* (2014) Building Water Models: A Different Approach. *J. Phys. Chem. Lett.*, **5**, 3863–3871.

Kingsley,L.J. and Lill,M.A. (2015) Substrate tunnels in enzymes: Structure–function relationships and computational methodology. *Proteins Struct. Funct. Bioinforma.*, **83**, 599–611.

Kokkonen,P. *et al.* (2019) Engineering enzyme access tunnels. *Biotechnol. Adv.*, **37**, 107386.

Magdziarz,T. *et al.* (2020) AQUA-DUCT 1.0: structural and functional analysis of macromolecules from an intramolecular voids perspective. *Bioinformatics*, **36**, 2599–2601.

Maier,J.A. *et al.* (2015) ff14SB: Improving the Accuracy of Protein Side Chain and Backbone Parameters from ff99SB. *J. Chem. Theory Comput.*, **11**, 3696–3713.

Marques,S.M. *et al.* (2017) Enzyme Tunnels and Gates As Relevant Targets in Drug Design. *Med. Res. Rev.*, **37**, 1095–1139.

Morris,G.M. *et al.* (2009) AutoDock4 and AutoDockTools4: Automated docking with selective receptor flexibility. *J. Comput. Chem.*, **30**, 2785–2791.

Pavelka,A. *et al.* (2016) CAVER: Algorithms for Analyzing Dynamics of Tunnels in Macromolecules. *IEEE/ACM Trans. Comput. Biol. Bioinform.*, **13**, 505–517.

Ryckaert,J.-P. *et al.* (1977) Numerical integration of the cartesian equations of motion of a system with constraints: molecular dynamics of n-alkanes. *J. Comput. Phys.*, **23**, 327–341.

Salomon-Ferrer,R. *et al.* (2013) Routine Microsecond Molecular Dynamics Simulations with AMBER on GPUs. 2. Explicit Solvent Particle Mesh Ewald. *J. Chem. Theory Comput.*, **9**, 3878–3888.

Wang,J. *et al.* (2004) Development and testing of a general amber force field. *J. Comput. Chem.*, **25**, 1157–1174.

Zwanzig,R. (1973) Nonlinear generalized Langevin equations. *J. Stat. Phys.*, **9**, 215–220.
